# Conformational Ensembles Reveal the Origins of Serine Protease Catalysis

**DOI:** 10.1101/2024.02.28.582624

**Authors:** Siyuan Du, Rachael C. Kretsch, Jacob Parres-Gold, Elisa Pieri, Vinícius Wilian D. Cruzeiro, Mingning Zhu, Margaux M. Pinney, Filip Yabukarski, Jason P. Schwans, Todd J. Martínez, Daniel Herschlag

## Abstract

Enzymes exist in ensembles of states that encode the energetics underlying their catalysis. Conformational ensembles built from 1231 structures of 17 serine proteases reveal atomic-level changes across their reaction states, identify molecular features that provide catalysis, and quantify their energetic contributions to catalysis. These enzymes precisely position their reactants in destabilized conformers, creating a downhill energetic gradient that selectively favors the motions required for reaction while limiting off-pathway conformational states. A local catalytic motif, the “nucleophilic elbow”, has repeatedly evolved, generating ground state destabilization in 50 proteases and 52 additional enzymes spanning 32 distinct structural folds. Ensemble–function analyses reveal previously unknown catalytic features, provide quantitative models based on simple physical and chemical principles, and identify motifs recurrent in Nature that may inspire enzyme design.

**One sentence summary:** Ensemble–function analyses provide a quantitative model for serine protease catalysis, reveal previously unknown conformational features that contribute to their catalysis, and identify a structural motif that underlie these features and has evolved in >100 different enzymes from 32 protein folds.

## Introduction

Understanding enzyme catalysis has been a central goal of biochemistry. Tremendous efforts in enzymology have led to high-confidence chemical models for the roles of active site residues and cofactors for the vast majority of known enzymes (*1*, *2*). Nevertheless, the long-standing challenge remains to account for the rate enhancements provided by enzymes, as reflected by our limited ability to predict enzyme rates and design efficient enzymes (*3*, *4*).

Over the past century, enzymologists have established foundational concepts that underlie enzyme catalysis (table S1) (*5–15*). Conformational ensembles are needed to connect these concepts to specific molecular mechanisms and to quantify their catalytic contributions. For example, the ability of enzymes to restrict motions and position groups for reaction is universally invoked as a catalytic feature, yet the extent of enzyme positioning and the amount of catalysis arising from the reduced conformational entropy remain elusive. Conformational ensembles define the distribution of states and thus can overcome this deficit. Most fundamentally, the statistics in ensembles encode free energies and thus are needed to quantify catalysis (*16*).

We investigate conformational ensembles of serine proteases, the classic example in biochemistry textbooks used to illustrate enzyme mechanisms (*17–20*). These descriptions highlight the nucleophilic serine, the oxyanion hole, and the catalytic triad (Fig. 1A), and it is known that mutation of the corresponding residues is deleterious to catalysis (*21*, *22*). However, it is not apparent *how* these catalytic groups furnish the observed ∼10^12^-fold rate enhancement (Fig 1B and table S2). The hydroxyl group of the nucleophilic serine has nearly identical reactivity as water, so it alone does not provide catalytic advantages. The oxyanion hole typically uses amide N–H hydrogen bond donors, also similar in strength as the hydrogen bonds from water (*23*). The histidine of the catalytic triad facilitates reaction by acting as a general base, but not nearly enough to account for the rate enhancement achieved by serine proteases (*24*, *25*). Several proposals for additional catalysis from the catalytic aspartate have not been supported by subsequent experiments, and its contribution remains to be determined (supplementary text S1).

**Fig. 1.**
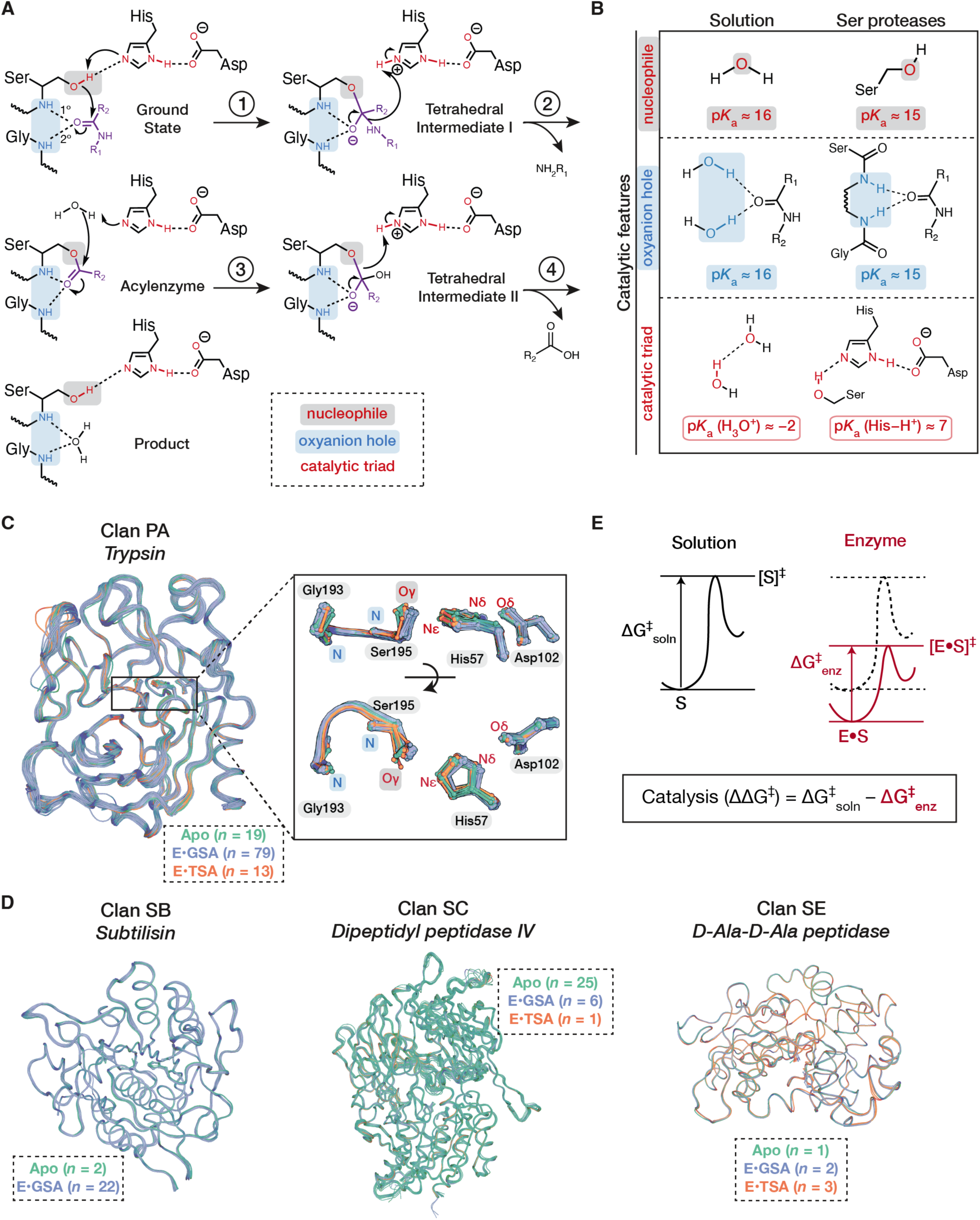
Serine proteases. **(A)** The serine protease reaction mechanism and catalytic groups. The atoms involved in catalytic triad hydrogen bonds are colored red, with the nucleophile shaded in gray; the oxyanion hole hydrogen bond donor atoms are colored and shaded light blue. The 1° hydrogen bond donor is the backbone amide neighboring nucleophilic group (*30*). The 2° hydrogen bond donor is a glycine backbone amide for trypsin and other clan PA serine proteases but varies across structural clans (table S3). Atoms from the substrate amide are purple. **(B)** Chemical features of the enzyme catalytic groups compared to analogous interactions in the solution reaction. p*K*_a_ values are used as proxies for electron density (*31–33*). Intrinsic p*K*_a_ values of water, hydronium ion, acetamide (oxyanion hole donors), methanol (serine sidechain) and imidazole (histidine sidechain) were taken from (*34*). **(C)** Pseudo-ensemble of trypsin (from clan PA), with the number of structures in the apo, GSA-bound and TSA-bound sub-ensembles annotated in the box. Structures were aligned on Cα atoms using the highest resolution structure as the reference (table S4). Labeled atom names follow the PDB format (*35*). **(D)** Pseudo-ensembles of serine proteases across three additional structural clans: subtilisin (clan SB), dipeptidyl peptidase (clan SC) and D-Ala-D-Ala peptidase (clan SE), with the number of structures in their apo, GSA-bound and TSA-bound sub-ensembles shown in the boxes. Figs. S1 to 4 show additional serine proteases in each clan and their active site ensembles. **(E)** Definition of enzyme catalysis. Free energy profiles for the solution reaction (black) and the enzymatic reaction (red) are shown. S and E•S represent the ground states and [S]^‡^ and [E•S]^‡^ represent the transition states of the solution and enzymatic reaction, respectively. The black and red arrows indicate the reaction free energy barriers in solution (ΔG^‡^_soln_) and on the enzyme (ΔG^‡^_enz_). The amount of catalysis is defined as the difference between the solution and enzyme reaction barriers (ΔΔG^‡^).

We brought together >1000 X-ray crystallographic structures of serine proteases to build “pseudo-ensembles” (*26–28*) and we asked the following questions: (1) *What are the molecular changes between reaction states?* (2) *How does the enzymatic reaction differ from the uncatalyzed reaction in solution?* and (3) *What are the energetic consequences of these differences?* Pseudo-ensembles in apo, ground state analog (GSA) and transition state analog (TSA) bound states revealed an altered energy landscape of the enzymatic reaction where shorter and more efficient reaction paths are favored compared to those in solution. Catalytic contributions from these and additional features provide a minimal, quantitative model that accounts for serine protease catalysis. Further, the catalytic conformational features are enabled by a local structural motif referred to as the “nucleophilic elbow” (*29*), and it has evolved in over 100 different enzymes across 32 distinct structural folds.

## Results and discussion

### Crystallographic data provide ensemble information for serine proteases across reaction states

To obtain ensemble information for serine proteases, we leveraged the vast structural data available in the Protein Data Bank (PDB) to build “pseudo-ensembles” (*26–28*, *36*, *37*). These ensembles are built by bringing together structures of the same proteins so that they capture different states around the local minima of the conformational landscape. Pseudo-ensembles are in excellent agreement with nuclear magnetic resonance (NMR) order parameters (*27*) and residual dipolar coupling (*38*, *39*), and with multi-conformer models from room temperature X-ray diffraction (*28*), supporting their ability to capture the low energy states present in crystalline and aqueous states. In particular, pseudo-ensembles provide three-dimensional atomic positions not accessible via NMR dynamics experiments and thus allow detailed interrogation of ensemble–function relationships. The extensive X-ray crystallographic data in the PDB further provided instantaneous access to ensemble information for multiple enzymes.

To build pseudo-ensembles and to compare serine proteases evolved within and across structural folds, we collected 1231 wild-type structures of 17 serine proteases across four structural superfamilies (also referred to as “clans”) from the PDB (Fig. 1C to D, figs. S1 to 4, and table S3). To examine serine protease ensembles across reaction states, we identified structures that are in the apo, GSA-bound, and TSA-bound states (fig. S5 and table S4). GSAs consist of non-covalent peptide (or peptide analog) inhibitors that bind in the same fashion as cognate substrates (supplementary text S2). TSAs are covalent inhibitors such as fluoromethyl ketones (*40*), boronic acids (*41–43*) and peptidyl aldehydes (*44*) that form a tetrahedral intermediate with the catalytic serine (Fig. 1A) but do not react further. Tetrahedral intermediates resemble the transition state geometry in our QM calculation for the solution reaction (fig. S6 to 7 and table S5). A late transition state resembling the tetrahedral intermediate is also found in prior quantum mechanics/molecular mechanics (QM/MM) calculations of serine proteases (*45*), supporting the use of TSAs as mimics for the transition state. Analyses below using a range of bound GSAs and TSAs for multiple serine proteases and other enzymes revealed consistent conformational properties, suggesting that differences between individual analogs or crystal forms did not significantly distort the ensembles or the conclusions obtained. We first focused our analyses on clan PA serine proteases, which include trypsin, the serine protease with the greatest number of available structures (*n* = 445; Fig. 1C and fig. S1). Additional serine proteases evolved within different structural clans (SB, SE and SC; Fig. 1D, figs. S2 to 4) provided tests for whether the identified conformational features have commonly emerged due to convergent evolution.

### Computational and crystallographic data provide information for the uncatalyzed solution reaction

Enzyme catalysis is quantified by the reduction in the free energy barrier relative to the corresponding uncatalyzed reaction (Fig. 1E). We used quantum mechanical calculations to obtain the reaction path for the analogous solution reaction, where water attacks the amide carbonyl of a substrate analog, *N*-methylacetamide (NMA) to form a tetrahedral intermediate. To sample the ensemble distribution of reacting molecules in solution, we conducted molecular dynamics (MD) simulations of NMA in water. We also used a knowledge-based approach in which we assembled crystallographic data of small molecule interactions between amide and hydroxyl groups that resemble the reacting groups in the solution reaction (*Methods* and table S6). Knowledge-based distributions have been widely used to determine conformational preferences and derive energy functions (*46–50*). Crystallographic distributions quantitatively match those from NMR experiments and QM calculations, supporting their ability to approximate ensembles of molecular interactions and determine conformational preferences (*51–55*) (supplementary text 3).

### Serine proteases precisely position reactants in the reactive orientation

To quantify the extent of positioning in serine proteases *versus* in solution, we compared the distributions of reactant positions in their ground state ensembles. While it is easy to appreciate the catalytic advantage of bringing together substrates that are present at low concentrations in cells (μM-mM), the advantage, if any, for hydrolysis reactions is less clear, since water (55 M) is always nearby.

The position of the serine nucleophile relative to the substrate amide carbonyl is defined by three parameters, a distance (*d*_attack_), an angle (*α*_attack_) and a dihedral (ϕ_attack_) (Fig. 2A). The observed distance and angles are narrowly distributed and highly similar for serine proteases across the four structural clans, with *d*_attack_ = 2.68 (*s.d.* = 0.14 Å), *α*_attack_ = 93° (*s.d.* = 7°) and ϕ_attack_ = 84° (*s.d.* = 8°) (Fig. 2B, fig. S8 and table S7). The same parameters in solution, measured using the knowledge-based approach and MD simulations, were much broader (Fig. 2B and fig. S8). In addition, the distances found in enzymes are shorter than those in solution, a feature common to serine proteases across the four distinct structural clans. We return to this observation in later sections.

**Fig. 2.**
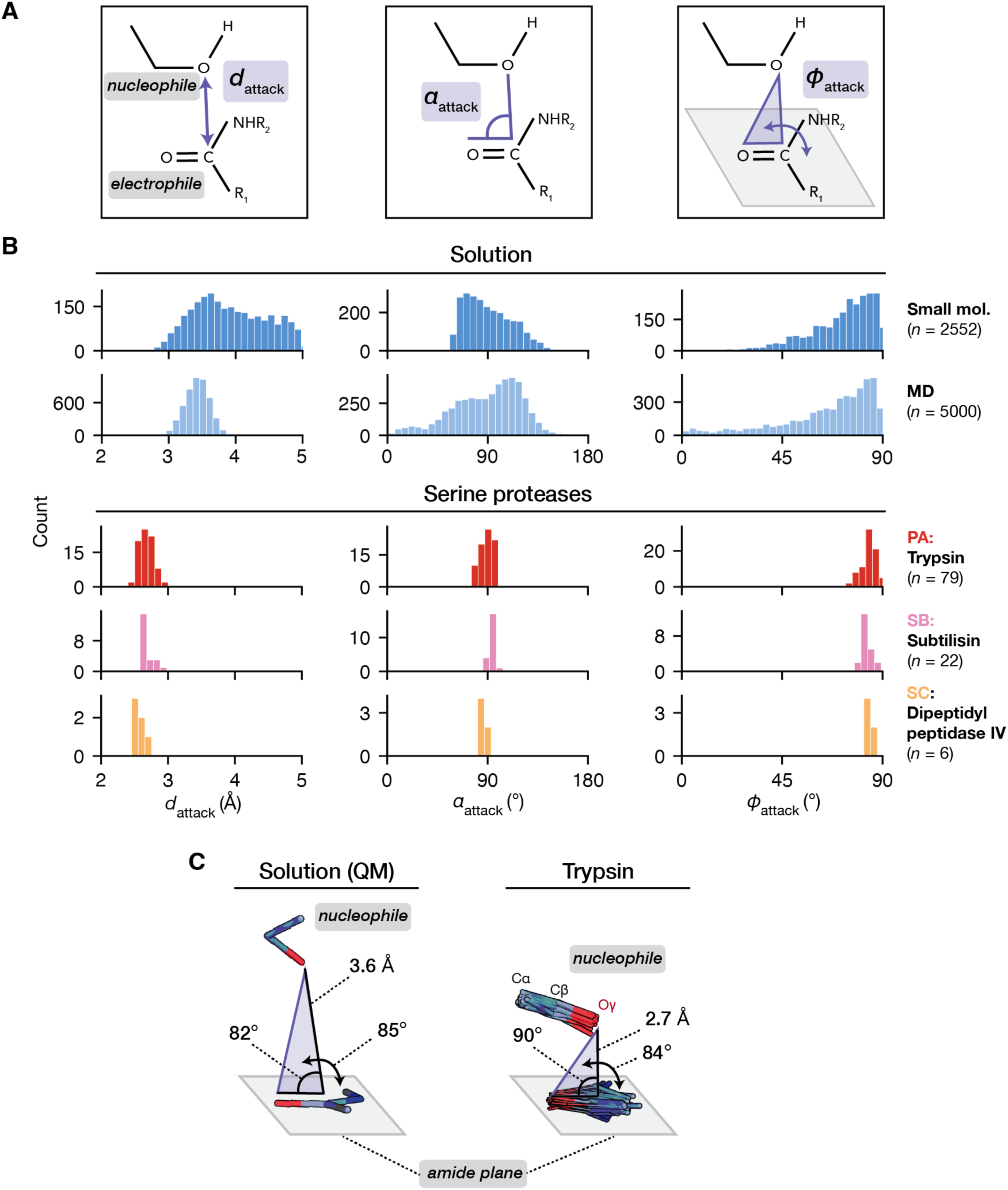
Positioning of the reacting groups in serine proteases. **(A)** The nucleophilic attack geometry is described by three parameters: the attack distance (*d*_attack_), defined by the distance between the nucleophilic and the electrophilic atom; the attack angle (α_attack_), defined by the angle between the nucleophilic atom, the electrophilic atom and the oxygen on the reactive carbonyl group; and ϕ_attack_, defined by the dihedral angle between two planes: (1) the plane defined by the nucleophilic atom, the electrophilic atom, and the oxygen on the amide carbonyl group (purple) and (2) the plane of the reactive amide carbonyl (gray). **(B)** Distributions of attack geometries in serine protease GSA-bound pseudo-ensembles *versus* in the reference ensembles for the solution reaction. The two solution distributions come from small molecule (“small mol.”) hydroxyl•amide interactions or MD simulations of *N*-methyl-acetamide (NMA) in water (“MD”, 5000 snapshots from 100 ns simulation). The differences in the tails of the MD and small molecule distributions arise from choosing only the water closest to the electrophile in each MD frame, which removes the more distal water molecules (see *Methods*). The data for one protease from each structural clan are shown: trypsin for clan PA (red), subtilisin for clan SB (pink), and dipeptidyl peptidase for clan SC (yellow). Additional proteases within each clan and MD replicates are shown in fig. S8. **(C)** Ground state conformation of the reactants from QM calculation of the solution reaction compared to those observed in the trypsin pseudo-ensemble (*n* = 79). The trypsin pseudo-ensemble is locally aligned on the sidechain atoms of the catalytic triad. Ground state ensembles of additional proteases are shown in fig. S9. In QM, ethanol is the nucleophile and NMA is the electrophile; the reaction ground state was found by searching for an energy minimum that is the closest to the tetrahedral intermediate state on the energy landscape.

For the restricted positioning to aid catalysis, two additional criteria need to be met: (1) the nucleophile and the substrate must be positioned in reactive orientations and (2) motions needed for bond formation must be allowed. As described below and in the following section, both criteria are met.

The enzyme angles *α*_attack_ (93°) and ϕ_attack_ (84°) agree with the preferred attack angles determined by Burgi and Dunitz from their analysis of small molecule O•C=O interactions (*α*_attack_ ≈ 100 ± 20°, ϕ_attack_ ≈ 90°) (*56*). The parameters obtained from our QM calculations for the solution ground state fall within the same ranges (*α*_attack_ = 82°, ϕ_attack_ = 84°; Fig. 2C). The attack distance in the QM ground state (3.62 Å) also agrees with the preferred distances observed in MD (3.44 Å, *s.d.* = 0.18 Å) and small molecule distributions (3.61 Å, *s.d.* = 0.54 Å) (table S7). In contrast, the enzymatic *d*_attack_ is ∼0.9 Å shorter (Fig. 2C and table S7), suggesting that the enzyme begins its reaction farther along the reaction coordinate.

### Serine proteases provide shortened, high-efficiency reaction paths

Enzyme positioning can increase the reaction rate by aligning groups in proximity so that the frequency of their reaction is higher than that in solution. However, over-positioning can limit reactant mobility and restrict the motions needed for reaction. Hammes *et al.* articulated this point: “For catalysis, flexible but not too flexible, as well as rigid but not too rigid, is essential.” (*57*). Ensemble–function analysis allowed us to transform this general axiom into atomic-level models for the magnitude, direction, and energetics of the motions involved in an enzymatic reaction.

#### A sidechain rotation of the catalytic serine brings the reacting atoms closer

To carry out an unbiased search for motions that may be part of the serine protease reaction path, we measured the backbone and sidechain torsion angles of each residue in the GSA- and TSA-bound states and compared the distributions between these states (supplementary text S4). We also compared the apo and GSA-bound enzyme ensembles to determine the changes associated with substrate binding. From these comparisons, we derived a minimal model for the motions along the enzymatic reaction path.

Among all buried residues in trypsin, there are 428 torsion angles, 32 of which have significantly different distributions upon substrate binding (apo *versus* GSA-bound state; *p* < 0.05 after multiple hypothesis correction), and 23 of which significantly change upon nucleophilic attack (GSA-*versus* TSA-bound state) (Fig. 3A and fig. S10). The residues identified are spread throughout the structure rather than contiguous, suggesting no large-scale or extensively coupled conformational changes in the serine protease reaction.

**Fig. 3.**
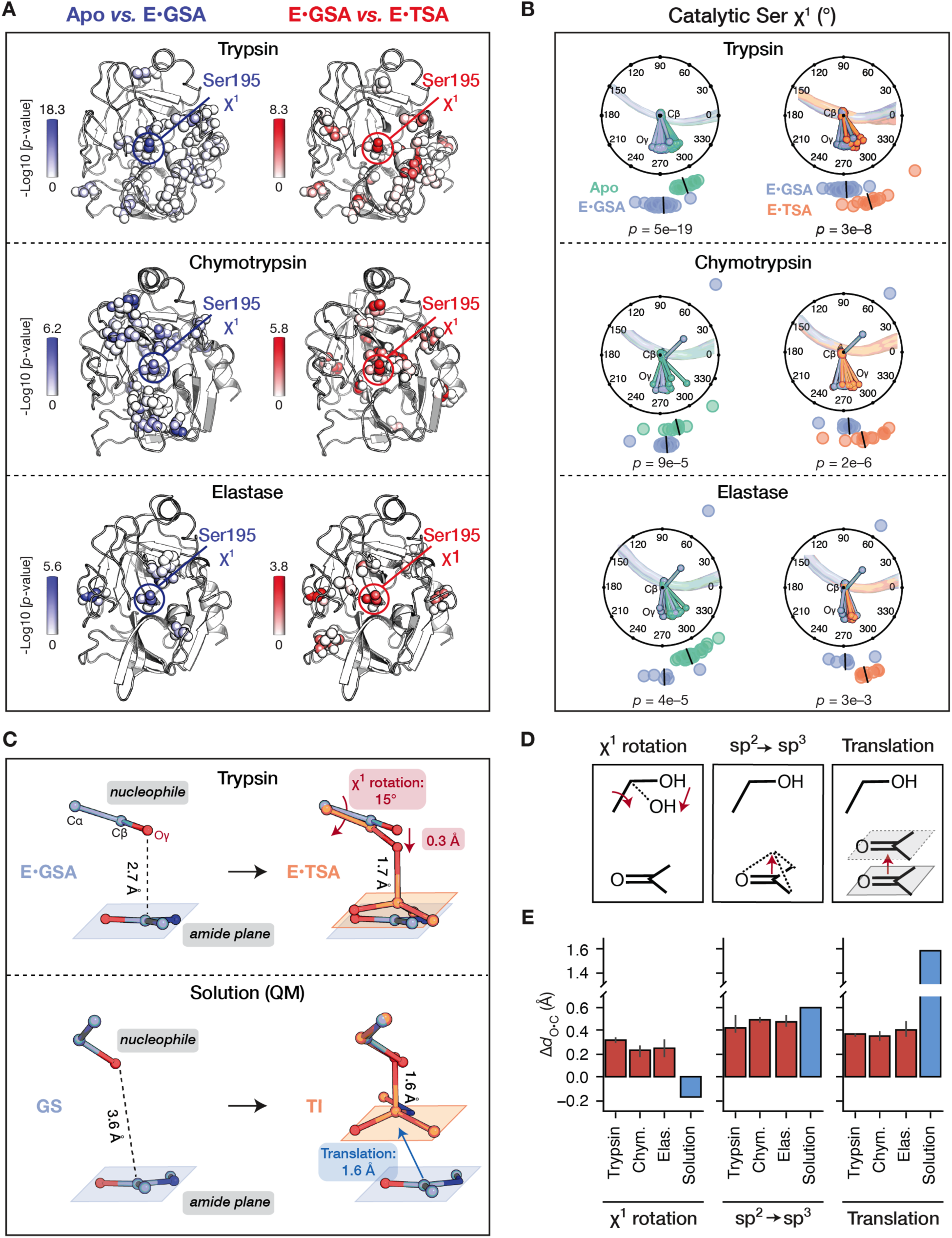
The enzymatic and solution reaction path. **(A)** Trypsin, chymotrypsin and elastase structures colored by the significance [-Log10(*p*-value)] of the differences in torsion angles between apo and GSA-bound (blue) and between GSA- and TSA-bound pseudo-ensembles (red), as determined by the Kolmogorov–Smirnov (K–S) test (*58*, *59*). Torsion angles with *p-*values <0.05 (after multiple hypothesis correction) are shown as spheres and colored; the color scales differ for each serine protease due to the differences in the *p*-value range between the serine proteases. **(B)** Catalytic serine (Ser195) χ^1^ angle distributions in serine protease pseudo-ensembles (apo: aqua; GSA-bound: blue; TSA-bound: orange), with black lines indicating medians. *P*-values are determined by the K–S test; additional tests are shown in fig. S11. Ser195 sidechains are shown in Newman projections along the Cα–Cý axis, aligned locally on the Ser195 backbone N, C, O and Ca and Cβ atoms. **(C)** Geometric changes of the reacting groups that accompany bond formation in the trypsin and solution reactions. For trypsin, only the center structure from each pseudo-ensemble is shown (E•GSA: 3M7Q, E•TSA: 1TPP) for clearer visualization. (The center structures were found by selecting the structure with the median Ser195 χ^1^ value in the ensembles and were locally aligned on sidechain atoms of the catalytic triad. See fig. S17 for full pseudo-ensembles.) The blue and orange plane indicate the amide “plane” (defined by the three substituents of the electrophile) in the GSA- and TSA-bound states. The red arrows indicate the rotation of the catalytic serine sidechain and the resulting distance change. For the QM-calculated solution reaction, ground state (GS) and tetrahedral intermediate (TI) were aligned on the ethanol, imidazole and acetate atoms. **(D)** Three types of motions that shorten the distance between reacting atoms to allow bond formation: the catalytic serine χ^1^ rotation, the sp^2^ to sp^3^ geometric transition of the substrate (sp^2^ → sp^3^), and the translational motion between reacting groups. For the solution reaction, χ^1^ rotation is defined as the H–C–C– OH torsion angle change of the ethanol. **(E)** Contribution from each motion to the nucleophile•electrophile distance shortening (Δ*d*_O•C_) calculated from trypsin, chymotrypsin and elastase pseudo-ensembles (red) and from the QM-determined solution reaction (blue). Positive values indicate that the motion brings reacting atoms closer; negative values indicate the opposite. Bar heights indicate median values and error bars indicate 95% confidence interval for the enzymes (red).

Intriguingly, the sidechain rotation of the catalytic serine (Ser195 χ^1^; rotation along the sidechain Cα–Cý axis) showed highly significant changes in both the substrate binding and the nucleophilic attack step (Fig. 3A and fig. S10). Upon substrate binding, the sidechain of the catalytic serine rotates by −14°, and it rotates back by the same amount (+14°) in going from the GSA-bound to the TSA-bound state (Fig. 3B and table S8). The Ser195 χ^1^ change is significant across a range of different TSAs (fig. S11), and the direction and magnitude of this change is consistent across subsets of structures with varying resolutions, model accuracy (R_free_), and for crystals with varying solvent contents and in different space groups (fig. S12 to 13).

The same analysis carried out for chymotrypsin and elastase, two serine proteases in the same clan as trypsin (PA), also revealed significant changes of the catalytic serine χ^1^ (Fig. 3A and fig. S10 to 12). This change is the only highly significant torsion change common to all three proteases in both the binding and chemical step (Fig. 3A and fig. S14 to 15), and the direction and magnitude of the rotation were also common (Fig. 3B and table S8).

#### Trypsin favors shortened reaction paths and uses a sidechain rotation of the catalytic serine to replace translational motion

The serine χ^1^ rotation from the GSA-to the TSA-bound state brings the nucleophilic oxygen closer to the electrophilic carbon by ∼0.3 Å, consistent with a motion along the reaction coordinate (Fig. 3C and fig. S16 to 18). The distance between the serine nucleophile and the substrate amide carbon progresses from ∼2.7 Å in the ground state to form the ∼1.6 Å covalent bond (table S9), requiring a total of ∼1.1 Å distance change.

The rest of the distance shortening must derive from geometric changes on the substrate and relative translational motion between the reacting groups. (Fig. 3D to E). The substrate amide “pyramidalization” from a sp^2^ carbonyl to a sp^3^ tetrahedral intermediate contributes 0.42 ± 0.09 Å to bond formation in trypsin (Fig. 3E and table S10), measured by converting the pyramidal TSA-bound structures to mimic their planar unreacted state. This amount of change agrees with the prediction from an empirical relationship derived by Burgi *et al.* (*56*). Translational motion contributes 0.36 ± 0.22 Å, determined by aligning pairs of GSA-bound and TSA-bound trypsin structures locally at the active site (Fig. 3E and table S10). The same types and extents of motions were found for other clan PA serine proteases (Fig. 3E, fig. S16 to 18 and table S10), providing additional evidence for a catalytic role and suggesting conservation of the core dynamic features of the enzymatic reaction path. Together, these three features account for the full ∼1.1 Å distance change.

For the solution reaction path, QM revealed a longer distance traveled by the reacting atoms (2.0 Å instead of 1.1 Å), as the reactants begin farther apart in the ground state yet form an essentially identical bond (Fig. 3C). The solution path does not involve bond rotation and is instead dominated by a 1.6 Å translational motion (80% of the total distance; Fig. 3C and E). Thus, the enzymes shorten their reaction paths and replace a portion of the remaining translational motion with a bond rotation along a single dimension that brings the reactants closer. As expected for a covalent change, substrate pyramidalization contributes the same amount in the enzymatic and solution reactions.

The differences identified suggest an altered energy landscape on the enzyme where shorter and more efficient reaction paths are favored, while the more diffuse translational motion that occurs in solution is suppressed (see also Fig. 7). Nevertheless, as the differences on the enzyme are derived from comparisons between ground state and transition state ensembles, we cannot determine the specific paths traveled or the rates of transitions between sub-states.

#### Common evolution of shortened enzymatic reaction paths via serine rotation across different structural contexts

We next turned to serine proteases evolved from four distinct structural clans (where there is sufficient structural data) to ask whether the structural convergence of their catalytic groups was accompanied by the convergence of their ensemble features and reaction paths. Torsion analysis for clans SB, SE, and SC proteases revealed changes along the reaction cycle analogous to those observed for clan PA (Fig. 4A, figs. S19 to 21 and tables S8).

**Fig. 4.**
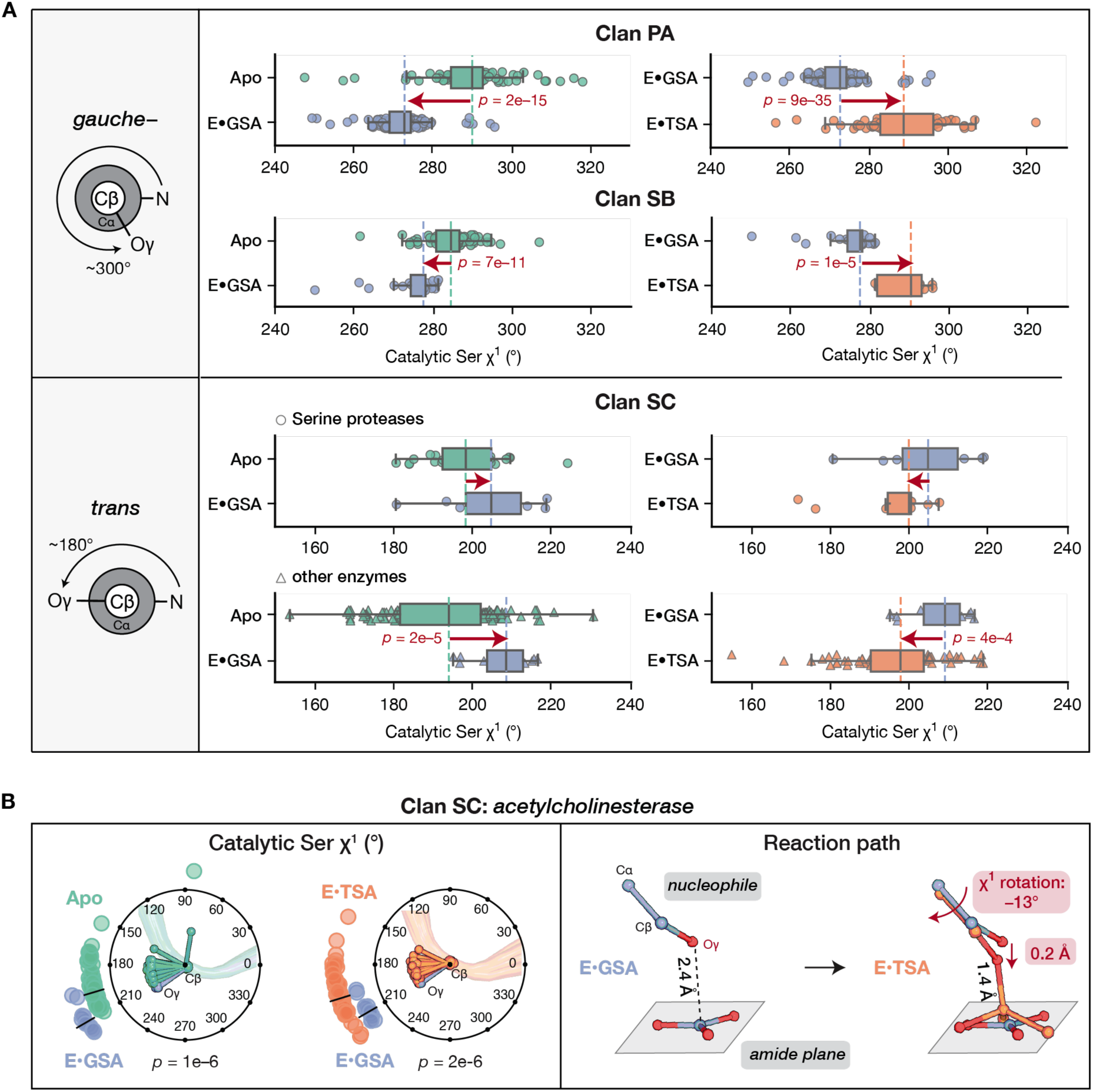
Common reaction path features across serine proteases and non-protease enzymes from multiple structural clans. **(A)** Torsion changes of the catalytic serine χ^1^ across reaction states from serine proteases and additional enzymes from distinct structural clans. The reactive rotameric state for each clan is shown (*gauche*– for clan PA and SB; *trans* for clan SC). Dashed lines indicate the median value of each distribution (apo: aqua; GSA-bound: blue; TSA-bound: orange). The full distributions including rare outliers in unreactive rotamer states are shown in fig. S19, and these outliers were included in *p*-value calculations. Arrows indicate the direction of change; *p*-values from the K–S tests are given for statistically significant changes (*p <* 0.05). Table S8 lists the individual enzymes included in each distribution. **(B)** Geometric changes of acetylcholinesterase across reaction states. Left: the catalytic serine χ^1^ angle distribution in the apo, GSA-bound and TSA-bound states; right: alignment of the center structures from each pseudo-ensemble shown (E•GSA: 2GYW, E•TSA:2HA0). Colors and symbols are as in Fig. 3B to C. Fig. S21 shows the full acetylcholinesterase pseudo-ensembles.

Clan SC, also known as the α/β hydrolase fold (*60*), contains enzymes with a catalytic triad and oxyanion hole like all serine proteases but uses a different oxyanion hole hydrogen bond donor that requires the serine to sit in a different rotameric state (*trans vs. gauche–,* as defined in Fig. 4A) (*30*). The clan SC catalytic serines undergo a rotation of the same magnitude as observed for other clans, but in the opposite direction (Fig. 4A to B and fig. S22). This opposite rotation achieves the same mechanistic outcome, bringing the nucleophilic oxygen closer to the electrophilic carbon for bond formation and thereby providing strong support for a common catalytic role of the rotation across multiple structural scaffolds.

The same reaction path was also found for non-protease members of clan SC that carry out nucleophilic attack on carbonyl centers. Acetylcholinesterase, a clan SC member whose many X-ray structures allowed us to evaluate its catalytic cycle, initiates its reaction from a short 2.42 Å (*s.d.* = 0.16 Å) distance in the ground state and undergoes the same ∼15° serine χ^1^ rotation (Fig. 4B). Data from additional clan SC non-protease enzymes, when combined, also revealed short attack distances (fig. S23) and the serine rotation (Fig. 4A).

Overall, the conservation and convergence of the enzymatic reaction paths strongly suggests common catalytic strategies that are provided by common conformational features. In the next sections, we quantitatively assessed their catalytic contributions.

### A minimal, quantitative model for serine protease catalysis

The functional groups in serine protease active sites have chemical properties similar to water, with the exception of the His general base that abstracts a proton from serine to increase its reactivity and the catalytic triad His•Asp hydrogen bond that is absent in the solution reaction (Fig. 1B). Linear free energy relationships (LFERs) are used to estimate the catalytic contributions from these two chemical features. The general base histidine and its positioning provides 8.2 ± 3.1 kcal/mol catalysis (“±” indicates upper and lower bounds), based on estimated Brønsted relationships and effective molarities of intramolecular general base catalysis (*61–64*), corresponding to a ∼10^6^-fold rate enhancement (supplementary text S5, fig. S24 and table S11). The His•Asp hydrogen bond strengthens in going from the ground to the transition state as positive charge accumulates on the histidine, providing an estimated 0.8 kcal/mol catalytic contribution (see *Catalysis from the catalytic triad hydrogen bonds*). These values emphasize the substantial catalysis possible by introducing chemical groups that stabilize the transition state more than water. Nevertheless, these features are not nearly sufficient to account for the total reduction in reaction barrier provided by serine proteases (9.0 kcal/mol predicted *vs*. 17.1 kcal/mol observed [table S2]), suggesting the presence of additional catalytic features.

We determined the energetic consequences from each conformational features of the enzymatic and solution reaction paths identified above. Most fundamentally, features that provide catalysis reduce the free energy difference between the ground and the transition state on the enzyme compared to that in solution (Fig. 1E). Our energetic analyses identified catalytic contributions arising from features of the enzymatic ground state––reduced conformational entropy and unfavorable interactions––that are absent in solution; these destabilizations are then relieved in the transition state. As such, we refer to these mechanisms as “ground state destabilization”. As exemplified by trypsin below, the sum of these mechanisms provides an additional 7.4 kcal/mol catalysis (Fig. 5A), leading to a minimal model that accounts for the total rate enhancement, within error.

**Fig. 5.**
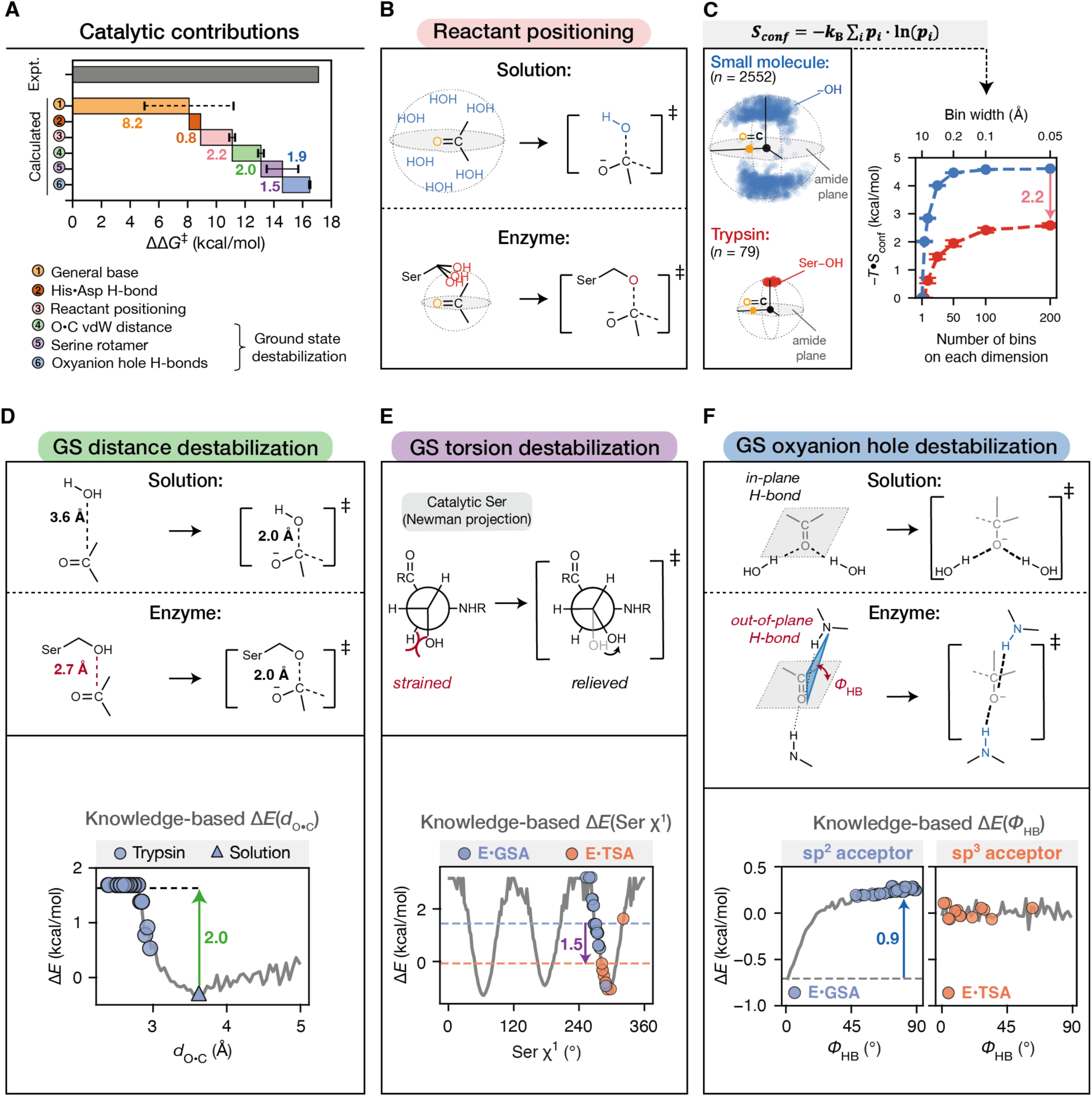
Catalytic features and quantitative estimates for their energetic contributions to trypsin catalysis. (See fig. S27 to 28 and S33 for additional serine proteases.) **(A)** The sum of the energetic contributions from the identified catalytic feature accounts for trypsin’s total rate enhancement from trypsin within error [17.1 kcal/mol from experiment (gray bar, table S2) *versus* 16.6 kcal/mol calculated (colored bars, with colors corresponding to each catalytic feature)]. The error bar for general base catalysis (dashed line) indicates lower and upper ranges; all other error bars (solid lines) indicate standard deviations determined as described in *Methods*. **(B)** Positioning of reactants on the enzyme compared to that in solution provides an entropic advantage for achieving the highly restricted transition state. **(C)** Catalytic contribution from enzyme positioning. Left: spherical distributions of nucleophile positions relative to the substrate amide group observed in the ensembles, with the electrophilic carbon placed at the origin and the substrate amide plane defining the *xy* plane. The colored circles represent positions that the nucleophile occupies (red: trypsin GSA-bound pseudo-ensemble; blue: small molecule hydroxyl oxygens that are in proximity to amide groups). Top: the formalism of Gibbs entropy used for the calculation of *S*_conf_ (equation in the gray box, where *k*_B_ is the Boltzmann constant and *p*_i_ is the probability of observing the system at a microstate *i*, and at *T* = 298 K). Right: conformational entropies (–*T•S*_conf_) calculated from the distributions using a range of bin number from 1^3^ to 200^3^, corresponding to bin widths from 10 Å to 0.05 Å or microstate volumes from (10 Å)^3^ to (0.05 Å)^3^. Error bars indicate standard deviations from bootstrapping (table S12). **(D)** Catalytic contribution from ground state destabilization of the short O•C van der Waals interaction, a feature that is not present in the solution reaction. Knowledge-based energy function for the O•C van der Waals distance (*d*_O•C_) built from 2,552 small molecule structures of amide•hydroxyl interactions (Fig. 2B) is shown as the gray solid line. The distances observed in the trypsin GSA-bound pseudo-ensemble (blue circles) and the solution ground state distance from our QM calculation (blue triangle) were mapped on the energy function. The black dashed line indicates the mean of calculated energies from the trypsin GSA-bound ensemble. The green arrow and number indicate the difference between the energies calculated for trypsin and for solution (from QM). **(E)** Catalytic contribution from ground state destabilization of the partially eclipsed torsion angle of the catalytic serine that is relieved in the transition state. Knowledge-based energy function for serine χ^1^ built from 41,237 serine residues from the BBDEP2010 database (*67*) is shown as the gray solid line. Catalytic serine χ^1^ values from the trypsin pseudo-ensembles were mapped on the energy function (GSA-bound: blue; TSA-bound: orange). Dashed blue and orange lines indicate the mean of the calculated energies for each pseudo-ensemble. The purple arrow and value indicate the difference between the means of the GSA-bound and TSA-bound energies. **(F)** Catalytic contribution from ground state destabilization of the 1° oxyanion hole hydrogen bond. Solution amide•carbonyl hydrogen bonds prefer an in-plane orientation whereas the enzymatic ground state is destabilized with out-of-plane oxyanion hole hydrogen bonds. The planarity of a hydrogen bond is quantified by ϕ_HB_ (red), the dihedral angle between the two planes: (1) the plane defined by the donor (N), the acceptor (carbonyl O) and the carbonyl C (blue) and (2) the carbonyl plane (gray). Knowledge-based energy functions of ϕ_HB_ built from 60,018 small molecule amide•carbonyl hydrogen bonds (sp^2^ acceptor) and from 5,823 hydrogen bonds between amide and sp^3^ oxygens (sp^3^ acceptor) are shown as gray solid lines. The ϕ_HB_ values from trypsin pseudo-ensembles for the 1° oxyanion hole hydrogen bonds were mapped on the energy function (GSA-bound: blue; TSA-bound: orange). See fig. S33 for the 2° hydrogen bonds. The blue arrow represents the amount of destabilization in the enzymatic ground state.

#### Catalysis from the positioning of reactants

The greater positioning of reacting groups on an enzyme compared to those in solution can increase the probability of reaching the highly restricted transition state and thereby contribute to catalysis (Fig. 5B). To quantify the difference in positioning, we considered the relative nucleophile•amide positions in three-dimensional space and calculated the conformational entropy (*S*_conf_) associated with the distributions, using the formalism of Gibbs entropy (*65*, *66*) (Fig. 5C). We assume that the enzymatic and solution transition states have the same *S*_conf_ that are similar and both less than or equal to the ground state values.

The trypsin and the “solution” distribution from small molecules differed in *T*ᐧ*S*_conf_ by 2.2 (*s.d.* = 0.2) kcal/mol (*T* = 298 K; Fig. 5B to C and table S12). The same difference was found between the trypsin and the solution distribution from MD (fig. S25). Bootstrap resampling and microstate binning controls indicated that the data provided consistent entropy values (fig. S25 and table S12). The 2.2 kcal/mol difference in *T*ᐧ*S*_conf_ between the enzyme and solution ground state corresponds to a ∼40-fold rate enhancement (Fig. 5A, pink), a modest catalytic contribution from enzyme positioning even relative to 55 M water.

#### Catalysis from distance and torsion destabilization in the enzymatic ground state

The nucleophile•electrophile distance on serine proteases is considerably shorter than that in solution (2.7 Å *vs.* 3.6 Å). To quantify the energetic difference, we constructed a knowledge-based energy function from the distance distribution of small molecule hydroxy•amide interactions, using Boltzmann statistics to convert the observed distance distribution into relative energies (Fig. 5D and supplementary text S3). This energy function gave a preferred distance of 3.61 Å, matching the distance found in the solution ground state from MD and QM (Fig. 2B to C and table S7). The shorter nucleophile•electrophile distances in trypsin corresponds to a destabilization of 2.0 (*s.d.* = 0.2) kcal/mol (Fig. 5D and table S13). Bootstrap analyses resampling datasets of smaller sample sizes from the small molecule data generated consistent energy functions that gave the same amount of destabilization, providing confidence for their statistical stability (fig. S26). However, since high energy regions are less sufficiently sampled in crystallographic distributions, the destabilization may be higher so that the catalytic contribution may be a lower limit.

This ground state destabilization is expected to be relieved in the transition state and the high-energy tetrahedral intermediate where partial and full covalent bonds are formed with the same bond lengths as in solution (Fig. 5D). Prior QM/MM calculation of serine proteases (*45*) and our solution QM calculations give the same transition state nucleophile•electrophile distance (2.0 Å); TSA-bound structures also gave the same bond lengths as that of the intermediate state in our QM calculation (1.56 Å *vs.* 1.60 Å; table S9). Relief of the destabilized nucleophile•electrophile distance in the enzymatic ground state is predicted to reduce the reaction barrier by 2.0 kcal/mol, corresponding to a 30-fold rate enhancement (Fig. 5D). Other serine proteases are estimated to provide the same contributions (fig. S27 and table S13).

The energetic consequence of the catalytic serine rotation was determined using an analogous approach. A knowledge-based energy function for serine sidechain χ^1^ angles was built using 41,237 serine residues from high quality PDB structures (*67*), reproducing the well-known preferences for staggered over eclipsed conformations (*68*, *69*) (Fig. 5E and fig. S28). Mapping the catalytic serine χ^1^ from pseudo-ensembles onto this energy function revealed a semi-eclipsed serine sidechain conformation in the ground state that is destabilized by 1.5 (*s.d.* = 1.1) kcal/mol (table S14). In the TSA-bound state, the serine rotates to the favored staggered conformation, thereby relieving the destabilization (Fig. 5E, fig. S28 and table S14). In contrast, no significant bond rotations occur for the nucleophile in the corresponding QM solution reaction (Fig. 3C). Thus, the serine torsion destabilization in the enzymatic ground state relative to the transition state contributes 1.5 kcal/mol to catalysis—a rate enhancement of ∼10-fold (Fig. 5A, purple).

In contrast to Ser195, other sidechain torsion changes from buried trypsin residues yield small energetic consequences (less than ±1 kcal/mol each; fig. S29), suggesting limited catalytic contributions from distal groups.

#### Catalysis from destabilization of the oxyanion hole hydrogen bonds in the enzymatic ground state

Previous studies by Simón and Goodman (*70*, *71*) found that enzyme oxyanion hole hydrogen bonds are generally out-of-plane from the carbonyl groups of the bound ligands and therefore unable to interact with the oxygen lone pairs in optimal orientations (Fig. 5F). Their computations suggested that this destabilization was lessened in the oxyanionic transition state where no lone pair directionality is expected. Our analysis of individual serine proteases with carefully curated GSA-bound structures supports the observation of out-of-plane hydrogen bonds for serine proteases across the four clans (supplementary text S6, figs. S30 to 35 and tables S15 to 16). These proteases have preferred dihedral angles (ϕ_HB_) of 76° (*s.d.* = 10°) and 76° (*s.d.* = 5°) for the two oxyanion hole hydrogen bonds, where ϕ_HB_ = 0° is in-plane and ϕ_HB_ = 90° is orthogonal to the carbonyl plane (Fig. 5F and fig. S30). As expected, knowledge-based distribution for small molecule amide•carbonyl hydrogen bonds donated to the sp^2^ hybridized oxygen prefer in-plane orientations (fig. S30 and table S12). The energy function for ϕ_HB_ derived from this distribution gave a destabilization of 0.94 ± 0.03 and 0.95 ± 0.02 kcal/mol from the suboptimal orientation for the two oxyanion hole hydrogen bonds in trypsin (Fig. 5F, sp^2^).

In contrast, no such destabilization is present for the TSA-bound states, as there is no angular preference for hydrogen bonds between amide–NH and sp^3^-hybridized oxygens (Fig. 5F, sp^3^ and fig. S32 to 33), presumably reflecting the near symmetric electron distributions around sp^3^ oxyanions. Thus, the two destabilized oxyanion hole hydrogen bonds in trypsin provide a catalytic contribution of 1.9 kcal/mol, corresponding to a rate enhancement of 25-fold (Fig. 5A, blue).

The same destabilized oxyanion hole hydrogen bonds were found for GSA-bound serine proteases across four structural clans (supplementary text S6, figs. S31 and 33 and tables S15 and 16C), as was also found for the distance and torsion destabilization described above.

#### Catalysis from the catalytic triad hydrogen bonds

Unlike the oxyanion hole hydrogen bonds, the catalytic triad hydrogen bonds (His•Ser and His•Asp) involve groups that are stronger hydrogen bond donors or acceptors than water, and therefore can provide catalysis based on their inherent properties without geometric destabilization. The catalytic serine and histidine undergo proton transfer concerted with the nucleophilic attack (*72*); thus, any catalytic contribution from the His•Ser hydrogen bond is in principle included as part of the general base catalysis term described above (supplementary text S5 and fig. S24). The His•Asp hydrogen bond strengthens going from the ground to the transition state as the histidine becomes protonated. The catalytic triad Asp is a stronger hydrogen bond acceptor than water, due to the higher electron density on a carboxylate than water oxygen atom. As the increase in hydrogen bond strength depends on the donor and acceptor strength, we expect the His•Asp hydrogen bond strengthens more than a His•water hydrogen bond during the reaction. This difference is estimated to be ∼0.8 kcal/mol (a ∼4-fold rate enhancement) from a linear free energy relationship for hydrogen bond energetics (*73–75*) (supplementary text S7 and fig. S36).

Pseudo-ensemble comparisons suggest little or no additional contribution from geometric features for either hydrogen bonds (supplementary text S8, figs. S37 to 46 and tables S17 to 20). While prior studies emphasized the potential catalytic role of a short His•Asp hydrogen bond (supplementary text S1), the short His•Asp hydrogen bond (2.71 Å, *s.d.* = 0.11 Å) in the TSA-bound state matches the optimal distance predicted from analogous imidazole•carboxylate hydrogen bonds in small molecules (2.69 Å, *s.d.* = 0.14 Å) and the average His•Asp hydrogen bonds in the PDB (2.73 Å, *s.d.* = 0.39 Å) (table S17A), providing no indication for special geometric features that contribute to catalysis.

#### The serine protease catalytic ledger

Our ensemble–function analyses allowed us to assemble an energetic ledger for the serine protease catalysis that sums to 16.6 kcal/mol, within error of the experimentally measured catalysis of 17.1 kcal/mol (table S2) for the acylation step. We expect these contributions to be additive, as each is obtained from a local, individual property (supplementary text S9 and fig S47). Overall, we found extensive catalysis (7.4 kcal/mol) provided by constraining reactants in a destabilized conformer in the ground state and relieving the destabilization in the transition state. Consistent conformational properties found across multiple crystallization conditions and crystal forms of the same enzyme and across many different enzymes provide strong support for the identified catalytic features.

This catalytic ledger is a *minimal* model; given the uncertainties, the possibility of modest catalytic contributions from additional features remains. In particular, the ability to detect catalytic features depends on the quality and amount of structural information available. Our pseudo-ensembles accurately recapitulated NMR chemical shift data, revealing subtle differences in catalytic triad hydrogen bond lengths (∼0.1 Å) between specific types of TSAs (supplementary text S8 and fig. S45). Remote changes identified from pseudo-ensembles yield minimal energetic consequences and are not conserved (fig. S15 and 29), so we do not invoke additional catalytic mechanisms arising from long-range effects [e.g. (*76–80*)]. Nevertheless, we cannot rule out smaller changes that cannot be confidently resolved or multiple small changes that sum to a non-negligible contribution.

This model relies on TSAs to provide adequate information for the transition state. In the case of serine proteases, the TSA-bound structures resemble the transition state geometry found in QM (fig. S6 and table S5), and consistent catalytic features are found regardless of specific TSA types (fig. S11). Limitations of TSAs will need to be evaluated in future ensemble studies of other reaction classes. There are also uncertainties associated with the energy functions used to calculate catalytic contributions; testing and improving these functions remains an important future challenge.

In addition, pseudo-ensembles do not describe barrier-crossing dynamics. The energy landscape and catalytic model are derived based on transition state theory (*81*) and thus do not account for these ultra-fast events [e.g. bond vibrations and hydrogen tunneling (*82–86*)] that might facilitate catalysis. Since our catalytic model accounts for serine protease catalysis within reasonable error (3∼4 kcal/mol), catalytic contributions from these dynamic features are expected to be modest.

The locally destabilized, catalytically productive conformers that we identified must be enforced by additional stabilizing interactions with the substrate and the structural scaffold. In the following sections, we investigate structural elements responsible for the catalytically competent active site conformer. Specifically, we describe a local motif, the “nucleophilic elbow”, that constrains the oxyanion hole hydrogen bond geometry to generate ground state destabilization and has evolved in >100 unique enzymes.

### The nucleophilic elbow underlies oxyanion hole ground state destabilization

A motif connecting the nucleophile and the oxyanion hole hydrogen bond donor, referred to as the “nucleophilic elbow”, was shown to be conserved in α/β hydrolases (*29*, *60*) and also found in additional proteases (*30*, *87*). This motif forms a pseudo-ring structure with the bound substrate carbonyl, via the serine nucleophile and its backbone amide (Fig. 6A, “N-type”) or that of its neighboring residue (“N+1”-type, fig. S48). As ring structures are geometrically constrained, we reasoned that the nucleophilic elbow pseudo-ring might be responsible for the ground state destabilization observed for the 1° oxyanion hole hydrogen bond (Fig. 5F). Such an effect would be analogous to ring strain commonly found in small molecule covalent rings (*88*), but in this case with a mixture of covalent and non-covalent connections and thus likely less severe energetic consequences (Fig. 6B).

**Fig. 6.**
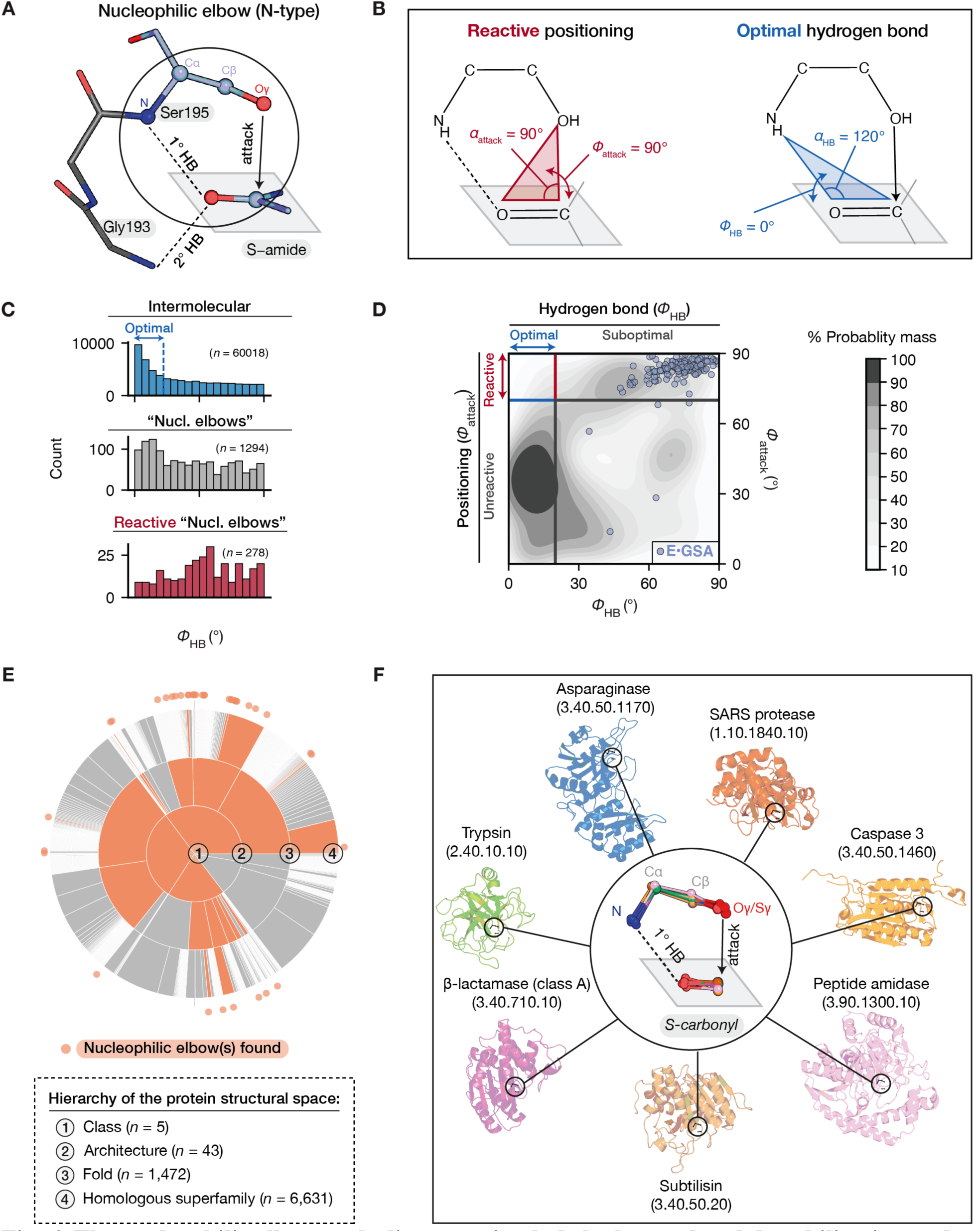
The nucleophilic elbow underlies oxyanion hole hydrogen bond destabilization and has repeatedly evolved. **(A)** The nucleophilic elbow (N-type) in GSA-bound trypsin (PDB: 3M7Q). See fig. S58 for the N+1-type. The “nucleophilic elbow” was first discovered as a conserved sheet-turn-helix motif in α/β hydrolases (*29*, *91*). This motif allows the backbone amide adjacent to the nucleophilic residue to be used as an oxyanion hole hydrogen bond donor (fig. S58). Similar atomic arrangements were later found across many serine and cysteine proteases (*30*, *87*). We use the term “nucleophilic elbow” to refer to this pseudo-ring atomic arrangement connecting the nucleophile and the oxyanion hole wherever it is found, extending the term “nucleophilic elbow” beyond its original usage for α/β hydrolases. **(B)** The reactive configuration (red; *α*_attack_ and ϕ_attack_ defined in Fig. 2A) and optimal hydrogen bond geometry (blue; *α*_HB_, defined as the angle between the hydrogen bond donor N, and the O and C atoms on the acceptor carbonyl group, and ϕ_attack_, defined in Fig. 5F). **(C)** Distributions of ϕ_HB_ in intermolecular amide•carbonyl hydrogen bonds (blue), small molecule N-type nucleophilic-elbow-like structures (gray), and the subset with reactive orientations (defined as ϕ_attack_ ≥ 70°, red). The blue arrow indicates the region of ϕ_HB_ ≤ 20° and is assigned as “optimal” for hydrogen bonding. **(D)** Knowledge-based conformational map for the hydrogen bond orientations (ϕ_HB_) and nucleophile•amide attack orientation (ϕ_attack_) in small molecule N-type nucleophilic-elbow-like motifs (*n* = 1294). The intensity of gray indicates the cumulative probability mass (%) at each contour level. Blue circles indicate the ϕ_HB_ and ϕ_attack_ values observed in GSA-bound serine proteases with N-type nucleophilic elbows. **(E)** Positions of protein structures where nucleophilic elbows are found among the hierarchy of structural classification of the known protein structural space. The structural hierarchy from the CATH database (*92*) was constructed from 536,613 structural domains in the PDB. Levels and the numbers of types of structures at each hierarchy level are annotated in the dashed box. The area of each slice at each level is proportional to the number of its members. The protein structural space at the class, architecture and fold levels is colored by whether the nucleophilic elbow is present (orange) or absent (gray). At the homologous superfamily level, only superfamilies with nucleophilic elbow-bearing enzymes are shown (as orange circles). **(F)** Alignment of the nucleophilic elbow in its GSA-bound state from 15 enzymes (inner circle). These enzymes are from seven structural folds, and the structure of one representative from each fold is shown on the outside. The enzymes shown have N-type nucleophilic elbows and their serine rotamers are in the *gauche–*state. (See fig. S55 for additional alignments for the N-type nucleophilic elbows in the GSA-bound, TSA-bound and acylenzyme states and the same alignments performed for N+1-type nucleophilic elbows.)

To evaluate this possibility, we searched the CSD for small molecule interactions with atomic arrangements that mimic the pseudo-ring structure of the nucleophilic elbow (Fig. 6B and fig. S48). The conformational distribution from 1294 small molecule “nucleophilic elbow” structures (N-type) revealed an overall preference for in-plane hydrogen bonds (ϕ_HB_ = 0°), as found for unconstrained, intermolecular amide•carbonyl hydrogen bonds (Fig. 6C and fig. S49). However, the subset of structures exhibiting the reactive conformation (ϕ_attack_ ≥ 70°) has out-of-plane, destabilized hydrogen bond orientations (Fig. 6C). Stated another way, conformers that simultaneously align reacting atoms for reaction and form a favorable hydrogen bond are low-probability and thus high-energy states (Fig. 6D). Similar constraints were found for N+1-type nucleophilic elbows (fig. S50). As a result of these constraints, the nucleophilic elbow provides a simple and efficient catalytic strategy—aligning the catalytic serine and the substrate for reaction simultaneously destabilizes the oxyanion hole hydrogen bonds, thereby augmenting catalysis.

The other serine protease oxyanion hole hydrogen bond (2°) is also out-of-plane and destabilized (fig. S33). Small molecule data indicates that when the 1° oxyanion hydrogen bond is out-of-plane, an out-of-plane 2° hydrogen bond is also favored (fig. S51), perhaps due to steric hindrance between the hydrogen bond donors. The acylenzyme state also exhibits geometric constraints and oxyanion hole destabilization, suggesting that the nucleophilic elbow also facilitates the deacylation step (fig. S52).

The catalytic conformation of the nucleophilic elbow requires binding interactions to position and constrain the substrate. Intriguingly, in some proteases, binding interactions remote from the cleavage site do not significantly enhance substrate binding but contribute to catalysis (increased *k*_cat_) (*7*, *89*, *90*), presumably by facilitating the ground state destabilization mechanisms identified above. Future ensemble-function studies may reveal the conformational and energetic underpinnings of these and other longer-range functional interconnections.

### Repeated evolution of the nucleophilic elbow

Catalysis from the nucleophilic elbow derives from local interactions. We therefore suspected that this motif could be used within different global structures and may have evolved multiple times. In particular, the analysis above predicts that the nucleophilic elbow can provide catalytic advantages for enzymes that perform nucleophilic attack and stabilize an oxyanionic covalent intermediate via hydrogen bonds. We searched the Mechanism and Catalytic Site Atlas (M-CSA) database (*93*) for enzymatic reactions matching these criteria and found 92 nucleophilic residues from 91 unique enzymes that carry out three types of reactions: nucleophilic addition on carbonyl compounds (*n* = 86), nucleophilic aromatic substitution (*n* = 1), and S_N_2 reaction on phosphoryl compounds (*n* = 5). Nucleophilic elbows were found in 69 of the 91 enzymes, including 50 that perform non-protease reactions (table S2). Combined with the serine and cysteine proteases described herein (*n* = 16; table S3) and in prior work by Buller and Townsend (*n* = 18; table S22) (*30*), the nucleophilic elbow was found in 102 different enzymes (table S23 and fig. S53).

We examined the distribution of the nucleophilic elbow-bearing enzymes across the known protein structural space using the CATH database (*92*). This set of 102 enzymes spans 32 structural folds spread across distinct branches structure space (Fig. 6E and fig. S54). Therefore, it appears that the nucleophilic elbow has repeatedly evolved, with multiple structural origins, presumably a result of the simplicity and the local nature of this motif, combined with its catalytic prowess.

### Nucleophilic elbow-bearing enzymes from multiple folds exploit common catalytic features

The wide presence of the nucleophilic elbow suggests an ability to maintain its catalytic function for multiple reactions when inserted in different structural scaffolds. To assess the generality of the catalytic function provided by the nucleophilic elbows, we turned to the enzymes that perform nucleophilic addition on carbonyl compounds, as they share similar mechanistic constraints. Within the set of 102 nucleophilic elbow-bearing enzymes, there are 66 enzymes (from 22 structural folds) performing this reaction type, and they act on an array of carbonyl compounds beyond peptide amides, including lactams (*n* = 3), amino acid sidechains (Gln or Asn; *n* = 8), esters (*n* = 16), thioesters (*n* = 9), carboxylate (*n* = 1), and aldehydes (*n* = 5) (table S23). The simplest prediction is that nucleophilic elbows provide the same catalytic advantages across these enzymes and reactions.

As nucleophilic elbow catalysis originates from its conformational properties, we assessed the nucleophilic elbow geometries for enzymes with GSA-bound and acylenzyme structures (34 enzymes from 13 structural folds, including 10 non-protease enzymes). Despite the diversity in overall structures, the nucleophilic elbow atoms of these enzymes are well aligned (Fig. 6F and fig. S55), with root-mean-square deviations from 0.16 to 0.23 Å (fig. S56 and table S24). In their GSA-bound states, the nucleophilic elbows from most of the non-protease enzymes achieve the same reactive configuration as observed for serine proteases, with ϕ_attack_ = 84° (*s.d.* = 21°) and *α*_attack_ = 88° (*s.d.* = 11°) and exhibit short, destabilized distances (2.77 Å, *s.d.* = 0.31 Å) (fig. S57 and table S25). In addition, their oxyanion hole hydrogen bonds are out-of-plane from the substrate carbonyl groups and destabilized, with ϕ_HB_ = 84 ° (*s.d.* = 20°) and 83° (*s.d.* = 29°) (fig. S58 and table S25), as they are in the serine proteases. Thus, reactive positioning and ground state destabilization appears to be common to nucleophilic elbow enzymes (supplementary text S10). Finally, and intriguingly, nucleophilic elbows were also found in two phosphoryl transfer enzymes, suggesting broader catalytic capabilities of this motif (fig. S59).

## Conclusions and implications

Our ensemble-function investigation identified catalytic features that were previously unknown and established a minimal, quantitative model for serine protease catalysis. These catalytic features are recurrent in Nature, having commonly evolved in serine proteases from distinct structural clans and in many enzymes catalyzing related reactions. This catalytic model does not invoke new concepts; instead, it is grounded in the most basic concepts of chemistry and physics—torsion angles, van der Waals interactions, hydrogen bonds, and entropy. The simplicity may inspire new ways of teaching enzyme catalysis, allowing instructors to reinforce the value of fundamental physical and chemical concepts and students to appreciate the tight connection between these concepts and the emergent, complex functions of biomolecules.

One of the most vexing questions in understanding enzymes has been the dichotomy between the potential catalysis from positioning and the motions that are needed for reactions to occur. Fig. 7 illustrates this paradox and how serine proteases have arrived at an elegant solution. An overly positioned enzyme (Enzyme 1) would hinder rather than facilitate reaction because the ground state is restricted to a deep energy minimum, rendering excursions to the transition state less likely than in solution. Uniformly increasing flexibility around this state allows the reaction to proceed but also increases the probability of occupying off-pathway conformations (Fig. 7, Enzyme 2). Serine proteases overcome this inefficiency, at least in part, by restricting motions to a sidechain rotation that aligns with the direction of bond formation (Fig. 7, Enzyme 3). This rotation replaces a portion of the translational motion that dominates the solution reaction, limiting the off-pathway conformational space and thereby increasing the probability of reaction.

**Fig. 7.**
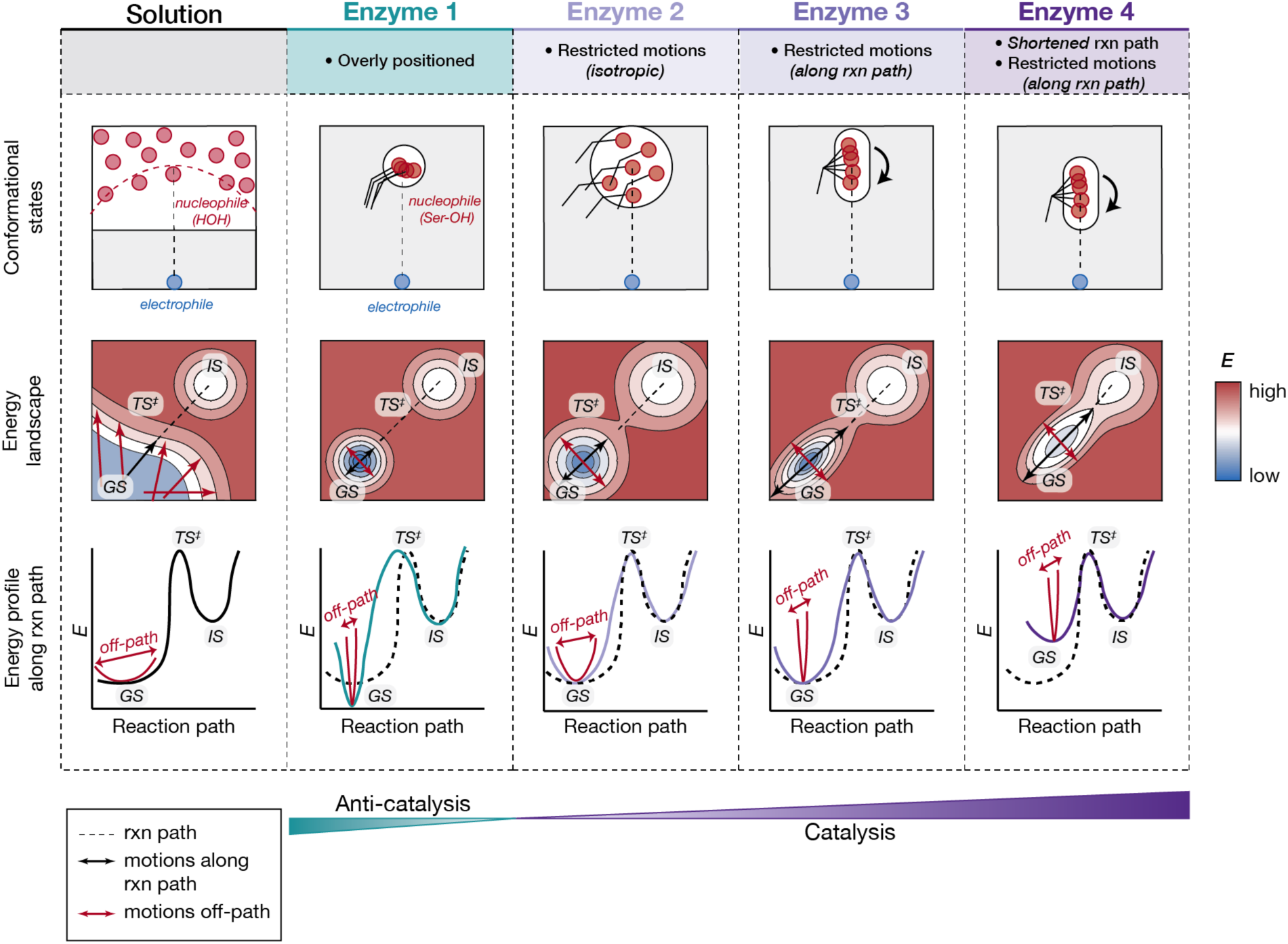
Synergy of positioning and motion in serine protease catalysis. Schematics for a solution reaction and four different enzyme examples are shown for comparison and described in the text. The top row shows the positioning between the nucleophile (red) and the electrophile (blue) for each reaction. The second row shows a two-dimensional energy landscape for the corresponding reaction. Dashed lines indicate the shortest reaction path; black arrows indicate motions along this reaction path whereas red arrows show a subset of possible off-pathway motions. The contour levels are colored by their energies, with blue to red corresponding to energies from low to high. The bottom row shows the one-dimensional “energy profile” of each reaction, corresponding to the slice of the energy landscape along the shortest reaction path. The energy wells and arrows colored in red represent a different “slice” of the energy landscape that is not along the reaction path (“off-path”). GS, ground state; TS^‡^, transition state; IS, intermediate state.

The synergy of positioning and motion is achieved by ground state destabilization. A partially eclipsed serine sidechain and a shorter-than-ideal van der Waals distance in the ground state allow the enzymatic reaction to commence from a higher energy state that is further along the reaction coordinate than the ground state in solution (Fig. 7, Enzyme 4). Ground state destabilization creates a downhill energetic gradient that selectively favors the serine rotation and bond formation and partially offsets the energetic barrier to reaction.

The energy landscape and catalytic model presented here are derived with a minimal set of features local to the active site. Given the ability of this model to account for the observed serine protease catalysis, there is no indication that long range effects contribute substantially to catalysis. Nevertheless, it will be important to test for and evaluate such effects in other systems, in particular those exhibiting allostery (*94*). The ensemble–function framework will be especially valuable for dissecting the interplay of conformational and energetic effects in these systems. Potential catalytic factors such as long-range electrostatic interactions (*78*, *80*), may be present in other systems and can be included in future ensemble-function models. In contrast, potential catalysis associated with barrier-crossing dynamics (*83*, *85*, *86*) cannot be accessed by this approach, as they are beyond the pseudo-equilibrium framework of transition state theory and our statistical mechanics treatment.

Textbook and literature descriptions of enzyme catalysis often emphasize the convergent evolution of the Ser•His•Asp catalytic triad across serine proteases (*19*, *20*, *95–97*). Our analyses revealed a still more general feature––the nucleophilic elbow––and its underlying mechanistic and evolutionary significance. This motif connects the nucleophile and the oxyanion hole and restrains their conformations in a catalytically adventitious manner. Additional groups from the structural scaffold are recruited to provide general base catalysis, the remainder of the oxyanion hole, and substrate binding and positioning; these groups differ in different enzymes and folds, with only a subset containing the classic Ser•His•Asp catalytic triad (*98*, *99*). Understanding how Nature has combined and elaborated upon catalytic motifs to generate catalytically productive active site features may lead to insights for enzyme design.

The ensemble–function approach developed here differs fundamentally from site-directed mutagenesis, which compares one enzyme to another, rather than an enzymatic reaction to its uncatalyzed counterpart. As a result, mutational rate effects are not a direct readout of catalytic contribution (*100*). In particular, mutations can have deleterious effects that extend beyond the loss of catalytic features due to structural rearrangements that stabilize alternative, unreactive conformations. For example, the large deleterious effects upon the mutation of trypsin’s catalytic triad aspartate to asparagine arises in part from flipping the catalytic histidine to a state that cannot act as a general base (*101*). In contrast, ensemble–function analyses compare the enzymatic and the uncatalyzed solution reaction, and thus can identify and quantify individual catalytic features.

The ensemble–function approach can be readily extended to the investigation of new systems. The PDB currently contains >300 proteins with >100 repetitive structures at high resolutions (≤ 2 Å) that can be built into pseudo-ensembles (July 2024). The feasibility of studying each system via pseudo-ensembles will depend on whether functionally relevant states or their mimics are available. Nevertheless, new atomic-level ensemble information can be obtained from room temperature X-ray crystallography (*28*) and from cryogenic electron microscopy at sufficiently high resolutions. Ensemble–function analyses can also be applied to molecular dynamics simulation data, to test their ability to reproduce atomic-scale properties observed in experiments and to generate new models that can be tested via subsequent experiments.

Most generally, conformational ensembles connect molecular properties to free energy, a connection that is needed to define enzyme catalysis and all aspects of protein function. The ensemble–function approach provides quantitative models for enzyme catalysis, and will be of value in understanding still more complex processes, including allostery (*94*) and the coordination of molecular machines (*102*).

## Acknowledgments

We thank members of the Herschlag lab for thoughtful discussions and manuscript feedback.

## Author contributions

Conceptualization: SD, RCK, JPG, MMP, DH

Methodology (*Ensemble*PDB): SD, RCK, JPG, FY

Data curation – serine protease pseudo-ensembles and additional crystallographic data: SD, RCK, JPG

Data curation – QM calculations and MD simulations: EP, VWDC, MZ

Investigation – serine protease pseudo-ensemble conformational analyses: SD, RCK, JPG, JPS

Investigation – serine protease catalytic model and energetic analyses: SD

Investigation – QM calculations and MD simulations: SD, EP, VWDC, MZ

Investigation – nucleophilic elbow analyses: SD

Visualization: SD

Funding acquisition: DH, TJM

Writing – original draft: SD, DH

Writing – review & editing: all authors

## Data and materials availability

All data and computer code used for the analyses are available in the supplementary materials or at 10.5281/zenodo.13336748. The *Ensemble*PDB package is available at: https://github.com/Herschlag-Lab/EnsemblePDB.

