## Supplemental Materials for "Conformational Ensembles Reveal the Origins of Serine Protease Catalysis"

**The PDF file includes:**

Methods

Supplementary Text

Figs. S1 to 61

Tables S1 to 26

References (*103*–*197*)

**Methods:**

*Building pseudo-ensembles of serine proteases.*

Pseudo-ensembles of serine proteases were built using the *Ensemble*PDB pipeline (available at: <https://github.com/Herschlag-Lab/EnsemblePDB>; see flowchart in fig. S60). A reference wild-type structure was chosen for each serine protease, and the PDB was searched for all structures with one or more chains with higher than 95% sequence identity to the reference sequence. (We used only wild-type sequences for building pseudo-ensembles; this search allowed us to capture mutant sequences that may be of interest, but we did not include these mutants in this study.) These structures were filtered by retrieving their metadata and excluding structures (1) at a resolution of >2.5 Å; (2) containing keywords such as “radiation damage”, “zymogen”, “proenzyme” or names associated with the zymogen form of that serine protease; or (3) from an organism different from the reference. Then, for each serine protease, the PDB files were reformatted such that the residue numbering and chain names are consistent. For PDB files with multiple chains aligned to the reference sequence, each chain was saved separately as an individual observation of the molecule. The renumbered PDBs were aligned on the C𝛼 atoms of the protease chain. All atoms that are symmetric in the PDB naming scheme were renamed consistently based on their distances to the same atoms in the reference PDB. (For example, the Oδ1 and Oδ2 atoms of aspartate residues are assigned randomly in different PDBs; if Oδ1 in the current PDB for a certain aspartate is closer to Oδ2 of the same residue in the reference PDB compared to Oδ1 in the aligned ensemble, then Oδ1 of the current PDB would be renamed to Oδ2, and *vice versa*.) The aspartate in the catalytic triad was renamed such that Oδ1 is the oxygen closer to the histidine Nδ1.

The resulting renamed PDBs were then manually and computationally examined and classified into different liganded states. A list of different types of ligands is shown in fig. S5. If no molecule was bound at the active site or only solvent molecules such as water and sulfate were bound, the structure was assigned to the apo class. If (1) a peptide or peptide analog was bound at the active site, (2) a carbonyl carbon was within 4.0 Å to the serine nucleophilic atom (O𝛾) but not covalently bound (distance longer than 2.2 Å), and (3) an oxygen atom was within 4.0 Å to both oxyanion hole hydrogen bond donors, the structure was assigned to GSA-bound class. For structures in the TSA-bound class, each must have a covalently bound ligand (within 2.2 Å to the serine nucleophile O𝛾) that is in the list of known TSA types (fig. S5). The acyl-enzyme class includes structures with ligands covalently bound to the serine nucleophile that are sp^2^ hybridized.

During this process, the ligand electrophilic group and its atoms were identified for use in subsequent geometric calculations. For the GSA-bound class, the electrophilic atom is the carbon atom from the non-covalently bound ligand that is the closest to the serine nucleophilic oxygen. For the TSA-bound and acylenzyme class, the electrophilic atom is the ligand atom covalently bound to the serine nucleophile.

After ligand classification, we built wild-type ensembles of each serine protease in each of the liganded states by selecting all PDBs with no mutations (based on PDB annotation) and aligning the structures within each ligand class. Different atoms were used in alignments to evaluate different conformational properties, e.g. catalytic triad atoms were aligned to assess local conformational changes at the active site, whereas all C𝛼 atoms were aligned to assess whether there are significant global changes. Details of each alignment are described in figure legends.

*Calculating interaction geometries in pseudo-ensembles.*

All the geometric parameters of interest, as defined in the main text and supplemental figures, were measured in the PDB structures using the *biopython* package (*103*). The residues and atoms involved in the interaction of interest were identified in each PDB structure and the distances, angles and dihedrals were calculated using the *Bio.PDB* module. All backbone 𝜓, 𝜑 and rotameric torsion angles were specified using the 0 to 360º range; non-rotameric torsion angles were normalized to ranges defined in (*2*). Other dihedral angles ($\phi$_attack_ for the reactant positions and $\phi$_HB_ for hydrogen bonds) were normalized such that they lie in the range of 0 to 90º, as follows:

$$normalized \phi=\left\{ \begin{aligned} \phi, & 0<\phi\leq90 \\ 180-\phi, &0<\phi\leq180 \\ \phi-180, &180<\phi\leq270 \\ 360-\phi, &270<\phi\leq360 \end{aligned} \right.$$

For each geometric parameter, the most probable (lowest energy) value (“mode”) was calculated by fitting the distribution into a probability density function (*scipy.stats.gaussian_kde*) and taking the highest probability value. Standard deviations (*s.d.*) were calculated using *numpy.std*.

*Obtaining knowledge-based distributions of interaction geometries.*

We searched for small molecule interactions that mimic interactions between groups in the enzyme in the Cambridge Structural Database (CSD) (*104*) using the CSD Python API. The interacting groups were specified using SMARTS strings (*105*) (table S6) and the matching substructures were searched using the *ccdc.search* module. The searches were constrained such that the interactions match distance and angular criteria as specified in table S6. Searches for van der Waals interactions can include cases where the atoms are shielded by another atom or interaction; therefore, we used the “line-of-sight” method to eliminate background that contains these shielded, indirect interactions (*106*). Duplicated entries found due to molecular symmetry were removed.

We obtained the knowledge-based distribution of amino acid sidechain torsion angles from the Dunbrack “Backbone-dependent Rotamer Library” dataset (*5*). For hydrogen bonds between amino acids (His•Asp and His•Ser), we searched the *Top2018* dataset (*107*), a set of curated high-quality, low-redundancy PDBs, for residues in proximity with their hydrogen bonding heavy atom distance $\leq$4 Å. For serines in contact with disulfide bonds (Ser•SS), we first searched for disulfide bonds where the sulfur atoms (S𝛾) between two cysteine residues are covalently bonded (within 3 Å). Then, we calculated the centroid of the two sulfur atoms and searched for serine O𝛾 atoms that are within 6 Å of the SS centroid.

*Quantum mechanical calculation for the solution reaction path.*

The initial tetrahedral intermediate structure for the system was obtained by extracting the coordinates from the PDB structure 1TYN. The structure was protonated using LEaP (*108*) and the protonation states of the titratable sites were determined by their p*K*_a_ values. The system included the sidechain of Asp102 (cut between C$\beta$ and C$\alpha$), His57 (cut between C$\beta$ and C$\gamma$), and Ser195 (cut between C$\alpha$ and C, and between C$\alpha$ and N), and a portion of cyclotheonamide molecule (atoms N8, C9, C10, C11 and O41). The cut bonds were capped with hydrogen atoms. This process yielded a system comprised of a *N*-methylacetamide (NMA) bonded to ethoxide, an imidazolium(+) and an acetate(–). Additionally, two water molecules hydrogen bonded to NMA’s carbonyl oxygen atom were added to mimic the oxyanion hole hydrogen bond donors.

To optimize the tetrahedral intermediate structure, the above coordinates were subjected to a careful series of constrained optimizations to prevent disrupting the sp^3^ hybridization and reverting to the ground state structure. The constraints were gradually lifted via the following procedure. First, we performed constrained optimization to relax the position of the water molecules while maintaining the tetrahedral structure and the hydrogen bonds. Restraints were placed on both N atoms and their bonded H atoms in the imidazole, the O and H atoms (hydroxyl moiety) in acetate, the ethoxide O atom, and the C atom of the carbonyl moiety in NMA. Second, we performed constrained optimization releasing the restraints previously placed on the acetate fragment and the N atom that is closer to acetate in the imidazole. Third, we performed constrained optimization releasing the restraints previously placed on the imidazole ring. Finally, an unconstrained optimization was performed to yield the minimum energy structure for the tetrahedral intermediate.

To obtain the reactant structure, we performed an unconstrained energy minimization from the coordinates extracted from the 1TYN PDB structure above. Frequency calculations were performed on the optimized structures to verify that they represent minimum energy structures.

The “reaction path” was defined as the minimum energy path connecting the tetrahedral intermediate and the ground state structure, and was calculated using the growing string method (*109*) implemented in the pyGSM python package (<https://github.com/ZimmermanGroup/pyGSM>). All calculations were performed using an implicit solvation model, COSMO (*110*) (with a dielectric constant of 78.39) and unrestricted B3LYP/6-31G*, on two GeForce GTX 970 GPU cards using TeraChem (*111*–*113*).

The actual reaction in water does not include the imidazole and acetate molecules, yet we were not able to obtain a reaction path when either the imidazole or the acetate was absent from the system, despite numerous attempts and different sequences of constrained optimizations; the calculations collapsed on the ground state structure instead of yielding a tetrahedral intermediate.

*Molecular dynamics simulations for the solution ground state.*

The initial structure of *N*-methylacetamide (NMA) was prepared using the LEaP program in AMBER 22 (*108*) with the ff19SB force field (*114*). We solvated the system with OPC water molecules (*115*) using a buffer of 15 Å, resulting in an initial box dimension of 39.88×40.01×38.13 Å^3^. We first performed energy minimization of the NMA molecule by posing harmonic constraints on the solvent. The system was gradually heated to 300 K over 6 ns under NVT conditions, and both the temperature (300 K) and density (1 bar) were equilibrated for another 8 ns. Our production trajectories were 100 ns long in the NVT ensemble (*T* = 300 K) with a timestep of 2 fs. We used the SHAKE algorithm (*116*) to constrain the H atoms and the Langevin thermostat with a collision frequency of 5 ps^-1^ to regulate temperature. The particle mesh Ewald method (*117*) with a real space cutoff of 8 Å was employed to describe long-range electrostatic interactions. We performed three replicates using the same procedures.

We measured geometric parameters (*d*_attack_, *α*_attack_, $\phi$_attack_, defined in Fig. 2A) using AMBER’s CPPTRAJ program (*108*) in 5000 snapshots sampled from the trajectory (with a 0.02 ns spacing). For each snapshot, the water molecule that is the closest to the electrophile (i.e. the carbonyl carbon of the NMA molecule) was designated as the nucleophile.

*Identifying torsion angle changes across reaction states.*

Torsion angles (backbone 𝜓, 𝜑 and sidechain 𝜒^1^, 𝜒^2^, 𝜒^3^, 𝜒^4^ whenever applicable) of all PDB structures in the pseudo-ensembles were calculated using the *Ensemble*PDB*.analyze.rotamer* module. For sidechains with alternative conformers, each conformer was counted as an individual observation.

Trypsin, chymotrypsin and elastase have at least five structures in each sub-ensemble (apo, GSA-bound and TSA-bound) and these three serine proteases were used in the following comparisons: (i) distributions of each torsion angle from the apo *versus* the GSA-bound ensemble; (ii) an analogous comparison for GSA-bound *versus* TSA-bound ensembles. Two-sided Kolmogorov–Smirnov (K–S) tests (*17*, *18*) were used to calculate­ *p*-values (*scipy.stats.kstest*). For each comparison, the significance of a torsion change was determined by a *p*-value threshold corrected for multiple hypothesis testing using the Bonferroni correction (*p*-value threshold = 0.05/number of all rotamers compared).

We assessed the accuracy of the sidechain conformers modeled in the PDB structures using *Ringer* (*118*), a program that samples electron density around modeled sidechain torsion angles. We identified the peaks of the *Ringer* electron density profiles (with 5º bins) and determined whether the modeled sidechain torsion angles match the *Ringer* peaks. We could perform *Ringer* analysis on a majority of the PDB structures (138 out of 196), as they have map coefficients deposited. We found that buried residues have 94% of the torsion angles within 15º of the *Ringer* peaks whereas solvent exposed residues have 85% (fig. S61). Therefore, we performed the same statistical tests described above using buried residues only (as our highest confidence dataset). The DSSP program (*119*, *120*) was used to calculate relative solvent accessibility of each residue, and all residues with a relative solvent accessibility of less than 0.25 were assigned as “buried”.

The statistical stability of the significance of the Ser195 𝜒^1^ changes observed were further tested by bootstrap analyses. The above tests were repeated 1000 times, with each repeat resampling from the Ser195 𝜒^1^ distributions (with the sample sizes maintained; see fig. S11), and *p-*value distributions were obtained. To determine whether the heterogeneity of the ligands included in the TSA class affects the significance of the Ser195 𝜒^1^ change, we performed the same bootstrap analyses for two additional comparisons between the GSA-bound ensembles and subsets of the TSA ensembles: TSAs that are reacted from (1) carbonyl compounds and (2) boronic acid derivatives. In addition to the K–S tests (*20*) which determine whether the distributions are different, we also performed Mann–Whitney U-tests (*121*) using *scipy.stats.mannwhitneyu* to determine whether one distribution is numerically greater than the other.

Torsion angle changes that were significant and conserved for the three serine proteases in clan PA were identified as follows. The *Bos taurus* trypsin sequence was used as the template sequence, and chymotrypsin (*Bos taurus*) and elastase (*Sus scrofa*) sequences were aligned to trypsin using the *biopython* package (*Bio.pairwise2.align.globalxx*) (*103*). For each aligned residue position excluding gaps and insertions, the *p-*values for each torsion angle comparison determined above (from K–S tests) were queried and compared. Comparisons were performed when the same sidechain torsion angle exists for all three serine proteases at the same residue position, regardless of whether the amino acid is conserved.

*Calculating the contributions to covalent bond formation arising from conformational changes between the GSA- to the TSA-bound state.*

Three types of motions are involved in forming the O–C bond between the catalytic serine and the substrate amide carbon: (1) bond rotation of the serine sidechain, (2) substrate geometric changes from sp^2^ to sp^3^, and (3) relative translational motion between the enzyme and the substrate (Fig. 3D).

To calculate the distance change between the reacting atoms resulting from the catalytic serine sidechain rotation, we used *PyMol* (*122*) to model the serine rotation from the ground state to the transition state while fixing all other atomic coordinates. Specifically, for each enzyme, we calculated the list of the catalytic serine 𝜒1 angles observed in the TSA-bound structures, and we iterated through the GS-bound structures to set their serine 𝜒1 angles to each of the 𝜒1 angles in the TSA-bound list using the *cmd.set_dihedral* tool. We then calculated the change in the distance between the serine O𝛾 and the electrophilic carbon on the substrate amide (Δ*d*_O•C_) resulting from the rotation.

We used the TSA-bound structures to model the substrate sp^2^ → sp^3^ transition. We first found the plane defined by the three atoms covalently bonded to the electrophilic atom on the TSA other than the catalytic serine, as these atoms belonged to the unreacted substrate. We then projected the electrophilic atom onto the plane to find its original position in the unreacted substrate. The distance change (*d*_O•C_) was calculated by the difference in the distance between the catalytic serine O𝛾 and the projected electrophile *versus* the observed bond length.

To calculate the contribution from the relative translational motion between the reactant and the enzyme, we paired every GSA-bound structure with every TSA-bound structure and locally aligned them on atoms of the catalytic triad residues. To remove effects from the serine rotation and the sp^2^ → sp^3^ change, we used the TSA-bound serine O𝛾 as a reference point and calculated its distance to the centroid of the three substituent atoms covalently bonded to the electrophilic atom on the substrate amide in the GSA-bound structure, as well as its distance to the centroid of the corresponding substituent atoms in the covalent TSA. The distance change (Δ*d*_O•C_) resulting from translational motion is calculated as the former distance subtracted from the latter.

The total distance change needed for bond formation was calculated by taking the difference in *d*_attack_ (defined in Fig. 2A) in the GSA-bound and the C–O bond lengths in the TSA-bound structures.

The above procedures were performed for serine proteases as well as additional non-protease enzymes in clan SC. Non-protease members of clan SC that perform nucleophilic addition on carbonyl compounds were collected by searching the Mechanism and Catalytic Site Atlas (M-CSA) database (*93*) (see *Determining the presence of nucleophilic elbows in additional enzymes*) and selecting enzymes that belong to the same homologous superfamily as clan SC proteases. Their structures were classified into apo, GSA-bound and TSA-bound states using approaches analogous to the ligand classification performed for serine protease structures, with details described below (*Classifying ligand bound states of nucleophilic elbow-bearing enzymes*).

*Calculating the catalytic contribution from reactant positioning.*

The catalytic contribution from positioning is quantified as the difference in conformational entropy (*S*_conf_) in going from the ground to the transition state on the enzyme *versus* that in the solution reaction. We made the following simplifying assumptions:

(1) The transition state ensemble is as narrow or narrower than the ground state ensemble for both the enzymatic and solution reactions; a narrow transition state relative to ground state ensemble is expected based on the orientational requirements for (partial) bond formation in the transition state.
(2) The transition state is restricted to the same extent in the enzymatic and the solution reactions.
(3) Groups that are not involved in direct bond formation (i.e. non-nucleophilic water molecules in the solution reaction and non-reacting residues in the enzymatic reaction) have, in sum, negligible changes in their conformational entropies in going from the ground to the transition state. This assumption is consistent with the absence of additional significant conformational changes beyond the groups directly involved in the reaction.

These assumptions reduce the calculation to a comparison of the conformational entropy of the reacting groups in the enzymatic and solution ground state.

For the enzymatic ground state, the atomic coordinates of the serine nucleophile (Oγ) and the substrate amide (C, O, Cα of the P1 residue and N of the P1′ residue) from the GSA-bound pseudo-ensembles were extracted. For the solution ground state, two datasets were used: the atomic coordinates of the nucleophilic oxygen and the amide carbonyl atoms extracted (1) from the 5000 snapshots of the MD simulation of water and NMA (see *Molecular dynamics simulations for the solution ground state*) and (2) from the knowledge-based distributions of hydroxyl•amide interactions in the CSD (see *Obtaining knowledge-based distributions for small molecule interactions*). The coordinates were then rotated such that for each entry the electrophilic carbon is at the origin, the substrate carbonyl oxygen is aligned along the +x axis, and the amide plane aligns with the *xy* plane. The resulting enzyme and solution distributions of the nucleophilic oxygen coordinates, with dimensions of (10 Å)^3^, were cut into three-dimensional histograms using *numpy.histogramdd* with the number of bins on each dimension (*n*_bins_) specified. We calculated entropy varying *n*_bins_ from 1^3^ to 200^3^, corresponding to bin volumes of (10 Å)^3^ to (0.05 Å)^3^, to determine the extent to which the bin volume affect entropy values (Fig. 5C and fig. S25). The probability of observing the reactants in each bin was calculated by dividing the histogram counts by the total number of data points. The *S*_conf_ associated with this discrete probability density function was then calculated following the formalism of Gibbs entropy, $S= -k_{B}\sum_{i} P_{i}ln(P_{i})$ where $k_{B}$ is the Boltzmann constant, *i* is a certain bin in the ensemble, and *P_i_* is the probability of observing the system in bin *i*. The value of $\sum_{i} P_{i}ln(P_{i})$ was calculated using *scipy.stats.entropy.* Bootstrap analyses were performed to quantify the variance associated with the entropy values by repeating the calculation 200 times, with each repeat randomly selecting a specified number of data points (*n*) from the distributions, and standard deviations of *S*_conf_ were calculated. We also confirmed that the sample sizes are appropriate and give stable entropy values by resampling the distributions using a range of *n* from 10 to 50000 and calculating the resulting *S*_conf_. (fig. S25).

Our calculations are based on crystallographic data, which do not contain information about vibrational entropies or the rotational entropy of the nucleophilic O–H bond (as hydrogens are typically not detected by X-ray diffraction). Vibrational entropy effects are not expected to yield substantial catalytic effects, as higher-energy vibrational states are rarely populated due to their larger gaps in energy levels (much larger than for translational and rotational states) (*9*, *123*). The serine O–H bond rotation can be more restricted than a water O–H bond in solution and thus may provide additional catalytic advantages. Completely freezing the rotation of one bond gives an entropic difference of ~0.8–1.3 kcal/mol (*24*), and we expect a smaller difference between serine proteases and water, as the water molecules are generally hydrogen bonded in solution and therefore not freely rotating.

*Energy calculation using knowledge-based distributions.*

Knowledge-based distributions for geometric parameters were converted into energy functions using the *Ensemble*PDB.*analyze.energy* module. The distributions were cut into a specified number of bins (*n*_bins_) and the number of samples within each bin (*n*) was calculated. The *n*_bins_ were set depending on the sample size of each distribution such that the bin widths are between 0.08 to 0.1 Å for distances and between 2 to 4° for angles and dihedrals and that the resulting energy functions were not overly rugged. The probability of observing a sample in a certain bin, *p*_i_, was calculated as *n* divided by the total sample size (*n*_total_). The reference state (*E*_ref_) was defined as a random distribution where the probability of observing any state is *P*_ref_ = 1/*n*_bins_ such that *E*_ref_ = 0_._ The energy of each bin relative to *E*_ref_ (Δ*E*) was calculated by

–*k*_B_*T*•ln(*P*_i_/*P*_ref_) where *k*_B_ is the Boltzmann constant and *T* = 298 K. The energy functions were saved as tables (*.csv* files) specifying each bin and its Δ*E* value. To calculate Δ*E* for a certain geometric measurement (*x*) using the corresponding energy function, the bin containing that value *x* was found and the corresponding Δ*E*(*x*) value was queried; if the geometric value was outside the range of geometries covered by the energy function (because the conformer in question was too rare to be present in the original crystallographic distribution), then the maximum Δ*E* value was assigned.

*Determination of errors for the trypsin catalytic model.*

The error for general base catalysis represents lower and upper limits of its catalytic contribution, calculated by Brønsted $\beta$ from 0.5 to 1 (see supplementary text S5). The error for reactant positioning is the standard deviation of entropy vales using a range of bin sizes from (2 Å)^3^ to (0.02 Å)^3^. The errors for ground state destabilization mechanisms, determined from knowledge-based energy functions, are standard deviations from the energies calculated from the distribution of geometric parameters observed in pseudo-ensembles.

We note that other sources of errors include assumptions made in each calculation, the accuracy of analogs in representing real reaction states, and the accuracy of knowledge-based energy functions. We discussed these potential sources of errors in the main text (*The serine protease catalytic ledger*) and in the *Methods* section corresponding to each calculation.

*Determining the presence of nucleophilic elbows in additional enzymes.*

We searched the Mechanism and Catalytic Site Atlas (M-CSA) database (*93*) for enzymes (*n*_total_ = 998) whose annotated catalytic residue contain the keyword “nucleophile” in their “role summary”, resulting in 229 enzymes (with a total of 256 nucleophilic residues, as some enzymes have more than one nucleophile). We filtered the enzyme list such that no enzyme is redundant with those already in the serine protease dataset collected in this work (table S3) and the serine and cysteine proteases collected in Buller and Townsend (table S22) (*28*, *29*) (*n* = 16 and 18, respectively). Next, the enzyme list was manually screened to determine whether each reaction involves a covalent oxyanion species by examining their arrow-pushing mechanisms in the M-CSA database and the cited references; 102 out of 229 enzymes met this criterion. For the 18 enzymes from Buller and Townsend and the 102 enzymes from M-CSA, we searched the PDB for structures with >95% sequence identity with the reference PDB annotated in M-CSA using the *Ensemble*PDB package and built pseudo-ensembles using the same procedures as described above for serine proteases. We examined ligand bound structures for each enzyme to determine the identity of enzyme groups consisting of the “oxyanion hole” that can stabilize the oxyanionic reaction intermediate. We were able to identify the oxyanion hole hydrogen bond donor(s) for all 18 enzymes from Buller and Townsend and for 91 of the 102 enzymes in our list from the M-CSA database. The rest of the M-CSA enzymes (11) do not have relevant ligand-bound structural data available, and thus the oxyanion hole groups could not be determined with confidence, and they were excluded from further analysis.

We combined our serine protease data, the Buller and Townsend proteases, and the M-CSA enzymes to obtain a dataset of 125 total enzymes (126 nucleophilic residues), and their oxyanion holes were annotated as follows: if one of the oxyanion hole donors is the backbone amide nitrogen of the nucleophilic residue, then the oxyanion hole is “N-type nucleophilic elbow” (67 of 126); if the backbone amide nitrogen from the residue immediately next to the nucleophilic residue (sequence number +1) is the hydrogen bond donor, then the oxyanion hole is “N+1-type nucleophilic elbow” (35 of 126); all other oxyanion holes were annotated as “other” (24 of 126).

*Classifying ligand-bound states of nucleophilic elbow-bearing enzymes.*

The ensembles of nucleophilic elbow-bearing enzymes were classified into different ligand-bound states (apo, GSA-bound, TSA-bound, *et cetera*) based on their reaction chemistry. We followed a similar approach as described for serine proteases using the *biopython* package to parse the PDBs. For each enzyme structure, the nucleophilic atom was found based on the M-CSA annotation of the corresponding enzyme. We determined whether a ligand is bound at the active site by looking for any non-enzymatic chains or compounds in the PDB that are within 4.5 Å to the nucleophilic atom, excluding cofactors and metal ions. To find the electrophilic atom, we identified the atom closest to the nucleophile on the ligand (if present) and the nucleophile•electrophile distance was calculated. The ligand was classified as covalently bound if this distance is smaller than 2.2 Å, and otherwise as non-covalently bound; for bonds between sulfur atoms this constraint was relaxed to 3.0 Å. Next, we determined all atoms that are covalently linked to the electrophile (using the same covalent constraints) and calculated all bond angles between the electrophilic atom and the substituent atoms. The hybridization state of the electrophilic group was determined as follows: (1) if the average bond angle is 120 ± 5°, then this group is sp^2^ hybridized; (2) if the average bond angle is 109.5 ± 5°, then this group is sp^3^ hybridized; (3) if there is at least one bond angle within 120 ± 5° and at least one bond angle within 90 ± 5°, then this group is dsp^3^ hybridized; (4) if the calculated bond angles match none of these criteria, then it is assigned as “other”. We determined potential oxyanion(s) by selecting oxygen atoms that are covalently bonded to the electrophile and calculated the distances between the potential oxyanion atoms to the oxyanion hole hydrogen bond donors that were manually identified above (*Determining the presence of nucleophilic elbows in additional enzymes*).

If the enzyme performs a nucleophilic addition reaction on a carbonyl group, then the ligand bound state of the enzyme structure was determined based on the following criteria: (1) if no ligand is bound, then the structure is in the apo state; (2) if a ligand is non-covalently bound and sp^2^ hybridized, and an oxyanion is within 4.0 Å to either of the oxyanion hole hydrogen bond donors, then it is GSA-bound; (3) if a ligand is covalently bound and sp^3^ hybridized, and an oxyanion is within 4.0 Å to either of the oxyanion hole hydrogen bond donors, then it is TSA-bound; (4) if a ligand is covalently bound and sp^2^ hybridized, and an oxyanion is within 4.0 Å to either of the oxyanion hole hydrogen bond donors, then it is in the acylenzyme state. If, instead, the enzyme performs S_N_2 nucleophilic attack, then the criteria were set as follows: (1) if no ligand is bound, then the structure is in the apo state; (2) if a ligand is non-covalently bound and sp^3^ hybridized, and at least one oxyanion is present, then it is GSA-bound; (3) if a ligand is covalently bound (<2.5 Å, constraint relaxed due to longer bonds in tungstate and vanadate TSAs) and dsp^3^ hybridized, and at least one oxyanion is present, then it is TSA-bound; (4) if a ligand is covalently bound and sp^3^ hybridized, and at least one oxyanion is present, then it is in the product state (“E–P”).

*Alignment of nucleophilic elbows across different enzymes.*

Structural alignments of nucleophilic elbows were performed for enzymes that perform nucleophilic addition on carbonyl compounds. Structures in the ensembles were separated into groups based on nucleophilic elbow type (N or N+1), the rotameric state of the nucleophilic residue sidechain (*gauche–*, *gauche+*, or *trans*), and their ligand-bound states (GSA-bound, TSA-bound, or acylenzyme). Within each group, structures were aligned on the atoms involved in the nucleophilic elbow motif using PyMol (*cmd.pair_fit*). For the N-type nucleophilic elbows, these atoms involve the N, Cα, Cβ and the nucleophilic oxygen or sulfur atom on the nucleophilic residue, the ligand electrophilic atom, and the oxyanion; for the N+1-type nucleophilic elbows, these atoms involve the C, Cα, Cβ and the nucleophilic oxygen or sulfur atom on the nucleophilic residue, the N atom on the residue next to the nucleophilic residue, the ligand electrophilic atom, and the oxyanion (fig. S47).

**Supplementary Text:**

**Text S1.** Literature perspectives on the serine protease catalytic triad.

Early catalytic proposals focused on a potential proton relay from the His general base to the Asp of the catalytic triad (*124*, *125*) and more recent ones on a short, strong hydrogen bond (SSHB) or low-barrier hydrogen bond (LBHB) between the His and Asp (*126*, *127*). NMR and structural studies provided strong evidence against the proton relay model, as the histidine remains protonated in TSA-bound complexes [see (*128*), *Where Are the Protons? One-Proton versus Two-Proton-Transfer Mechanisms*]. The second proposal, the SSHB/LBHB model, emphasizes a short His•Asp hydrogen bond that arise from a “nonaqueous” enzyme environment and special energetic contribution from the matched p*K*_a_ values of the hydrogen bond donor and acceptor (when Δp*K*_a_ = 0). Subsequent experiments refuted this model:

1. The enzyme hydrogen bonds do not have unusual geometric properties compared to those in aqueous solution, as hydrogen bond lengths follow a linear relationship with donor/acceptor Δp*K*_a_ that is insensitive to environment (*129*, *130*).
2. The p*K*_a_ values of the histidine and the aspartate are not matched in the TSA-bound state (*131*, *132*) and the proton is associated with the histidine instead of equally shared (*133*–*135*).
3. Even enzyme hydrogen bonds with matched p*K*_a_ values do not provide special energetic contributions––a linear relationship was found between hydrogen bond energy and Δp*K*_a_ in model compounds, with no special energetic effects at Δp*K*_a_ = 0 (*136*).

Our pseudo-ensemble analyses provided additional evidence against special geometric properties of the His•Asp hydrogen bond that would contribute to catalysis (see *Catalysis from the catalytic triad hydrogen bonds* and supplementary text S7 to 8).

**Text S2.** Serine protease ground state analogs (GSAs) bind in substrate-like conformations.

To follow the catalytic cycle, we classified serine protease structures into different ligand-bound states, including the apo, ground-state analog (GSA)-bound, transition-state analog (TSA)-bound and acylenzyme states (fig. S5). TSA-bound states and acylenzymes are covalent species and their covalent constraints limit the ability to explore alternative conformations; our analyses also indicate that these species are bonded with the expected catalytic groups. GSAs are crystallized in their unreacted form, so it is critical to ask whether their bound structures reflect the true ground state geometry. Our collected GSAs include bovine pancreatic trypsin inhibitors (BPTIs) [or Kunitz-type inhibitors] (*137*, *138*), Bowman-Birk inhibitors (BBIs) (*139*, *140*), pancreatic secretory trypsin inhibitor [or Kazal-type inhibitors] (*141*, *142*), *Streptomyces* subtilisin inhibitor (SSI) (*143*, *144*), *et al*. These GSAs are competitive inhibitors of serine proteases that follow the same inhibition mechanisms (*145*). Previous studies along with our analyses strongly suggest that they bind in the same conformation as cognate substrates.

The GSAs undergo a fast acylation step, indicative of ready access to a reactive E•S conformer when bound. An equilibrium between the acylenzyme and the E•S complex is established within seconds wherein the E•S complex is thermodynamically favored (*146*). The back reaction is favored over deacylation, presumably due to interactions that stabilize the leaving group in the active site and other features that increase the probability of the reverse reaction. These GSAs typically contain disulfide bonds (*147*–*149*), extensive hydrogen bond networks (*146*, *150*, *151*) and/or cyclic backbones (*152*), and their removal leads to decreased inhibition (and increased overall reaction) (*148*, *153*). These structural features stabilizing the uncleaved state are conserved within evolutionarily-related inhibitors and have convergently evolved across different inhibitor types, supporting their role in the inhibition mechanism (*145*, *154*). Overall, substantial literature evidence suggests the same inhibition mechanism across these GSAs––they are acylated in a substrate-like manner yet the E•S complex is favored over deacylation.

In addition, our (and prior) structural analyses provided no evidence for unreactive binding modes formed by these GSAs. A consistent active site conformation was found across GSA-bound serine proteases (fig. S9 and table S7); the relative orientation between the reactants matches the ground state in the solution reaction in our QM calculation and the predictions for reactive nucleophilic attack conformation from Burgi and Dunitz (*60*, *61*). Further, an unreactive binding conformation would involve interactions that stabilize the ground state such that the reaction barrier is higher than that for a canonical substrate; destabilizing interactions in alternative unreactive states are not expected. We found destabilized sidechain torsion angles of the catalytic serine and a shorter-than-ideal distance between the reactants in the GSA-bound state, both of which are relieved in the TSA-bound state, consistent with a reactive conformer facilitating reaction instead of an unreactive conformer that favors binding.

In contrast to the GSAs that binds in reactive conformers, a subset of noncovalent inhibitors was found to bind in alternative modes, and they were not included as GSAs:

(1) Peptides (or peptide analogs) with C-terminal tyrosine, phenylalanine, or phenol groups that form CH–𝜋 interactions at the S1 or S2 binding site (*155*, *156*). These inhibitors adopt a different binding conformation where the nucleophilic serine is displaced and far from the bond to be cleaved. These inhibitors were not included in the GSA-bound class, as enforced by the criteria that the reacting carbon need to be within 4.0 Å to the serine nucleophilic atom (O𝛾).
(2) Small molecule aromatic inhibitors (e.g. cinnamates, phenolates). These molecules are mobile and form other favorable interactions in the binding site in unreactive modes. For these reasons, all small molecule non-peptide-like inhibitors were excluded as GSAs.

**Text S3.** Knowledge-based energy functions.

Knowledge-based or statistical energy functions are derived from empirical databases of three-dimensional structures of molecules based on the standard formalism of Boltzmann statistics: the probability of observing a system in a certain state is (inversely) related to its energy:

$P_{i}$ = $\frac{e^{-E_{i}}/k_{B}T}{Z}$ (eq. 3.1).

where $P_{i}$ is the probability of observing state *i*, $E_{i}$ is the total energy of the microstate *i*, *k* is the Boltzmann factor, *T* is the temperature, and *Z* is the partition function ($Z = \sum_{j\in(0, n)} e^{-E_{j}/k_{B}T}$). This formalism describes a system at thermal equilibrium (a canonical ensemble); therefore, an assumption that underlie all knowledge-based energy functions is that a collection of experimental structures, while individually determined, resembles the physical ensemble of a system at thermal equilibrium.

Knowledge-based distributions and their derived energy functions quantitatively match those obtained from NMR experiments and QM calculations, supporting their ability to model ensembles of molecular interactions (*67*–*69*). They have been widely applied to determine pairwise atomic or residue distances (*47*, *157*, *158*) and to discriminate solvent accessibility (*159*), secondary structures (*160*), hydrogen bonding (*73*) and torsion angle (*161*) energetics. The accuracy and capabilities of crystallographic knowledge-based energy functions suggest that the assumption that they resemble a canonical ensemble holds. In other words, although each crystal favors a particular state, these preferences are randomized, and their combined effects do not significantly bias the overall ensemble distributions.

Below we describe the method we used to derive energy functions for molecular interactions (torsion angles, van der Waals, and hydrogen bonds) used in this work and the underlying assumptions and limitations.

1. **Boltzmann-like distribution.**

To derive a knowledge-based energy function for an interaction (e.g., the sidechain torsion angle of a serine residue, χ^1^), we searched for all crystallographic structures that contain the interaction of interest and collected their geometric parameter(s), in this case the serine χ^1^ angle. We defined a reference state with no conformational preferences, where all χ^1^ angles have equal probabilities, so that its energy $E_{ref}$ = 0. From eq. 1, we can calculate the difference in energy between a certain state where χ^1^ = *i* and the reference state:

$\Delta E =E_{i} - E_{ref} = -k_{B}T($ln$\frac{P_{i}}{P_{ref}} -$ln$\frac{Z_{i}}{Z_{ref}}) = -k_{B}T$ln$\frac{P_{i}}{P_{ref}} +$ constant (eq. 3.2).

where $E_{i}$ is the energy of state *i*, $E_{ref}$ is the energy of the reference state*,* $k_{B}$ is the Boltzmann constant, $P_{i}$ is the probability of observing state *i*, $P_{ref}$ is the probability of observing the reference state, $Z_{i}$ and $Z_{ref}$ is the partition function of state *i* and the reference state.

To obtain $P_{i}$, we equally split the distribution of χ^1^ to a number of bins (*n*_bins_) and calculate the probability of observing outcomes in each bin ($P_{i} = \frac{n_{i}}{n_{total}})$, the number of structures in bin *i* divided by the number of total structures), which gives a discrete probability function. A large *n*_bins_ and a large *n* allow this discrete function to approximate a smooth probability density function. Nevertheless, there may not be a sufficient number of unfavorable states present in the collected datasets. In such cases the energy function can be discontinuous or inaccurate in high energy regions, resulting in higher uncertainties or limits for these regions.

Since all $\chi_{1}$ values are equally probable in the reference state, $P_{ref} =\frac{1}{n_{bins}}$. The relative energy between each microstate and the reference state becomes:

$\Delta E = -k_{B}T$ln$\frac{P_{i}}{1/n_{bins}}$ (eq. 3.3).

2. **Bayesian Statistics.**

Bayesian statistics have been used as an alternative framework to establish the use of knowledge-based approaches to derive scoring functions for structural prediction (*162*, *163*). Here, we also considered the construction of knowledge-based energy functions from a Bayesian approach and show its mathematical equivalence to the Boltzmann statistics approach in section **1**. Following the standard Bayesian interpretation, the conditional probability of observing a molecular geometry *i* in a canonical ensemble *C* is:

$P(i|C)P(C) = P(C|i)P(i)$ (eq. 3.4).

where $P(i|C)$ is the probability of observing a geometric parameter *i* (e.g. a certain serine χ^1^ value), given microstates of a canonical ensemble *C* of a system (e.g. a free serine) under certain condition, $P(C)$ is the probability that any random structure is a member of the ensemble, $P(C|i)$ is the probability of observing a serine in the ensemble given that it has a χ^1^ = *i*, and $P(i)$ is the probability of observing χ^1^ = *i* with no chemical or ensemble constraints (in other words, the probability of observing *i* when no chemical forces are at play, which would be completely random). The energy function can be represented using $P(i|C)$ following eq. 3.1:

$E_{i} = -k_{B}T$ln$P(C|i) - k_{B}T$ln$Z = -k_{B}T$ln$\frac{P(i|C)P(C)}{P(i)} +$constant (eq. 3.5)

Similarly, with the assumption that our distribution collected from experimental data approximates a canonical ensemble, we equally separate our observations into a number of bins (*n_bins_*). $P(i|C)$ would then be the probability of observing state *i* in the distribution, or *P*_i_ as defined in section **1**. $P(C)$ is a constant; under the assumption that the collected structures resemble canonical ensemble *C,* $P(C)$ = 1. Lastly, $P(i)$ = $\frac{1}{n_{bins}}$, since there will be no preference for any atomic arrangement. Therefore, our expression reduces to:

$E_{i} = -k_{B}T$ln$\frac{P_{i}}{1/n_{bins}}$	+ constant (eq. 3.6).

To consider relative energies, we again determine the difference in energy for state *i* and the reference state as defined in section **1**:

${\Delta E =E}_{i} - E_{ref} = -k_{B}T$ln$\frac{P_{i}}{1/n_{bins}}$	(eq. 3.7).

which is the same as eq. 3.3.

**Text S4.** Limitations of Cartesian-based metrics (RMSD, B-factors, *etc.*) for structural comparisons.

Protein structural data is universally parametrized by atomic coordinates based on a cartesian system, and thus structural analyses often rely on metrics based on atomic coordinates. Root-mean-square deviation of atomic coordinates (RMSD) and related metrics (e.g. root-mean-square fluctuation) are commonly used to evaluate the similarities and differences between structures (*164*–*167*). In addition, B-factors describe harmonic deviations of the electron density distribution from the modeled coordinates and are widely used as a proxy for the rigidity or flexibility of atoms in structural comparisons (*168*).

While useful for many applications, cartesian-based metrics alone do not provide the information needed to evaluate individual atomic-level changes that are relevant to and responsible for the reaction and its energetics. All non-covalent conformational changes in proteins essentially arise from bond rotations (*169*, *170*), with the bond rotations coupled to changes in the geometries of their surrounding interactions such as hydrogen bonds and van der Waals interactions*.* Cartesian-based metrics reduce the complex, multidimensional conformational changes to a single scalar value describing displacements of atomic coordinates. Because of this disconnect between atomic displacements and the changes in fundamental chemical interactions, Cartesian-based metrics cannot be directly related to the energetics that define catalysis. We instead turned to torsion angles as they describe the essential bond rotations and overcome the above limitations.

**Text S5.** Calculation of general base catalysis and associated errors.

General base catalysis was calculated using Brønsted coefficients (β) and effective molarity (EM) values according to the Brønsted relationship (*171*):

$$Log10 \left( k \right)=\beta\times pK_{a}+\mathrm{constant}$$

where *k* is the second order rate constant of the amide hydrolysis reaction and the p*K*_a_ refers to that of the conjugate acid of the base catalyst. Therefore,

$rate enhancement=\frac{k_{enz}[\mathrm{His}]}{k_{soln}[H_{2}O]}=\frac{{10}^{\beta(pK_{a}\left[ His-H^{+} \right])}[\mathrm{His}]}{{10}^{\beta(pK_{a}\left[ H_{3}O^{+} \right] )}[H_{2}O]}$.

The Brønsted β value corresponds to the extent of proton transfer in the transition state. A β < 0.5 indicates an early transition state where the proton is mostly associated with the acid; conversely, β > 0.5 indicates a late transition state where the proton transfer is mostly complete. Our QM calculation of the solution reaction as well as prior QM/MM on serine proteases (*84*) suggests a late transition state of the acylation reaction resembling the tetrahedral intermediate. Therefore, we used a range of β from 0.5 to 1 for our calculation. The p*K*_a_ value of the catalytic triad histidine was determined to be ~7 (*172*, *173*). Using the same Brønsted β values for the enzyme and solution reaction assumes that the reaction in water goes through the same reaction path where the proton transfer happens. The EM of the catalytic histidine ([His]) was estimated to be ~10 M. Across a range of intramolecular base catalysis for hydrolysis and aminolysis reactions, the highest EM was determined to be 80 M, and the great majority are less than 10 M, with an overall low sensitivity to the molecular structure of the reactants (*62*–*64*, *174*). Therefore, we used EM = 10 M as a first approximation and also used 1 to 100 M to calculate the range of effects. The p*K*_a_ of hydronium ion is –2 and [H_2_O] is 55 M in the solution reaction. The above values give:

$$rate enhancement=\frac{{10}^{0.75\times7}\times10 M}{{10}^{0.75 (-2)}\times55 M}=1.0\times{10}^{6}$$

corresponding to 8.2 ± 3.1 kcal/mol at *T* = 298 K (table S11).

In principle, one could obtain a more accurate EM by simulating imidazole (the general base) and ethanol (the nucleophile) in aqueous phase, determining the fraction of time where an imidazole is aligned to abstract a proton from ethanol, and comparing this fraction to that in the enzyme ground state ensembles. However, we are unable to perform this analysis due to current limitations in modeling and computational power, as we would need to define the reactivity for each imidazole•ethanol configuration while considering their interactions with surrounding water molecules. This calculation involves obtaining a multi-dimensional potential energy surface via quantum mechanical treatments, as is needed due to the quantum character of proton transfer and hydrogen bonds.

While there are considerable errors associated with the calculation of general base catalysis, it is apparent that it alone cannot account for the rate enhancements provided by serine proteases. Even using overestimates EM = 100 M and β = 1 give 12.6 kcal/mol from general base catalysis, insufficient to account for the 17.1 kcal/mol catalysis observed for trypsin (table S2 and 11).

**Text S6.** Oxyanion hole hydrogen bonds contribute to catalysis via ground state destabilization.

*Systematic assessment of oxyanion hole hydrogen bond geometry in serine proteases*

Hydrogen bonds donated to sp^2^ hybridized groups such as carbonyls prefer to orient themselves towards the lone pair (𝛼_HB_ = 120° and $\phi$_HB_ = 0°, as defined in fig. S30) (*46*, *175*). Contrary to this preference, a previous study by Simón and Goodman (*93*) found that enzymatic oxyanion hole hydrogen bonds are generally out of the plane of the carbonyl group of the bound ligand. The suboptimal hydrogen bond geometry is expected to destabilize the ground state but not the transition state, where the sp^3^ oxyanion has a largely symmetric electron density (see below). Theoretical calculation of simple model compounds mimicking the enzymatic oxyanion hole structure gave a 2.4 kcal/mol catalytic advantage from the ground state destabilization of the two hydrogen bonds (*93*). Nevertheless, this calculation, performed using two fixed water molecules as hydrogen bond donors and in the gas phase, may overestimate the catalytic contribution (*176*). Our pseudo-ensemble analyses and knowledge-based energy functions strongly support catalysis via this ground state destabilization mechanism and provided a similar but somewhat smaller estimate for its energetic contribution (1.7 to 1.9 kcal/mol) to catalysis.

The dataset used previously in Simón and Goodman (*93*) consists of PDB structures from multiple enzymes where an oxyanion hole is bound to a carbonyl ligand (*93*). Analysis of this database found that (1) some of the carbonyl ligands are not expected to be ground state analogs, and (2) a small number of enzymes are over-represented, skewing the distributions of hydrogen bond geometries to possibly represent a small subset of enzymes rather than a general phenomenon.

To overcome these limitations, we curated serine proteases so that each enzyme can be examined separately, and we ensured that GSA-bound structures mimic the planar reaction ground state geometry (supplementary text S2). We observed out-of-plane orientations for all 10 serine proteases for which we found GSA-bound structures, with $\phi$_HB_ = 76° (*s.d.* = 10°) and 78° (*s.d.* = 11°) for the 1° and 2° hydrogen bond, respectively (1° and 2° defined in Fig. 1A; table S15). These serine proteases represent four structural clans (fig. S31, color-coded). Further, the distributions of $\phi$_HB_ in our GSA-bound pseudo-ensembles are narrow, with no in-plane population observed (fig. S31), suggesting a robust out-of-plane geometry and an inability to conformationally relax to avoid the ground state destabilization without a substantial energetic penalty. In contrast and as expected, we found that small molecule amide•carbonyl hydrogen bonds prefer to be in-plane ($\phi$_HB_ = 5°, *s.d.* = 27°) (table S15 and fig. S31). Our results strongly support the experimental findings of Simón and Goodman (*93*).

For the oxyanion hole ground state destabilization to contribute to catalysis, the hydrogen bonds must be less destabilized (or not destabilized) in the transition state. We therefore also compared hydrogen bond geometries in TSA-bound pseudo-ensembles to the analogous small molecule structures with amide groups donating hydrogen bonds to sp^3^ hybridized oxygens attached to a carbon atom, again using a knowledge-based approach. No preferences were observed for $\phi$_HB_ in the small molecule distribution, suggesting that the hydrogen bond orientational destabilization is relieved in the transition state (fig. S32). This observation is consistent with the trends observed in the calculations performed by Simón and Goodman (*93*).

*Calculation of the catalytic contribution from oxyanion hole destabilization*

To estimate catalytic contribution from ground state destabilization of the oxyanion hole hydrogen bond, we constructed knowledge-based energy functions for $\phi$_HB_ from small molecule amide•carbonyl hydrogen bonds (sp^2^; resembling GS) and amide hydrogen bonds to sp^3^ oxygen atoms (resembling TS) (Fig. 5F and fig. S33). As described in the main text, the suboptimal ground state hydrogen bond orientation ($\phi$_HB_) and the absence of any orientational destabilization in the sp^3^–like transition state gave ΔΔG^‡^ (= ΔG_GS_ – ΔG_TS_) = 0.94 and 0.95 kcal/mol for the 1° and 2° oxyanion hydrogen bond, with small standard deviations on the scale of ~0.05 kcal/mol (table S16C).

This energetic effect is the same across 10 serine proteases from four structural clans (1.7 to 1.9 kcal/mol for the two hydrogen bonds combined) (fig. S33 and table S16C). This quantitative consistency across enzymes and folds is unlikely to arise from chance, as it would be highly unlikely for artifactual configurations to arise uniformly in all the different enzymes and folds. The commonality of this catalytic strategy appears to originate from a common catalytic motif, the nucleophilic elbow, as described in the main text (***The nucleophilic elbow underlies ground state destabilization of the oxyanion hole hydrogen bonds***).

*Minimal energetic effects from changes in oxyanion hole hydrogen bond lengths (d_HB_) and angles (𝛼_HB_)*

To comprehensively assess the energetic changes of the oxyanion hole hydrogen bonds in going from the ground state to the transition state, we performed the comparisons as described above for the other two hydrogen bond parameters, the length *d*_HB_ and the angle 𝛼_HB_ (defined in fig. S30). Our results suggest little or no catalytic effects from changes in these two geometric parameters.

For trypsin (with the largest *n* and therefore highest confidence), small differences in *d*_HB_ were found for the ground state, where the 1° hydrogen bonds are the same as those found in small molecules (2.9 Å) and the 2° hydrogen bonds are slightly shorter (2.7 Å *vs.* 2.9 Å) (fig. S31 and table S15A). Coupling in lengths for a pair of hydrogen bonds donated to the same acceptor is expected on this scale (|Δ*d*_HB_| ~ 0.05 to 0.2 Å) (*177*). The TSA-bound state has the same 1° and 2° hydrogen bond lengths (2.9 Å and 2.7 Å, respectively, fig. S32 and table S15B). The ground state *versus* transition state differences yield a minimal net energetic difference of 0.2 kcal/mol for the two hydrogen bonds together (table S16A and fig. S34). Other serine proteases (with smaller numbers of structures) gave a wide range of net differences ranging from ΔΔG^‡^ = –2.8 to +2.4 kcal/mol (table S16A and fig. S34). The large deviations (*s.d*. = 0.9–2.3 kcal/mol) associated with these energy values and the differences across serine proteases do not allow conclusions to be drawn about potential catalytic effects from changes in hydrogen bond lengths for these serine proteases.

The hydrogen bond angles (𝛼_HB_, defined in fig. S30) in enzyme oxyanion holes match the optimal angle observed in the small molecule distribution for the GSA-bound state (fig. S31 and table S15A); for the TSA-bound states, the 𝛼_HB_ distributions deviate from the optimal angle by 15 – 20° (fig. S32 and table S15B). Knowledge-based energy functions for 𝛼_HB_ revealed a shallow minimum (fig. S35), resulting in little energetic consequence from this deviation. For trypsin, we found ΔΔG^‡^ values of –0.3 ± 0.5 and –0.4 ± 0.5 kcal/mol for its 1° and 2° hydrogen bonds, and for other serine proteases a range of ΔΔG^‡^ from –0.8 to 0.2 kcal/mol were found (table S16B). Similar to *d*_HB_, these ΔΔG^‡^ values are associated with large variations (larger than the differences between the GSA-bound and TSA-bound states) and are not consistent across serine proteases, presumably resulting from the small sample sizes (table S16B).

*Assessing potential catalytic contribution from reduced conformational entropy*

Oxyanion hole hydrogen bonds could provide additional catalytic advantage from increased positioning compared to the reaction in solution. MD simulation suggests a very small entropic contribution of ~0.2 kcal/mol from these hydrogen bonds. Our simulation for the substrate analog NMA in water showed that, on average, there are 1.8 water molecules hydrogen bonded to the amide carbonyl, a small difference compared to the two oxyanion hole hydrogen bonds on the enzyme. The catalytic advantage from reduced positioning is:

$$-T\Delta\Delta S_{\mathrm{conf}}=T\left( \Delta S_{\mathrm{soln}}^{\ddagger}- {\Delta S}_{\mathrm{enz}}^{\ddagger} \right)$$

Since the enzyme has two hydrogen bonds formed throughout the reaction, ${\Delta S}_{\mathrm{enz}}^{\ddagger}=0$. The solution term can be estimated considering the probability (1.8 out of 2) where the hydrogen bonds are formed in the ground state and assuming the hydrogen bonds are always formed in the transition state ($S_{\mathrm{soln}}^{\mathrm{TS}}=0$):

$T\Delta S_{\mathrm{soln}}^{\ddagger}$ = $T{(S}_{\mathrm{soln}}^{\mathrm{GS}}- S_{\mathrm{soln}}^{\mathrm{TS}})$= $TS_{\mathrm{soln}}^{\mathrm{GS}}=$ $k_{B}T\left( \sum_{i} P_{i}\ln P_{i} \right)$ = $k_{B}T[\left( \frac{1.8}{2}\ln\left( \frac{1.8}{2} \right)+\frac{0.2}{2}\ln\left( \frac{0.2}{2} \right) \right)$

$$= 0.19\frac{kcal}{mol} (at T=298 K)$$

Additional catalysis may result from a more restricted hydrogen bonded state in the enzymatic ground state than that in solution. Nevertheless, for this entropic difference to contribute to catalysis, the hydrogen bonds in the transition state must be more restricted than those in the ground state (for both the solution and enzymatic reaction). In contrast to the positioning between reactants where in the transition state a partial bond is formed and a higher level of positioning is expected, we do not know whether the oxyanion hole hydrogen bonds become more restricted in the transition state *a priori*. While these hydrogen bonds are stronger in the transition state as partial charge accumulates on the oxygen, the ground state orientational restrictions are relieved in the transition state (Fig. 5F and fig. S33, sp^2^ *vs.* sp^3^). Therefore, we do not expect significant catalytic contribution from reduced conformational entropy of the oxyanion hole hydrogen bonds.

**Text S7.** Evaluating catalytic contributions from increased catalytic triad His•Asp hydrogen bond strength in going from the ground to the transition state.

The catalytic triad histidine becomes protonated during the reaction. As positive charge accumulates on the histidine from the ground state to the transition state, the His•Asp hydrogen bond becomes stronger. This increase in hydrogen bond strength is expected to be larger for the enzymatic His•Asp hydrogen bond than an imidazole•water hydrogen bond in the base-catalyzed solution reaction, as the aspartate has a higher electron density than water and thus is a stronger hydrogen bond acceptor (*96*–*98*). Catalysis from the His•Asp hydrogen bond derives from the difference in the amount of hydrogen bond strengthening during the reaction between the enzymatic and solution reaction.

We used the empirical linear free energy relationship known as the Hine equation (*96*–*98*) to quantify this energetic difference. As outlined in the thermodynamic cycle in fig. S36, the rate enhancement from the enzymatic His•Asp hydrogen bond ($\Delta\Delta G_{His\bullet Asp}^{\ddagger}$) is defined as:

$\Delta\Delta G_{His\bullet Asp}^{\ddagger}= \Delta G_{His\bullet Asp}^{\ddagger}- \Delta G_{sol}^{\ddagger}= -k_{B}T\times ln\left( \frac{K_{His\bullet Asp}}{K_{sol}} \right) = -k_{B}T\times ln(\frac{K_{HB, TS}^{f}}{K_{HB, GS}^{f}})$.

where$K_{His\bullet Asp}$ is a pseudo-equilibrium constant representing the change in hydrogen bond energetics in course of reaction for the His•Asp hydrogen bond on the enzyme; $K_{sol}$ is the corresponding term for aqueous solution; $K_{HB, GS}^{f}$ and $K_{HB, TS}^{f}$ are the pseudo-equilibrium constants for His•Asp hydrogen bond formation in the ground state and in the transition state, respectively.

The Hine equation describes the sensitivity of hydrogen bond strength in response to the charge densities of the hydrogen bonding partners, typically approximated by p*K*_a_ values (*31*, *33*, *178*):

$$logK_{HB}^{f}= \tau\left( pK_{a}(HOH)- pK_{a}(HA \right))\left( pK_{a}\left( BH^{+} \right)- pK_{a}\left( {[H}_{3}{O]}^{+} \right) \right)$$

$-log(2\times55M)$.

where HA is the hydrogen bond donor, BH^+^ is the conjugate acid of the hydrogen bond acceptor and $K_{HB}^{f}$ is the equilibrium constant for hydrogen bond formation between HA and B. Based on the Hine equation, the rate enhancement provided by the His•Asp hydrogen bond is:

$$log \frac{K_{HB, TS}^{f}}{K_{HB, GS}^{f}} = \tau\left[ \left( pK_{a}(HOH)- pK_{a}\left( [His-N^{\delta}{H]}^{+} \right) \right)\left( pK_{a}\left( Asp-OH \right)- pK_{a}({[H}_{3}{O]}^{+}) \right)- \left( pK_{a}(HOH)- pK_{a}(His-N^{\delta}H \right)\left( pK_{a}\left( Asp-OH \right)- pK_{a}\left( {[H}_{3}{O]}^{+} \right) \right) \right]$$

where the $\tau$ value is a constant that describes the sensitivity of hydrogen bond strength in response to the change in charge densities of the hydrogen bonding partners.

As the enzyme active site differs from the aqueous solution, we cannot *a priori* predict its exact $\tau$ value (*98*). Nevertheless, prior studies revealed similar energetic behavior of solution hydrogen bonds and active site hydrogen bonds of a model enzyme, ketosteroid isomerase (*179*, *180*). These studies found a shallow (but significant) dependence of hydrogen bond strength and donor/acceptor p*K*_a_ values on the enzyme, similar to the relationship in solution. A similarity for proteases is also reasonable given their exposed active sites that solvent molecules can readily access.

Based on the observations above, we used a $\tau$ value of 0.013 measured for aqueous solution in our calculation (*102*–*105*), which gives:

$$log \frac{K_{HB, TS}^{f}}{K_{HB, GS}^{f}} =0.013 \times\left[ \left( 16-7 \right)\left( 4-(-2) \right)-\left( 16-14 \right)\left( 4-(-2) \right) \right]=0.55$$

$$\Delta\Delta G_{His\bullet Asp}^{\ddagger}= -k_{B}T\times log\left( \frac{K_{HB, TS}^{f}}{K_{HB, GS}^{f}} \right)\times\ln\left( 10 \right)= -0.592\frac{kcal}{mol} \times0.55 \times2.30=-0.75 \frac{kcal}{mol}$$

While we found no indication for an active site environment that is substantially different from solution, we cannot rule out the possibility that the active site provide a higher $\tau$ value than that in water that enhance the contribution from the His•Asp hydrogen bond. A range of $\tau$ values from 0.013 to 0.033 gives $\Delta\Delta G_{His\bullet Asp}^{\ddagger}$ from –0.75 to –1.90 kcal/mol so that a modest catalytic contribution from the strengthening of His•Asp hydrogen bond is expected even if $\tau$ is substantially perturbed.

**Text S8.** Evaluating potential catalytic contributions from geometric properties of the catalytic triad hydrogen bonds.

Above, we evaluated the catalytic contributions from general base catalysis and from the strengthening of a hydrogen bond (the His•Asp hydrogen bond), chemical features that are not present in the solution reaction. Geometric properties of the catalytic triad hydrogen bonds can provide additional catalysis if they stabilize the transition state more than the ground state, as occurs in the oxyanion hole (see main text *Catalysis from ground state destabilization of oxyanion hole hydrogen bonds*). To assess this possibility, we compared geometric distributions of the catalytic triad hydrogen bonds in serine protease pseudo-ensembles to analogous small molecule hydrogen bonds. As described below, these comparisons provide no evidence for special geometric properties of these hydrogen bonds that would provide additional catalysis.

*The His*•*Asp hydrogen bond*

The distributions of His•Asp hydrogen bond geometric parameters (defined in fig. S37) in most serine proteases match the optimal values of the knowledge-based distributions of small molecule imidazole•carboxylate hydrogen bonds, and the geometries do not change from the GSA- to the TSA-bound states (fig. S38 to 39 and table S17). Correspondingly, knowledge-based energy functions derived from small molecule distributions suggest little or no catalytic effects from hydrogen bond geometries (on the order of <0.5 kcal/mol, table S18).

Literature and biochemistry textbooks emphasize potential catalytic contribution from a short His•Asp hydrogen bond (supplementary text S1). However, the His•Asp hydrogen bond length is expected to be shortened as the histidine becomes protonated in going from the ground to the transition state, and there is no indication that its length is shorter-than-normal in the transition state. The His•Asp lengths were found to be 2.71 Å (*s.d.* = 0.11 Å) in the TSA-bound pseudo-ensembles, matching the preferred length found in small molecules (2.69 Å, *s.d.* = 0.14 Å) and in proteins in the PDB (2.73 Å, *s.d.* = 0.39 Å) (table S17A).

Crystallographic distributions of hydrogen bond lengths revealed the expected change in His•Asp length in going from the ground to the transition state. Protonated imidazole•corboxylate hydrogen bonds in small molecules are 0.06 Å (*s.d.* = 0.18 Å) shorter than those with neutral imidazoles, matching the change from the ground to the transition state obtained in our QM calculation. There is a similar difference in His•Asp lengths comparing the GSA- and TSA-bound serine proteases (0.08 Å for trypsin; table S17A), although the error in atomic coordinates is significant on this scale and the change was not statistically significant. Nevertheless, when only high resolution (<1.5Å) structures were included in the comparisons, the same change was observed and is statistically significant (see *Assessing potential sources of variations in hydrogen bond lengths in pseudo-ensemble comparisons.*).

*The His•Ser hydrogen bond*

The same analyses performed for the His•Ser hydrogen bond in the catalytic triad revealed differences in hydrogen bond lengths (*d*_HB_) and angles (𝛼_HB_) on serine proteases *versus* in small molecules (fig. S40 to 42 and table S19 to 20). In the GSA-bound state, the His•Ser hydrogen bond length is 0.1 Å longer than the optimal length predicted by small molecule (neutral) imidazole•hydroxyl hydrogen bonds (2.67 Å observed *vs.* 2.77 predicted), and it lengthens by 0.2 Å in going to the TSA-bound state, such that it becomes 0.1 Å longer than the optimum predicted by small molecule (protonated) imidazole•hydroxyl hydrogen bonds (2.92 Å observed *vs.* 2.81 Å predicted, table S19A). Little or no energetic difference result from this change, as the lengths are suboptimal in both the GSA- and the TSA-bound states, (<0.1 kcal/mol, fig. S42 and table S20). Similarly, the 𝛼_HB_ angles deviate from the predicted optimal values to the same extent in the GSA- and TSA-bound states, giving negligible energetic differences (fig. S42 and table S20).

The increase in the His•Ser hydrogen bond length appears to result from the sidechain rotation of the catalytic serine while fixing the histidine in place (fig. S43). The inability for the catalytic histidine to move and relieve to the optimal His•Ser length provides additional support for a highly positioned active site that prioritizes motion along the reaction path (i.e. the serine rotation) while limiting other motions.

*Assessing potential sources of variations in hydrogen bond lengths in pseudo-ensemble comparisons.*

The ability of pseudo-ensemble comparisons to resolve geometric changes relies on the amount and quality of the structural data. Subtle changes on the scale of <0.1 Å, such as for hydrogen bond lengths, may be obscured by the heterogeneity of structures included in the ensembles and by the errors in the structural models. Therefore, we assessed whether these factors caused systematic errors on the observed pseudo-ensemble geometries. Specifically, we assessed three potential sources of variations that may affect hydrogen bond lengths: (1) errors in atomic coordinates from X-ray crystallographic structural models, (2) different crystallographic pH values that may affect residue protonation states, and (3) differences between different types of bound analogs. We performed these analyses for the His•Asp hydrogen bond as data from NMR experiments are available for comparisons. Overall, we found that pseudo-ensembles from high-resolution structures (<1.5 Å) capture subtle geometric changes (<0.1 Å) better than those built from lower resolution structures and that the different chemical nature of TSAs results in systematic differences on the His•Asp lengths, consistent with NMR data.

(1) Comparisons of high-resolution structures (<1.5 Å) gave significant differences of 0.05 – 0.1 Å for the His•Asp *d*_HB_ between the GSA and TSA-bound states (*p* = 0.03 for trypsin; *p* = 0.02 when 12 serine proteases with available data were combined; fig. S44), matching differences predicted by small molecule structures and from our QM calculations (0.06 Å). This observation is consistent with a shortened His•Asp hydrogen bond upon histidine protonation (as expected based on small molecule data; table S17A). This and other small differences may be masked by model errors in lower resolution crystallographic data.

(2) Different protonation states of the histidine are expected to give different His•Asp hydrogen bond lengths. Prior NMR data showed that the histidine protonation state is pH-independent in the GSA-bound and TSA-bound states (*181*–*184*). Consistent with the NMR results, there are no pH-dependent differences in His•Asp lengths for the GSA-bound ensembles (fig. S45). An apparent increase in His•Asp length was observed for higher crystallization pH in the TSA-bound ensembles, but the variance at each pH is large and this trend could be artifactual due to the aggregation of data from multiple proteases and difference types of TSAs [see (3)], as there is no apparent relationship between length and pH within the same TSA class (for the limited data for each TSA class across pH (fig. S45). Therefore, it is unlikely that crystallographic pH systematically affects the His•Asp hydrogen bond length.

(3) TSAs differ from one another in their electronic properties, and those with nonpolar *versus* polar substituents are expected to interact with the catalytic histidine differently (fig. S46A). Bound TSAs containing polar substituents (e.g. boronic acids, phosphonates and sulfonates) attract the histidine on one side (Nε), thus lengthening the hydrogen bonding distance between its nitrogen on the other side (Nδ) and the aspartate; non-polar TSA (e.g. aldehydes and various ketones) do not exhibit this effect (fig. S46B). Notably, this observation is consistent with prior NMR results that the trifluromethyl ketone-bound chymotrypsin, compared to boronic acid-bound, gives a more downfield chemical shift for the catalytic triad histidine (*181*, *184*, *185*). The agreement between pseudo-ensembles and NMR measurements supports the ability of crystallography-based pseudo-ensembles to capture subtle geometric differences that occur in the solution phase.

**Text S9.** Energetic additivity of identified catalytic features.

Catalytic contributions are expected to be independent and additive for the catalytic features described in the main text, which are (1) general base catalysis, (2) the catalytic triad His•Asp hydrogen bond, (3) positioning of the reactants, and (4) ground state destabilization of the serine rotamer, (5) the reactant distance and (6) the oxyanion hole hydrogen bonds.

Contributions from the His general base catalysis (1) and the His•Asp hydrogen bond (2) in the catalytic triad are derived by separately evaluating the chemical properties of these interactions. Coupling effects seen in protein hydrogen bond networks are small—and considerably smaller than the uncertainties associated with the empirical LFERs (*129*, *136*, *177*).

Catalytic contribution from the positioning of the reactants (3) is independent from all other energy calculations. It is derived from the spread of the ensemble distributions, distinct from the other energy terms derived by mapping the observed geometries on knowledge-based energy functions.

Catalytic contributions from ground state destabilization mechanisms [(4) to (6)] are also expected to be independent, as each calculation is based on the geometry of an individual chemical interaction and a knowledge-based energy function for that specific interaction. We evaluated the correlation between these energy terms to determine whether they are independent of one another, and we found minimal coupled effects, as follows.

A two-dimensional knowledge-based distribution collected for serine residues in proximity to a carbonyl group revealed little or no coupling between serine χ^1^ angles and the O•C distance that would affect our calculations (fig. S47, built from 162,706 interactions collected from 13,149 high quality PDB structures). The global preference for rotameric state (*gauche–, trans, gauche+*) slightly shifts at different distances, but the same local minima are preferred in each well. Therefore, the energetic destabilization from the serine rotamer *within* the reactive rotameric state (4) and from the nucleophile•electrophile distance (5) are expected to be independent and additive.

We do not have sufficient data to systematically evaluate the coupling between the three oxyanion hole hydrogen bond parameters (*d*_HB_, 𝛼_HB_ and $\phi$_HB_; defined in fig. S30). As we only found an energetic difference for one parameter ($\phi$_HB_) from the ground to the transition state, we only considered this term in our minimal model (supplementary text S6).

**Text S10.** Mechanistic and structural considerations for the use of nucleophilic elbow in enzyme catalysis.

Nucleophilic elbows perform the same catalytic function when they are used in different enzymes performing nucleophilic addition to carbonyl reactions, as they showed the same catalytic conformational features (main text, *Nucleophilic elbow-bearing enzymes from multiple folds exploit common catalytic features*). Conversely, nucleophilic elbows are not expected to be present in enzymes where structural or mechanistic constraints prevent the formation and catalytic use of its pseudo-ring structure, in particular in the following scenarios: (1) use of nucleophilic residue with more than one rotatable sidechains (e.g. His, Glu); (2) formation of an oxyanion that is multiple covalent bonds from the electrophile instead of adjacent, as occurs in nucleophilic aromatic substitution; (3) use of the backbone nitrogen of the nucleophilic residue as a general base instead of a hydrogen bond donor, an alternative strategy used by N-terminal proteases where the nitrogen is an amine instead of an amide; or (4) nucleophile attack internally at a neighboring residue such that the amide group of the neighbor cannot (also) be used for hydrogen bonding.

Eighteen of the 23 enzymes that do not use a nucleophilic elbow follow these criteria (table S27), whereas no nucleophilic elbow-bearing enzymes have these mechanistic features. The remaining five enzymes have limited structural and mechanistic data so we cannot evaluate potential mechanistic reasons for their lacking a nucleophilic elbow. Two of these enzymes use stronger hydrogen bond donors (Lys, Arg), suggesting possible alternative strategies to achieve catalysis from an oxyanion hole.

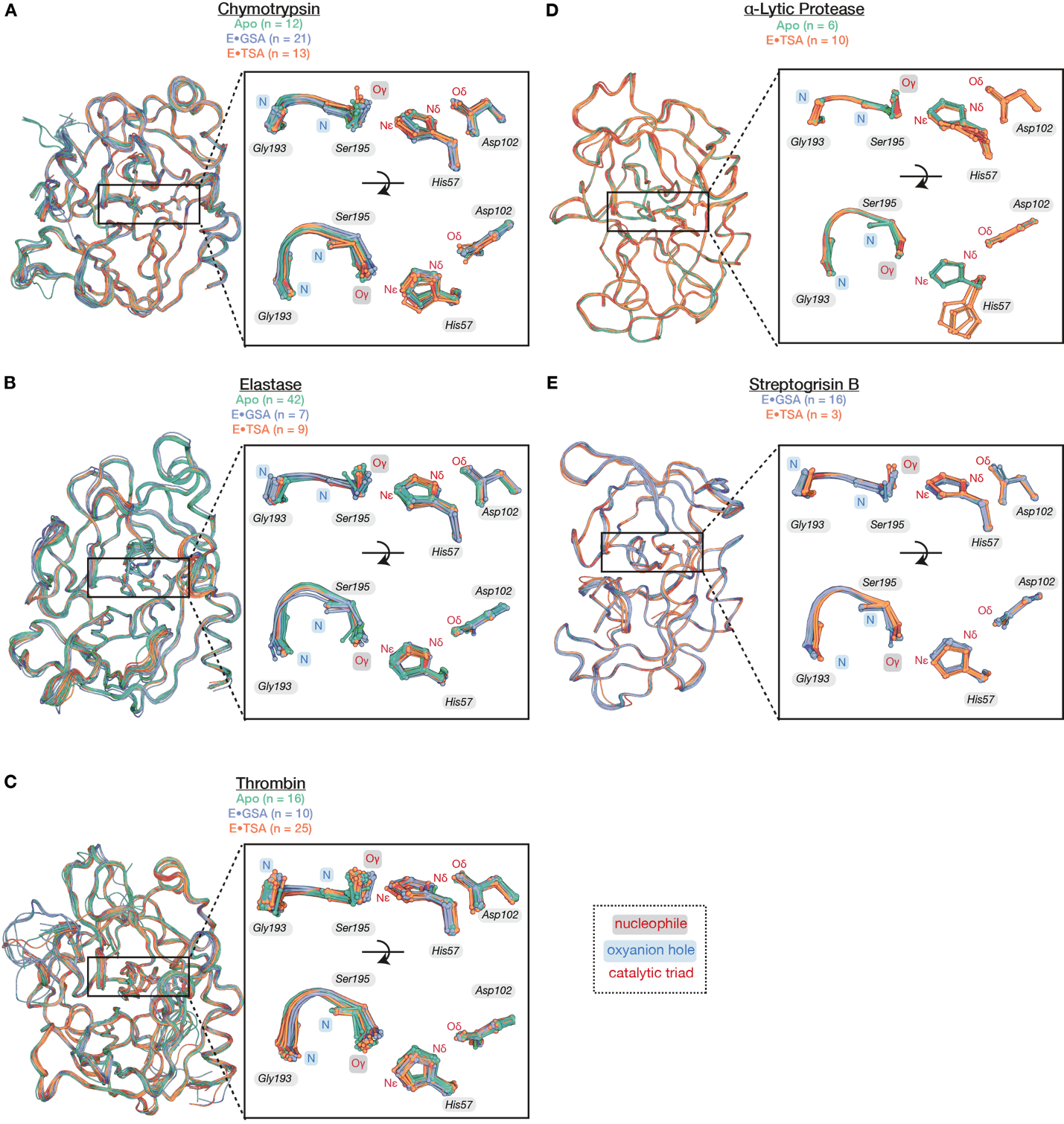
**Fig. S1.** Pseudo-ensembles of serine proteases in clan PA (except for trypsin, shown in Fig. 1C): chymotrypsin **(A)**, elastase **(B)**, thrombin **(C)**, α-Lytic protease **(D)**, and streptogrisin B **(E)**. Structures were aligned on the C𝛼 atoms of the protease chains using the highest resolution structure as the reference PDB (table S3). Labeled residue numbers are taken from the reference structure; all clan PA serine proteases here follow the trypsin numbering scheme. Labeled atom names follow the PDB format.

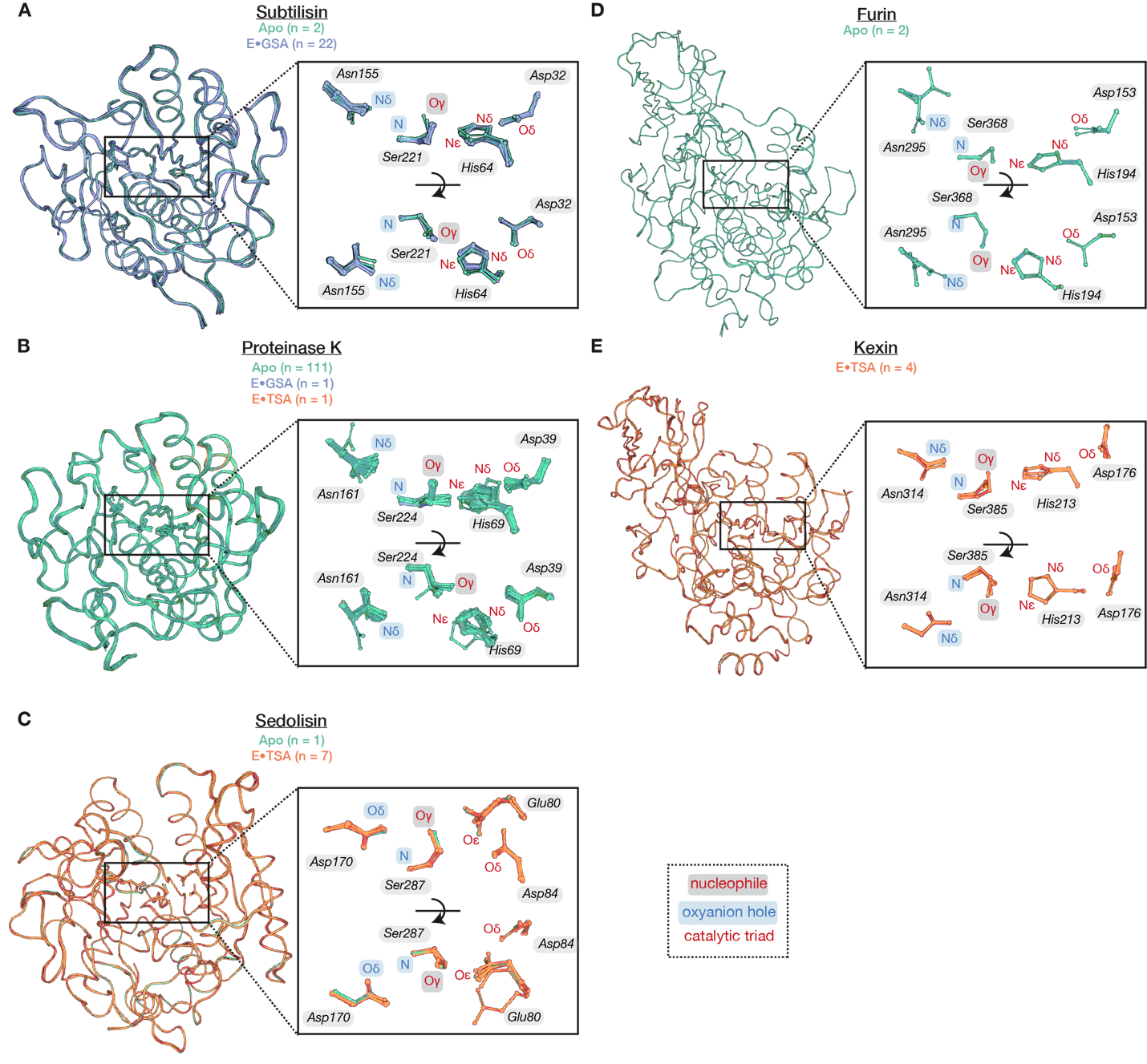

**Fig. S2.** Pseudo-ensembles of serine proteases in clan SB, **(A)** subtilisin, **(B)** proteinase K, **(C)** sedolisin, **(D)** furin and **(E)** kexin. Structures were aligned on the C𝛼 atoms of the protease chains using the highest resolution structure as the reference PDB (table S3). Labeled residue numbers are taken from the reference structure. Labeled atom names follow the PDB format.

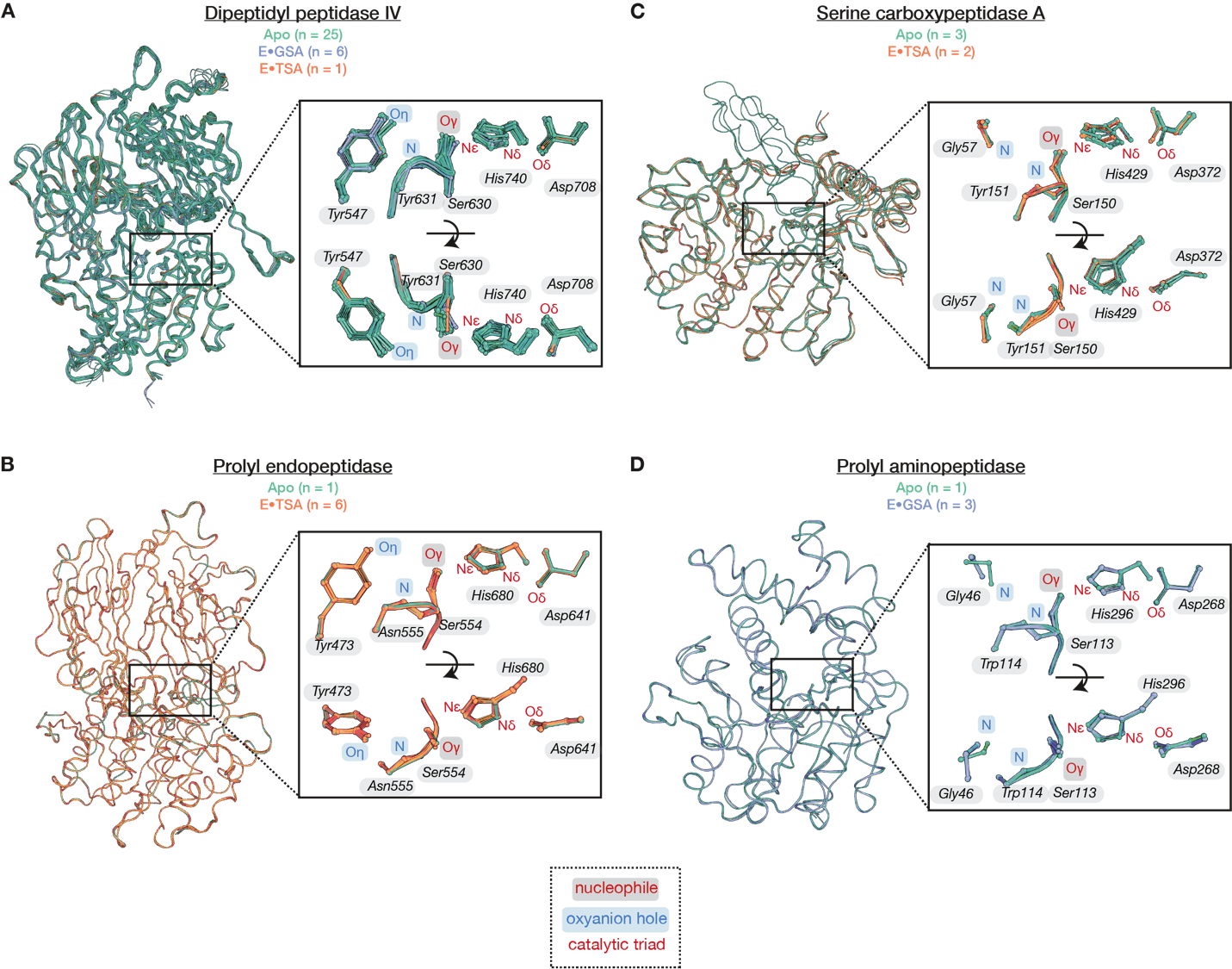
**Fig. S3.** Pseudo-ensembles of serine proteases in clan SC, **(A)** dipeptidyl peptidase IV, **(B)** prolyl endopeptidase, **(C)** serine carboxy peptidase A and **(D)** Prolyl aminopeptidase. Structures were aligned on the C𝛼 atoms of the protease chains using the highest resolution structure as the reference PDB (table S3). Labeled residue numbers are taken from the reference structure. Labeled atom names follow the PDB format.

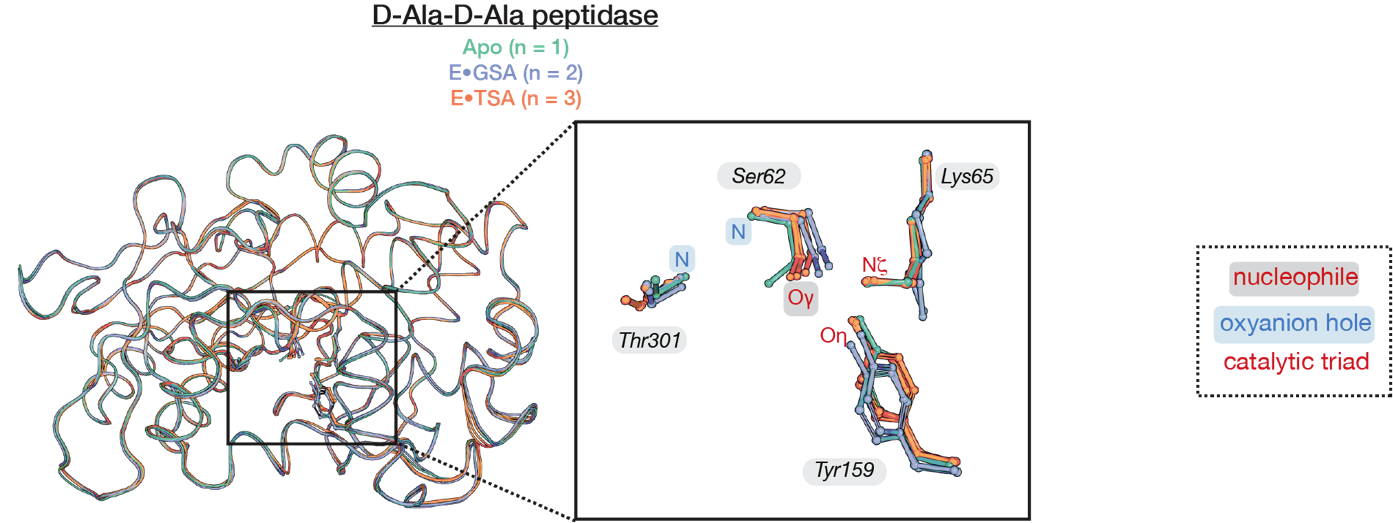
**Fig. S4.** Pseudo-ensemble of D-Ala-D-Ala peptidase, a serine protease in clan SE. Structures were aligned on the C𝛼 atoms of the protease chains using the highest resolution structure as the reference PDB (table S3). Labeled residue numbers are taken from the reference structure. Labeled atom names follow the PDB format.

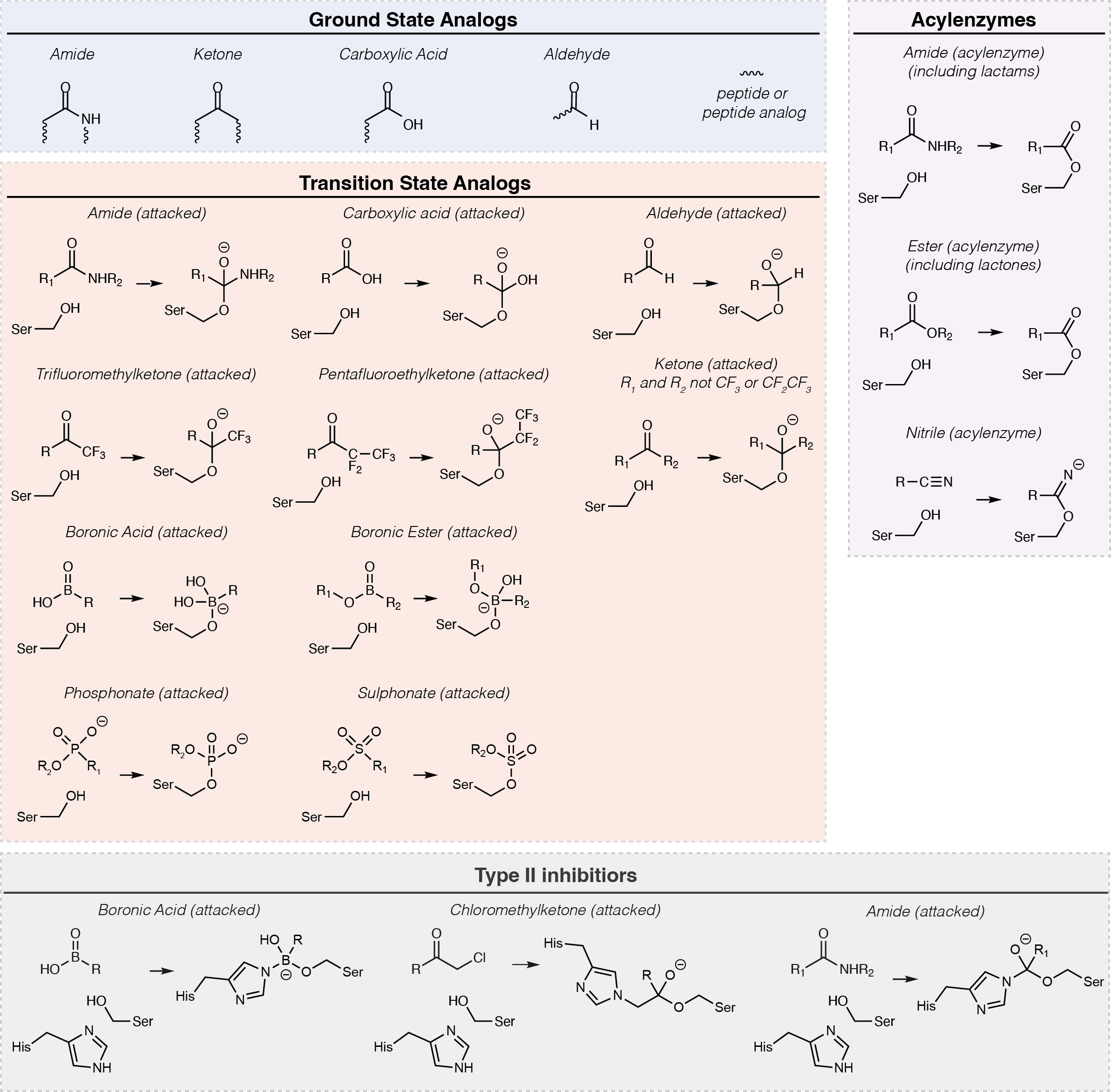

**Fig. S5.** Classification of serine protease ligands by their chemical groups. Ground state analogs (GSAs) are peptide or peptide analogs. Transition state analogs (TSAs) are compounds that react with the catalytic serine and form a stable covalent tetrahedral intermediate. Type II inhibitors include ligands that form a covalent bond with the general base histidine and were not included as transition state analogs. The acylenzyme class contains ligands covalently attached to the catalytic serine with a planar geometry. (See methods for detailed description of the criteria used for ligand classification.)

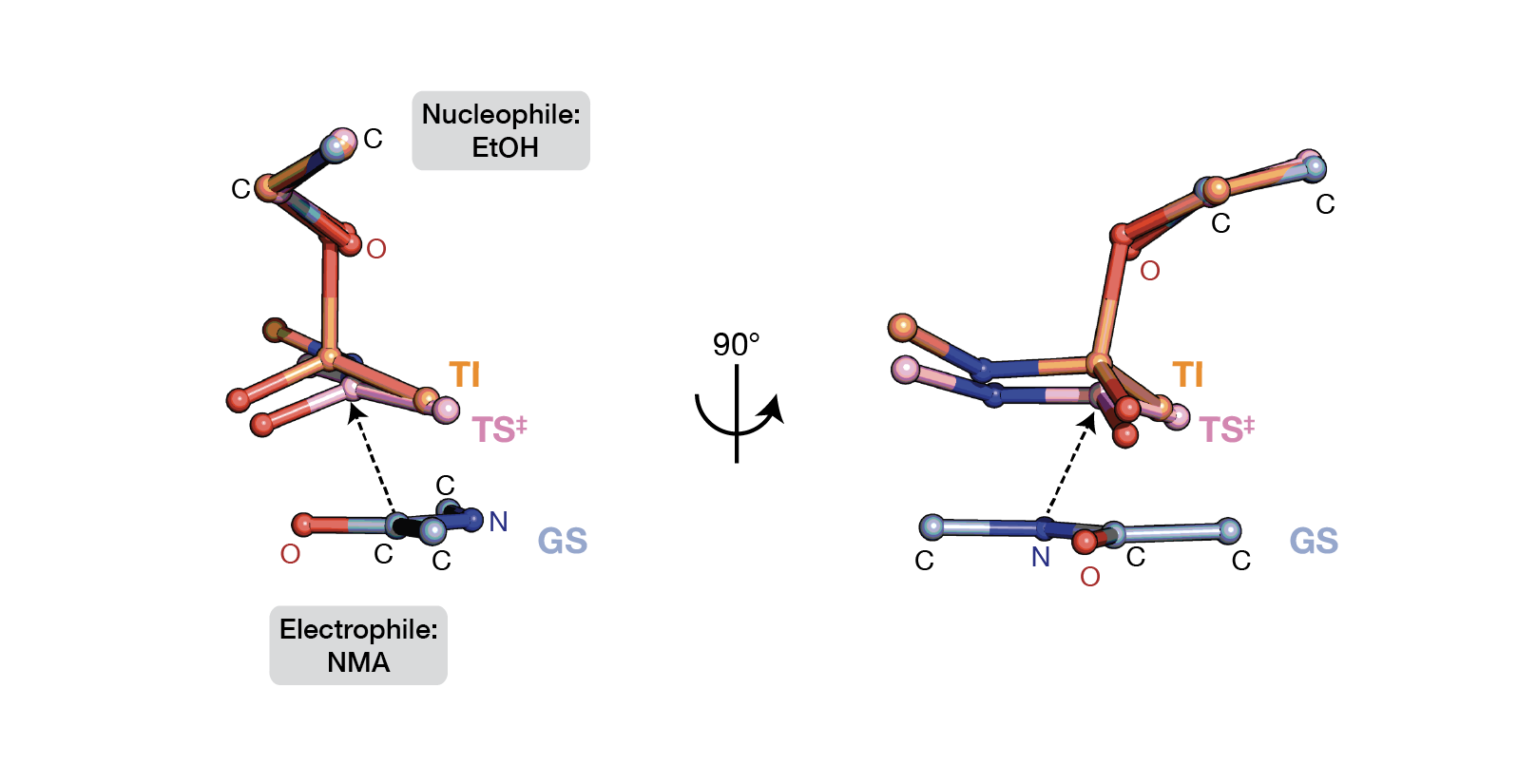

**Fig. S6.** The ground state (GS, blue), transition state (TS^‡^, pink) and tetrahedral intermediate state (TI, orange) structures determined by QM calculation of the solution reaction of nucleophilic addition on amide, using *N*-methylacetamide (NMA) as electrophile and ethanol (EtOH) as nucleophile. The tetrahedral intermediate state is highly similar to the transition state, as found for the reaction on serine proteases from prior QM/MM calculation (*109*). See also fig. S7 and table S5.

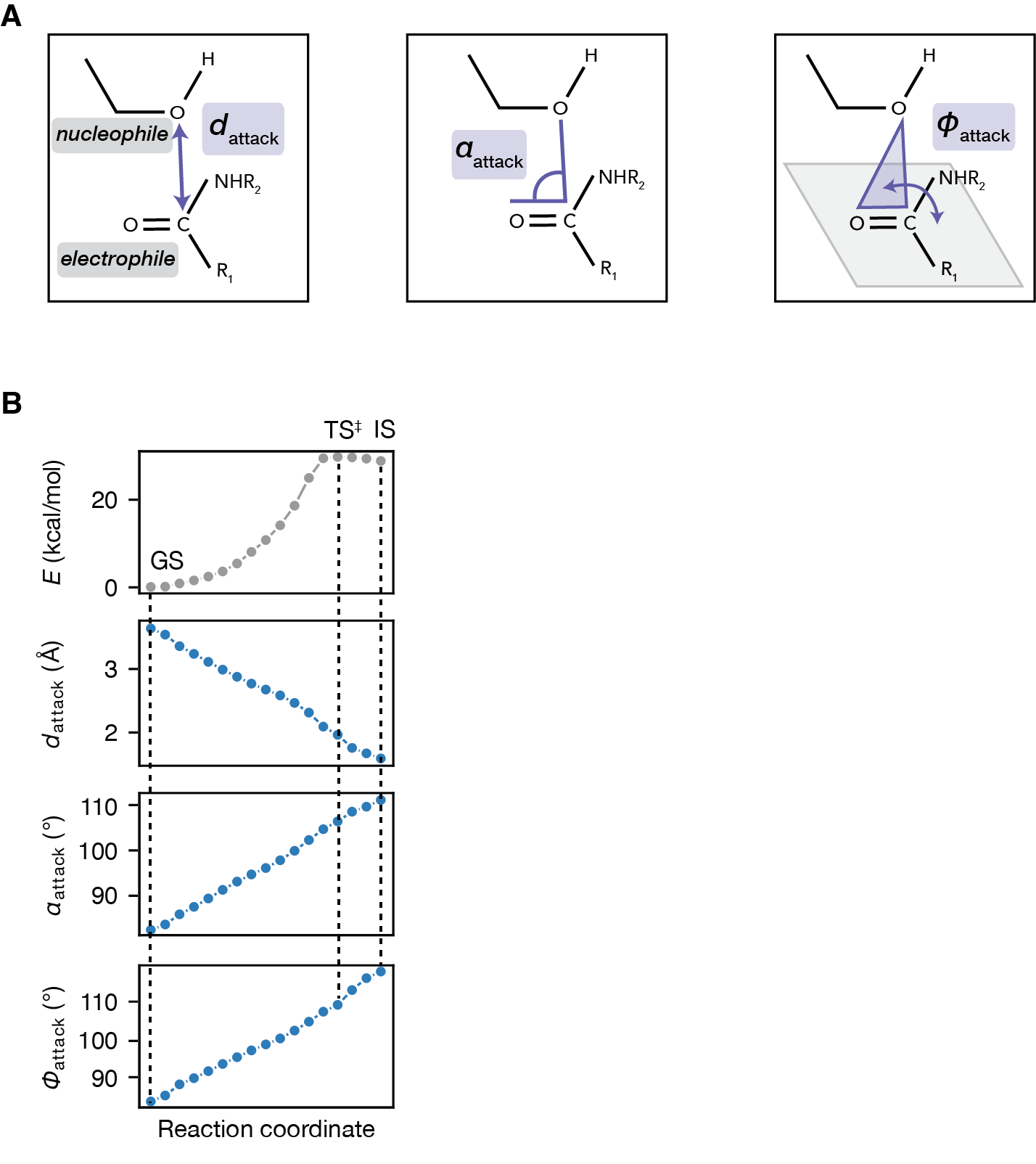

**Fig. S7.** Reaction coordinate for the nucleophilic addition on amide reaction in solution, determined by QM calculation using *N*-methylacetamide (NMA) as electrophile and ethanol (EtOH) as nucleophile. **(A)** The nucleophilic attack geometry is described by three parameters: the attack distance (*d*_attack_), defined by the distance between the nucleophilic and the electrophilic atom; the attack angle (𝛼_attack_), defined by the angle between the nucleophilic atom, the electrophilic atom and the oxygen on the reactive carbonyl group; and 𝜙_attack_, defined by the dihedral angle between two planes: (1) the plane defined by the nucleophilic atom, the electrophilic atom, and the oxygen on the amide carbonyl group (purple) and (2) the plane of the reactive amide carbonyl (gray). The same definitions are used in Fig. 2A. **(B)** Potential energy (*E*) and the changes in nucleophile•electrophile geometric parameters along the reaction coordinate. GS, ground state; TS^‡^, transition state; IS, tetrahedral intermediate state.

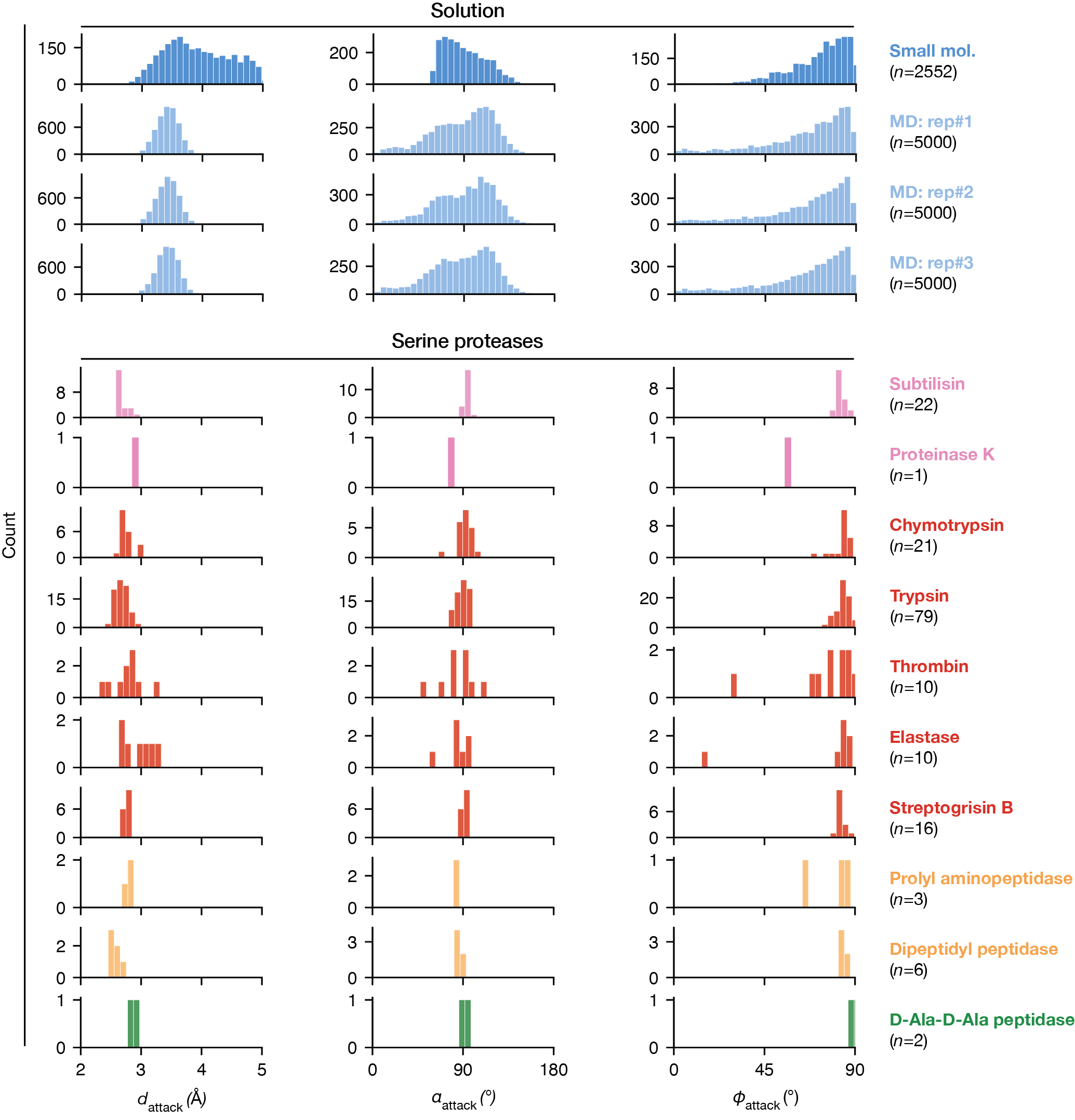

**Fig. S8.** Distributions of nucleophilic attack geometric parameters in “solution” (blue) and across serine proteases from clans SB (pink), PA (red), SC (yellow) and SE (green). The distributions in solution were obtained from small molecule structures (dark blue) and MD simulations (light blue). Small molecule distributions contain crystallographic structural fragments of hydroxyl•amide van der Waals interactions from the CSD after filtering irrelevant interactions that are shielded from the “line-of-sight” (see *Methods*). The MD distributions contain water•amide interactions of the water molecule that is the closest to the substrate analog NMA in each frame. Three replicates of 100 ns simulations produced the same distributions. Two serine proteases, thrombin and elastase, appear to differ from the others, with less positioning (see also fig. S9). The broader thrombin distribution may be related to the ability of its active site to respond to allosteric ligands (*186*). For elastase, the outliers with longer distances may arise from the absence of substrate binding interactions that facilitate substrate positioning (e.g. PDB: 1FLE is an outlier lacking a hydrogen bond between elastase Thr41and the substrate P2' site).

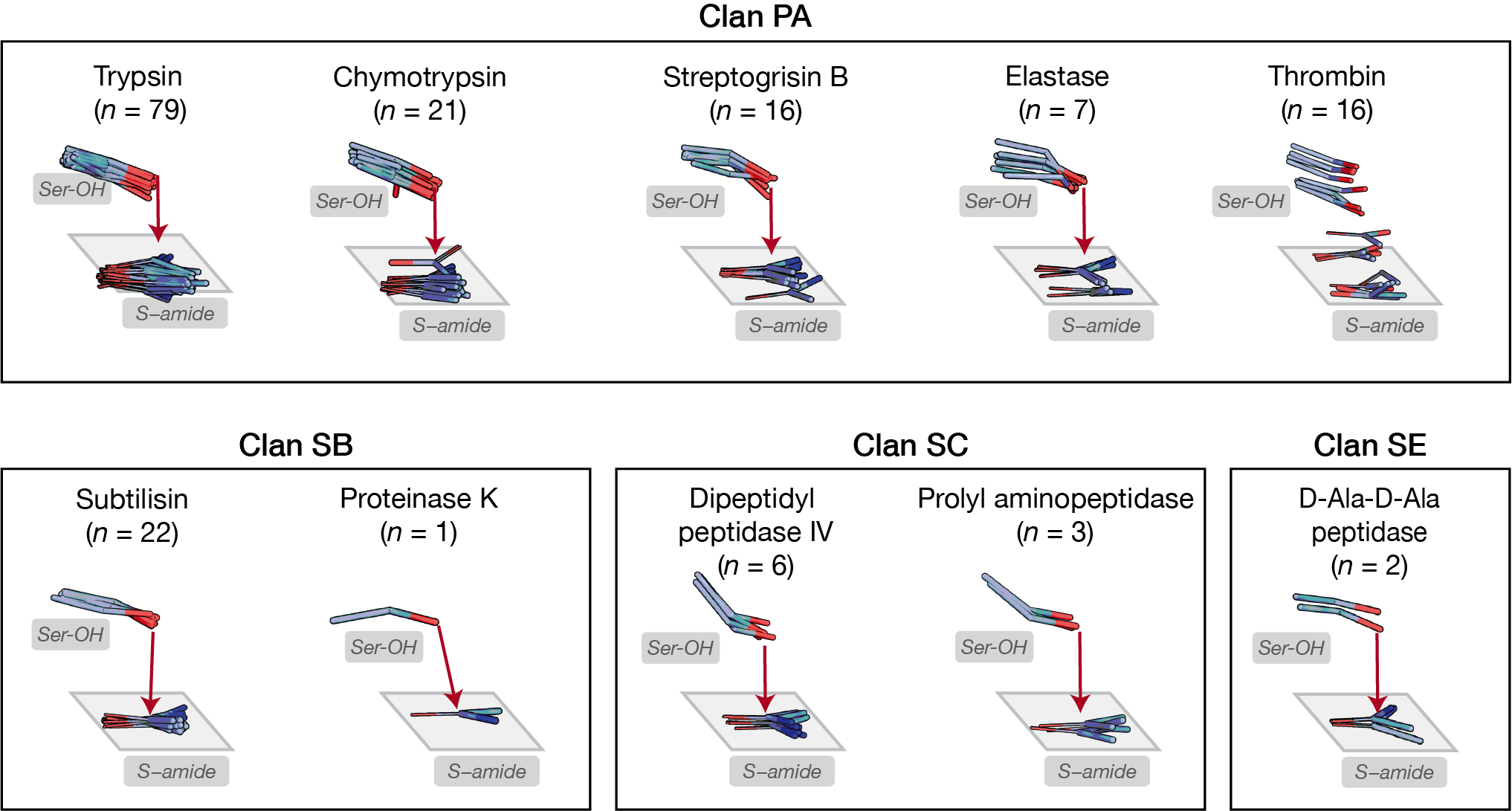
**Fig. S9.** Nucleophile•electrophile positions in the GSA-bound pseudo-ensembles of serine proteases. The catalytic serine sidechain (“Ser–OH”) and the substrate P1–P1' amide (“S–amide”) of each structure are shown. Structures were locally aligned on the sidechain atoms of the catalytic triad residues for each protease (table S3). The broader distribution in thrombin may be related to the ability of its active site to respond to allosteric ligands (*186*).

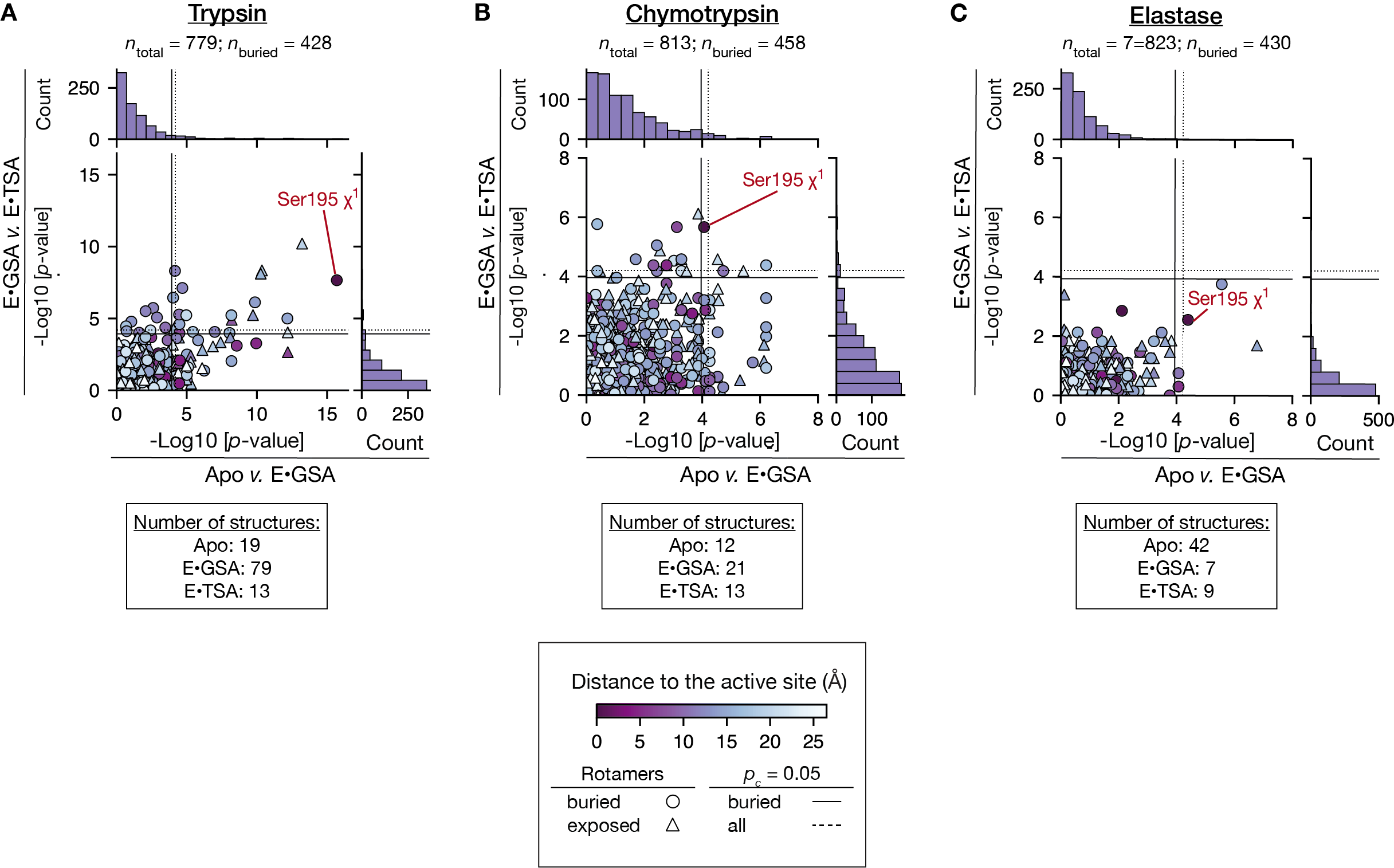
**Fig. S10.** Analysis of torsion angle changes upon substrate binding (apo *versus* GSA-bound) and nucleophilic attack (GSA-bound *versus* TSA-bound) and their significance. **(A)** Trypsin: among a total of 428 buried torsion angles (circles), 32 angles change significantly from apo to GSA-bound state and 23 from GSA-bound to TSA-bound states. *P*-values were determined by the Kolmogorov–Smirnov (K–S) tests (*24*) and significance threshold is set at a corrected *p*–value (*p*_c_) = 0.05 after multiple hypothesis (Bonferroni) correction. When all torsion angles including the solvent exposed ones (triangles) were included, 49 angles change significantly from apo to GSA-bound state and 22 from GSA-bound to TSA-bound states. Solid lines indicate the *p_c_* = 0.05 threshold for buried residues only; dashed lines indicate the *p_c_* = 0.05 threshold when all residues are considered. **(B)** Chymotrypsin: among a total of 458 buried torsion angles (circles), 24 angles change significantly from apo to GSA-bound state and 14 from GSA-bound to TSA-bound states. When all torsion angles including the solvent exposed ones (triangles) were included, 27 angles change significantly from apo to GSA-bound state and 15 from GSA-bound to TSA-bound states. **(C)** Elastase: among a total of 823 buried torsion angles (circles), 4 angles change significantly from apo to GSA-bound state and none from GSA-bound to TSA-bound states. When all torsion angles including the solvent exposed ones (triangles) were included, 3 angles change significantly from apo to GSA-bound state and none from GSA-bound to TSA-bound states. The smaller number of significant changes for elastase compared to chymotrypsin and trypsin presumably arises from the smaller number of structures in its GSA- and TSA-bound pseudo-ensembles (table S4).

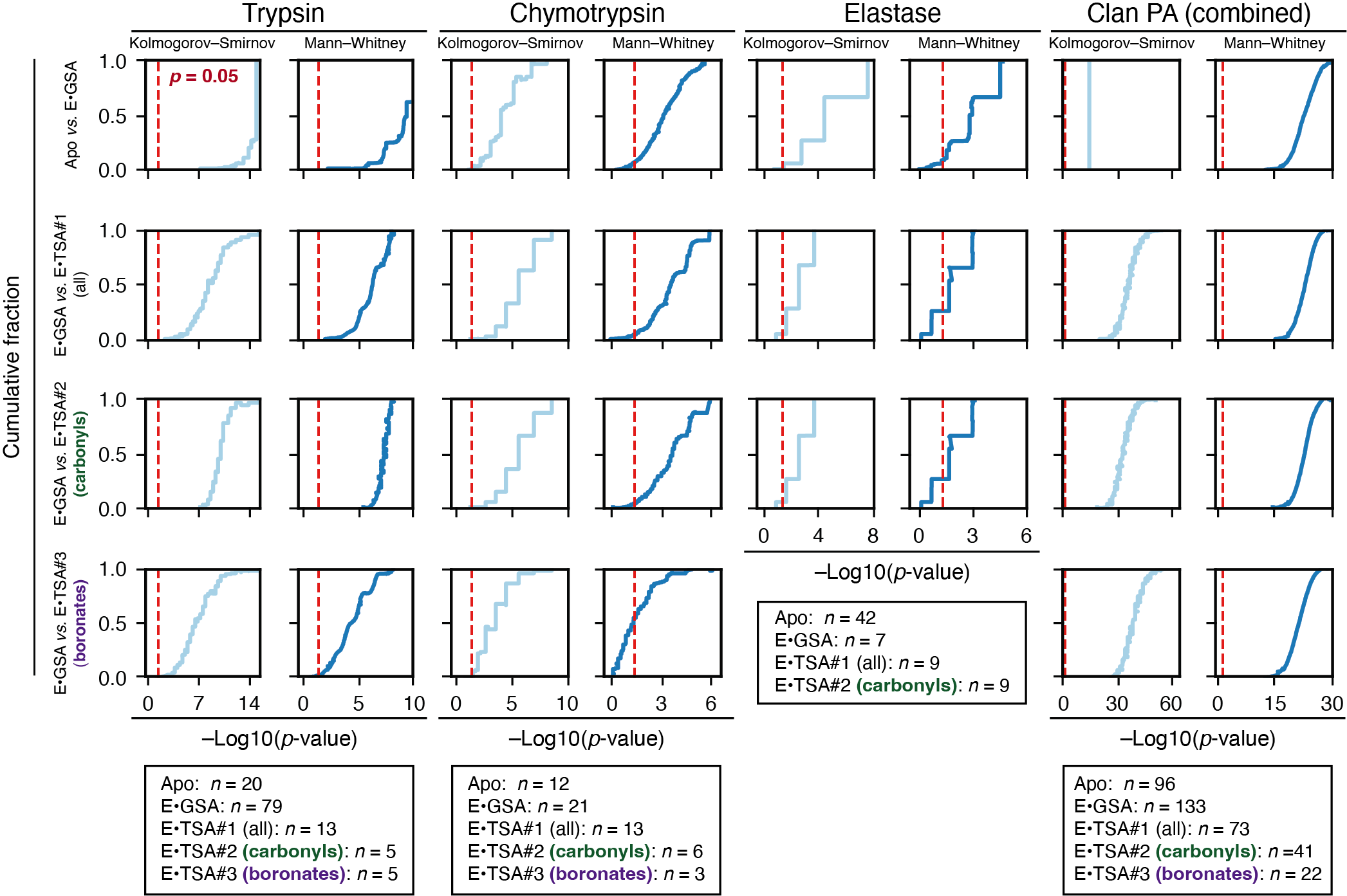

**Fig. S11.** Cumulative fraction of *p*-values from bootstrap analysis to determine the statistical stability of the observed changes in the catalytic serine (Ser195) χ^1^ and whether the significance of the changes depend on the types of ligands in the pseudo-ensembles. Two different statistical tests were used for each bootstrap analysis, the Kolmogorov–Smirnov test (*26*) (light blue) and the Mann–Whitney U test (*121*) (dark blue). The former tests whether the two distributions are *different*, and the latter tests whether one distribution is *greater* than that of the other. The tests were performed for the catalytic serine (Ser195) χ^1^ distribution of the apo *versus* GSA-bound pseudo-ensembles, for GSA-bound *versus* TSA-bound pseudo-ensembles (#1, including all TSA-bound structures), for GSA-bound versus subsets of TSA-bound pseudo-ensembles (#2, including all tetrahedral intermediate formed from a carbonyl compound; #3, including those formed from a boronic acid derivative). For each test, bootstrap resampling of the data and statistical tests were performed 1000 times and the cumulative distributions of their *p*-values [–Log10(*p*-value)] were plotted. The red dashed lines indicate the threshold of *p* = 0.05. In most cases, the change in χ^1^ is significant regardless of the type of test performed or the subset of TSA-bound structures included in the analysis.

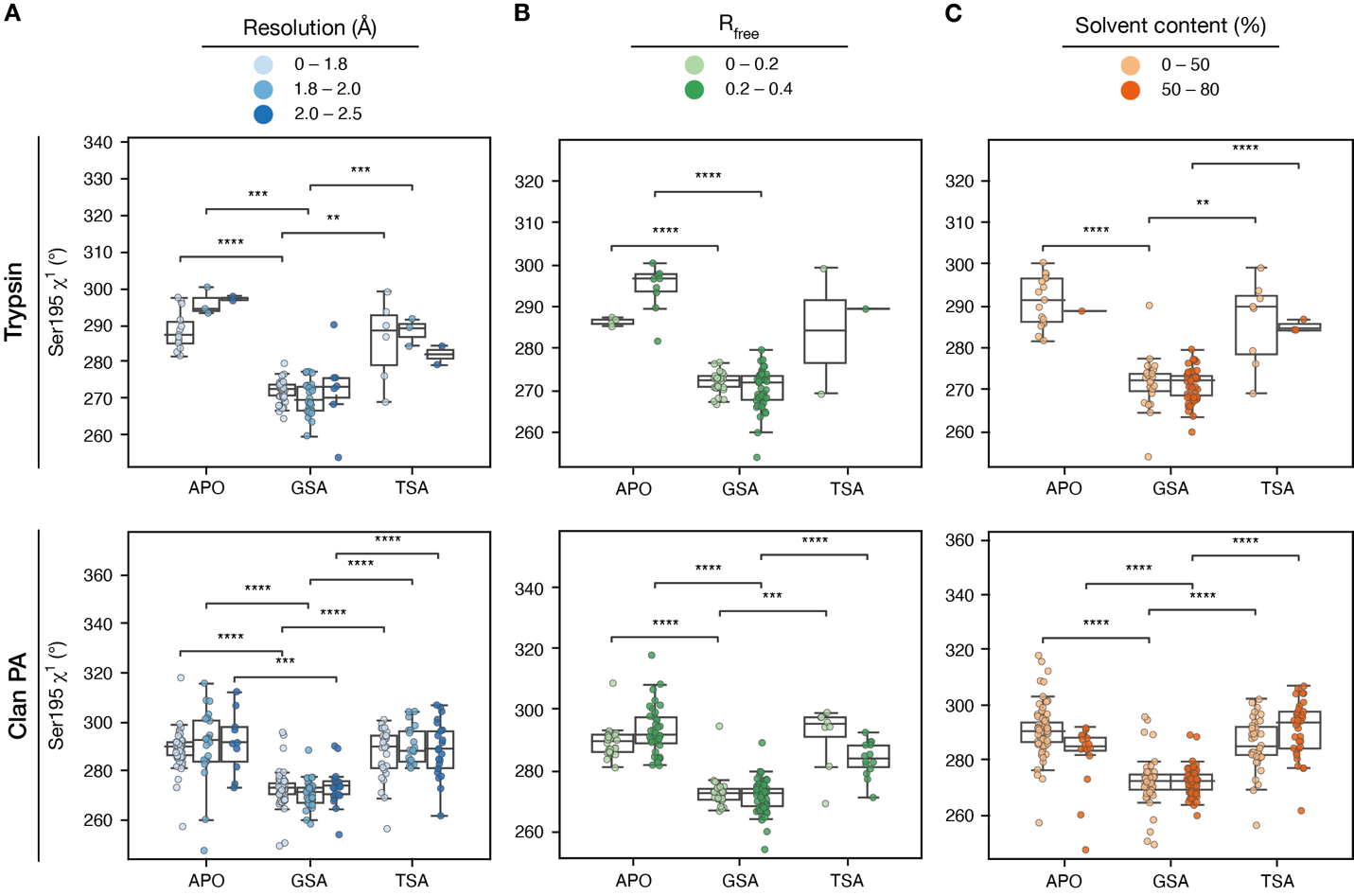

**Fig. S12.** Comparisons of the Ser195 χ^1^ distribution in the apo, GSA-bound and TSA-bound pseudo-ensembles of trypsin (top) and combined clan PA proteases (trypsin, chymotrypsin and elastase; bottom) across a range of resolution of the diffraction data (**A**), model R_free_ **(B)** and crystal solvent content (**C**). Number of * symbols indicate *p*-value determined by the Mann-Whitney test. Asterisk symbols indicate significance levels, with one asterisk corresponding to *p*-value $\leq$ 0.05 and with any number (*n*) of asterisks larger than one corresponding to $1\times{10}^{-n}\leq$ *p*-value$\leq1\times{10}^{-(n+1)}$. The direction and magnitude of Ser195 χ^1^ change across reaction states are consistent regardless of the variations in data and model quality of structures in the pseudo-ensembles.

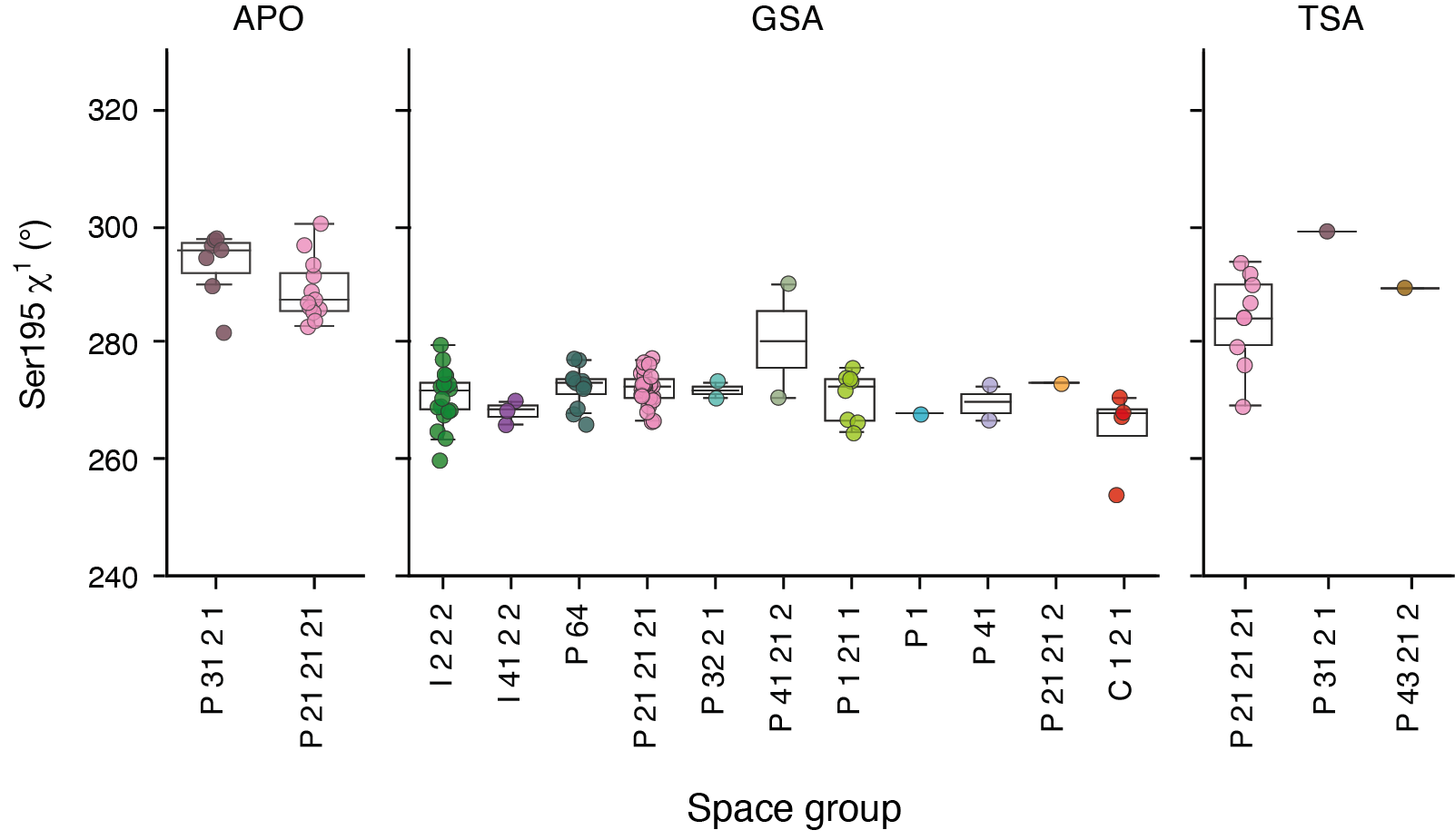

**Fig. S13.** Comparisons of the Ser195 χ^1^ distribution in the apo, GSA-bound and TSA-bound pseudo-ensembles of trypsin across a range of different crystal forms (space groups). The differences between reaction states are greater than the variations within the same reaction state between different space groups.

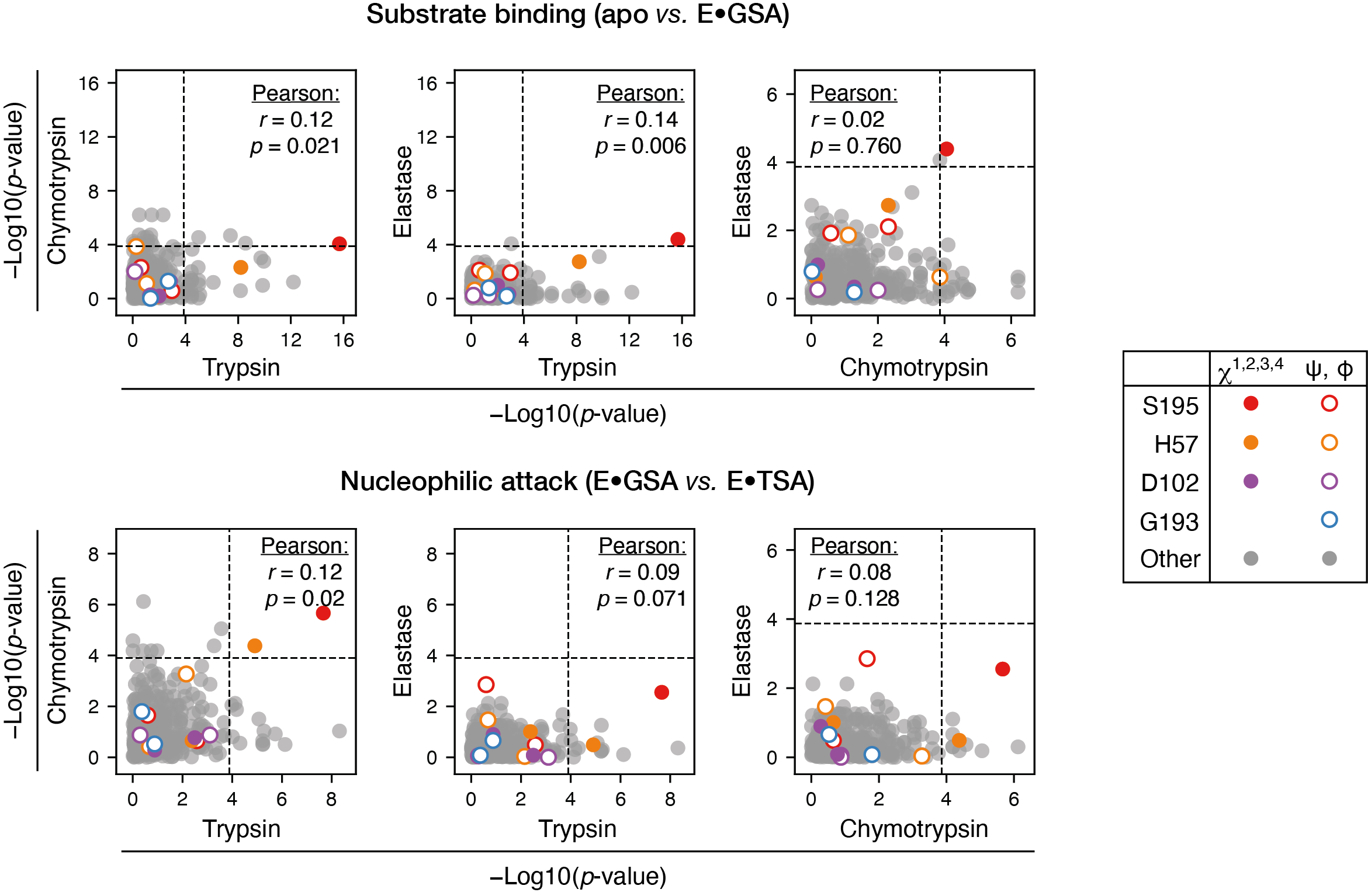
**Fig. S14.** Comparisons of the significance of the same torsion changes for three serine proteases in clan PA, highlighting catalytic residues. Trypsin, chymotrypsin and elastase sequences were aligned and the torsion distributions (backbone ϕ, ϕ, and sidechain χ angles) for each aligned position were calculated and compared across pseudo-ensembles representing different reaction states (apo *vs.* GSA-bound and GSA-bound *vs.* TSA-bound states). Only residues that can be aligned for all three proteases were included in the comparison. The significance of change [as indicated by -Log10(*p*-value)] for the same rotamers for different proteases are plotted. Dashed lines indicate the corrected *p ≤* 0.05 thresholds after Bonferroni correction. Pearson correlation was used to determine the relationship of torsion change significance between serine proteases (*r*, Pearson correlation coefficient; *p*, *p*-value of the correlation). The catalytic serine (Ser195) χ^1^ showed highly significant changes for both steps across all three enzymes and its conservation suggests a functional role. In contrast, no significant correlations were found overall, suggesting that additional motions along the reaction path for the three proteases are idiosyncratic and not conserved.

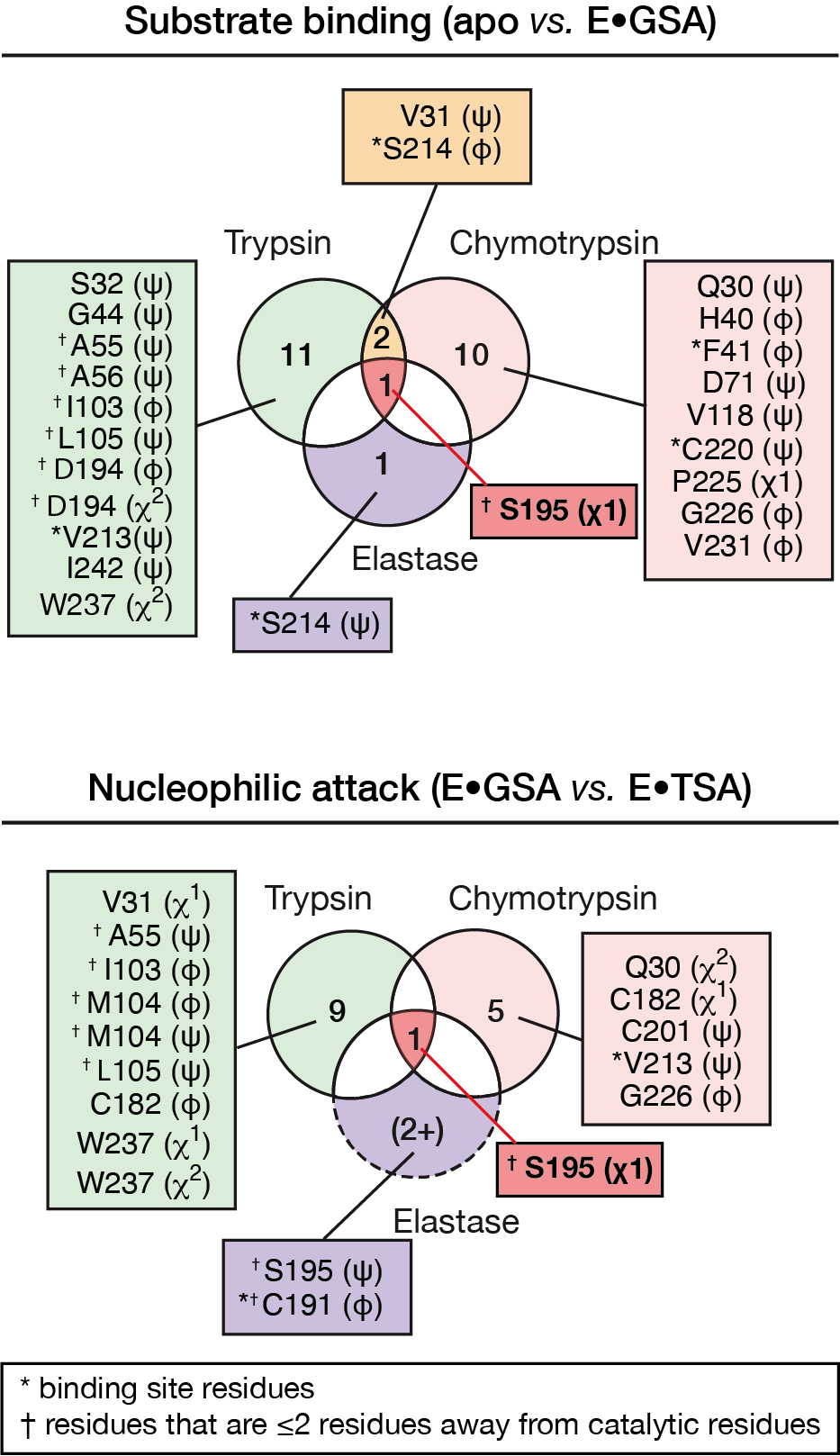

**Fig. S15.** Comparison of significant torsion changes for clan PA proteases (trypsin, chymotrypsin and elastase) in their buried residues. Comparisons are shown for apo *versus* GSA-bound (upper) and GSA-bound *versus* TSA-bound states (lower). Only residues that can be aligned for all three proteases were included in the comparison. Residue numbers follow the trypsin numbering scheme (and follow reference structures specified in table S4). Torsion changes that are significant are annotated in parenthesis. For elastase, no torsion changes have a corrected *p*-value below 0.05 after Bonferroni correction; for it, the three changes with their *p*-values ranked lowest were included (*p* < 0.008).

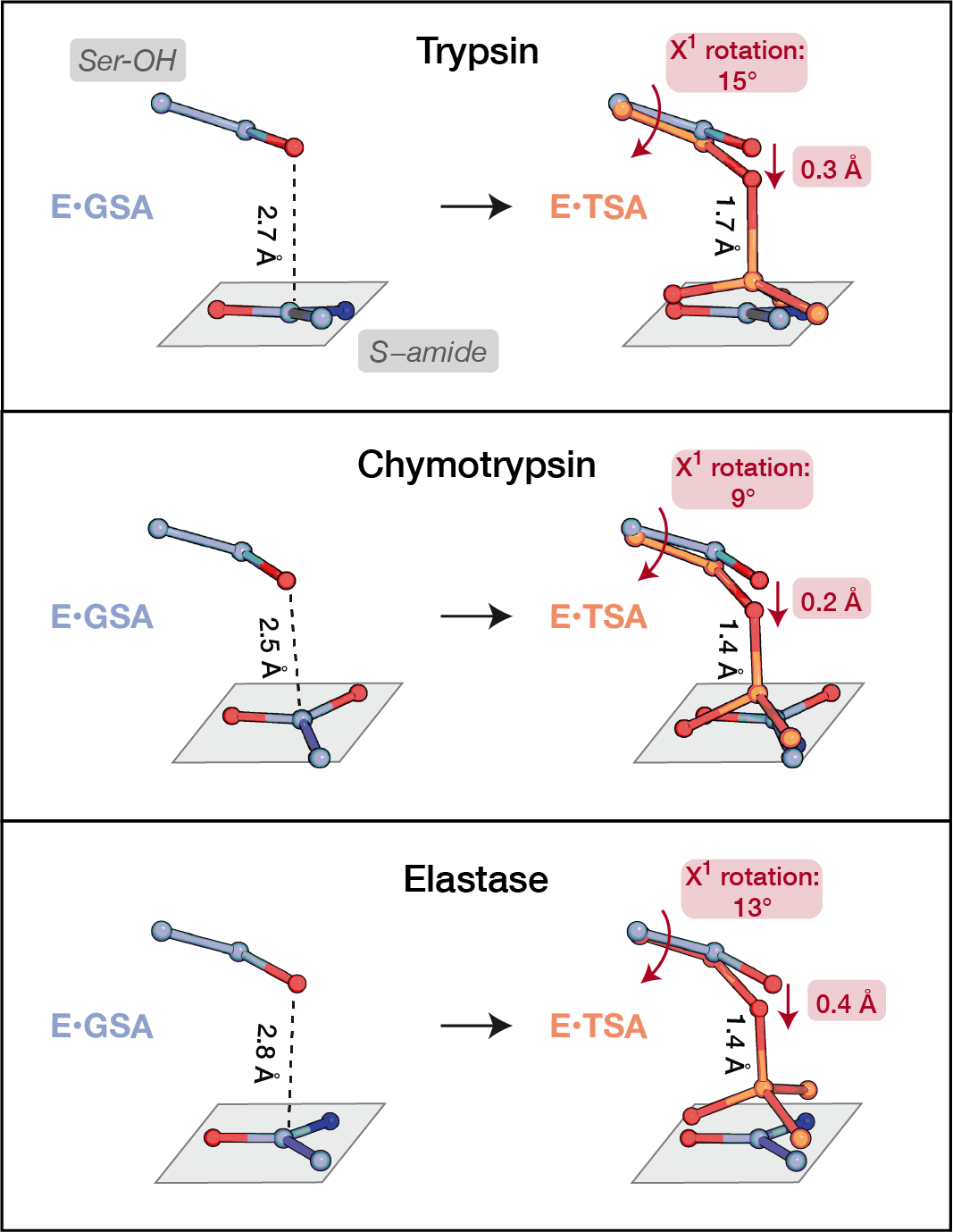

**Fig. S16.** A common reaction path involving the catalytic serine χ^1^ rotation for clan PA serine proteases. The “center” structures of the GSA-bound and TSA-bound pseudo-ensembles (defined as the structure with the median Ser195 χ^1^ value) were aligned locally on their catalytic triad residue atoms (Trypsin GSA-bound: 3M7Q, TSA-bound: 1TPP; chymotrypsin GSA-bound: 1GHA, TSA-bound: 1GMD; elastase GSA-bound: 3UOU; TSA-bound: 1EAU). See full ensembles in fig. S17.

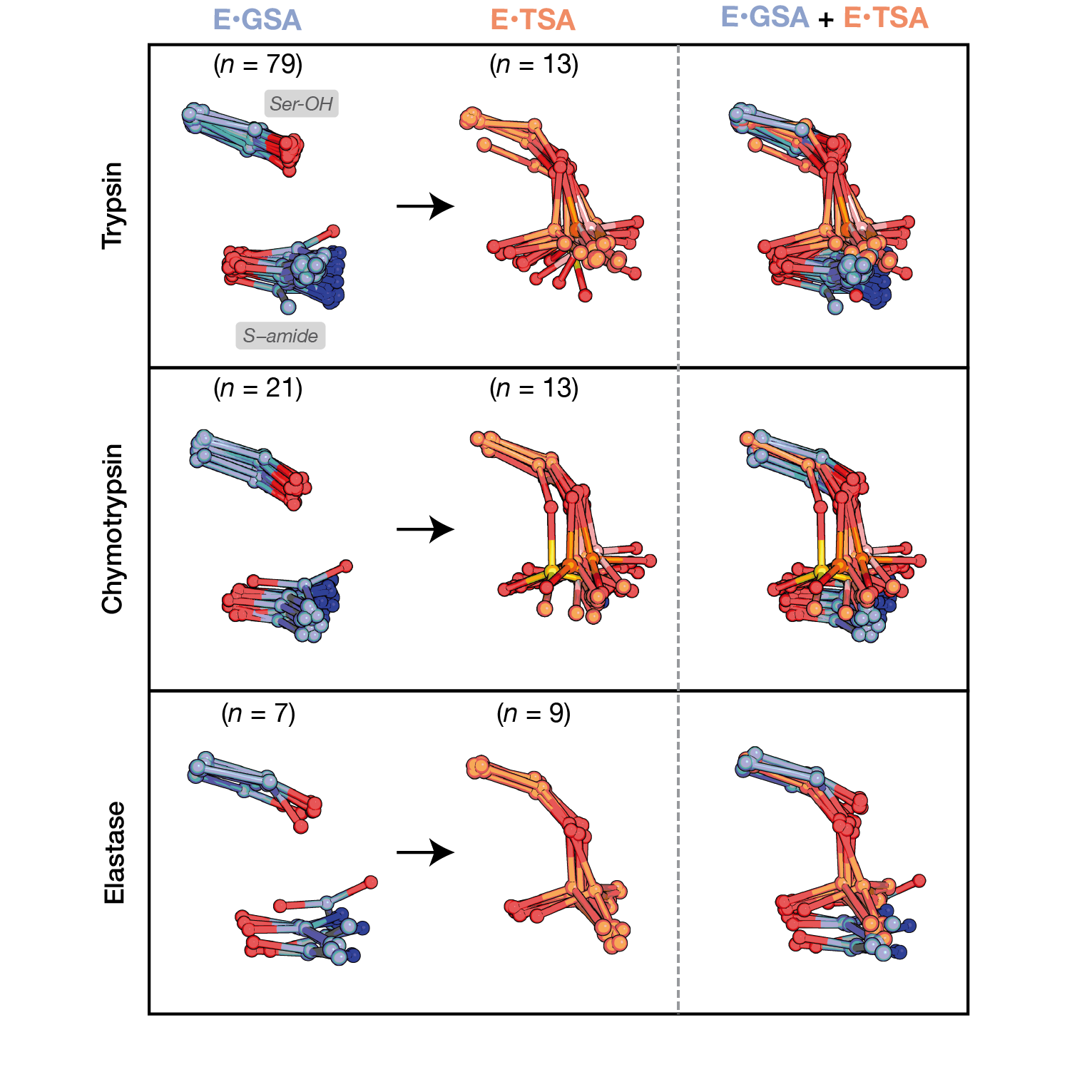

**Fig. S17.** A common reaction path involving the catalytic serine χ^1^ rotation for clan PA serine proteases (trypsin, chymotrypsin, and elastase). The full GSA-bound and TSA-bound pseudo-ensembles of clan PA serine proteases are shown, with the structures aligned locally on the catalytic triad residue atoms.

**_­_­**
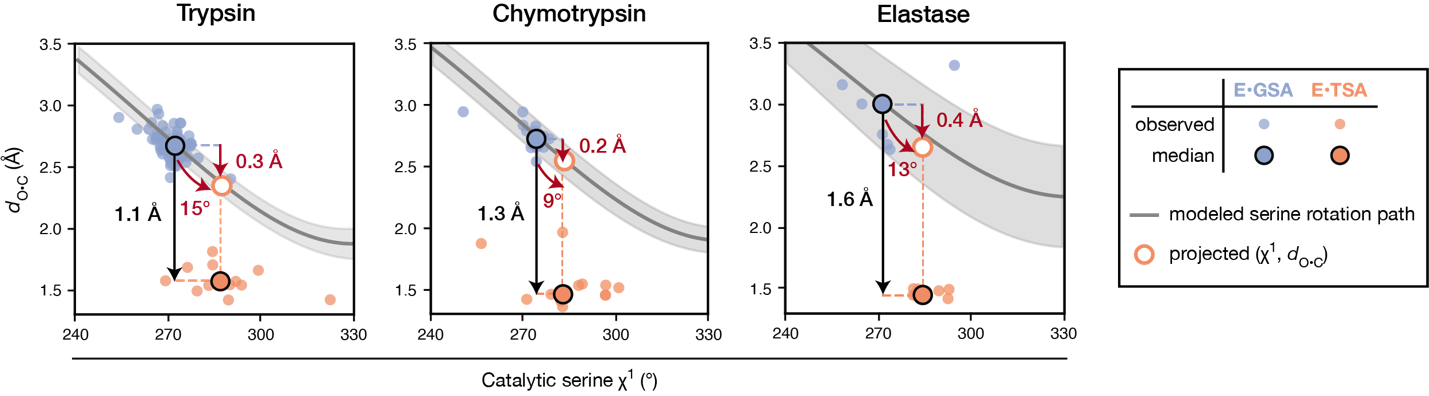
**Fig. S18.** Rotation of the catalytic serine χ^1^ shortens the distance between the reacting atoms (*d*_O•C_) in clan PA serine proteases (trypsin, chymotrypsin and elastase). The catalytic serine χ^1^ angles were rotated within the reactive *gauche*– rotameric well (from 240° to 330°) in each structure of the GSA-bound ensembles while keeping other atoms fixed. The *d*_O•C_ shortens as the serine χ^1^ is rotated (gray lines, with the gray shaded area indicating the 95% confidence intervals based on the ensemble of conformations). This observation supports the distance shortening shown in fig. S16 as a result of the serine rotation instead of other structural rearrangements that might occur but be obscured by structural alignment. Measurements of *d*_O•C_ and the catalytic serine χ^1^ angles from pseudo-ensembles are shown as small circles (blue: GSA-bound; orange: TSA-bound); their median values are indicated by the large outlined circles. The net Δ*d*_O•C_ resulting from the serine χ^1^ rotation was calculated by projecting the median TSA-bound χ^1^ on the modeled path (white circle with orange outline). Black arrows indicate the total change in median *d*_O•C_ values from the GSA- to the TSA-bound ensembles. Red curved arrows indicate median Δχ^1^ from GSA- to TSA-bound states; red straight arrow indicate the amount of Δ*d*_O•C_ resulting from Δχ^1^.

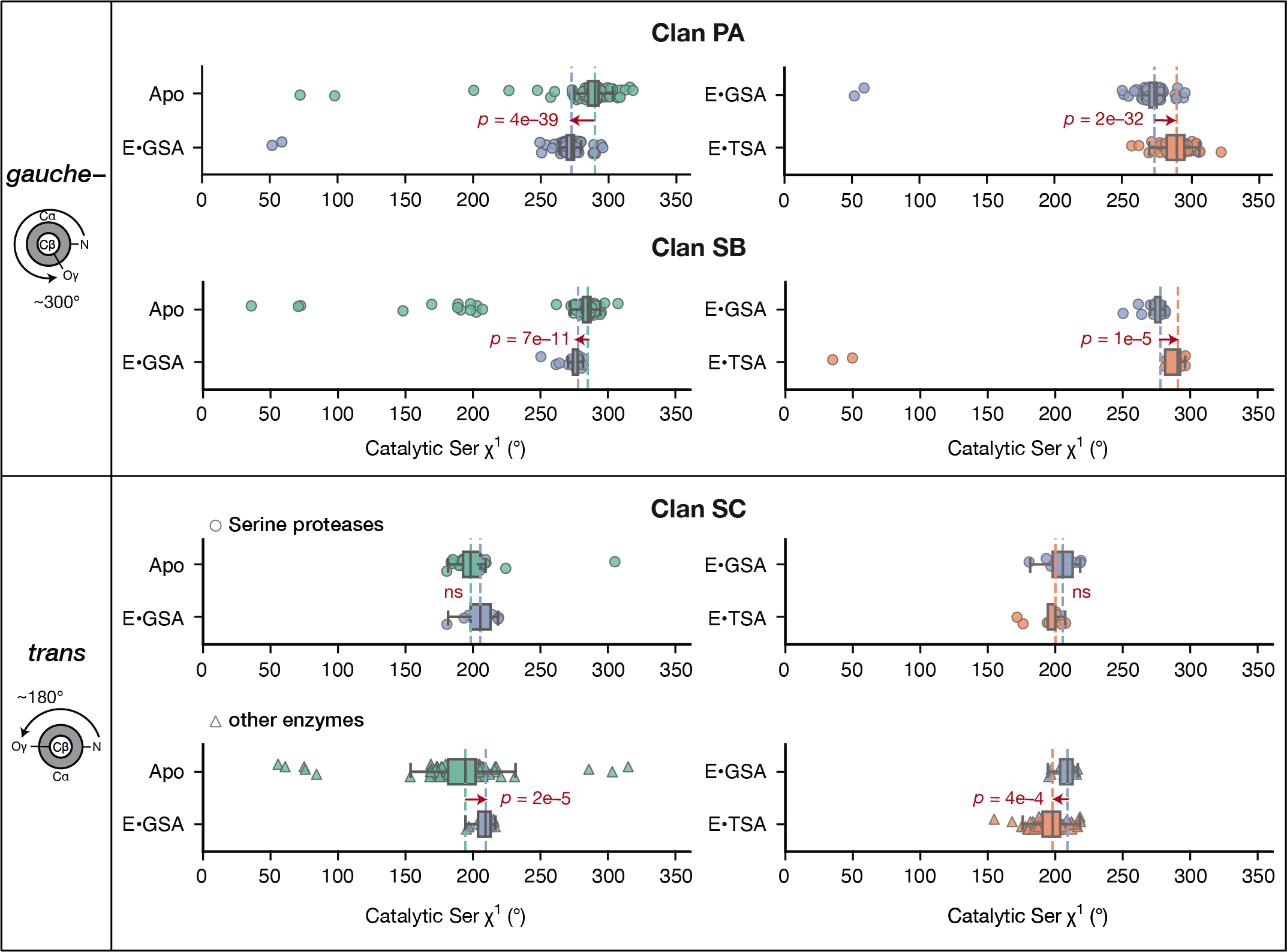

**Fig. ­­S19.** Changes of the catalytic serine χ^1^ across reaction states from serine proteases and additional enzymes in four structural clans (PA, SB, SE and SC). Clan PA, SB and SC distributions are the same as in Fig. 4A with the entire range of torsion angles (0 to 360°) shown. Dashed lines indicate the median value of each distribution (aqua: apo; blue: GSA-bound; orange: TSA-bound) (table S8). Arrows indicate the direction of change; *p*-values for changes that are significant (*p <* 0.05) are annotated beside the arrows.

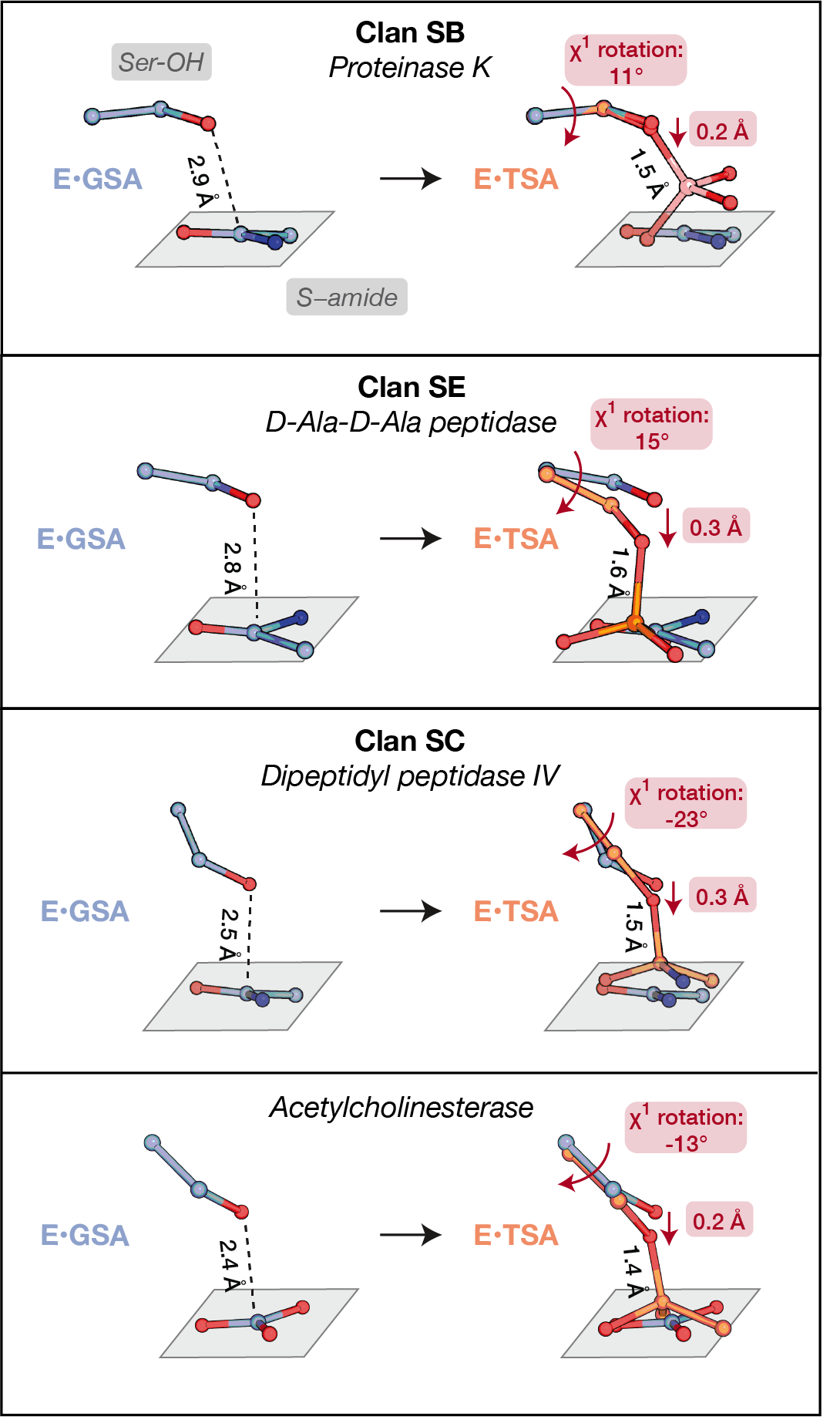

**Fig. S20.** A common reaction path involving the catalytic serine χ^1^ rotation for serine proteases from clan SB (proteinase K), SE (D-Ala-D-Ala peptidase) and SC (Dipeptidyl peptidase IV), as well as acetylcholinesterase, a non-protease enzyme from clan SC (also shown in Fig. 4B). The “center” structures of the GSA-bound and TSA-bound pseudo-ensembles (defined as the structure with the median catalytic serine χ^1^ value) were aligned on catalytic triad residue atoms (see table S2 for serine proteases; residues Ser203, His447 and Asp333 for acetylcholinesterase). The center structures are: 2DUJ (GSA-bound) and 2ID8 (TSA-bound) for proteinase K; 1PW1 (TSA-bound) and 1SCW (GSA-bound) for D-Ala-D-Ala peptidase; 1NU8 (TSA-bound) and 1WCY (GSA-bound) for dipeptidyl peptidase IV; 2HA0 (TSA-bound) and 2GYW (GSA-bound) for acetylcholinesterase. (See fig. S21 for full ensembles.) For proteinase K, the only available TSA-bound structure has a boron electrophile (pink); its electronic properties are different from other TSAs (with carbon electrophiles) due to its negative charge, potentially resulting in its slightly different conformation than other TSAs (see also supplementary text S8, *Assessing potential sources of variations in hydrogen bond lengths in pseudo-ensemble comparisons*). Nonetheless, a 11° rotation was observed for the catalytic serine of Proteinase K comparing the two structures (see also table S8).

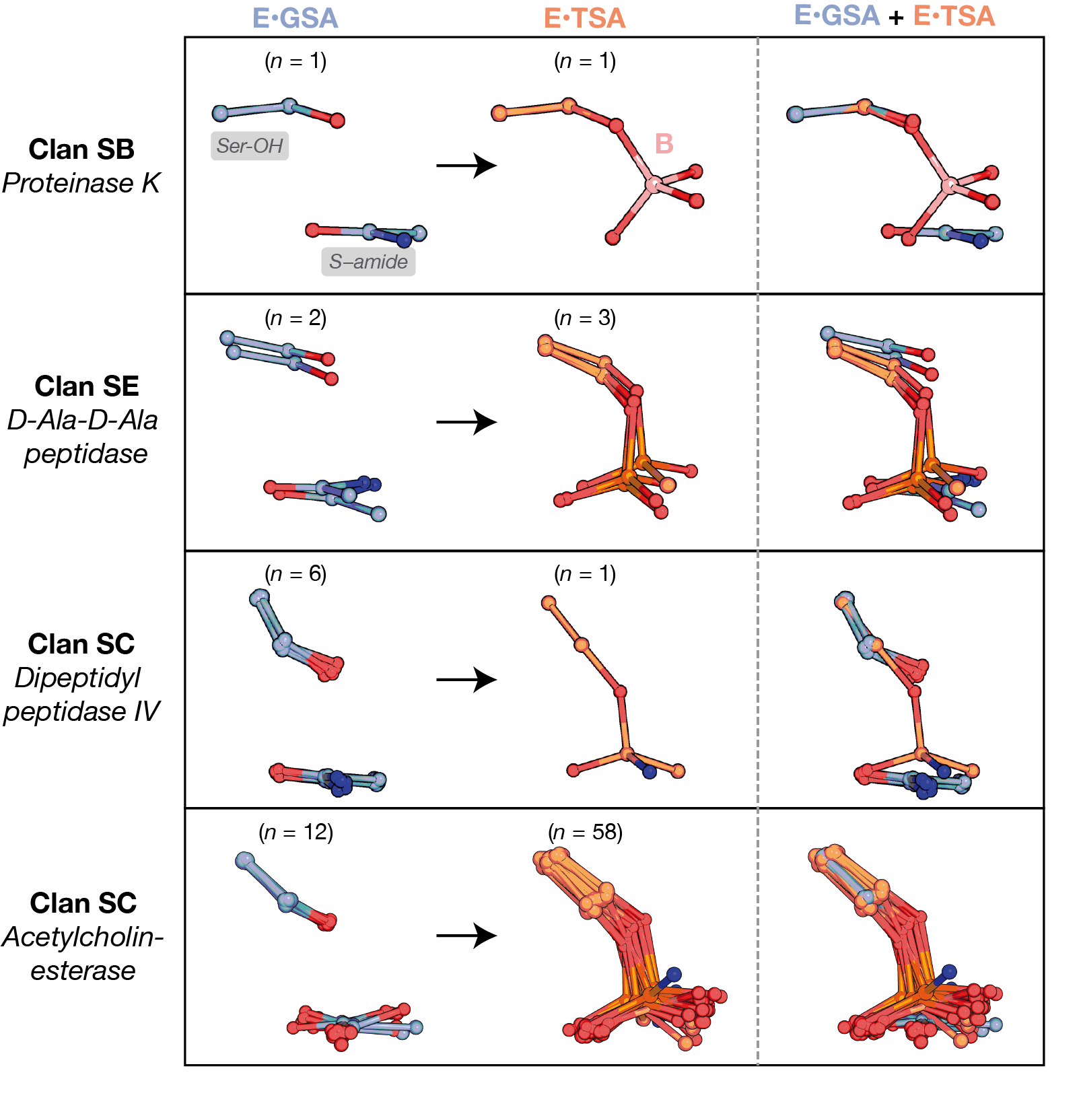

**Fig. S21.** A common reaction path involving the catalytic serine χ^1^ rotation for serine proteases in clan SB (proteinase K), clan SE (D-Ala-D-Ala peptidase) and clan SC (dipeptidyl peptidase), as well as acetylcholinesterase, a non-protease enzyme from clan SC. The full GSA-bound and TSA-bound pseudo-ensembles of clan PA serine proteases are shown, with the structures aligned locally on the catalytic triad residue atoms.

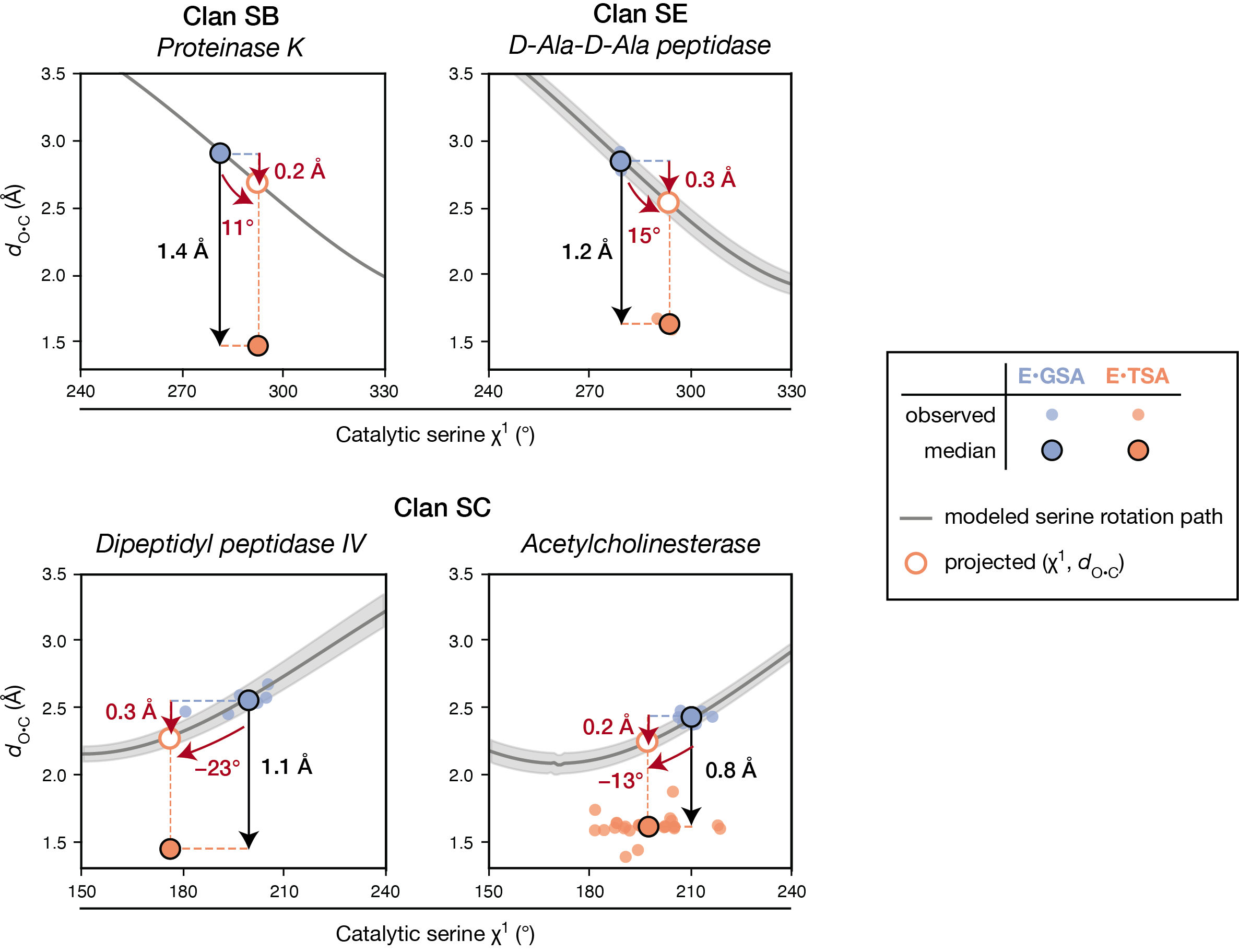

**Fig. S22**. Rotation of the catalytic serine χ1 shortens the distance between the reacting atoms (*d*_O•C_) in serine proteases from clan SB (proteinase K), SE (D-Ala-D-Ala peptidase) and SC (Dipeptidyl peptidase IV), as well as in acetylcholinesterase, a non-protease enzyme from clan SC. The catalytic serine χ^1^ angle was rotated in each structure of the GSA-bound ensembles while keeping other atoms fixed. The catalytic serines were rotated within the reactive *gauche*– rotameric well (from 240° to 330°) for clan SB and SE proteases and within the *trans* rotameric well (from 150° to 240°) for clan SC enzymes. The *d*_O•C_ shortens as the serine χ^1^ is rotated (gray lines, with the gray shaded area indicating the 95% confidence intervals based on the ensemble of conformations). This observation supports the distance shortening shown in fig. S20 as a result of the serine rotation instead of other structural rearrangements that might occur but be obscured by structural alignment. Measurements of *d*_O•C_ and the catalytic serine χ^1^ angle from pseudo-ensembles are shown as small circles (blue: GSA-bound; orange: TSA-bound); their median values are indicated by the large outlined circles. The net Δ*d*_O•C_ resulting from the serine χ^1^ rotation was calculated by projecting the median TSA-bound χ^1^ on the modeled path (white circle with orange outline). Black arrows indicate the total change in median *d*_O•C_ values from the GSA- to the TSA-bound ensembles. Red curved arrows indicate median Δχ^1^ from GSA- to TSA-bound states; red straight arrow indicate the amount of Δ*d*_O•C_ resulting from Δχ^1^.

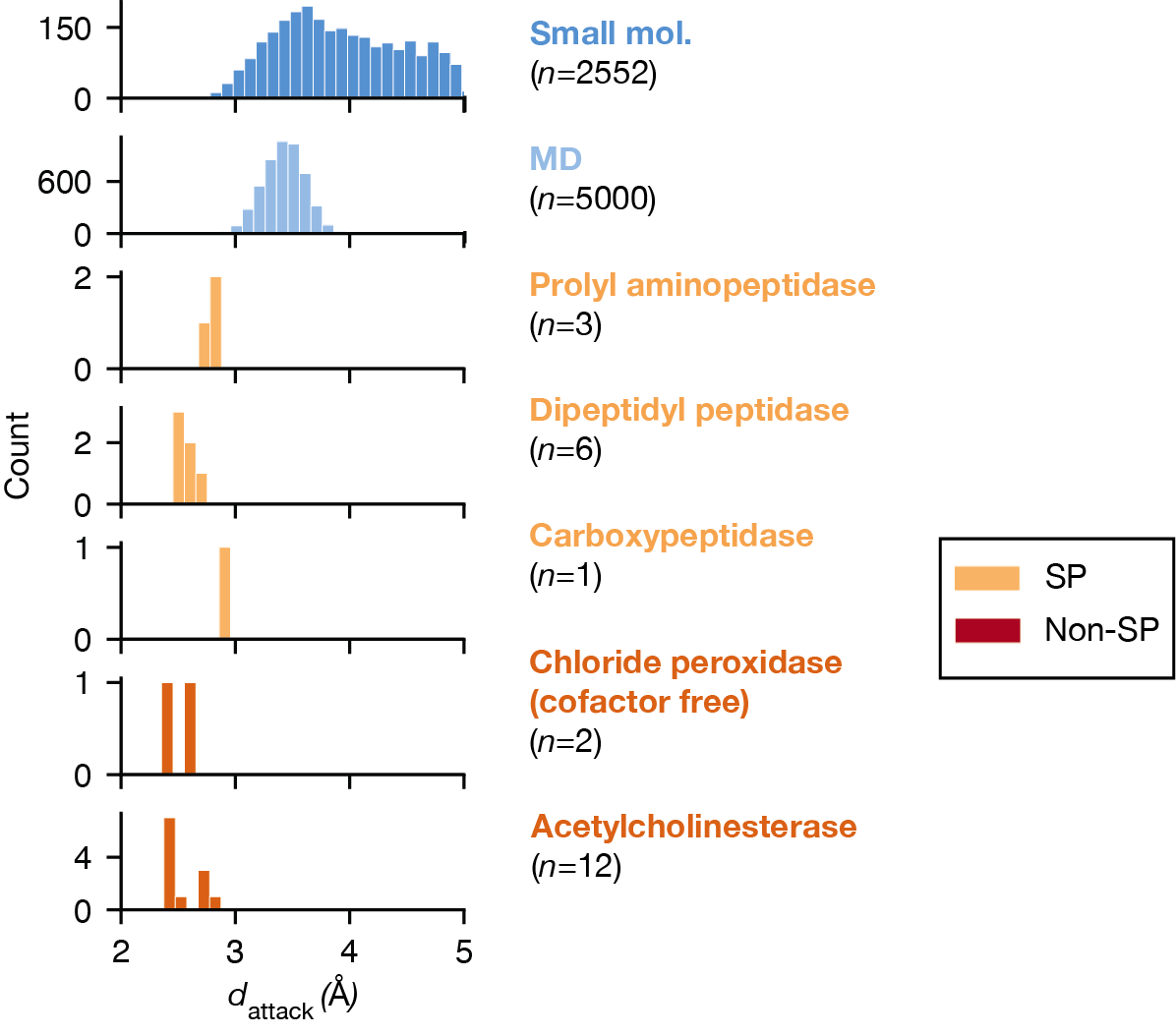

**Fig. S23.** Nucleophile•electrophile distances (*d*_attack_, defined in Fig. 2A) for clan SC serine proteases (“SP”, light orange) and non-proteases (“Non-SP”, dark orange). All enzymes have short *d*_attack_ compared to analogous interactions in solution from hydroxyl•amide interactions found in small molecules (dark blue) and MD simulations of water and NMA (light blue). The small molecule and MD distributions are also shown in fig. S8.

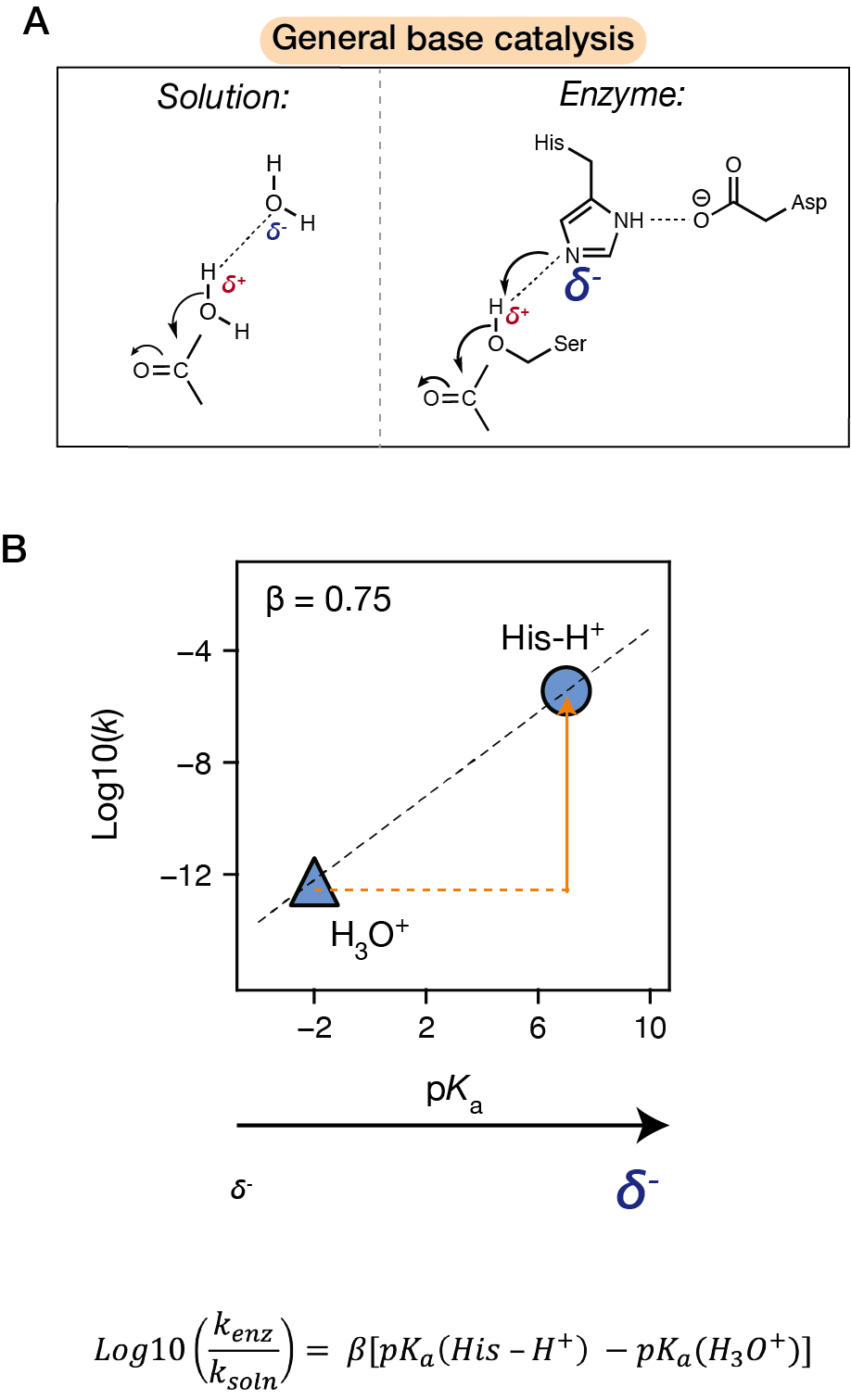

**Fig. S24.** Catalytic contribution from the general base. **(A)** The histidine general base has larger negative partial charge than water and thus favors proton abstraction from Ser−OH more than water. **(B)** A linear free energy relationship between general base p*K*_a_ and rate [Log10(*k*)] with a slope (brønsted β) of 0.75. The general base His, with a p*K*_a_ $\boldsymbol{\approx}$ 7 (*172*, *173*), provides 6 × 10^6^-fold rate enhancement (orange arrow) compared to reaction in water. See supplementary text S5 for details of the calculation.

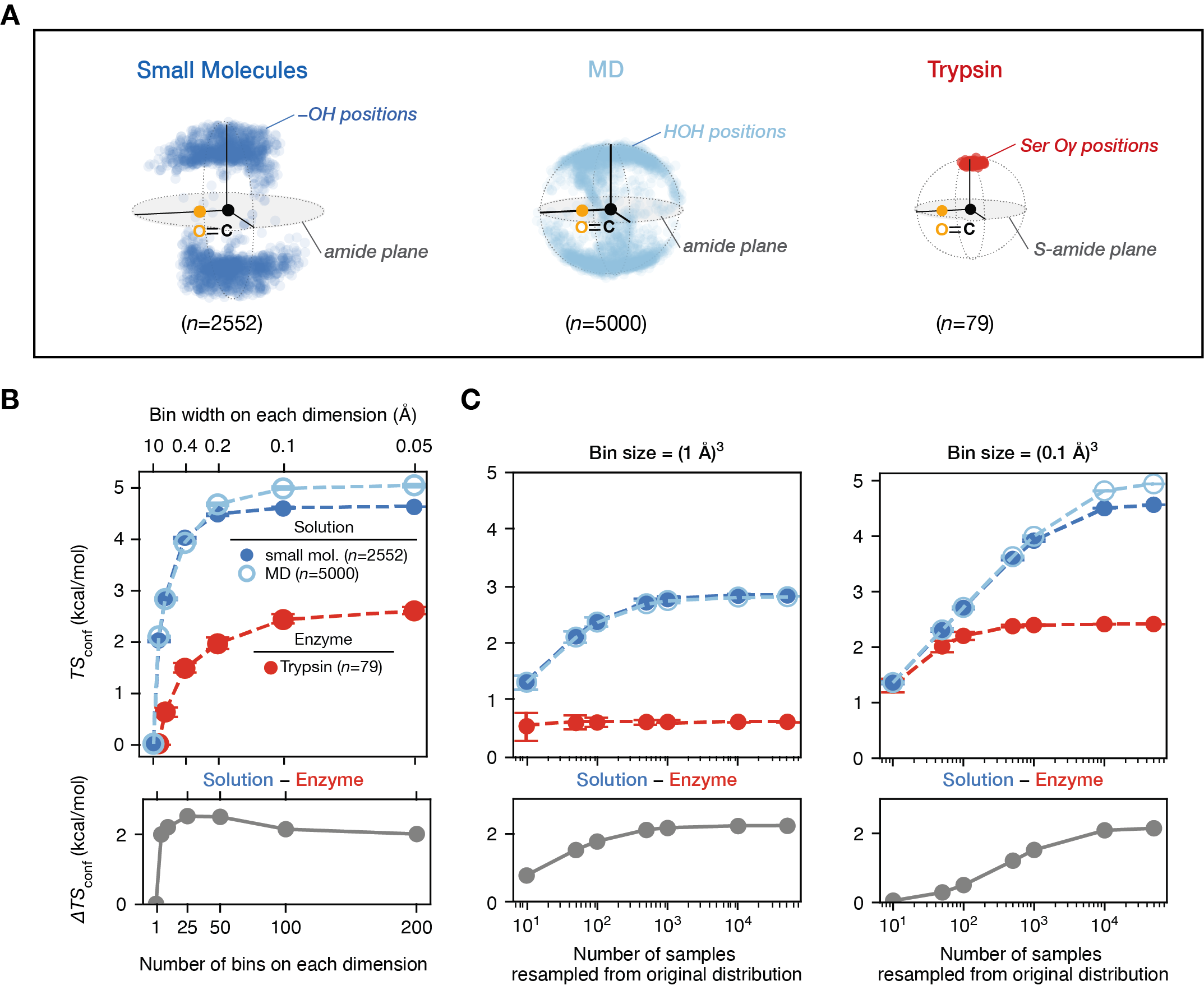
**Fig. S25.** Conformational entropies *(T•S*_conf_) calculated from ensemble data. **(A)** Small molecule hydroxyl•amide interaction distributions (dark blue), water•amide interactions from MD simulations (5000 snapshots from 100 ns simulation, light blue), and nucleophile•electrophile interactions in trypsin GSA-bound pseudo-ensemble (red). These distributions were used for the entropy calculations in (B) and (C). **(B)** *T•S*_conf_ values for the solution and enzyme distributions calculated using a range of bin size from (0.05 Å)^3^ to (10 Å)^3^ (upper, also shown in Fig. 5C) and the difference in *T•S*_conf_ (Δ*T•S*_conf_) between the small molecule and the trypsin distributions at each bin size (lower). The total considered volume is (10 Å)^3^, and the number of bins determines the size of each “microstate”, e.g. 10 bins on each dimension gives a microstate volume of (1 Å)^3^. Absolute entropy values are dependent on the bin size, and a bin size that match the actual microstates provides the most accurate entropy value. While we do not know the best bin size *a priori*, Δ*T•S*_conf_ values are stable across a range of bin sizes from (2 Å)^3^ to (0.02 Å)^3^. Circles indicate the mean from bootstrapping (200 repeats); error bars indicate the standard deviations. **(C)** *T•S*_conf_ values for the solution and enzyme distributions calculated across a range of sample size (from 10 to 50,000) by bootstrap resampling from the original distribution [with bin sizes of (1 Å)^3^ on the left and (0.1 Å)^3^ on the right]. Similar to the values calculated in (A), Δ*T•S*_conf_ converge to ~2 kcal/mol when large enough sample sizes are reached.

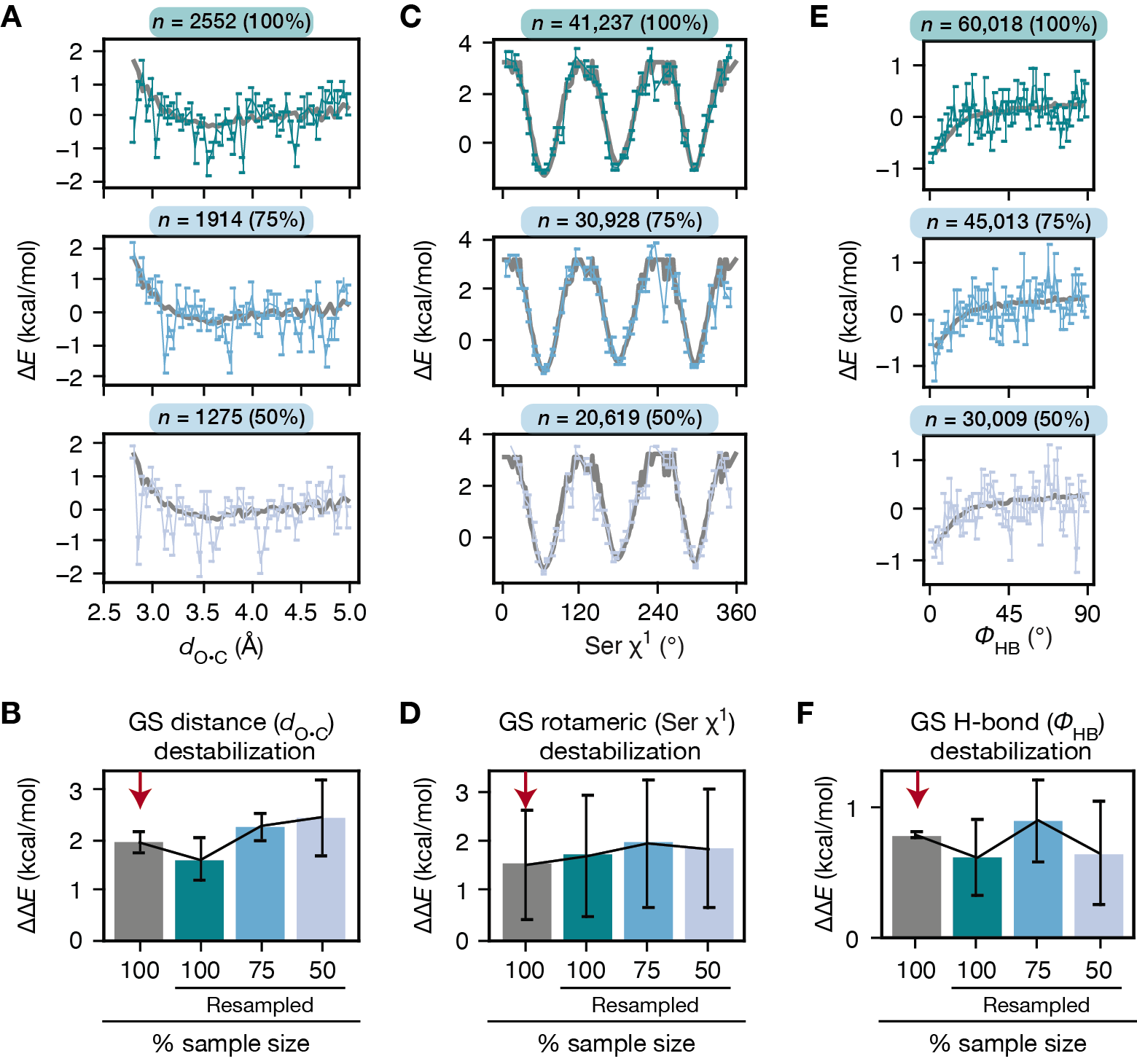

**Fig. S26.** Knowledge-based energy functions derived from bootstrapped datasets of a range of sample sizes (50% to 100% of the original dataset) and the catalytic contributions calculated from each function. The consistency in the amount of ground state destabilization derived from these energy functions, even when using only 50% of the data, provides confidence in the calculated catalytic contributions. **(A)** Knowledge-based energy function for O•C van der Waals distance (*d*_O•C_) built from small molecule structures of amide•hydroxyl interactions. The gray lines indicate the energy function derived from the entire collected dataset (without resampling). The dataset was randomly resampled to sample sizes that are 100% (dark green), 75% (medium blue) and 50% (light blue) of the original sample size. Energy functions were derived using each resampled dataset using the same method, and this process was repeated 1000 times. The colored lines represent mean energy values from the 1000 repeats, and the error bars indicate the standard deviations. **(B)** The amount of ground state destabilization from the short O•C van der Waals distance in trypsin, calculated using the energy functions derived from the full collected dataset (gray) and from the resampled datasets of 100% (dark green), 75% (medium blue) and 50% (light blue) of the original sample size. The *d*_O•C_ values from the trypsin GSA-bound ensemble and from solution QM (3.62 Å) were mapped onto the energy functions to calculate their energies, and the amount of ground state destabilization is calculated by subtracting the solution energy from the trypsin energies. The bar heights indicate the mean values calculated from all trypsin GSA-bound structures (*n* = 79) and the error bars indicate standard deviations. The red arrow indicates the value used in the main text (Fig. 5A). **(C)** Knowledge-based energy function for serine χ^1^ built from serine residues from the BBDEP2010 database (*27*); the same approach as described in (A) were used. **(D)** The amount of ground state destabilization from a partially eclipsed serine χ^1^ in trypsin, calculated using the same approach as described in (B). Serine χ^1^ from the trypsin GSA-bound ensemble and the TSA-bound ensemble were mapped onto the energy functions, and the amount of ground state destabilization is calculated by subtracting the TSA-bound from the GSA-bound energies. The bar heights indicate the mean values calculated from all trypsin GSA-bound (*n* = 79) and TSA-bound (*n* = 13) structures and the error bars indicate standard deviations. **(E)** Knowledge-based energy function of hydrogen bond $\boldsymbol{\phi}$_HB_ (defined in Fig. 5H) built from small molecule amide•carbonyl hydrogen bonds; the same approaches as described in (A) were used. **(F)** The amount of ground state destabilization from the suboptimal $\boldsymbol{\phi}$_HB_ in trypsin, calculated using the same approaches as described in (B).

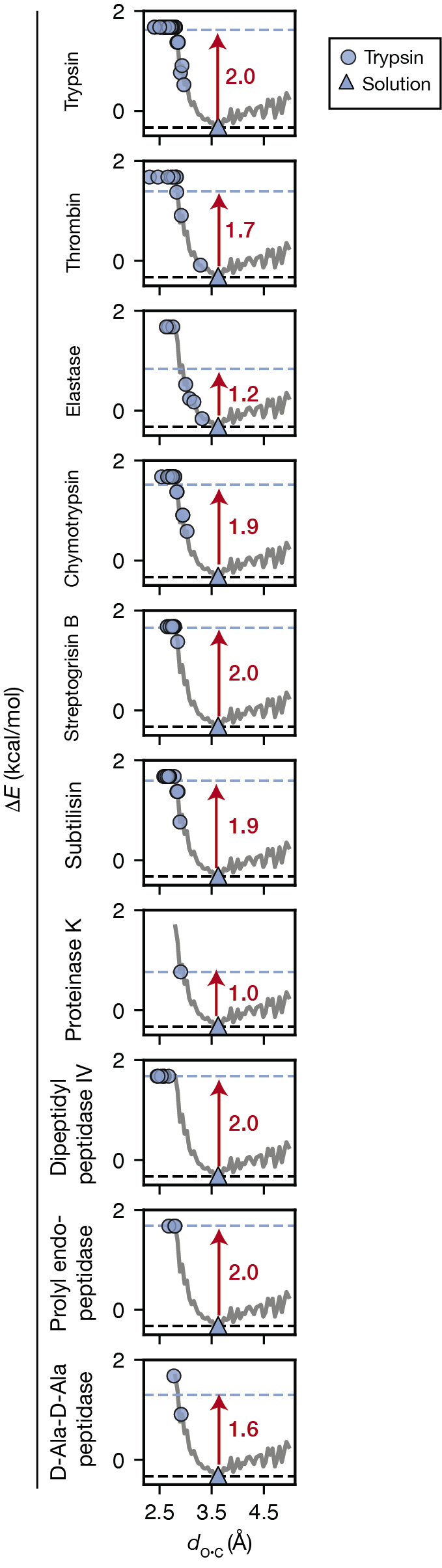

**Fig. S27.** Nucleophile•electrophile distances (*d*_O•C_) in serine protease pseudo-ensembles (blue circles) and in solution (QM; blue triangles) mapped on the knowledge-based energy function of *d*_O•C_ (derived from small molecule distribution shown in Fig. 2B). Blue dashed lines indicate the mean Δ*E*(*d*_O•C_) for the enzyme; black dashed line indicate the minimum of the knowledge-based energy function. Red arrows and values indicate the difference in mean Δ*E*(*d*_O•C_) between the enzyme and the solution distances (see also table S13).
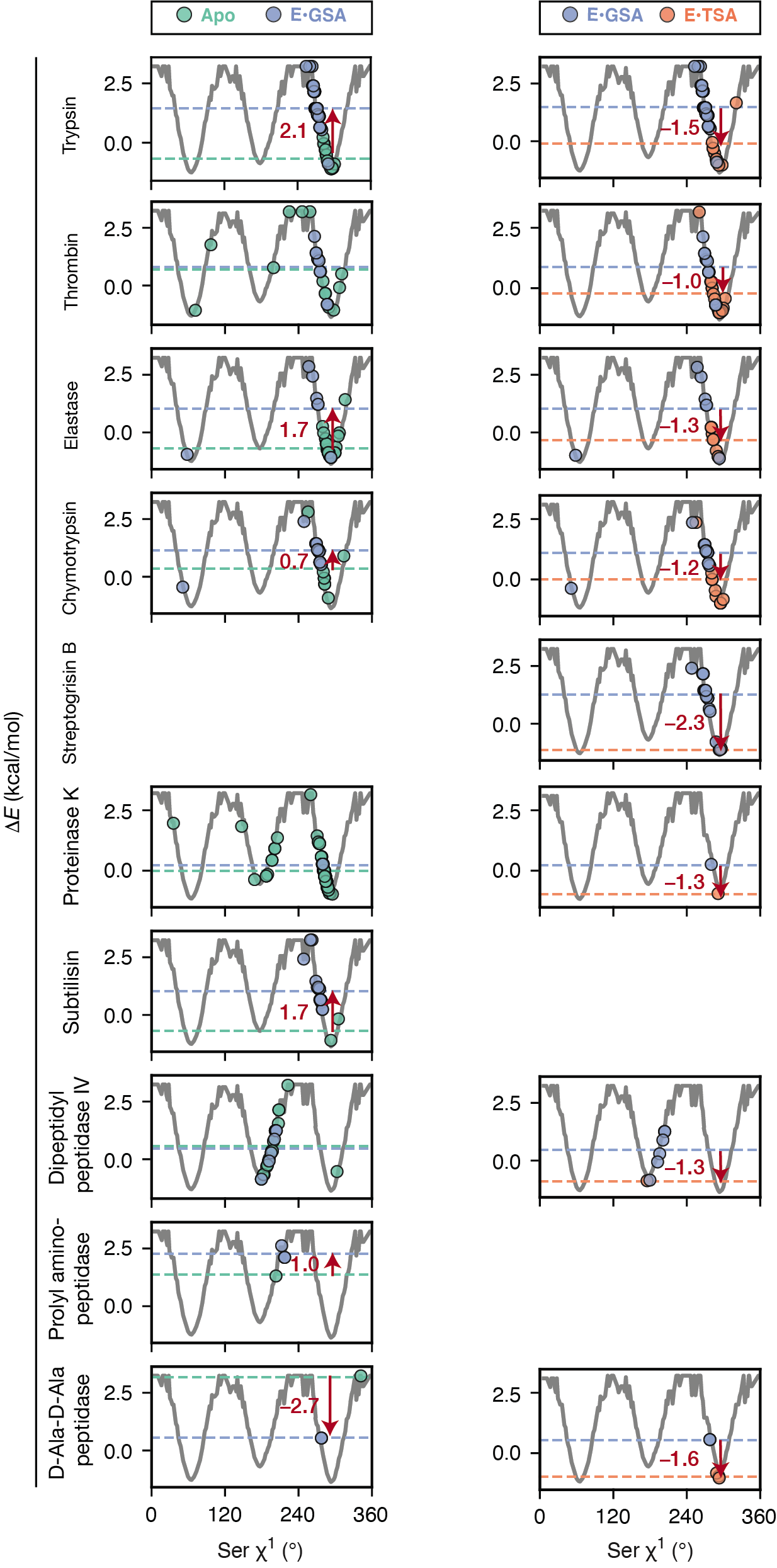

**Fig. S28.** Catalytic serine χ^1^ angles in serine protease apo (aqua circles), GSA-bound (blue circles) and TSA-bound (orange circles) pseudo-ensembles mapped on the knowledge-based energy function of serine χ^1^. Colored dashed lines indicate the mean Δ*E*(χ^1^) values for the corresponding states. Red arrows and values indicate the difference in mean Δ*E*(χ^1^) between states (see also table S14).

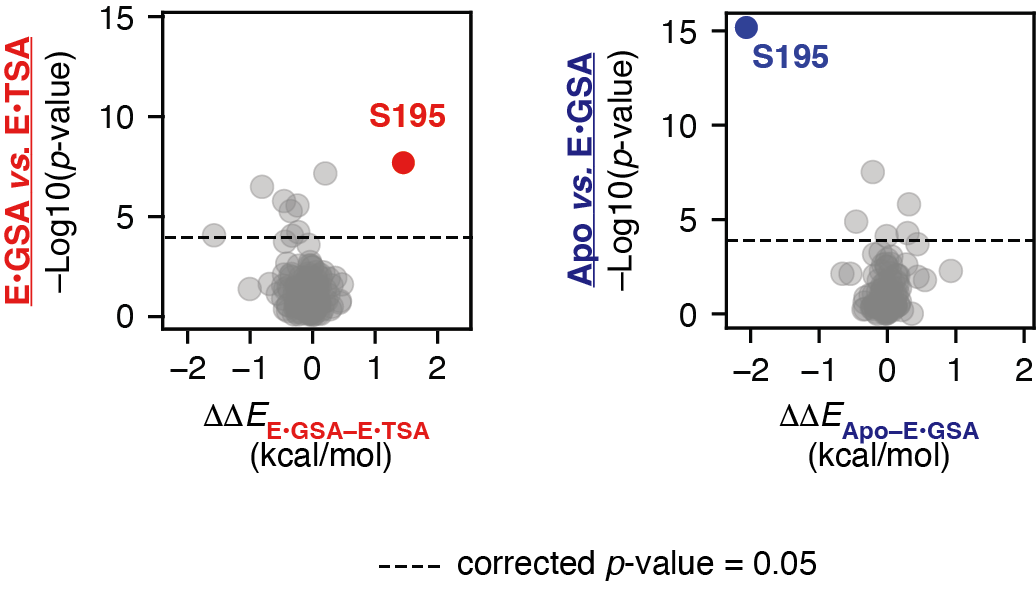

**Fig. S29.** The significance of each torsion change [–Log10(*p*-value) from the K–S tests] between pseudo-ensembles of different reaction states (apo *vs*. GSA-bound: blue; GSA-bound *vs.* TSA-bound: red) and the resulting energetic change (ΔΔ*E*) from each torsion change. The sidechain torsion angles from buried residues of trypsin (*n* = 181) were considered. Energy values were calculated from knowledge-based energy functions derived for each type of torsion angle using the same approaches as described for the serine χ^1^ angle. The solid lines indicate the *p*-value threshold of 0.05 after multiple hypothesis (Bonferroni) correction. Catalysis is derived from the GSA-bound *vs.* TSA-bound comparisons (red). The catalytic serine (Ser195) χ^1^ appears to be the only and the most significant change with >1 kcal/mol energetic difference. Additional changes appear to contribute minimally to the reaction barrier, with some apparently contributing to catalysis (ΔΔE_E•GSA–E•TSA_ > 0) and some apparently diminishing catalysis (ΔΔE_E•GSA–E•TSA_ < 0). Thus, we do not consider these changes as catalytic features in our minimal model.

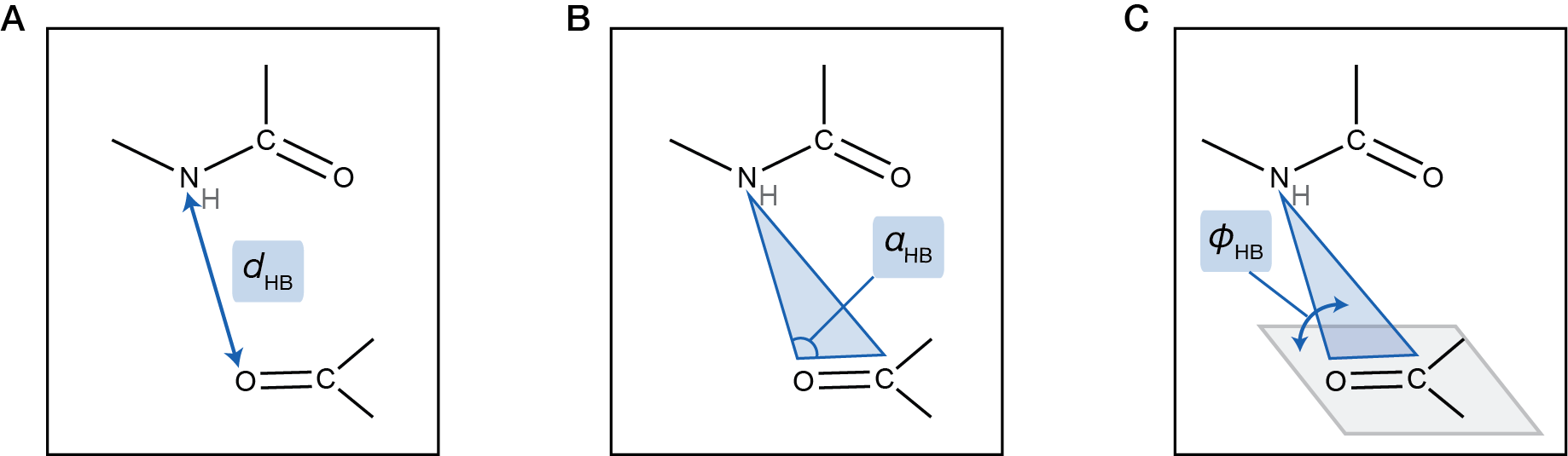

**Fig. S30.** Oxyanion hole hydrogen bond geometric parameters. **(A)** The hydrogen bond distance (*d*_HB_), defined by the distance between the heavy atoms, i.e. the donor (amide nitrogen) and the acceptor (carbonyl oxygen). **(B)** The hydrogen bond angle (𝛼_HB_), defined by the angle between the donor, the acceptor, and the carbon atom on the accepting carbonyl group. **(C)** 𝜙_HB_, defined by the dihedral angle between the two planes: (1) the plane defined by the donor, the acceptor, and the carbon on the accepting carbonyl group (blue), and (2) the plane of the accepting carbonyl (gray).

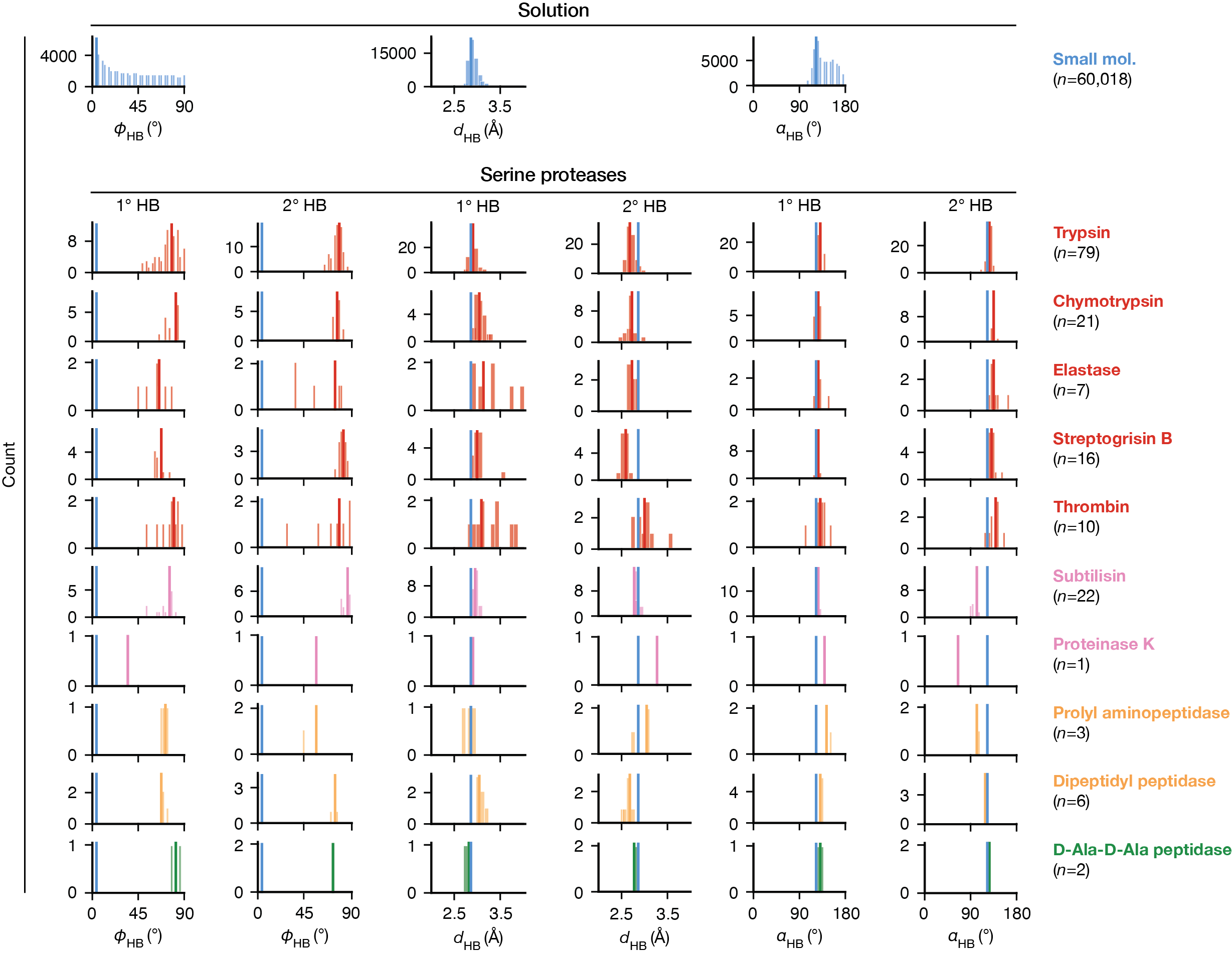

**Fig. S31**. Distributions of hydrogen bond geometries in small molecule amide•carbonyl hydrogen bonds *versus* in the oxyanion holes of GSA-bound serine proteases. The two hydrogen bonds, 1° and 2°, are defined in Fig. 1A. Geometric parameters are defined in fig. S30. For serine protease distributions, structural clans are denoted by colors (red: PA, pink: SB, yellow: SC, green: SE). The blue lines indicate the most probable value from the small molecule distribution and are repeated in each panel for comparison.

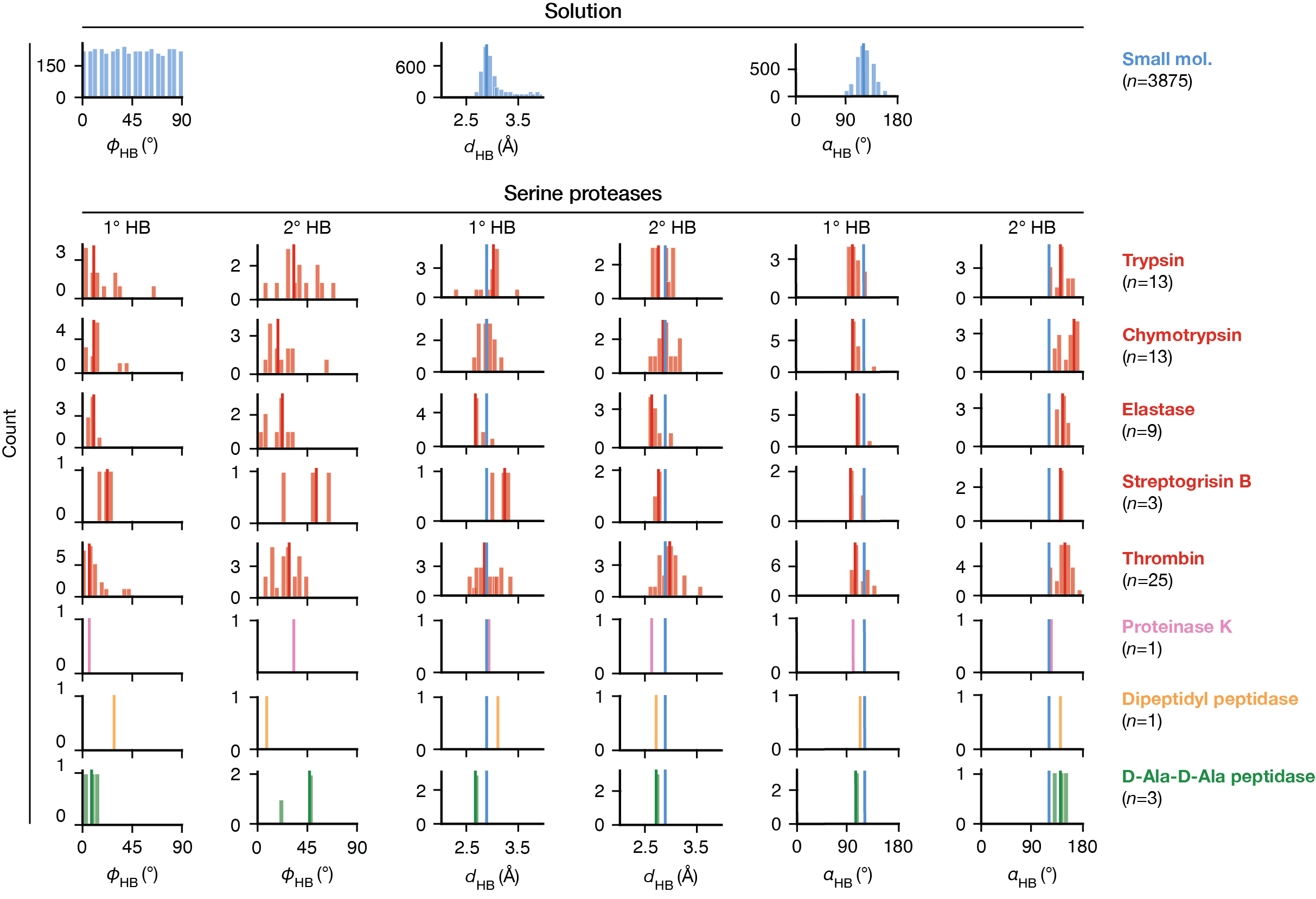

**Fig. S32.** Distributions of hydrogen bond geometries in small molecule amide•O(sp^3^)–C hydrogen bonds *versus* in the oxyanion holes of TSA-bound serine proteases. The two hydrogen bonds, 1° and 2°, are defined in Fig. 1A. Geometric parameters are defined in fig. S30. For serine protease distributions, structural clans are denoted by colors (red: PA, pink: SB, yellow: SC, green: SE). The blue lines indicate the most probable value from the small molecule distribution and are repeated in each panel for comparison.

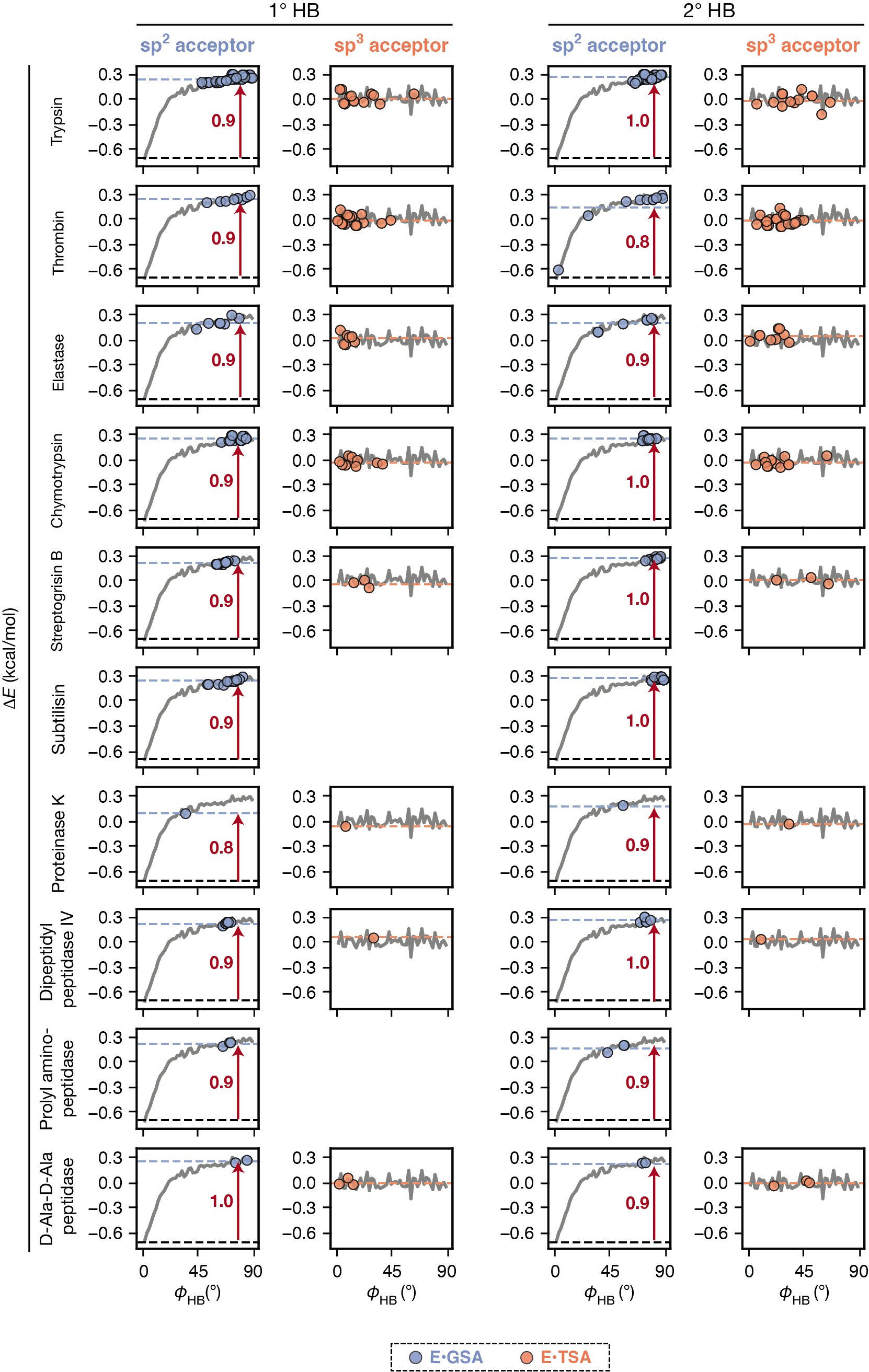

**Fig. S33.** Oxyanion hole hydrogen bond orientations (*ϕ*_HB_, as defined in fig. S30) in serine proteases pseudo-ensembles mapped on the knowledge-based energy functions of *ϕ*_HB_ [Δ*E*(*ϕ*_HB_)]. The two hydrogen bonds, 1° and 2°, are defined in Fig. 1A. The GSA-bound observations (blue circles) were mapped on a Δ*E*(*ϕ*_HB_) derived from amide•carbonyl small molecule interactions (fig. S31); the TSA-bound observations (orange circles) were mapped on a Δ*E*(*ϕ*_HB_) derived from amide•O(sp^3^)–C interactions (fig. S32). Blue and orange dashed lines indicate the mean Δ*E*(*ϕ*_HB_) values calculated for the enzyme GSA-bound and TSA-bound states, respectively, and the black dashed lines indicate the minima of the knowledge-based energy functions.

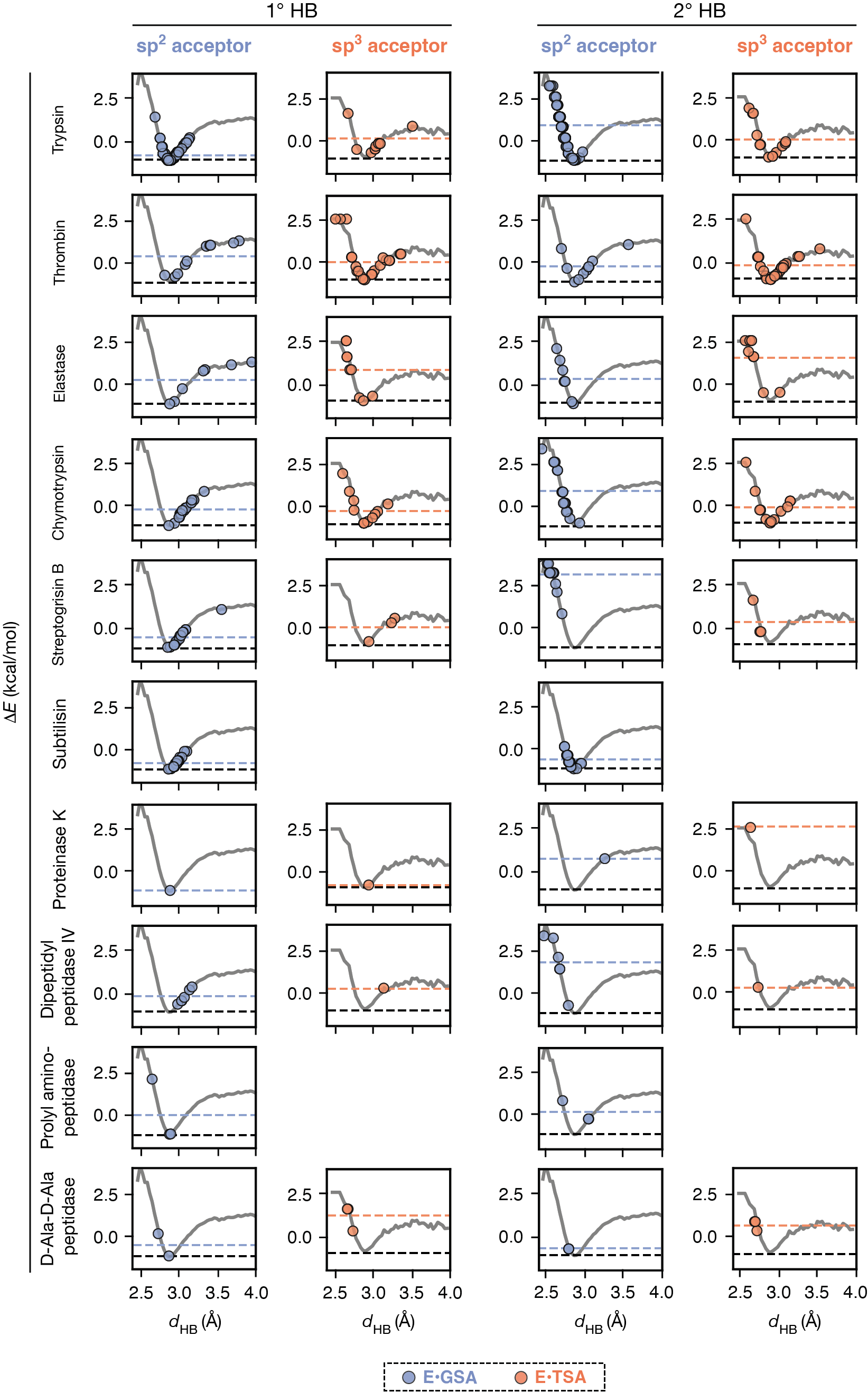

**Fig. S34.** Oxyanion hole hydrogen bond lengths (*d*_HB_, as defined in fig. S30) in serine proteases pseudo-ensembles mapped on the knowledge-based energy functions of *d*_HB_ [Δ*E*(*d*_HB_)]. The two hydrogen bonds, 1° and 2°, are defined in Fig. 1A. The GSA-bound observations (blue circles) were mapped on a Δ*E*(*d*_HB_) derived from amide•carbonyl small molecule interactions (fig. S31); the TSA-bound observations (orange circles) were mapped on a Δ*E*(*d*_HB_) derived from amide•O(sp^3^)–C interactions (fig. S32). Blue and orange dashed lines indicate the mean Δ*E*(*d*_HB_) values calculated for the enzyme GSA-bound and TSA-bound states, respectively, and the black dashed lines indicate the minima of the knowledge-based energy functions.

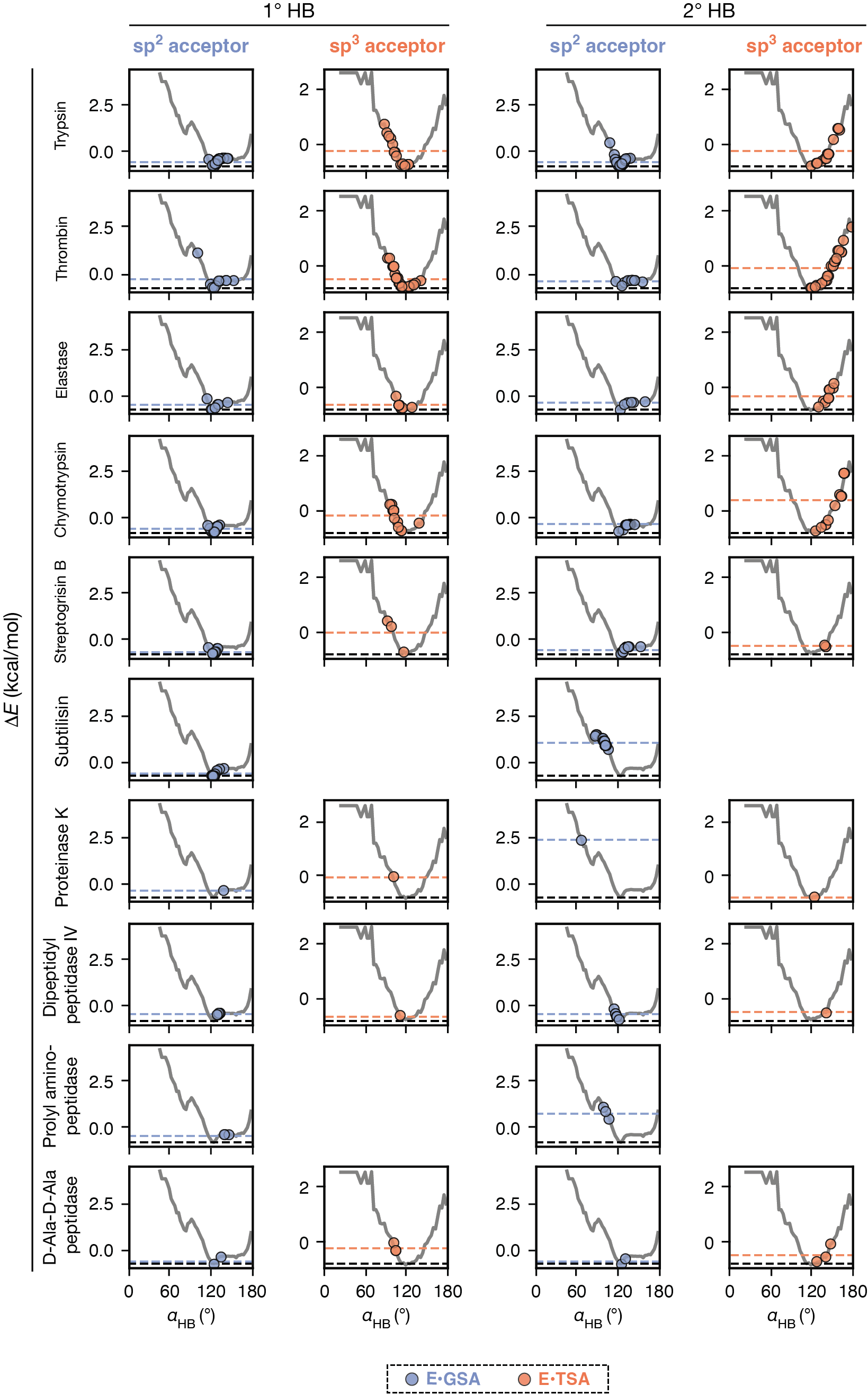

**Fig. S35.** Oxyanion hole hydrogen bond angles (𝛼_HB_, as defined in fig. S30) in serine proteases pseudo-ensembles mapped on the knowledge-based energy functions of 𝛼_HB_ [Δ*E*(𝛼_HB_)]. The two hydrogen bonds, 1° and 2°, are defined in Fig. 1A. The GSA-bound observations (blue circles) were mapped on a Δ*E*(𝛼_HB_) derived from amide•carbonyl small molecule interactions (fig. S31); the TSA-bound observations (orange circles) were mapped on a Δ*E*(𝛼_HB_) derived from amide•O(sp^3^)–C interactions (fig. S32). Blue and orange dashed lines indicate the mean Δ*E*(𝛼_HB_) values calculated for the enzyme GSA-bound and TSA-bound states, respectively, and the black dashed line indicate the minima of the knowledge-based energy functions.

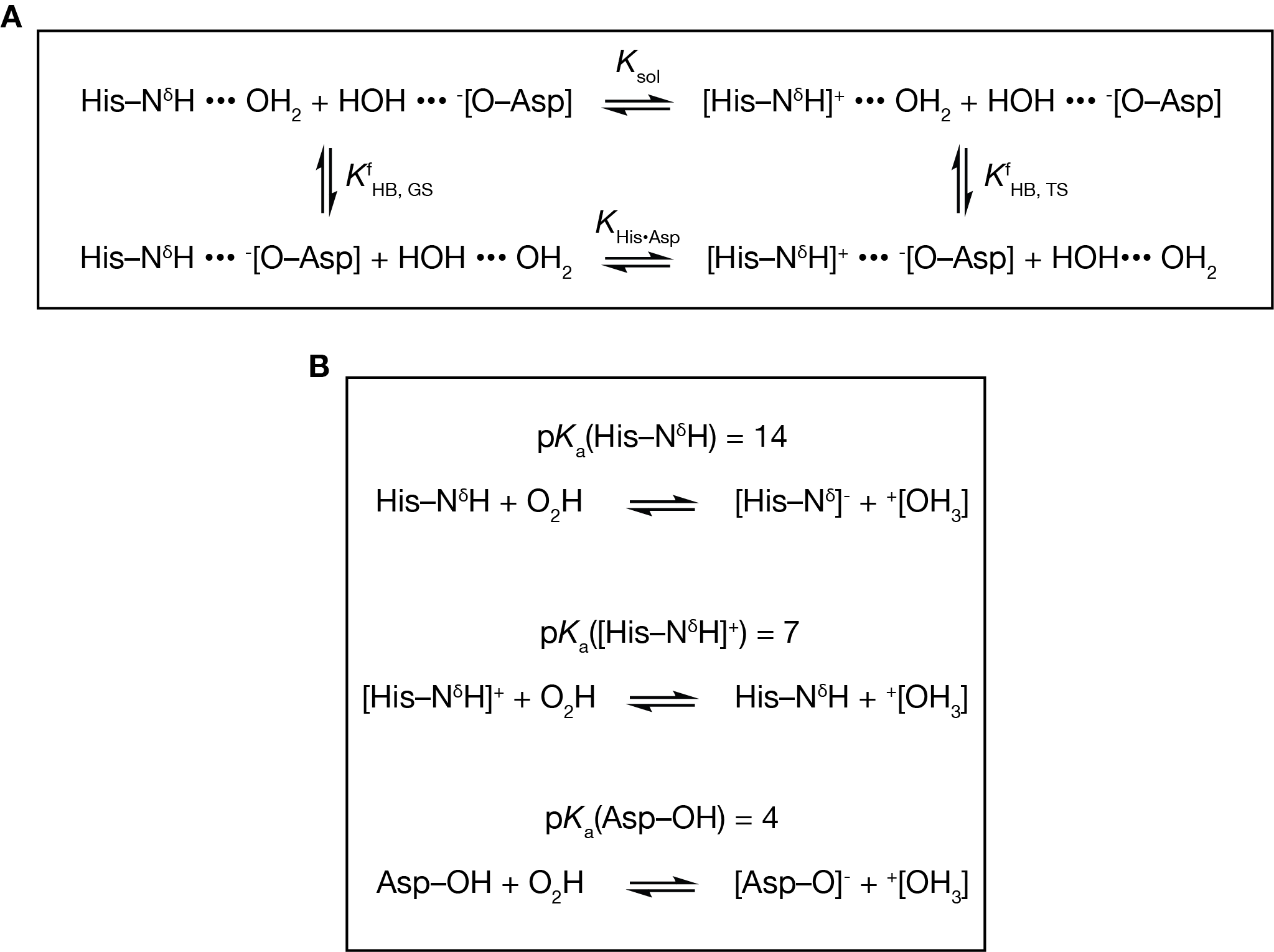
 **Fig. S36.** **(A)** Equilibria of the catalytic triad His•Asp hydrogen bond in the enzymatic reaction *versus* in solution. $K_{His\bullet Asp}$ is the equilibrium constant for the reaction in the presence of the His•Asp hydrogen bond and $K_{sol}$ is that for the aqueous solution reaction without the His•Asp hydrogen bond; $K_{HB, GS}^{f}$ and $K_{HB, TS}^{f}$ are the equilibrium constants for His•Asp hydrogen bond formation in the ground state and in the transition state, respectively. (**B)** Definition of p*K*_a_ values used in the calculations in supplementary text S7.

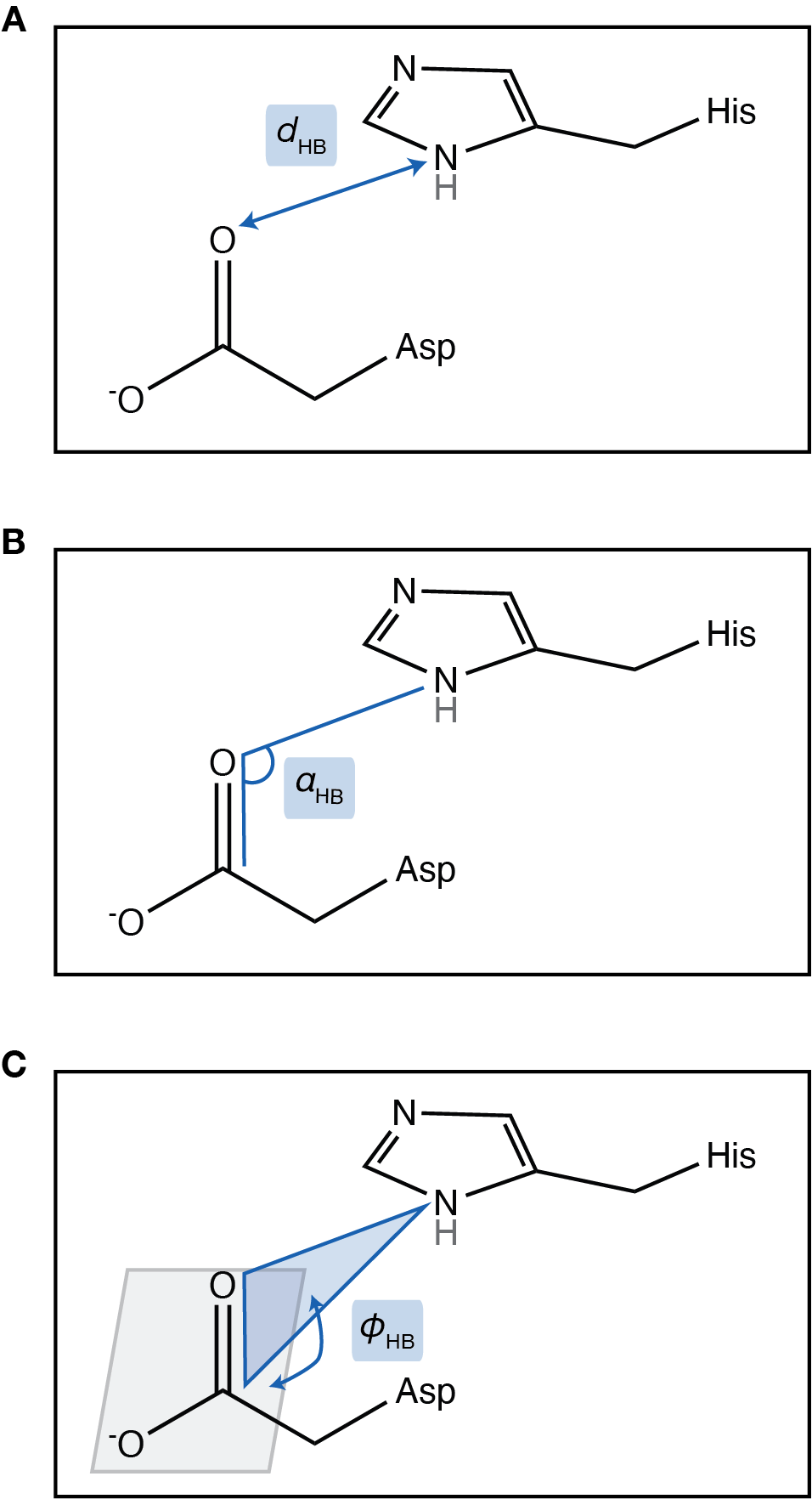

**Fig. S37.** Definition of His•Asp hydrogen bond geometric parameters. **(A)** The hydrogen bond distance (*d*_HB_), defined by the distance between the heavy atoms, i.e. the donor (histidine Nδ1) and the acceptor (aspartate Oδ1). **(B)** The hydrogen bond angle (𝛼_HB_), defined by the angle between the donor, the acceptor, and the carbon atom adjacent to the acceptor (aspartate C𝛾). **(C)** 𝜙_HB_, defined by the dihedral angle between the two planes: (1) the plane defined by the donor (histidine Nδ1), the acceptor (aspartate Oδ1), and the carbon adjacent to the acceptor (aspartate C𝛾) (blue), and (2) the plane of the accepting carbonyl (defined by aspartate Oδ1, C𝛾 and Oδ2; gray).

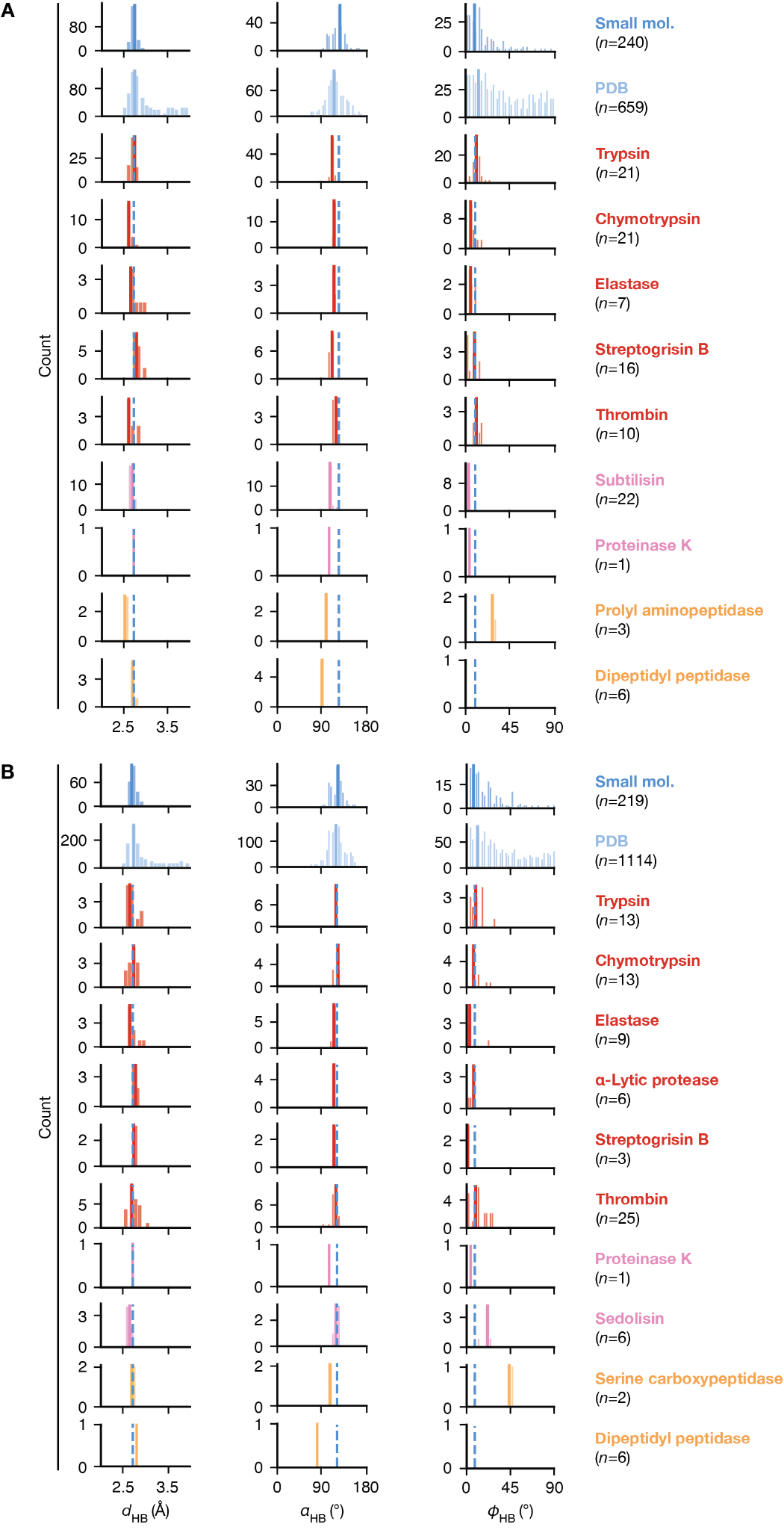

**Fig. S38.** Distributions of the His•Asp hydrogen bond geometries in small molecule structures *versus* in serine proteases. Geometric parameters are defined in fig. S37. **(A)** Comparisons of the ground state distributions, where the small molecule interactions are neutral imidazole•carboxylate hydrogen bonds in the CSD (*104*), the PDB interactions are His•Asp hydrogen bonds taken from *Top2018* (*107*) PDB structures that are crystallized at pH > 8, and the enzymatic distributions are from GSA-bound pseudo-ensembles. For the enzymatic distributions, structural clans are denoted by colors (red: PA, pink: SB, yellow: SC, green: SE). The solid lines of corresponding colors indicate the most probable value of each distribution; the blue dashed lines indicate the most probable value from the small molecule distribution and are repeated in each panel for comparison. **(B)** Comparisons of the transition state distributions, where the small molecule interactions are protonated imidazole•carboxylate hydrogen bonds in the CSD (*104*), the PDB interactions are His•Asp hydrogen bonds taken from high quality structures in *Top2018* (*107*) crystallized at pH < 6, and the enzymatic distributions are from TSA-bound pseudo-ensembles.

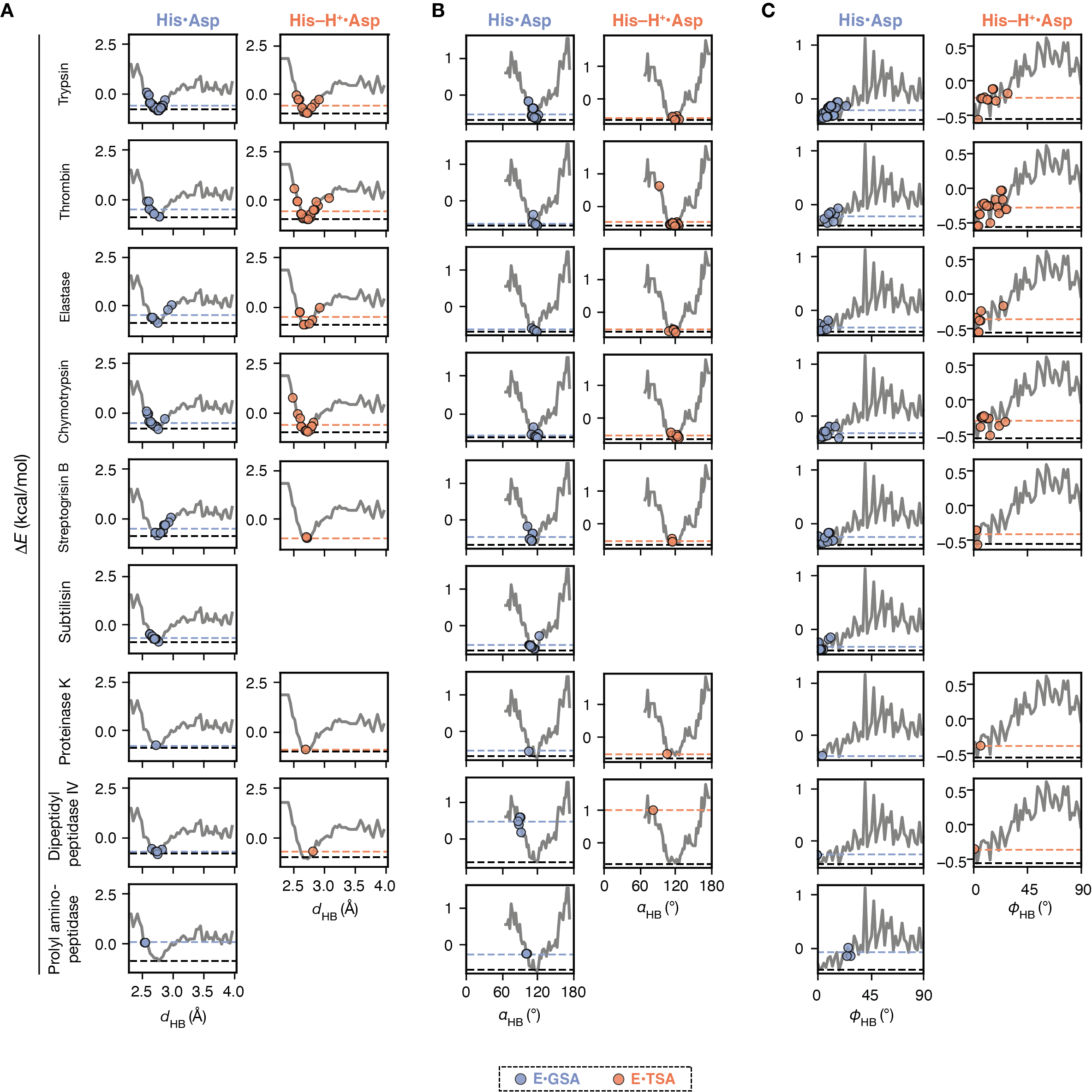

**Fig. S39.** His•Asp hydrogen bond geometries, **(A)** *d*_HB_, **(B)** 𝛼_HB_, and **(C)** 𝜙_HB_ (defined in fig. S37) in serine proteases pseudo-ensembles, mapped on their corresponding knowledge-based energy functions. The energy functions are derived from the PDB distributions of His•Asp hydrogen bonds shown in fig. S38 using structures crystallized at pH > 8 (His•Asp) and pH < 6 (His–H^+^•Asp). The PDB distributions are used instead of small molecule distributions in this case because the sample sizes are larger, and the distributions are highly similar to the small molecule ones. Observations from GSA-bound pseudo-ensembles are shown as blue circles; those from TSA-bound pseudo-ensembles are shown as orange circles. Blue and orange dashed lines indicate the mean energy values calculated for the enzyme GSA-bound and TSA-bound states, respectively, and the black dashed line indicate the minima of the knowledge-based energy functions.

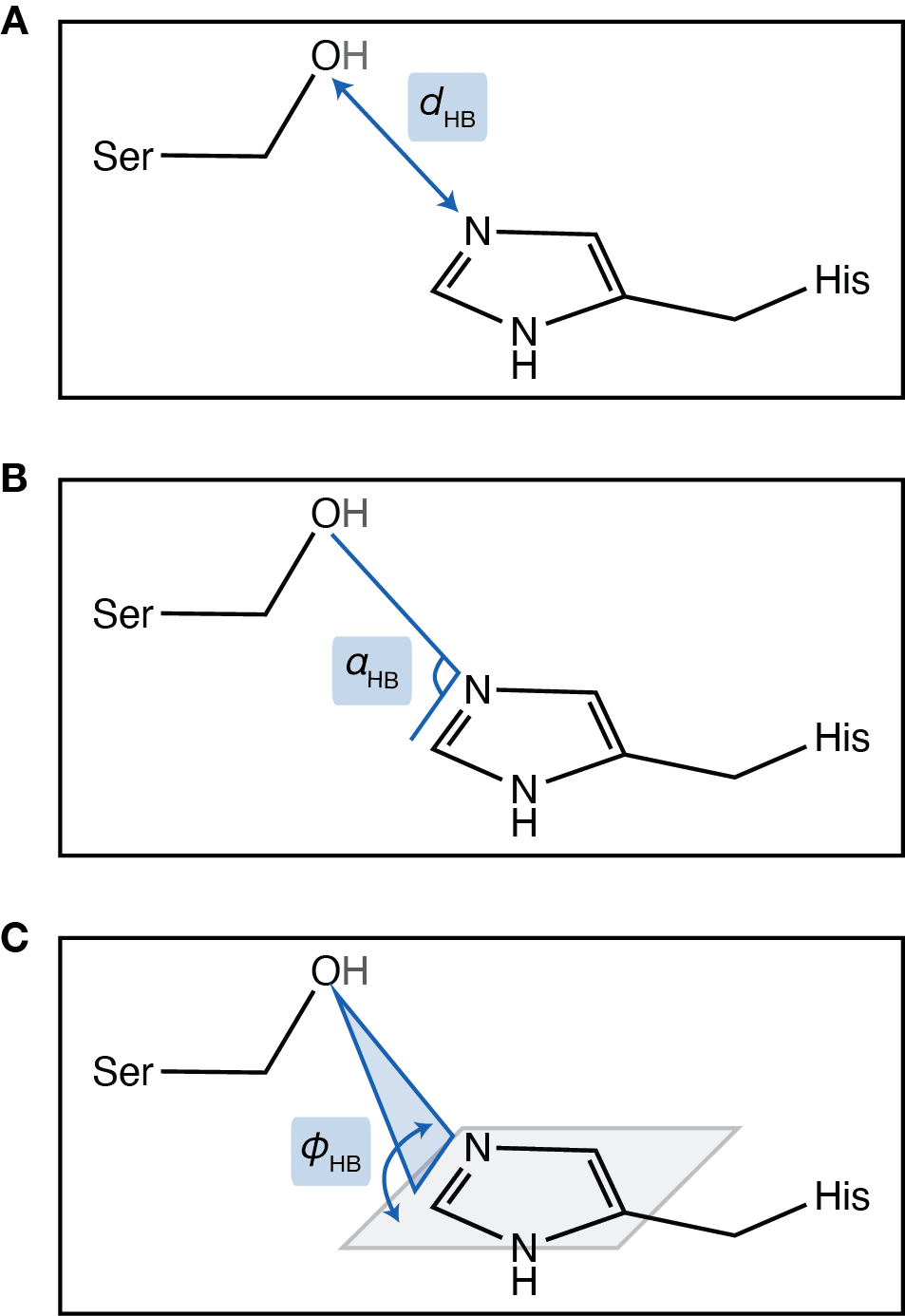

**Fig. S40.** Definition of His•Ser hydrogen bond parameters. **(A)** The hydrogen bond distance (*d*_HB_), defined by the distance between the heavy atoms, i.e. the donor (histidine Nε2) and the acceptor (serine O𝛾). **(B)** The hydrogen bond angle (𝛼_HB_), defined by the angle between the donor (histidine Nε2), the acceptor (serine O𝛾), and the carbon atom adjacent to the acceptor (histidine Cε1). **(C)** 𝜙_HB_, defined by the dihedral angle between the two planes: (1) the plane defined by the donor (histidine Nε2), the acceptor (serine O𝛾), and the carbon atom adjacent to the acceptor (histidine Cε1) (blue), and (2) the plane of the accepting carbonyl (defined by the histidine Nε2, Cε1 and Nδ1; gray).

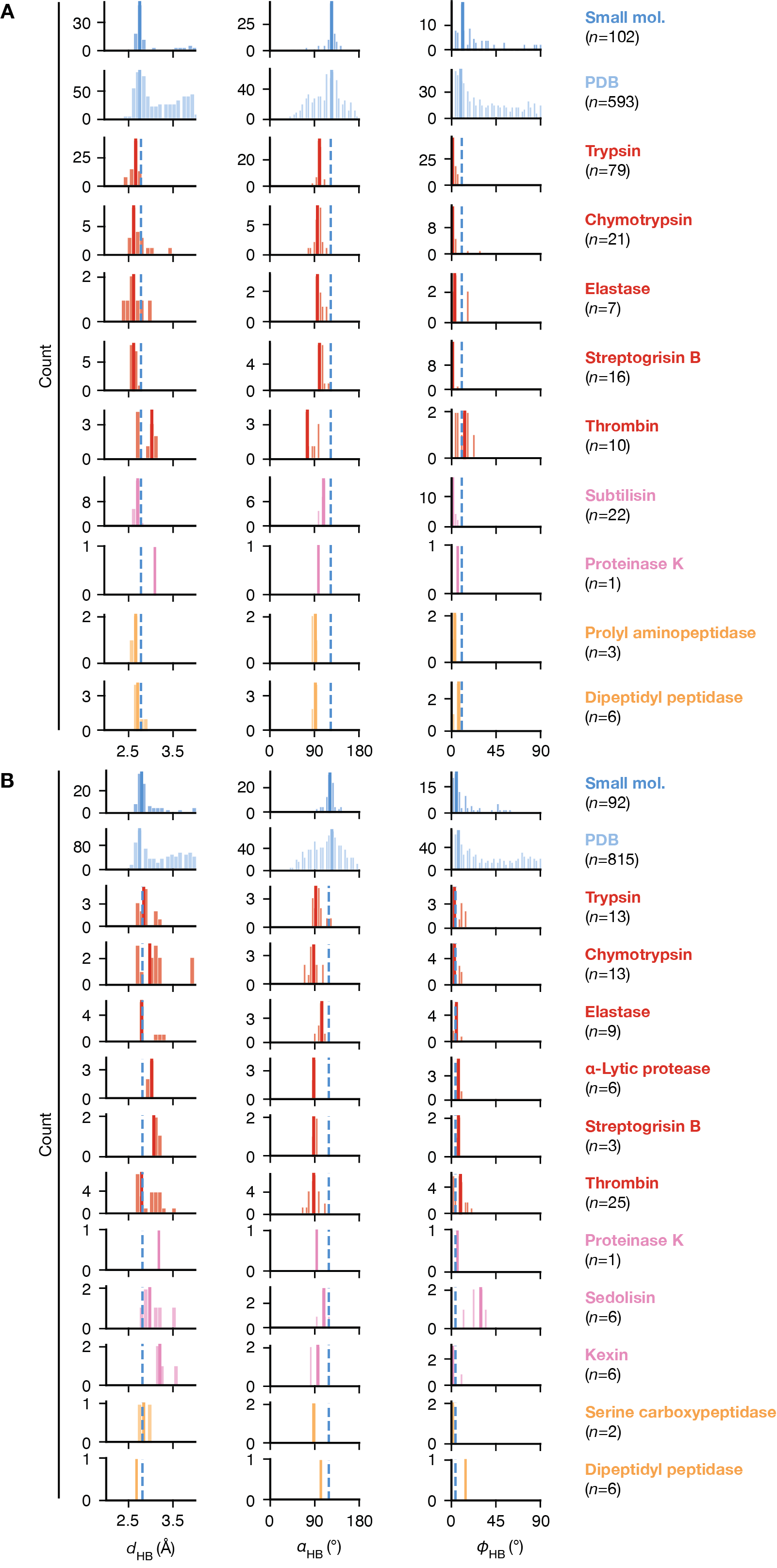

**Fig. S41.** Distributions of the His•Ser hydrogen bond geometries in small molecule structures *versus* in serine proteases. Geometric parameters are defined in fig. S40. **(A)** Comparisons of the ground state distributions, where the small molecule interactions are neutral imidazole•hydroxyl hydrogen bonds in the CSD (*104*), the PDB interactions are His•Ser hydrogen bonds taken from *Top2018* (*107*) PDB structures that are crystallized at pH > 8, and the enzymatic distributions are from GSA-bound pseudo-ensembles. For the enzymatic distributions, structural clans are denoted by colors (red: PA, pink: SB, yellow: SC, green: SE). The solid lines of corresponding colors indicate the most probable value of each distribution; the blue dashed lines indicate the most probable value from the small molecule distribution and are repeated in each panel for comparison. **(B)** Comparisons of the transition state distributions, where the small molecule interactions are protonated imidazole•ether or hydroxyl hydrogen bonds in the CSD (*104*), the PDB interactions are His•Ser hydrogen bonds taken from *Top2018* (*107*) PDB structures that are crystallized at pH < 6, and the enzymatic distributions are from TSA-bound pseudo-ensembles.

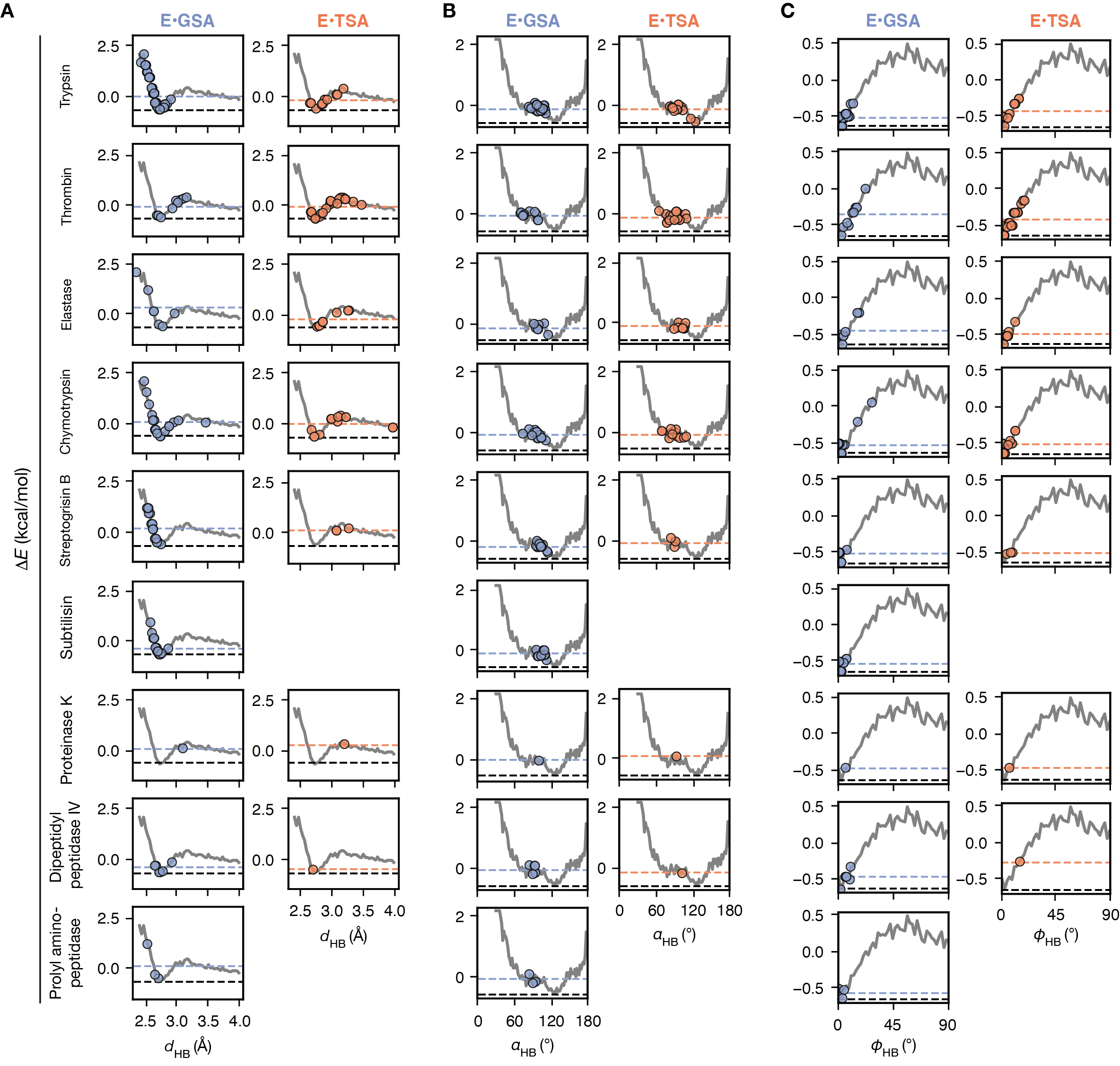

**Fig. S42.** His•Ser hydrogen bond geometries, **(A)** *d*_HB_, **(B)** 𝛼_HB_, and **(C)** 𝜙_HB_ (defined in fig. S40) in serine proteases pseudo-ensembles, mapped on their corresponding knowledge-based energy functions. The energy functions are derived from the PDB distribution of His•Ser hydrogen bonds collected from *Top2018* (*n* = 3408). PDB structures are used instead of the small molecule distributions in this case as the larger sample sizes provided smoother energy functions and the distributions are highly similar to the small molecule ones and across pH (fig. S41). Observations from GSA-bound pseudo-ensembles are shown as blue circles; those from TSA-bound pseudo-ensembles are shown as orange circles. Blue and orange dashed lines indicate the mean energy values calculated for the enzyme GSA-bound and TSA-bound states, respectively, and the black dashed lines indicate the minima of the knowledge-based energy functions.

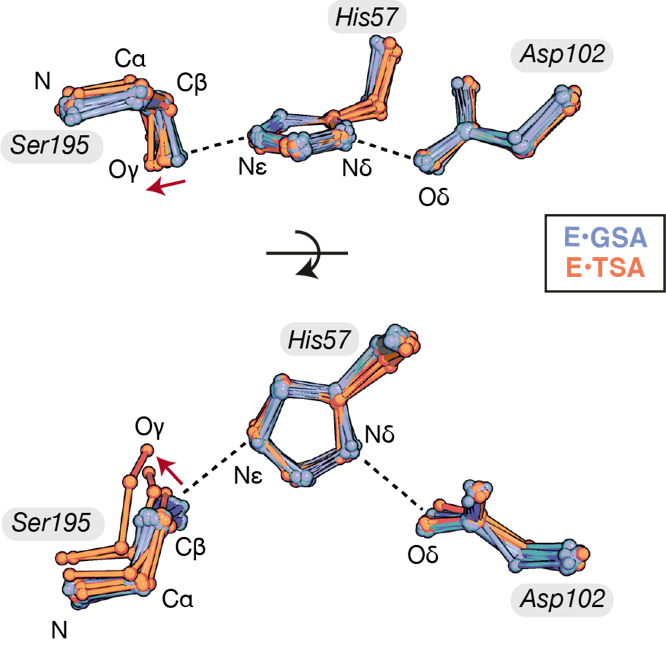

**Fig. S43**. Trypsin GSA- and TSA-bound pseudo-ensembles [blue (*n* = 79) and orange (n = 13), respectively] aligned on the catalytic triad sidechain atoms. The catalytic serine rotates from the GSA-bound to the TSA-bound state, whereas the histidine and aspartate maintained their positions. The serine rotation lengthens the His•Ser hydrogen bond (red arrow), with no significant accompanying change in the His•Asp hydrogen bond.

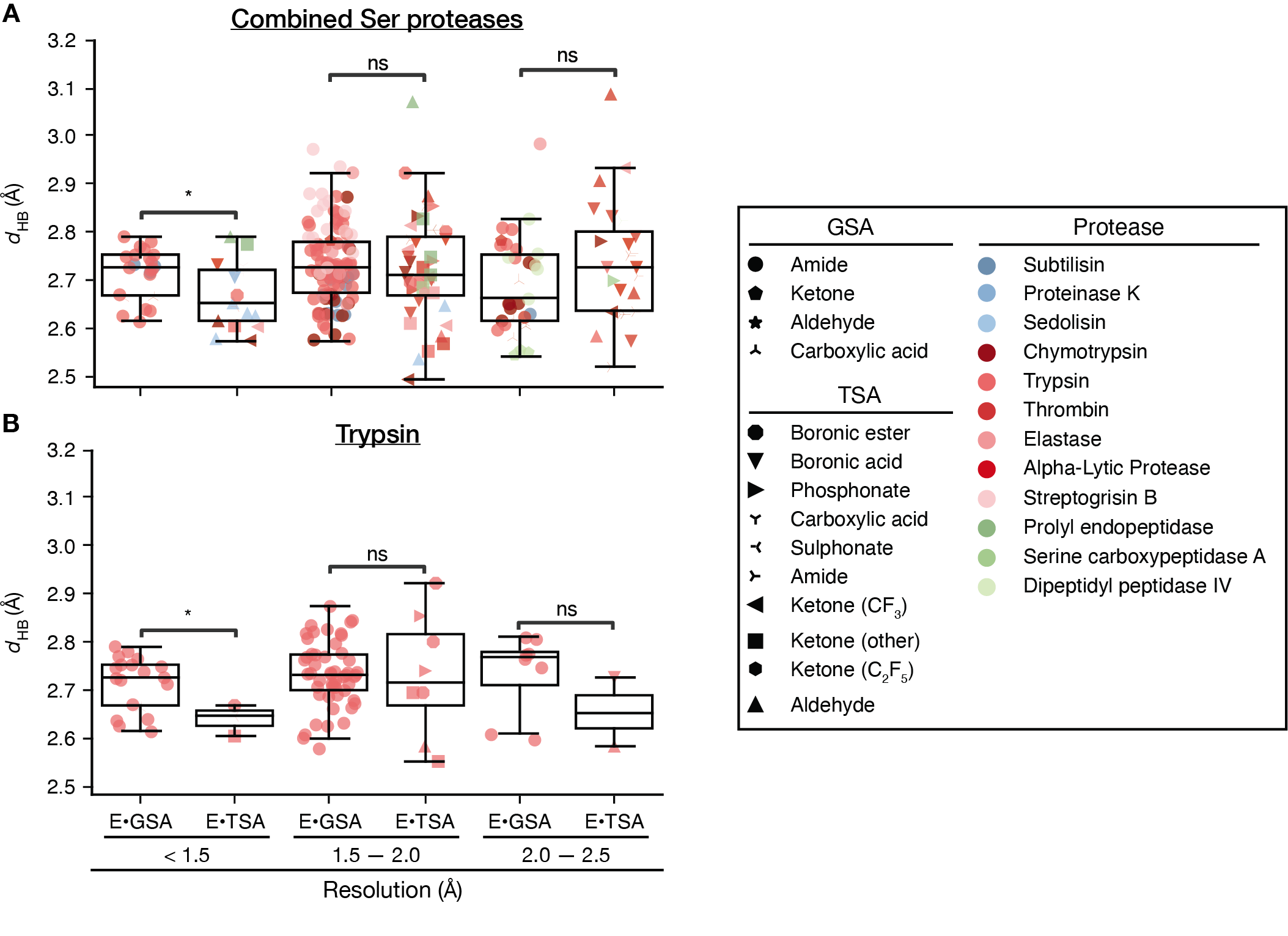

**Fig. S44**. His•Asp hydrogen bond lengths observed in structures of different resolution ranges for combined serine proteases **(A)** and trypsin only **(B)**; the two panels share the same x-axis. * indicate significant changes (*p*-value < 0.05). The list of GSA and TSAs are defined in fig. S5. At the high-resolution range (<1.5 Å), there is a significant difference of ~0.1 Å between the His•Asp hydrogen bond lengths in GSA-bound and TSA-bound states.
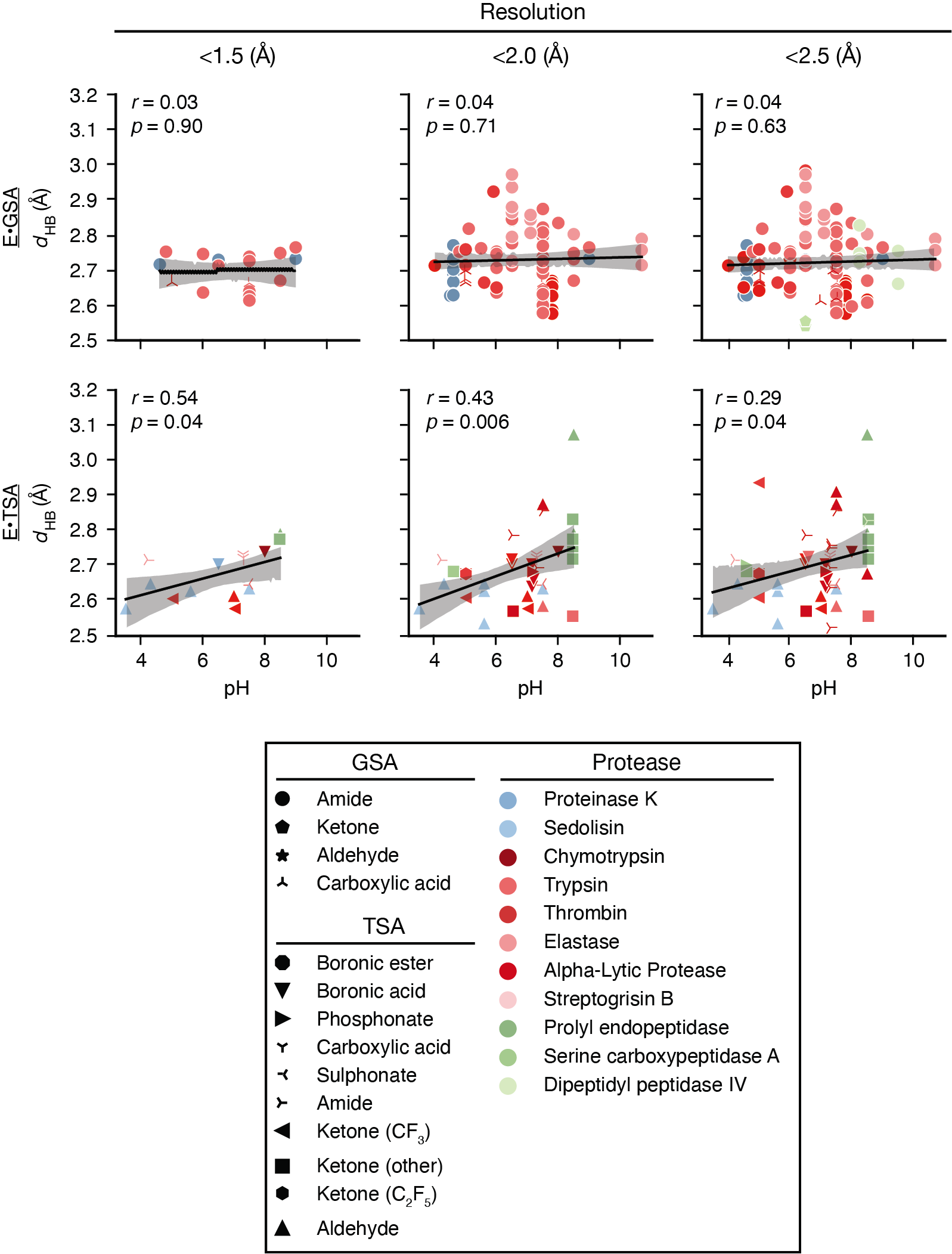

**Fig. S45.** His•Asp hydrogen bond length *versus* crystallographic pH in GSA-bound (upper) and TSA-bound (lower) pseudo-ensembles for subsets of data at different resolution ranges. The black line indicates the linear fit to the data and the gray shaded area indicates 95% confidence intervals (*r*, Pearson coefficient; *p*, *p*-value of the correlation). The GSA and TSAs listed are defined in fig. S5. There is no apparent increase in lengths at higher pH for GSA-bound structures; for TSA-bound structures, an apparent increase is observed, but the lengths observed at each pH are associated with large variations, and the correlations are not significant (except for the subset of structures at <2.0 Å resolution).

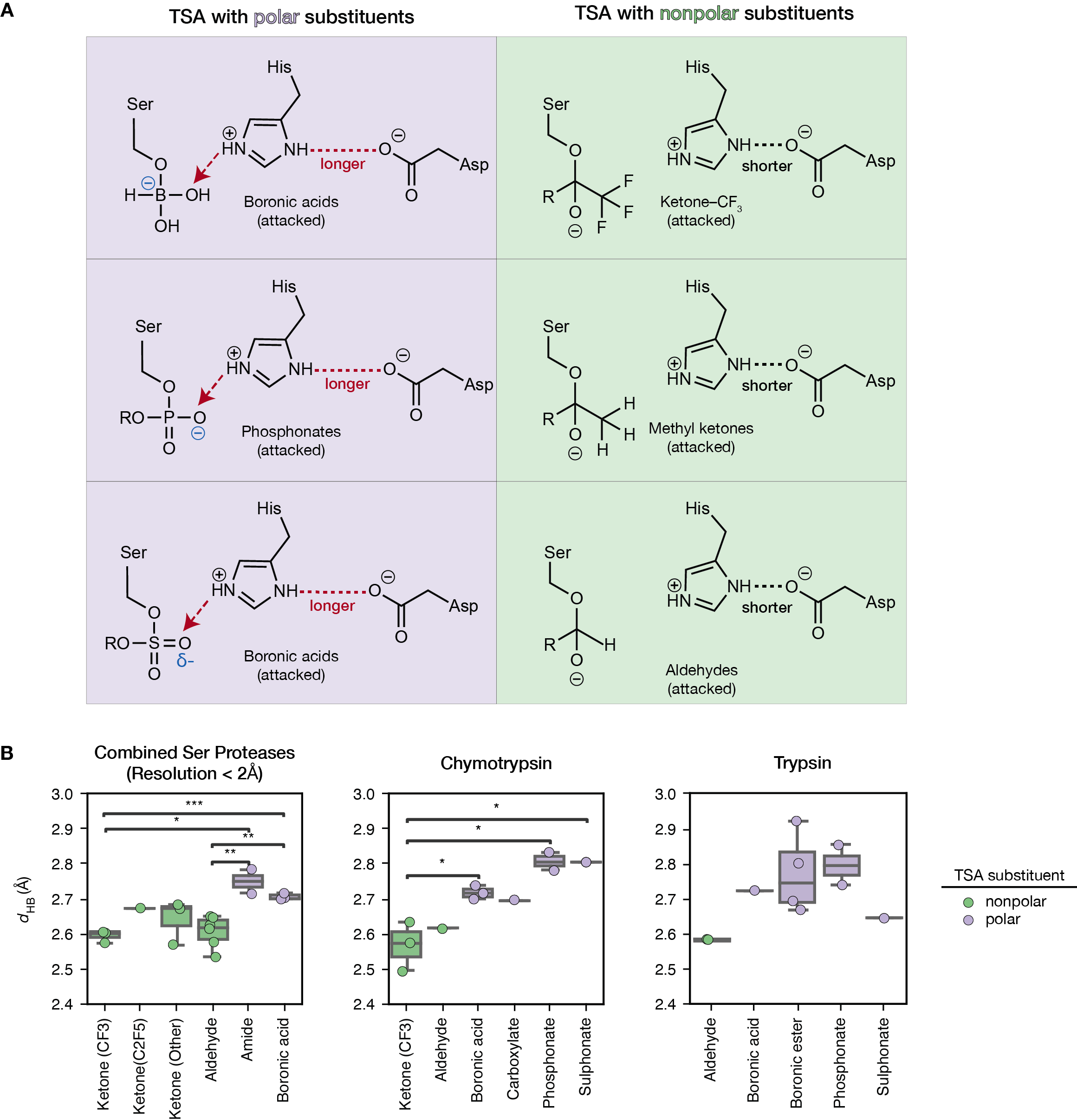

**Fig. S46.** Comparisons of His•Asp hydrogen bond lengths when serine proteases are bound with different type of TSAs. **(A)** The TSAs with polar *versus* nonpolar substituents may result in different His•Asp lengths when bound to the active site, as those with polar substituents can form hydrogen bonds with the catalytic histidine, whereas those with nonpolar substituents cannot. **(B)** His•Asp hydrogen bond lengths (*d*_HB_) in combined serine protease structures at high resolutions (< 2 Å, left), in chymotrypsin (middle) and in trypsin (right) when the enzymes are bound with different types of TSAs (defined in fig. S5). TSAs with nonpolar substituents are colored in greens and those with polar substituents colored in purples. Asterisk symbols indicate significance levels, with one asterisk corresponding to *p*-value $\leq$ 0.05 and with any number (*n*) of asterisks larger than one corresponding to $1\times{10}^{-n}\leq$ *p*-value$\leq1\times{10}^{-(n+1)}$.

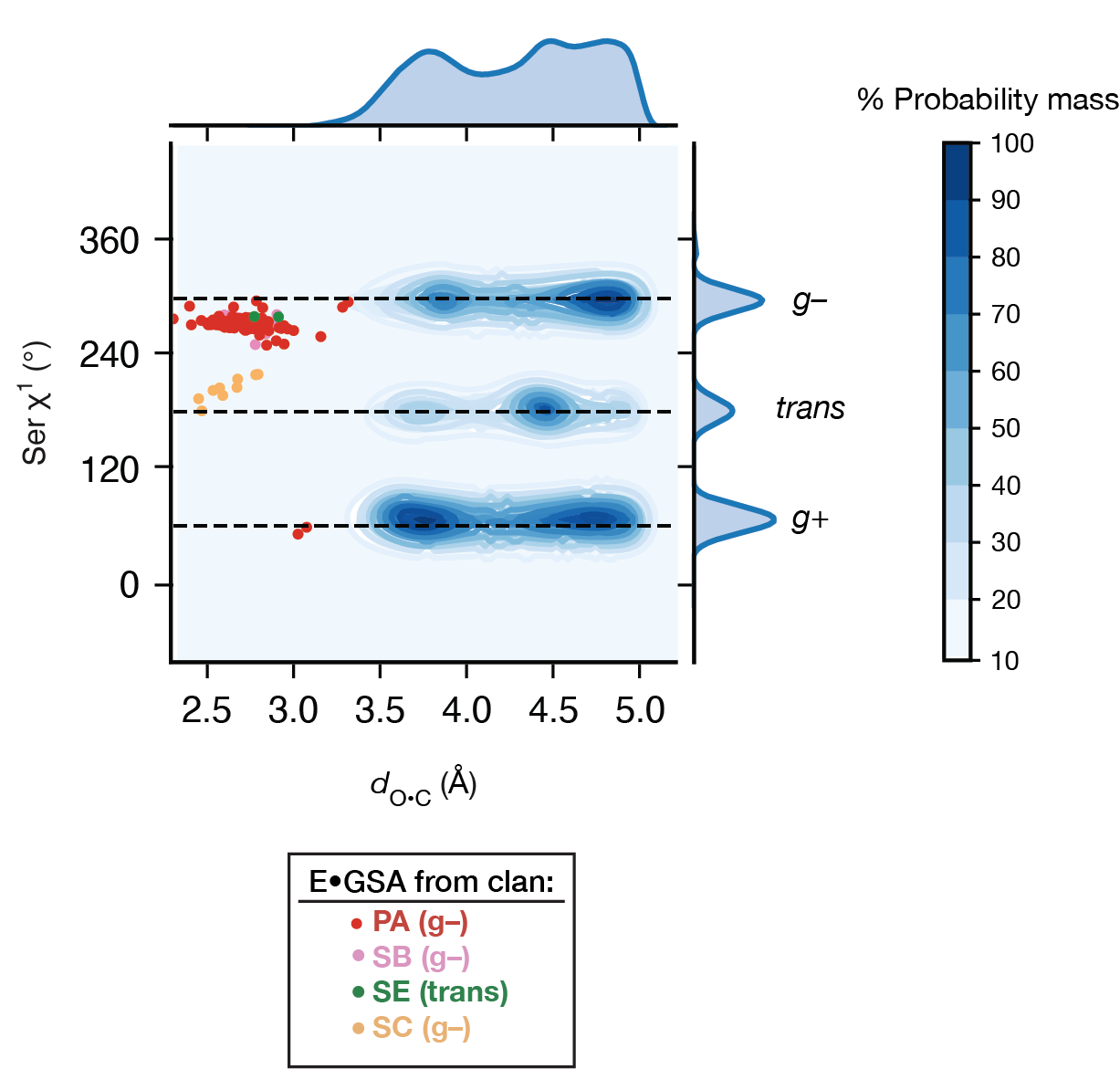

**Fig. S47.** Two-dimensional knowledge-based distribution of serine χ^1^ angles *versus* the distance between the serine Oγ and the carbon from a nearby amide group (*d*_O•C_), built from 162,706 interactions collected from 13,149 high quality PDB structures from the *Top2018* dataset (*107*). The dataset was searched for serine residues in proximity to a backbone amide carbon with *d*_O•C_ $\leq$ 5 Å; the population where the serine Oγ is hydrogen bonded to the amide oxygen (O) was removed by selecting the instances where *d*_O•C_ (the interaction of interest) is less than *d*_O•O_ (the putative hydrogen bond distance). The map is contoured and colored by cumulative probability mass (in %) with increased intensity of the blue color indicating higher probability. Observations from serine protease GSA-bound pseudo-ensembles were plotted on the map as dots and colored by their structural clans (PA: red; SB: pink; SE: green, SC: yellow). For clan PA, SB and SE, the reactive rotamer state is *gauche–* (*g–*); for clan SC, the reactive rotameric state is *trans*. The three local minima of serine χ^1^ angles, *g–, g+ and trans* are indicated by black dashed lines on the map. The three local minima have the same χ^1^ values across *d*_O•C_, as expected for energetically independent parameters.

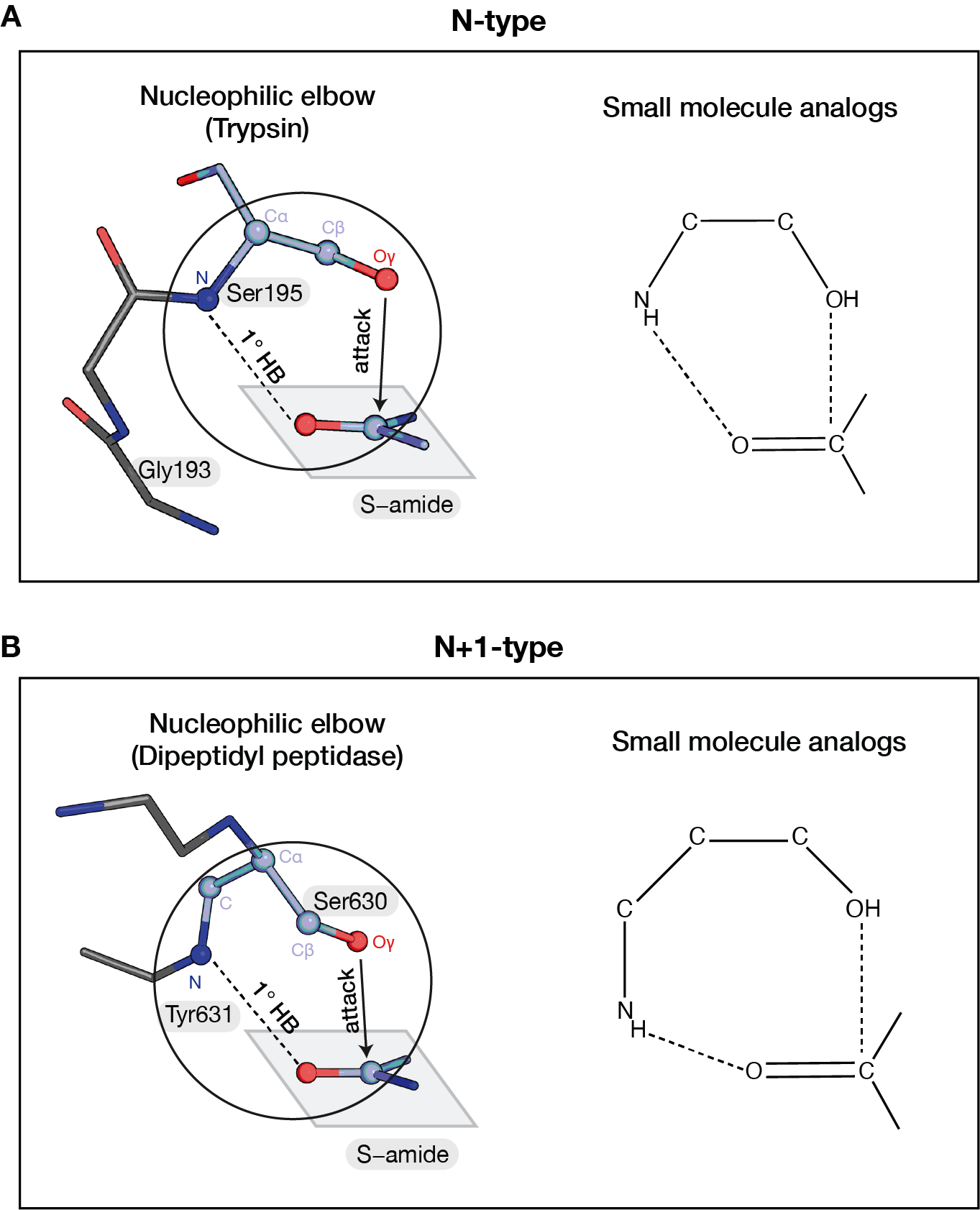

**Fig. S48.** The nucleophilic elbow geometries. **(A)** The N-type nucleophilic elbow in GSA-bound trypsin (left, PDB: 3M7Q) and analogous small molecule interactions (right). **(B)** The N+1-type nucleophilic elbow in GSA-bound dipeptidyl peptidase (left, PDB: 1WCY) and analogous small molecule interactions (right). For enzyme nucleophilic elbows, the atoms consisting of the nucleophilic elbow “ring” are shown as spheres; for the small molecule structures, corresponding atoms are shown, and additional hydrogen atoms and substituents of the molecules are omitted. The small molecule structures were used to construct the knowledge-based distributions shown in Fig. 6D and figs. S49 to 50. See table S6 for entries used for the small molecule search.

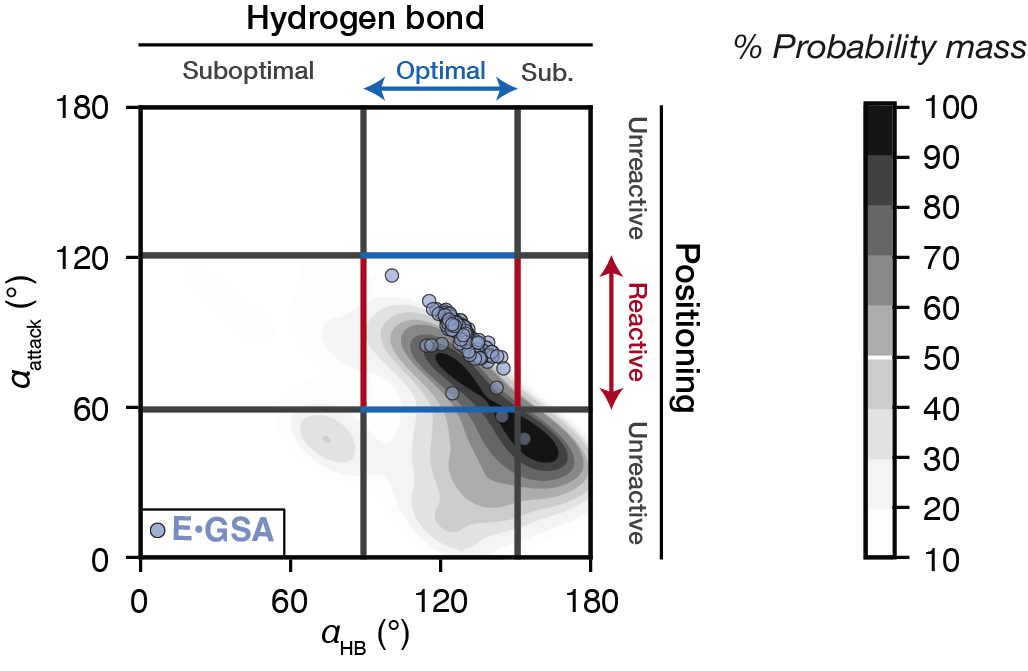

**Fig. S49.**  Knowledge-based conformational map for hydrogen bond angles (*α*_HB_, defined in Fig. 6B) and nucleophile•amide (*α*_attack_, defined in Fig. 2A) in small molecule N-type nucleophilic-elbow-like motifs. (*n* = 1294). The intensity of gray indicates the cumulative probability mass (%) at the contour level. Blue circles indicate the observed *α*_HB_ and *α*_attack_ in GSA-bound serine proteases with N-type nucleophilic elbows. The region of optimal hydrogen bond angles (blue arrow) is defined as 90° $\leq$ *α*_HB_ $\leq$150°, and the region of optimal attack angles (red arrow) is defined as 60° $\leq$*α*_attack_ $\leq$120°, each allows a 30° deviation from the most favorable angle (*α*_HB_ = 120° and *α*_attack_ = 90°, Fig. 6B). Favorable regions for the *α*_HB_ and *α*_attack_ angles can be mutually achieved, and they match the angles observed on GSA-bound serine proteases, suggesting no geometric constraints for α_HB_ and α_attack_.

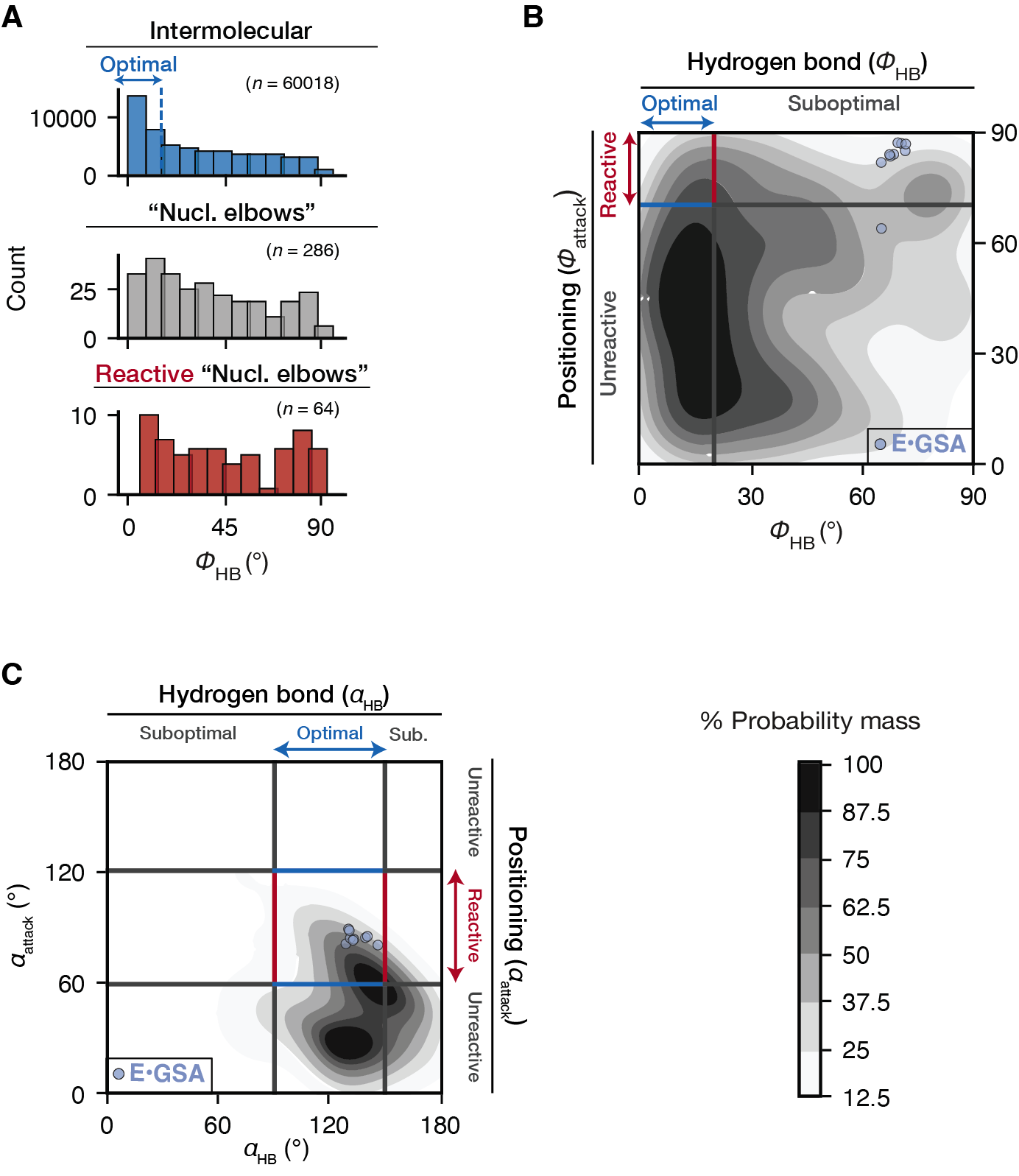
**­­**

**Fig. S50.** Conformational landscape of the N+1-type nucleophilic elbow (defined in fig. S48)­­­. **(A)** Distributions of $\phi$_HB_ in intermolecular amide•carbonyl hydrogen bonds (blue), small molecule N+1-type nucleophilic-elbow-like structures (gray), and its subset where the reactant orientations are reactive (defined as $\phi$_attack_ ≥ 70°, red). The blue arrow indicates the region of $\phi$_HB_ $\leq$ 20° and is assigned as “optimal”. **(B)** Knowledge-based conformational map for the hydrogen bond orientations ($\phi$_HB_) and nucleophile•amide attack orientation ($\phi$_attack_) in small molecule N+1-type nucleophilic-elbow-like motifs (*n* = 286). (B) and (C) share the same color scale with the intensity of gray indicates the cumulative probability mass (%) at the contour level. Blue circles indicate the observed $\phi$_HB_ and $\phi$_attack_ in GSA-bound serine proteases with N+1-type nucleophilic elbows. **(C)** Knowledge-based conformational map for the hydrogen bond angle (*α*_HB_) and nucleophile•amide attack angle (*α*_attack_) in small molecule N+1-type nucleophilic-elbow-like motifs, in the same format as (B). The region of optimal hydrogen bond angles (blue arrow) is defined as 90°$\leq$*α*_HB_$\leq$150°, and the region of optimal attack angles (red arrow) is defined as 60°$\leq$*α*_attack_$\leq$120°, each allows a 30° deviation from the most favorable angle (*α*_HB_ = 120° and *α*_attack_ = 90°, Fig. 6B).

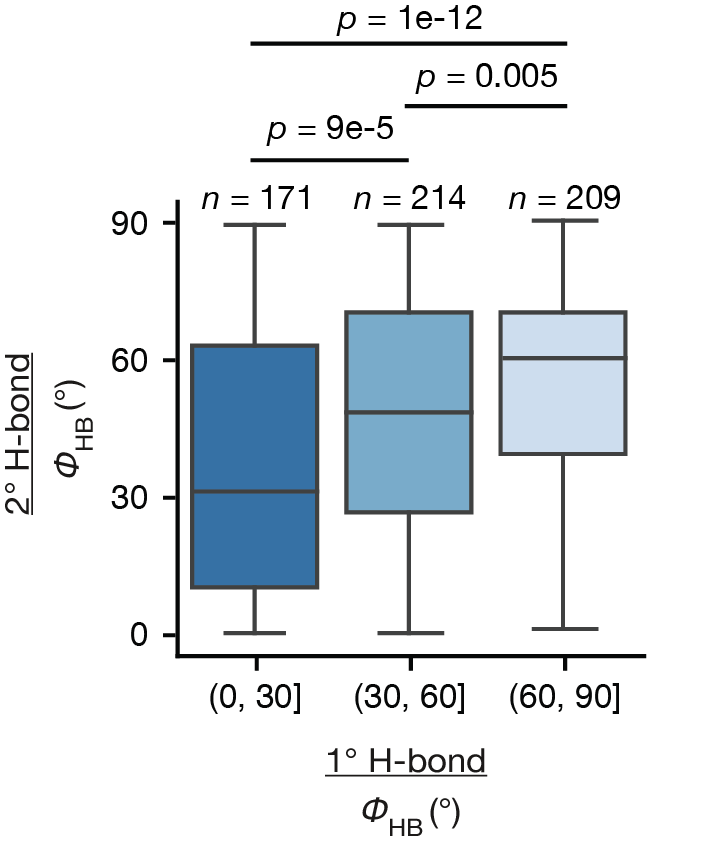

**Fig. S51.** Hydrogen bond orientations ($\phi$_HB_) in small molecule structures where a carbonyl C=O group accepts hydrogen bonds from two amide groups (1° and 2° H-bond). The orientations of the two hydrogen bonds are coupled such that a non-planar 1° hydrogen bond is associated with a non-planar 2° hydrogen bond, and *vice versa.* *P*-values are obtained from *t*-tests.

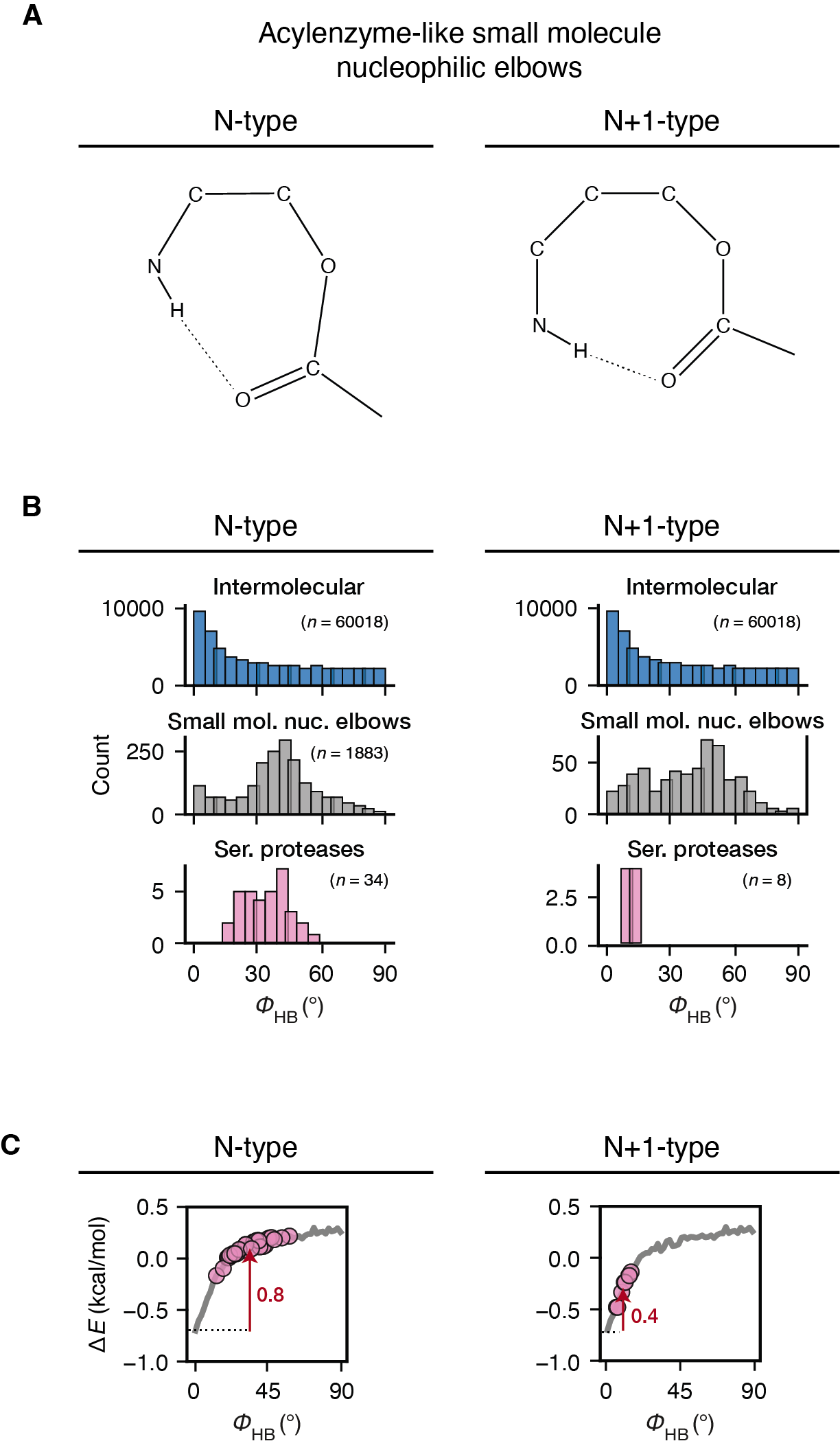

**Fig. S52.** Nucleophilic elbows in the acylenzyme state have destabilized hydrogen bonds. **(A)** Configurations of the N- and N+1-type nucleophilic elbow in the acylenzyme state. The small molecule structures were used to construct the knowledge-based distributions shown in B in gray. **(B)** Distributions of $\phi$_HB_ in intermolecular amide•carbonyl hydrogen bonds (blue), small molecule nucleophilic-elbow-like structures (gray) for N-type acylenzyme-like (left) and for N+1-type acylenzyme-like (right), and in serine protease acylenzyme ensembles (pink). **(C)** $\phi$_HB_ in the serine protease acylenzyme pseudo-ensembles mapped on the knowledge-based energy function for $\phi$_HB_ (also shown in Fig. 5F, sp^2^). Dashed lines indicate the energy minimum; red arrows and values indicate the amount of destabilization.

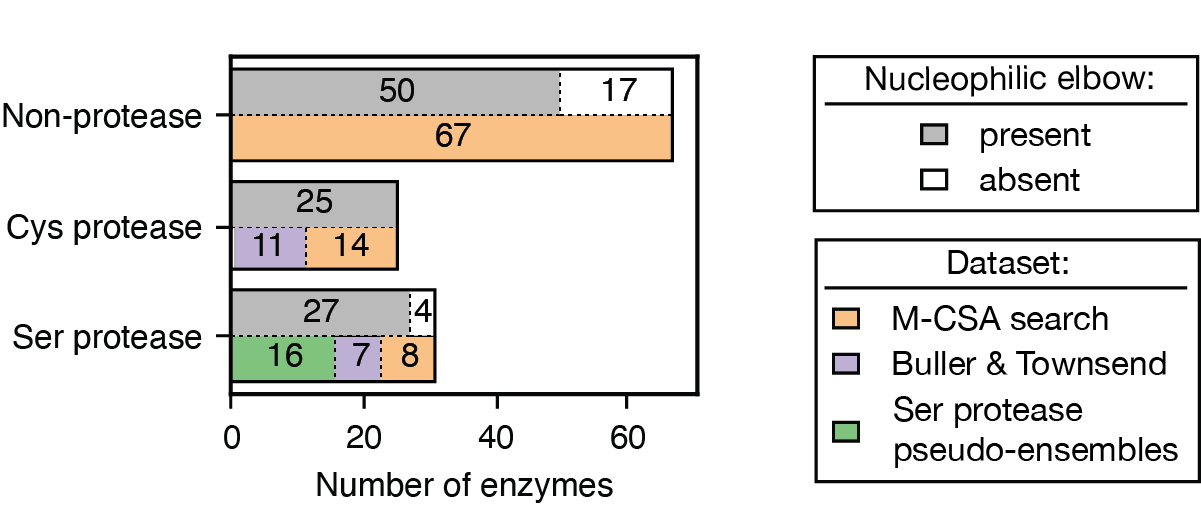

**Fig. S53.** Number of enzymes found from different datasets in the search for nucleophilic elbow-bearing enzymes. The upper bars for each group indicate the number of enzymes with and without nucleophilic elbows (gray and white, respectively); the lower bars indicate the source of the collected enzymes. Serine proteases investigated in this work are colored the green (table S3), serine and cysteine proteases collected by Buller and Townsend (table S22) (*115*) are colored in purple, and the enzymes collected from M-CSA (excluding enzymes repetitive with the other two datasets; table S21) are colored in orange.

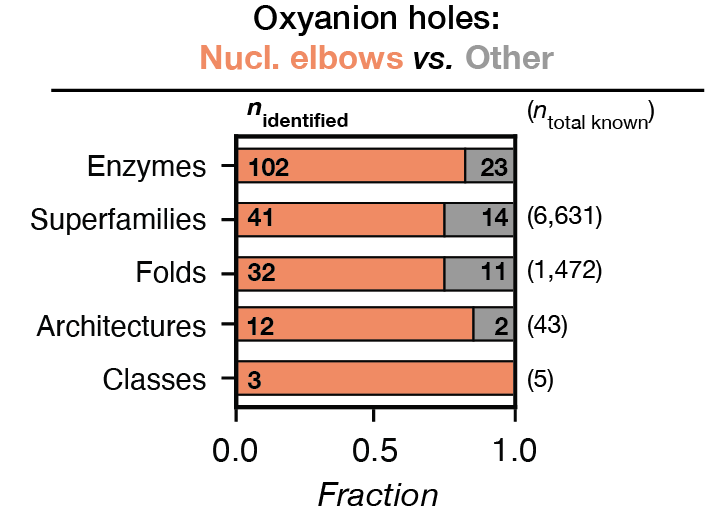

**Fig. S54.** Most enzymes containing oxyanion holes are found to have nucleophilic elbows (orange) rather than other types of oxyanion holes (gray). The set of 125 enzymes from fig. 53 was classified by the hierarchy of structural classification of the known protein structural space (classes > architectures > folds > superfamilies > enzymes). The structural hierarchy was taken from the CATH database (*92*) and was constructed from 536,613 structural domains in the PDB. This set of 125 enzymes consists of all serine proteases in this work (*n* = 16), additional serine and cysteine proteases from Buller and Townsend (*n* = 18) (*116*) and other additional enzymes from our M-CSA search (*n* = 91).

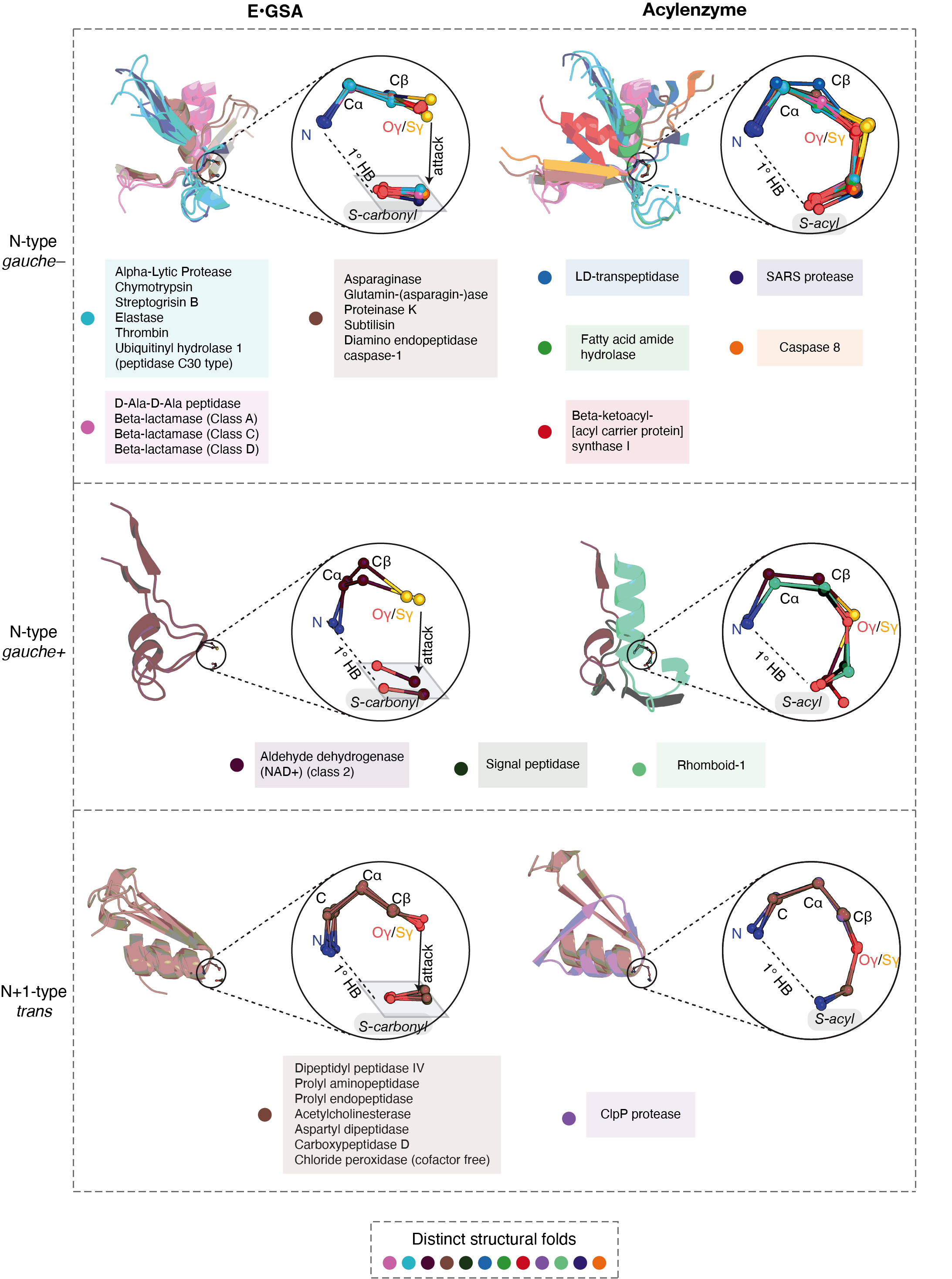

**Fig. S55.** Nucleophilic elbows can be inserted in different structural folds within various types of secondary structural elements, suggesting high evolvability. Shown are local alignments of nucleophilic elbows from different enzymes performing nucleophilic addition on carbonyl compounds (see table S24). The enzymes are separated into groups by their nucleophilic elbow configuration (N- and N+1-type), ligand-bound state (GSA-bound and acylenzyme), and the rotameric state of the nucleophilic residue (*gauche–*, *gauche+* and *trans*). For each group, one structure with the lowest alignment RMSD with the reference structure (table S24) from each enzyme is shown; the structural region where the nucleophilic elbow is located ($\pm$10 residues) is shown as cartoons on the left, and the nucleophilic elbow atoms are shown on the right in the circle (“S-carbonyl” indicates the substrate electrophilic carbonyl plane and “S-acyl” indicates the ligand acyl group that is connected to the enzyme). The nucleophilic elbows are colored by the structural folds of their corresponding enzymes.

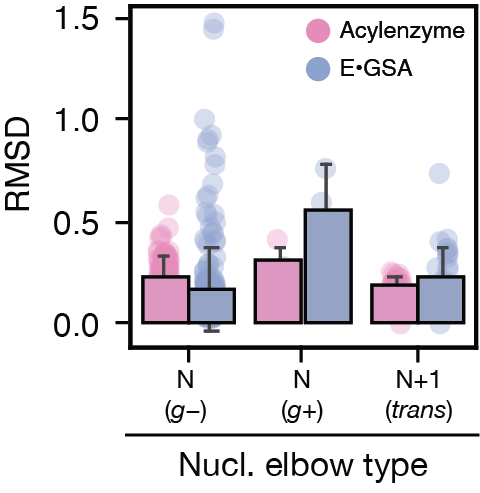

**Fig. S56.** Distributions of pairwise RMSD values obtained by aligning nucleophilic elbow atoms from enzymes that perform nucleophilic addition on carbonyl compounds. The enzymes are separated into groups by their nucleophilic elbow configuration (N- and N+1-type), ligand-bound state (GSA-bound: blue; acylenzyme: pink), and the rotameric state of the nucleophilic residue (*gauche–*, *gauche+* and *trans*). See table S24 for the number of structures and the reference structure of each group. Error bars indicate standard deviations.

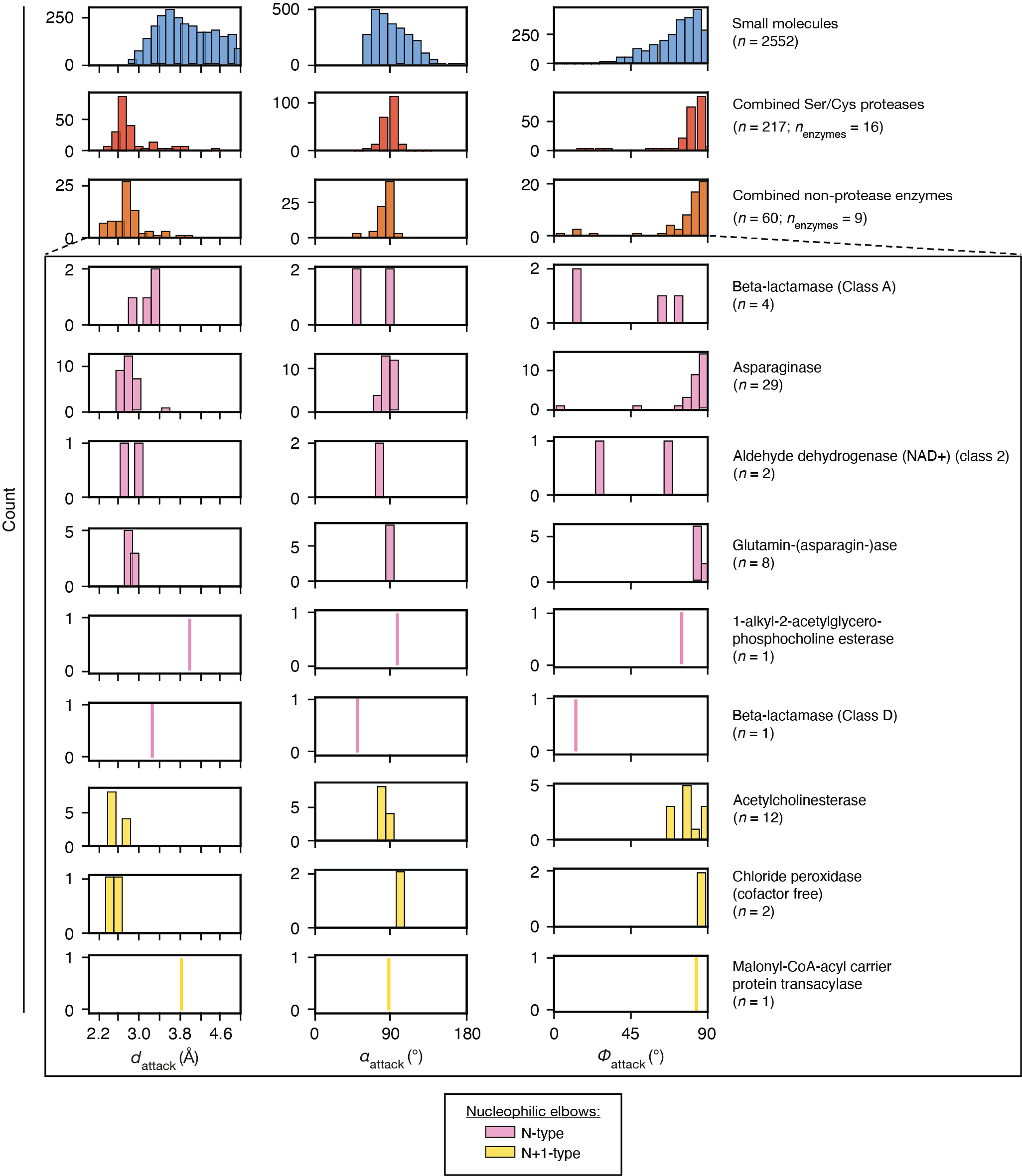

**Fig. S57.** Distributions of attack geometries (*d*_attack_, 𝛼_attack_ and 𝜙_attack_, defined in Fig. 2A) in small molecule hydroxyl•amide interactions (blue), in GSA-bound protease structures (“Combined Ser/Cys proteases”, red) and in additional nucleophilic elbow-bearing enzymes that perform nucleophilic attack on carbonyl compounds (“Combined non-protease enzymes”, orange). Enzymes in the “combined non-protease enzymes” group are plotted individually in the box and colored by their nucleophilic elbow types (N-type: pink; N+1-type: yellow).

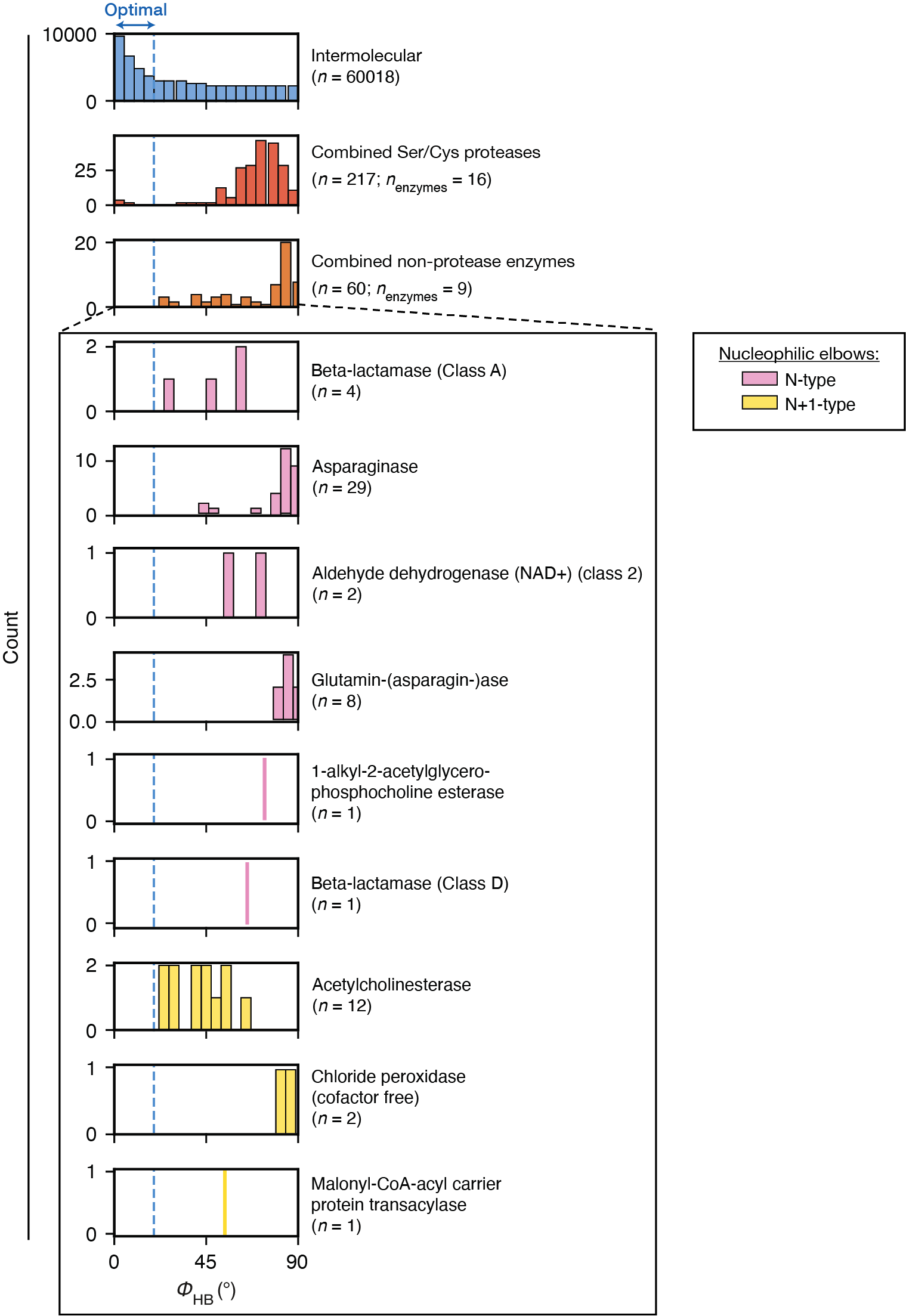

**Fig. S58.** Distributions of oxyanion hole hydrogen bond orientations (𝜙_attack_, defined in Fig. 5F) in small molecule intermolecular amide•carbonyl hydrogen bonds (blue), in GSA-bound protease structures (“Combined Ser/Cys proteases”, red) and in additional nucleophilic elbow-bearing enzymes that perform nucleophilic attack on carbonyl compounds (“Combined non-protease enzymes”, orange). Enzymes in the “combined non-protease enzymes” group are plotted individually in the box and colored by their nucleophilic elbow types (N-type: pink; N+1-type: yellow). The blue arrow and dashed lines indicate the region of 𝜙_HB_ ≤ 20° assigned as “optimal”.

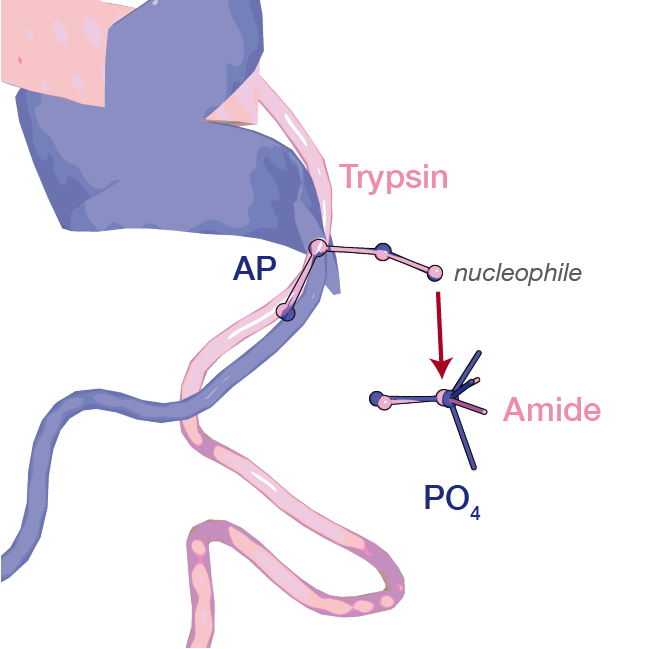

**Fig. S59.** Alkaline phosphatase (“AP”, PDB:1Y6V, purple) aligned with trypsin (PDB: 3M7Q, pink) on nucleophilic elbow atoms (shown in spheres) in their GSA-bound states.

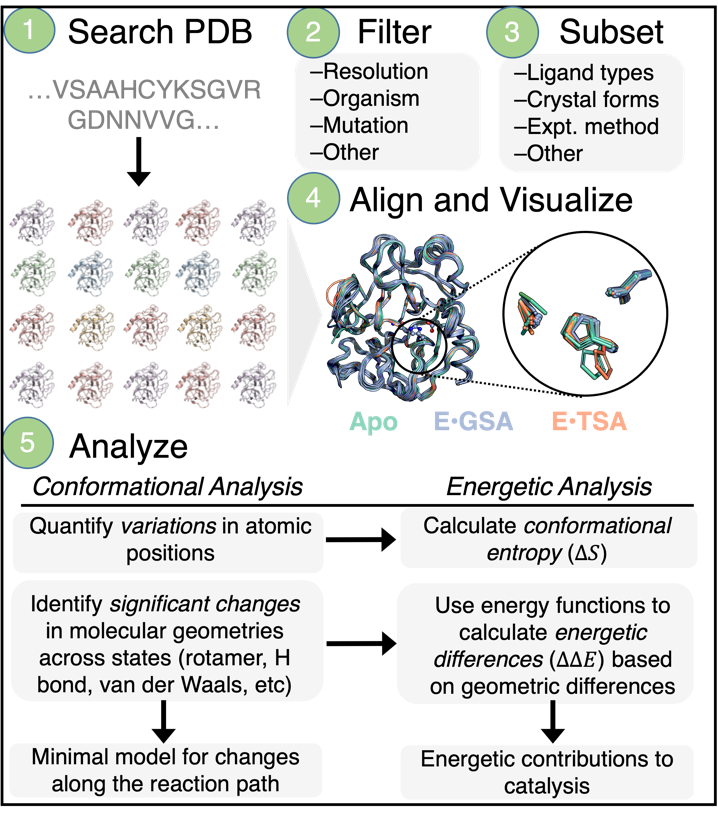

**Fig. S60.** The *Ensemble*PDB pipeline and functionalities.

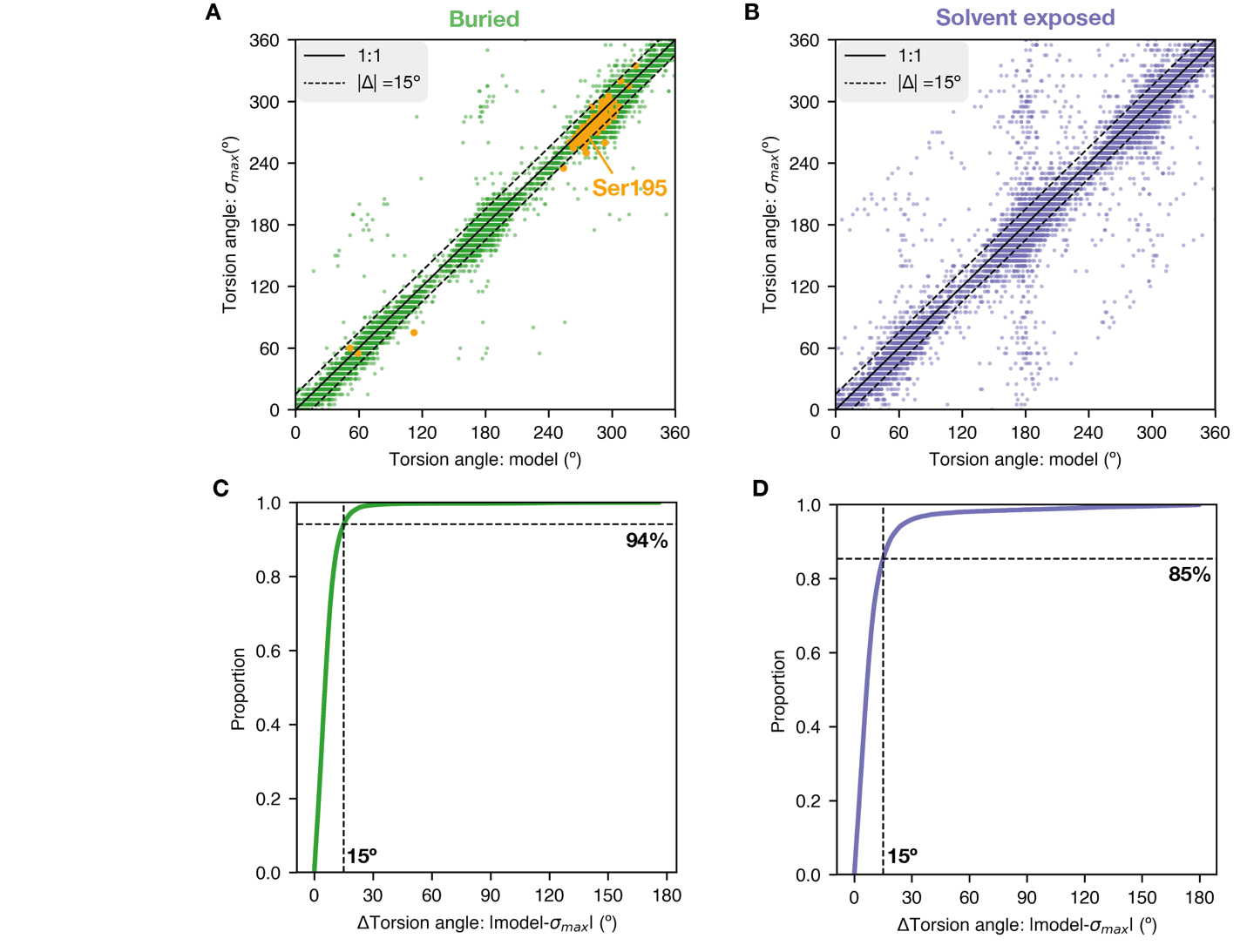

**Fig. S61.** The agreement between modeled sidechain torsion angles and the electron density distributions obtained from *Ringer* (*118*). **(A)** The sidechain torsion angles in the model *versus* the peak of local electron density distribution (σ_max_) for buried residues in trypsin, chymotrypsin and elastase pseudo-ensembles in the apo, GSA-bound and TSA-bound states (138 structures, 26,358 angles). Orange points indicate Ser195 χ^1^ and green points indicate all other torsion angles. The solid line indicates perfect agreement (1:1) and dashed lines indicate a difference within 15º between the modeled angle and the peak of the *Ringer* distribution (σ_max_). **(B)** The same analyses as (A) performed for solvent exposed residues (23,083 angles). **(C)** Cumulative probability density function for the difference in modeled torsion angles and the electron density peak (ΔTorsion angle: |model – σ_max_**|**) for buried residues; 94% of the torsion angles are accurately modeled, with <15º deviation from σ_max_. **(D)** Cumulative probability density function for ΔTorsion angle for solvent exposed residues. Solvent-exposed torsion angles are modeled less accurately than the buried ones, with 85% of the modeled angles showing a deviation from σ_max_ smaller than 15º.

**Table S1.** A subset of models for enzyme catalysis that invoke the positioning of enzyme groups.

| **Classic Catalytic Models Invoking Positioning** | |
| --- | --- |
| *Transition state complementarity* | Haldane (1930) (*117*); Pauling (1946) (*118*–*120*); Jencks (1975) (*121*) |
| *“Entatic states” in enzyme active sites* | Vallee & Williams (1968) (*122*) |
| *Orbital Steering* | Storm & Koshland (1970) (*123*) |
| *Binding entropy* | Page & Jencks (1971) (*124*) |
| *Freezing at reaction centers on enzyme* | Mildvan (1974) (*110*) |
| *Ground state destabilization* | Jencks (1975) (*106*) |
| *Electrostatic Preorganization* | Warshel & Levitt (1976) (*108*) |
| *Spatiotemporal effects* | Menger (1985) (*13*, *14*, *187*) |
| *Near-attack conformers* | Hur & Bruice (2002) (*188*) |

**Table S2.** Peptide hydrolysis rate constants in solution **(A)** and on serine proteases **(B)**.

| (A) | **Molecule** | **pH** | **T (°C)** | **Rate constant (s^-1^)** | | **Ref** |
| --- | --- | --- | --- | --- | --- | --- |
| *Solution* | Glycylglycine | 6.8 | 25 | 6.3 × 10^-11^ | | (*189*)† |
|  | N-acetyl glycyl glycine | 6.8 | 25 | 4.4 × 10^-11^ | |  |
|  | N-acetylglycylglycine  N′-methylamide¶ | 6.8 | 25 | 3.6 × 10^-11^ | |  |
|  | hippuryl phenylalanine | 9 | 25 | 1.3× 10^-10^ | | (*190*) |
|  | (phenylacetyl)glycyl bond of *N*-(phenylacetyl)-  glycyl-D-valine | 7 | 37 | 4.65 (0.44) × 10^-11^ | | (*24*)† |
|  | Glycyl-D-valyl bond  of *N*-(phenylacetyl)-  glycyl-D-valine | 7 | 37 | 2.69 (0.13) × 10^-11^ | |  |
| **(B)** | **Substrate**§ | **pH** | **T (°C)** | **Rate constant (s^-1^)** | | **Ref** |
|  |  |  |  | *Acylation* | *k_cat_* |  |
| *Trypsin* | Soybean inhibitor  (ground state analog)¶ | 8 | 21 | 140 | – | (*192*) |
|  | **YLVGHRGFFYDA** | 8 | 25 | – | 16 | (*193*) |
|  | **YLVGPRGFFYDA** | 8 | 25 | – | 31 |  |
|  | Insulin, beta-chain | 8 | 25 | – | 40 | (*194*) |
|  | **RPPPHFSPFRSVQ** | 9 | 25 | 33* | 33 | (*195*) |
|  | **FRSVQ** | 9 | 25 | 55* | 55 |  |
|  | Ac-**FRSV**-NH_2_ | 9 | 25 | 153* | 153 |  |
| *Chymotrypsin* | Ac-**PAPFAA**-NH_2_ | 8 | 30 | 59.5* | 59.5 | (*196*) |
|  | Ac-**PAPFAAA**-NH_2_ | 8 | 30 | 60.6* | 60.6 |  |
| *α-Lytic Protease* | Ac-**PAPAAA**-NH_2_ | 9 | 37 | – | 17.5 | (*90*) |
| *Elastase* | Ac-**PAPA**-NH_2_ | 9 | 37 | – | 8.5 | (*89*) |
| *Subtilisin* | Ac-**PAPFAA**-NH_2_ | 8 | 25 | – | 40 | (*104*) |
| *Proteinase K* | Ac-**PAPF**-NH_2_ | 8 | 25 | – | 64 |  |

† There is no significant buffer catalysis, as the increase in buffer concentration and in ionic strength did not affect the rate of hydrolysis.

§ Peptide sequences are represented by one-letter amino acid code (bolded). Substrates with the fastest rates are listed; see references for the rates of additional substrates.

*Deacylation rates were inferred by measuring the *k_cat_* of the same sequence with an ester leaving group and were shown to be faster or similar to *k_cat_*. Thus, the acylation step is expected to be rate-limiting.

¶ Values used in Fig. 5A for the experimentally measured catalysis. N-acetylglycylglycine

N′-methylamide mimics a regular peptide amide bond that is not at the N or C terminal. The soybean inhibitor is a ground state analog that follows the same reaction mechanism as all other ground state analogs in our pseudo-ensembles (supplementary text S2) and thus its acylation rate is the best comparison to the predictions from pseudo-ensemble analyses.

$\Delta\Delta G^{\ddagger}=-RTln\left( \frac{3.6\times{10}^{-11}}{140 s^{-1}} \right)= 0.59\frac{kcal}{mol}*29.0=17.1 kcal/mol$

**Table S3.** Serine proteases investigated in this study.

| **Clan**†  **(CATH**‡**)** | **Protease** | **Family**† | **Sub-**  **family**† | **Organism** | **Catalytic groups** | | **P1 specificity**† |
| --- | --- | --- | --- | --- | --- | --- | --- |
|  |  |  |  |  | *Catalytic Triad* | *Oxyanion Hole*§ |  |
| PA  (2.40.10.10) | Alpha-Lytic Protease | S1 | E | *Lysobacter enzymogenes* | S195 - H57 - D102 | S195:N + G193:N | T/A/S/V |
|  | Chymotrypsin | S1 | A | *Bos taurus* | S195 - H57 - D102 | S195:N + G193:N | F/Y/W/L |
|  | Elastase | S1 | A | *Sus scrofa* | S195 - H57 - D102 | S195:N + G193:N | F/L/Y |
|  | Streptogrisin B | S1 | E | *Streptomyces griseus* | S195 - H57 - D102 | S195:N + G193:N | F/L/A |
|  | Thrombin | S1 | A | *Homo sapiens* | S195 - H57 - D102 | S195:N + G193:N | R/K |
|  | Trypsin | S1 | A | *Bos taurus* | S195 - H57 - D102 | S195:N + G193:N | R/K |
| SB  (3.40.50.20) | Furin | S8 | B | *Homo sapiens* | S368 - H194 - D153 | S368:N + N295:Nδ2 | R |
|  | Kexin | S8 | B | *Saccharomyces cerevisiae* | S385 - H213 - D176 | S385:N + N314:Nδ2 | R |
|  | Proteinase K | S8 | A | *Parengyodontium album* | S224 - H69 - D39 | S224:N + N161:Nδ2 | Neutral amino acids |
|  | Sedolisin | S53 | / | *Pseudomonas sp. 101* | S287 - E80 - D84 | S287:N + D170:Oδ2 | F |
|  | Subtilisin | S8 | A | *Bacillus amyloliquefaciens* | S221 - H64 - D32 | S221:N + N155:Nδ2 | Neutral amino acids |
| SC  (3.40.50.1820) | Dipeptidyl peptidase IV | S9 | B | *Homo sapiens* | S630 - H740 - D708 | Y631:N + Y547:Oη | P |
|  | Prolyl aminopeptidase | S33 | / | *Serratia marcescens* | S113 - H296 - D268 | W114:N + G46:N | P |
|  | Prolyl endopeptidase | S9 | A | *Sus scrofa* | S554 - H680 - D641 | N555:N + Y473:Oη | P |
|  | Serine carboxypeptidase A | S10 | / | *Homo sapiens* | S150 - H429 - D372 | Y151:N + G57:N | F |
| SE  (3.40.710.10) | D-Ala-D-Ala peptidase | S12 | / | *Streptomyces sp. R61* | S62 - Y159 - K65 | S62:N + T301:N | A |

† Clan, family, subfamily and P1 specificity were obtained from the MEROPS (*105*) database. P1 specificity is indicated by amino acid one-letter code or amino acid properties if the specificity is broad.

‡ CATH ids were obtained from the CATH Protein Structure Classification database (*107*).

§ Atom names for the hydrogen bond donors are specified after the colon symbol and follow the PDB format.

**Table S4.** Number of serine protease structures in each class. All structures are wild type (WT), with 100% sequence identity with the reference PDB chain(s).

| **Clan** | **Protease** | **Reference**  **PDB Chain**† | **Number of WT Structures in each Subensemble**‡ | | | | | | |
| --- | --- | --- | --- | --- | --- | --- | --- | --- | --- |
|  |  |  | *APO* | *GSA* | *TSA* | *Type II*  *Inhibitors* | *Acyl-*  *enzyme* | *Other* | *Total* |
| PA | Alpha-Lytic Protease | 2H5C_A | 6 | – | 10 | – | – | 1 | 17 |
|  | Chymotrypsin | 1ACB_E | 12 | 21 | 13 | 2 | 4 | 5 | 57 |
|  | Elastase | 1QNJ_A | 42 | 7 | 9 | – | 11 | 33 | 102 |
|  | Streptogrisin B | 2QAA_A | – | 16 | 3 | – | – | – | 19 |
|  | Thrombin | 1C5L_L_H | 16 | 10 | 25 | 25 | 5 | 175 | 256 |
|  | Trypsin | 3MI4_A | 19 | 79 | 13 | – | 8 | 326 | 445 |
| SB | Furin | 5JXG_A | 2 | – | – | 2 | – | 23 | 27 |
|  | Kexin | 2ID4_A | – | – | 4 | 1 | – | – | 5 |
|  | Proteinase K | 1IC6_A | 111 | 1 | 1 | 2 | – | 25 | 140 |
|  | Sedolisin | 1GA4_A | 1 | – | 7 | – | – | 1 | 9 |
|  | Subtilisin | 1ST2_A | 2 | 22 | – | – | – | – | 24 |
| SC | Dipeptidyl peptidase IV | 1PFQ_A | 25 | 6 | 1 | – | 6 | 54 | 92 |
|  | Prolyl aminopeptidase | 1QTR_A | 1 | 3 | – | – | – | – | 4 |
|  | Prolyl endopeptidase | 1H2W_A | 1 | – | 6 | – | 2 | 2 | 11 |
|  | Serine carboxypeptidase A | 1IVY_A | 3 | – | 2 | – | – | 3 | 8 |
| SE | D-Ala-D-Ala peptidase | 3PTE_A | 1 | 2 | 3 | – | 6 | 2 | 14 |
| *total* | | | 243 | 167 | 97 | 32 | 42 | 650 | 1231 |

† PDB code followed by chain names in the original PDB.

‡ Subensembles are defined in fig. S5.
**Table S5.** Comparison of the ground state, transition state and tetrahedral intermediate state geometries from QM calculation of the solution reaction using *N*-methylacetamide (NMA) as electrophile and ethanol (EtOH) as nucleophile.

|  | **Ground state** | **Transition state** | **Tetrahedral intermediate** |
| --- | --- | --- | --- |
| *d*_attack_† (Å) | 3.62 | 1.97 | 1.60 |
| *a*_attack_† (°) | 82 | 107 | 111 |
| *ϕ*_attack_† (°) | 84 | 109 | 118 |

† Defined in fig. S7.

**Table S6.** Search criteria used to collect small molecule interactions from the CSD (*107*). Substructures are specified as SMARTS strings (*107*), and atoms are specified by the index (0-based) for terms the SMARTS string.

| **Interaction** | | **Substructure and atom index** | | | | | | | **Constraints^#^** | | |
| --- | --- | --- | --- | --- | --- | --- | --- | --- | --- | --- | --- |
|  |  | ***Group1*** | ***Atom***  ***1.1*** | ***H***  ***atom*** | ***Atom***  ***1.2*** | ***Group2*** | ***Atom***  ***2.1*** | ***Atom***  ***2.2*** | ***Distance***  ***A1.1–A2.1*** | ***Angle***  ***(A1.1–H–A2.1)*** | ***Distance***  ***A1.2–A2.2*** |
| ***vdW*** | Hydroxyl•carbonyl* | [CX4&+0]  [OX2&+0][H] | 1 | - | - | [CX3&+0]  (=[OX1&+0])  ([*])([*]) | 0 | - | ≤ 5Å | - | - |
|  | Hydroxyl•amide* |  |  |  |  | [NX3&+0]([H])  ([CX3]=[OX1]) | 2 |  |  | - | - |
| ***H-bond*** | Amide•carbonyl† | [NX3&+0]([H])  ([*+0])  ([CX3]=[OX1]) | 0 | 1 |  | [CX3&+0]  (=[OX1&+0])  ([*]) | 1 |  | ≤ 4Å | ≥ 120° | - |
|  | Amide•O(sp3)-C‡ |  |  |  |  | [CX4&+0]  (-[OH])([*]) | 1 |  |  |  | - |
|  | (neutral)  imidazole•hydroxyl**§** | N1(-[H])C=NC=C1 | 0 | 1 |  | [CX4&+0]  [OX2&+0][H] | 1 |  | ≤ 4Å | ≥ 120° | - |
|  | (protonated)  imidazole•ether/hydroxyl¶ | N1(-[H])C=[N&+1]  (-[H])C=C1 | 0, 3 | 1, 4 |  | [CX4&+0]  [OX2&+0][C,H] | 1 |  |  |  | - |
|  | (neutral)  imidazole•carboxylate ** | N1(-[H])C=[N&+0]  C=C1 | 0 | 1 | - | CC(-[O&-1])=O | 2 | - | ≤ 4Å | ≥ 120° | - |
|  | (protonated)  imidazole•carboxylate†† | N1(-[H])C=[N&+1]  (-[H])C=C1 | 0 | 1 | - |  |  |  |  |  | - |
| ***Nucl.***  ***Elbow*** | N-type intermolecular**§§** | N(-[H])  (-[C](~[*]))  CC(-[OH]) | 0 | 1 | 6 | [CX3&+0]]  (=[OX1&+0])  ([*])([*]) | 1 | 0 | ≤ 5Å | ≥ 120° | ≤ 5Å |
|  | N+1-type intermolecular**§§** | N(-[H])  (-[C](~[*]))  CCC(-[OH]) | 0 | 1 | 7 |  |  |  |  |  |  |
|  | N-type covalent¶¶ | N([H])  (CC[N,O,C]C=O) | 0 | 1 | 4 | - | 6 | 5 |  |  |  |
|  | N+1-type covalent¶¶ | N([H])  (CCC[N,O,C]C=O) | 0 | 1 | 6 |  | 7 | 8 |  |  |  |

**#** Constraints used in searching criteria; interactions not matching these criteria were filtered.

* Small molecule analog for the reacting nucleophile•electrophile in the enzymatic reaction.

**†** Small molecule analog for the oxyanion hole hydrogen bond in the enzymatic reaction ground state.

‡ Small molecule analog for the oxyanion hole hydrogen bond in the enzymatic reaction transition state.

§ Small molecule analog for the catalytic triad His•Ser hydrogen bond in the enzymatic reaction ground state.

¶ Small molecule analog for the catalytic triad His•Ser hydrogen bond in the enzymatic reaction transition state.

** Small molecule analog for the catalytic triad His•Asp hydrogen bond in the enzymatic reaction ground state.

†† Small molecule analog for the catalytic triad His•Asp hydrogen bond in the enzymatic reaction transition state.

§§ Small molecule analog for the nucleophilic elbow structure in the enzymatic ground state.

¶¶ Small molecule analog for the nucleophilic elbow structure in the enzymatic acylenzyme state. Because the covalent bond forms between the nucleophile and the electrophile in this reaction state, the hydrogen bond becomes intramolecular and both the donor and acceptor atoms are contained in group1. All atom indices indicate their positions in group1.

**Table S7.** Geometric parameters defining the reaction geometry for serine proteases and solution.

|  |  | **Protease** | ***n*** | ***d*_attack_**† **(Å)** | | ***a*_attack_**† **(°)** | | ***ϕ*_attack_**† **(°)** | |
| --- | --- | --- | --- | --- | --- | --- | --- | --- | --- |
|  |  |  |  | *mode*‡ | *s.d* | *mode* | *s.d* | *mode* | *s.d* |
| *Enzyme* | PA | Chymotrypsin | 21 | 2.71 | 0.12 | 93 | 7 | 85 | 4 |
|  |  | Elastase | 7 | 3.07 | 0.26 | 88 | 13 | 85 | 27 |
|  |  | Streptogrisin B | 16 | 2.77 | 0.05 | 92 | 3 | 83 | 2 |
|  |  | Thrombin | 10 | 2.78 | 0.26 | 89 | 18 | 82 | 18 |
|  |  | Trypsin | 79 | 2.65 | 0.11 | 92 | 6 | 85 | 3 |
|  |  | *combined* | 133 | 2.70 | 0.15 | 93 | 8 | 85 | 9 |
|  | SB | Proteinase K | 1 | 2.91 | – | 78 | – | 57 | – |
|  |  | Subtilisin | 22 | 2.65 | 0.09 | 94 | 3 | 83 | 2 |
|  |  | *combined* | 23 | 2.65 | 0.10 | 94 | 4 | 83 | 6 |
|  | SC | Dipeptidyl peptidase IV | 6 | 2.55 | 0.08 | 83 | 3 | 84 | 2 |
|  |  | Prolyl aminopeptidase | 3 | 2.78 | 0.06 | 84 | 3 | 84 | 13 |
|  |  | *combined* | 9 | 2.58 | 0.12 | 84 | 3 | 85 | 7 |
|  | SE | D-Ala-D-Ala peptidase | 2 | 2.84 | 0.10 | 89 | 5 | 88 | 2 |
|  | *combined* | | 167 | 2.68 | 0.14 | 93 | 7 | 84 | 8 |
| *Solution* | MD (HOH-NMA) rep#1 | | 5000 | 3.43 | 0.18 | 109 | 30 | 83 | 21 |
|  | MD (HOH-NMA) rep#2 | | 5000 | 3.44 | 0.18 | 109 | 29 | 82 | 22 |
|  | MD (HOH-NMA) rep#3 | | 5000 | 3.45 | 0.18 | 111 | 31 | 82 | 22 |
|  | CSD (OH•amide) | | 2552 | 3.61 | 0.54 | 72 | 21 | 84 | 15 |
|  | CSD (OH•carbonyl) | | 13690 | 3.63 | 0.55 | 83 | 19 | 85 | 16 |
|  | QM (ground state) | | 1 | 3.62 | – | 82 | – | 84 | – |

†	Defined in Fig. 2A.

‡ Mode is reported instead of mean, as it is the highest probability value, corresponding to the most energetically favored geometry.

**­_­­­_Table S8.** Catalytic serine χ^1^ angles in apo, GSA-bound and TSA-bound states and their changes across states for all proteases (P) and additional non-serine proteases (N).

| **Clan** | **Type**† | **Enzyme** | ***Apo***  **Ser χ^1^** | | | ***E●GSA***  **Ser χ^1^** | | | ***E●TSA* Ser χ^1^** | | | ***E●GSA – Apo***  **ΔSer χ^1^** | | ***E●TSA – E●GSA***  **ΔSer χ^1^** | |
| --- | --- | --- | --- | --- | --- | --- | --- | --- | --- | --- | --- | --- | --- | --- | --- |
|  |  |  | *n* | *mode*‡  (°) | *s.d.*  (°) | *n* | *mode*‡  (°) | *s.d.*  (°) | *n* | *mode*‡  (°) | *s.d.*  (°) | Δ¶  (°) | *p*-value§ | Δ¶  (°) | *p*-value§ |
| PA | P | Alpha-Lytic  Protease | 6 | 287 | 2 | – | – | – | 10 | 304 | 5 | – | – | – | – |
|  |  | Chymotrypsin | 12 | 283 | 13 | 21 | 274 | 47 | 13 | 286 | 12 | -9 | 9E-05 | 12 | 2E-06 |
|  |  | Elastase | 42 | 291 | 7 | 7 | 272 | 75 | 9 | 283 | 4 | -19 | 4E-05 | 11 | 3E-03 |
|  |  | Streptogrisin B | – | – | – | 16 | 271 | 10 | 3 | 297 | 1 | – | – | 26 | – |
|  |  | Thrombin | 16 | 281 | 69 | 10 | 273 | 7 | 25 | 287 | 10 | -8 | 3E-01 | 14 | 3E-03 |
|  |  | Trypsin | 19 | 287 | 6 | 79 | 273 | 5 | 13 | 287 | 12 | -14 | 5E-19 | 14 | 3E-08 |
|  |  | *Combined* | 95 | 290 | 33 | 133 | 272 | 27 | 73 | 286 | 11 | -17 | 4E-39 | 13 | 2E-32 |
| SB | P | Furin | 2 | 72 | 1 | – | – | – | – | – | – | – | – | – | – |
|  |  | Kexin | – | – | – | – | – | – | 4 | 252 | 119 | – | – | – | – |
|  |  | Proteinase K | 111 | 285 | 36 | 1 | – | – | 1 | – | – | – | – | – | – |
|  |  | Sedolisin | 1 | – | – | – | – | – | 7 | 294 | 4 | – | – | – | – |
|  |  | Subtilisin | 2 | 301 | 6 | 22 | 277 | 7 | – | – | – | -23 | – | – | – |
|  |  | *combined* | 116 | 285 | 44 | 23 | 278 | 7 | 12 | 289 | 92 | -7 | 7E-11 | 12 | 1E-05 |
| SC | N | 2-hydroxy-muconate-  semialdehyde hydrolase | 1 | – | – | – | – | – | – | – | – | – | – | – | – |
|  |  | 6-deoxyerythronolide-B  synthase | 6 | 194 | 57 | – | – | – | 1 | – | – | – | – | – | – |
|  |  | acetylcholinesterase | 78 | 199 | 17 | 12 | 211 | 7 | 58 | 199 | 11 | 12 | 1E-06 | -12 | 2E-06 |
|  |  | alpha-amino-acid esterase | – | – | – | – | – | – | – | – | – | – | – | – | – |
|  |  | Carboxylesterase | – | – | – | – | – | – | 2 | 201 | 11 | – | – | – | – |
|  |  | cephalosporin-C  deacetylase | 2 | 203 | 9 | – | – | – | – | – | – | – | – | – | – |
|  |  | chloride peroxidase  (cofactor free) | 62 | 181 | 8 | 2 | 200 | 3 | – | – | – | 19 | – | – | – |
|  |  | cutinase | 35 | 197 | 17 | – | – | – | 13 | 195 | 10 | – | – | – | – |
|  |  | esterase | 3 | 200 | 7 | – | – | – | – | – | – | – | – | – | – |
|  |  | myristoyl-ACP-specific  thioesterase | 2 | 192 | 1 | – | – | – | – | – | – | – | – | – | – |
|  |  | Palmitoyl [protein]  hydrolase (type 2) | 1 | – | – | – | – | – | – | – | – | – | – | – | – |
|  |  | para-nitrobenzyl  esterase | 2 | 202 | 3 | – | – | – | – | – | – | – | – | – | – |
|  |  | triacylglycerol lipase  (EstA) | 28 | 200 | 48 | – | – | – | 4 | 209 | 1 | – | – | – | – |
|  |  | triacylglycerol lipase  (Pseudomonas family) | 8 | 216 | 6 | – | – | – | – | – | – | – | – | – | – |
|  |  | triacylglycerol lipase  (pancreatic) | 1 | – | – | – | – | – | – | – | – | – | – | – | – |
|  |  | Triacylglycerol lipase  (type B  carboxylestrase) | 1 | – | – | – | – | – | – | – | – | – | – | – | – |
|  |  | *combined* | 230 | 196 | 25 | 14 | 211 | 7 | 78 | 198 | 11 | 14 | 2E-05 | -13 | 4E-04 |
|  | P | Dipeptidyl peptidase IV | 25 | 197 | 23 | 6 | 201 | 8 | 1 | – | – | 4 | 1E+00 | – | – |
|  |  | Prolyl aminopeptidase | 1 | – | – | 3 | 218 | 2 | – | – | – | – | – | – | – |
|  |  | Prolyl endopeptidase | 1 | – | – | – | – | – | 6 | 200 | 2 | – | – | – | – |
|  |  | Serine carboxypeptidase A | 3 | 196 | 2 | – | – | – | 2 | 202 | 6 | – | – | – | – |
|  |  | Carboxypeptidase D | – | – | – | 1 | – | – | 2 | 188 | 16 | – | – | – | – |
|  |  | *combined* | 260 | 196 | 25 | 24 | 209 | 9 | 89 | 198 | 11 | 13 | 1E-05 | -11 | 3E-04 |
|  | *combined* | | 30 | 198 | 21 | 10 | 205 | 11 | 11 | 200 | 11 | 7 | 2E-01 | -6 | 9E-02 |
| SE | N | beta-lactamase  (Class A) | 96 | 287 | 51 | 5 | 297 | 15 | 15 | 287 | 5 | 9 | – | -10 | – |
|  |  | beta-lactamase  (Class C) | 3 | 293 | 7 | – | – | – | 3 | 294 | 11 | – | – | – | – |
|  |  | beta-lactamase  (Class D) | – | – | – | 1 | – | – | – | – | – | – | – | – | – |
|  |  | *combined* | 99 | 287 | 50 | 6 | 296 | 13 | 18 | 288 | 8 | 8 | 3E-01 | -7 | 7E-01 |
|  | P | D-Ala-D-Ala peptidase | 1 | – | – | 2 | 279 | 0 | 3 | 294 | 2 | – | – | 15 | – |
|  | *combined* | | 100 | 288 | 51 | 8 | 281 | 13 | 21 | 291 | 7 | -7 | 2E-01 | 10 | 7E-02 |

† “P” for proteases and “N” for non-protease enzymes.

‡ The highest probability value, corresponding to the most energetically favored geometry.

§ *P*-values were calculated via the two-sided Kolmogorov–Smirnov test when both the GSA-bound and TSA-bound ensembles contain more than 5 structures.

¶ The difference between the most probable values (modes) of the two states.

**Table S9.** Bond lengths (*d*_O–X_) between the nucleophile and the attacked electrophile in the TSA-bound enzymes or in the tetrahedral intermediate state in the solution reaction.

|  | Electrophilic  atom | ***d*_O–X_ (Å)** | |
| --- | --- | --- | --- |
|  |  | *mean*‡ | *s.d.* |
| Enzyme† | B | 1.56 | 0.11 |
|  | P | 1.58 | 0.06 |
|  | S | 1.52 | 0.12 |
|  | C | 1.56 | 0.19 |
|  | *combined* | 1.56 | 0.16 |
| Solution (QM) | C | 1.60 | – |

† From TSA-bound structures combining all serine proteases. These TSA structures are subsetted by the identity of their attacked electrophilic atom (B: boron, P: phosphorus, S:sulfur, C: carbon).

**Table S10.** Distance changes between the reacting atoms on the reaction path (Δ*d*_O•C_) resulting from each of three types of motions (defined in Fig. 3D).

|  | **Clan** | **Enzyme** | **Δ*d*_O•C_ (Å)** | | | | | | **Total Δ*d*_O•C_**  **required**† **(Å)** | |
| --- | --- | --- | --- | --- | --- | --- | --- | --- | --- | --- |
|  |  |  | **χ^1^ rotation** | | **Translation** | | **sp^2^🡪 sp^3^** | |  |  |
|  |  |  | *median* | *s.d.* | *median* | *s.d.* | *median* | *s.d.* | *median* | *s.d.* |
| *Enzyme* | PA | Trypsin | 0.32 | 0.22 | 0.36 | 0.22 | 0.42 | 0.09 | 1.10 | 0.15 |
|  |  | Chymotrypsin | 0.24 | 0.27 | 0.36 | 0.20 | 0.50 | 0.15 | 1.25 | 0.21 |
|  |  | Elastase | 0.24 | 0.22 | 0.41 | 0.40 | 0.48 | 0.13 | 1.55 | 0.25 |
|  |  | Alpha-Lytic Protease | – | – | 0.28 | 0.19 | 0.40 | 0.09 | 1.25 | 0.10 |
|  |  | Streptogrisin B | 0.49 | 0.19 | 0.32 | 0.12 | 0.26 | 0.12 | 1.08 | 0.17 |
|  |  | Thrombin | 0.20 | 0.23 | -0.38 | 0.89 | 0.39 | 0.14 | 1.12 | 0.32 |
|  | SB | Proteinase K | 0.23 | – | 2.03 | – | 0.50 | – | 1.43 | – |
|  | SE | D-Ala-D-Ala peptidase | 0.32 | 0.04 | 0.19 | 0.18 | 0.41 | 0.04 | 1.21 | 0.08 |
|  | SC | Dipeptidyl peptidase IV | 0.27 | 0.12 | 0.23 | 0.07 | 0.02 | – | 1.11 | 0.08 |
|  |  | Prolyl endopeptidase | – | – | 0.22 | 0.13 | 0.47 | 0.03 | 1.29 | 0.07 |
|  |  | Acetylcholinesterase | 0.18 | 0.11 | – | – | – | – | – | – |
| *Solution (QM)*‡ | | | -0.18 | – | 1.58 | – | 0.60 | – | 1.93 | – |

†Determined by subtracting the covalent bond lengths in the TSA-bound structures from the starting *d*_O•C_ in the GSA-bound structures.

‡In QM the nucleophile is ethanol, and the electrophilic group is *N*-methylacetamide.

**Table S11.** Rate enhancements provided by general base catalysis calculated using a range of Brønsted coefficients (β) and effective molarities (EM).

| EM (M) | **Rate enhancement** | | | | | |
| --- | --- | --- | --- | --- | --- | --- |
|  | β = 0.50 | | β = 0.75 | | β = 1.0 | |
|  | fold | kcal/mol† | fold | kcal/mol | fold | kcal/mol |
| 1 | 5.8$\times$10^2^ | 3.8 | 1.0$\times$10^5^ | 6.8 | 1.8$\times$10^7^ | 9.8 |
| 10‡ | 5.8$\times$10^3^ | 5.1 | 1.0$\times$10^6^ | 8.2 | 1.8$\times$10^8^ | 11.2 |
| 100 | 5.8$\times$10^4^ | 6.5 | 1.0$\times$10^7^ | 9.5 | 1.8$\times$10^9^ | 12.6 |

† Determined for *T* = 298 K.

‡ Value used to calculate the catalytic contribution of general base catalysis.

**Table S12.** Conformational entropies (*T*•*S*_conf_, with T = 298 K) calculated from the ensemble distributions of reactants in trypsin, from small molecule hydroxyl•amide interactions, and from MD simulations of NMA in water (5000 snapshots from 100 ns simulation). *T*•*S*_conf_ values were calculated **(A)** varying the bin width, and varying the sample size using two fixed bin widths at **(B)** 1 Å and **(C)** 0.1 Å.

| **(A)** | | ***T•S*_conf_**  (kcal/mol) | | | | | | **Δ*T*•*S*_conf_**  (kcal/mol) | |
| --- | --- | --- | --- | --- | --- | --- | --- | --- | --- |
| **Number of bins**† | **Bin width**† **(Å)** | **(1) Trypsin**  **(Ser–OH•amide)** | | **(2) Small molecule**  **(hydroxyl•amide)** | | **(3) MD**  **(water•amide)** | | **(2) – (1)** | **(3) – (1)** |
|  |  | *mean*‡ | *s.d.*‡ | *mean*‡ | *s.d.*‡ | *mean*‡ | *s.d.*‡ | *mean* | *mean* |
| 5 | 2 | 0.0 | 0.0 | 2.0 | 0.0 | 2.1 | 0.0 | 2.0 | 2.1 |
| 10 | 1 | 0.6 | 0.0 | 2.8 | 0.0 | 2.8 | 0.0 | 2.2 | 2.2 |
| 25 | 0.4 | 1.5 | 0.0 | 4.0 | 0.0 | 3.9 | 0.0 | 2.5 | 2.4 |
| 50 | 0.2 | 1.9 | 0.1 | 4.5 | 0.0 | 4.7 | 0.0 | 2.5 | 2.7 |
| 100 | 0.1 | 2.4 | 0.0 | 4.6 | 0.0 | 5.0 | 0.0 | 2.2 | 2.6 |
| 200 | 0.05 | 2.6 | 0.0 | 4.6 | 0.0 | 5.0 | 0.0 | 2.0 | 2.4 |
|  | | | | | | | ***mean***¶ | **2.2** | 2.4 |
|  |  |  |  |  |  |  | ***s.d.*** ¶ | **0.2** | 0.2 |

| **(B)** | ***T•S*_conf_**  (kcal/mol) | | | | | | **Δ*T•S*_conf_**  (kcal/mol) | |
| --- | --- | --- | --- | --- | --- | --- | --- | --- |
| **Number of samples**  **(resampled)** | **(1) Trypsin**  **(Ser–OH•amide)** | | **(2) Small molecule**  **(hydroxyl•amide)** | | **(3) MD**  **(water•amide)** | | **(2) – (1)** | **(3) – (1)** |
|  | *mean*‡ | *s.d.*‡ | *mean*‡ | *s.d.*‡ | *mean*‡ | *s.d.*‡ | *mean* | *mean* |
| 10 | 0.5 | 0.1 | 1.3 | 0.1 | 1.3 | 0.1 | 0.8 | 0.8 |
| 50 | 0.6 | 0.1 | 2.1 | 0.1 | 2.1 | 0.1 | 1.5 | 1.5 |
| 100 | 0.6 | 0.0 | 2.4 | 0.0 | 2.4 | 0.0 | 1.8 | 1.8 |
| 500 | 0.6 | 0.0 | 2.7 | 0.0 | 2.7 | 0.0 | 2.1 | 2.1 |
| 1000 | 0.6 | 0.0 | 2.8 | 0.0 | 2.7 | 0.0 | 2.2 | 2.1 |
| 10000 | 0.6 | 0.0 | 2.8 | 0.0 | 2.8 | 0.0 | 2.2 | 2.2 |
| 50000 | 0.6 | 0.0 | 2.8 | 0.0 | 2.8 | 0.0 | 2.2 | 2.2 |

| **(C)** | ***T•S*_conf_**  (kcal/mol) | | | | | | **Δ*T•S*_conf_**  (kcal/mol) | |
| --- | --- | --- | --- | --- | --- | --- | --- | --- |
| **Number of samples**  **(resampled)** | **(1) Trypsin**  **(Ser–OH•amide)** | | **(2) Small molecule**  **(hydroxyl•amide)** | | **(3) MD**  **(water•amide)** | | **(2) – (1)** | **(3) – (1)** |
|  | *mean*‡ | *s.d.*‡ | *mean*‡ | *s.d.*‡ | *mean*‡ | *s.d.*‡ | *mean* | *mean* |
| 10 | 1.3 | 0.1 | 1.4 | 0.0 | 1.4 | 0.0 | 0.1 | 0.1 |
| 50 | 2.0 | 0.1 | 2.3 | 0.0 | 2.3 | 0.0 | 0.3 | 0.3 |
| 100 | 2.2 | 0.0 | 2.7 | 0.0 | 2.7 | 0.0 | 0.5 | 0.5 |
| 500 | 2.4 | 0.0 | 3.6 | 0.0 | 3.6 | 0.0 | 1.2 | 1.3 |
| 1000 | 2.4 | 0.0 | 3.9 | 0.0 | 4.0 | 0.0 | 1.5 | 1.6 |
| 10000 | 2.4 | 0.0 | 4.5 | 0.0 | 4.8 | 0.0 | 2.1 | 2.4 |
| 50000 | 2.4 | 0.0 | 4.6 | 0.0 | 4.9 | 0.0 | 2.1 | 2.5 |

†Number of bins and bin width are specified for dimension and are the same for the three dimensions, e.g. a 1 Å bin width corresponds to a bin volume of (1Å)^3^.

‡ Calculated via bootstrap analyses (200 repeats).

¶ Calculated from the range of values above from 0.05 to 2 Å bin widths.

**Table S13**. Energy differences (ΔΔ*E*_enz–soln_) resulting from the differences in nucleophile•electrophile distances (*d*_C•O_) between GSA-bound serine proteases and the QM-determined solution ground state, calculated using the knowledge-based energy function (fig. S25).

| **Clan** | **Protease** | ***n*** | **ΔΔ*E*_enz–soln_**  **(kcal/mol)** | |
| --- | --- | --- | --- | --- |
|  |  |  | *mean* | *s.d.* |
| PA | Trypsin | 79 | 2.0 | 0.2 |
|  | Thrombin | 10 | 1.7 | 0.6 |
|  | Elastase | 7 | 1.2 | 0.8 |
|  | Chymotrypsin | 21 | 1.9 | 0.3 |
|  | Streptogrisin B | 16 | 2.0 | 0.1 |
| SB | Subtilisin | 22 | 1.9 | 0.2 |
|  | Proteinase K | 1 | 1.0 | – |
| SC | Dipeptidyl peptidase IV | 6 | 2.0 | 0.0 |
|  | Prolyl aminopeptidase | 3 | 2.0 | 0.0 |
| SE | D-Ala-D-Ala peptidase | 2 | 1.6 | 0.4 |

**Table S14.** Energy differences resulting from the catalytic serine χ^1^ rotation from the apo to the GSA-bound state (ΔΔ*E*_GS–apo_) and from the GSA-bound to the TSA-bound state (ΔΔ*E*^‡^), calculated using the knowledge-based energy function (fig. S26).

| **Clan** | **Protease** | **Apo**  *n* | **E•GSA**  *n* | **E•TSA**  *n* | **ΔΔ*E*_GS–apo_**  **(kcal/mol)** | | **ΔΔ*E*^‡^**  **(kcal/mol)** | |
| --- | --- | --- | --- | --- | --- | --- | --- | --- |
|  |  |  |  |  | *mean* | *s.d.* | *mean* | *s.d* |
| PA | Trypsin | 19 | 79 | 13 | 2.12 | 0.76 | -1.52 | 1.14 |
|  | Thrombin | 16 | 10 | 25 | 0.14 | 1.71 | -1.05 | 1.33 |
|  | Elastase | 42 | 7 | 9 | 1.70 | 1.47 | -1.30 | 1.57 |
|  | Chymotrypsin | 12 | 21 | 13 | 0.71 | 1.03 | -1.16 | 1.12 |
|  | Streptogrisin B | – | 16 | 3 | – | – | -2.29 | 0.98 |
| SB | Subtilisin | 2 | 22 | 0 | 1.70 | 0.94 | – | – |
|  | Proteinase K | 111 | 1 | 1 | 0.31 | 0.68 | -1.25 | – |
| SC | Dipeptidyl peptidase IV | 25 | 6 | 1 | -0.07 | 1.27 | -1.32 | 0.75 |
|  | Prolyl aminopeptidase | 1 | 3 | 0 | 0.96 | – | – | – |
| SE | D-Ala-D-Ala peptidase | 1 | 2 | 3 | -2.67 | – | -1.55 | 0.29 |

**Table S15.** Oxyanion hole hydrogen bond geometries **(A)** in the GSA-bound pseudo-ensembles (“Enzyme”) and in small molecule amide•carbonyl hydrogen bonds (“Reference”) and **(B)** in the TSA-bound pseudo-ensembles (“Enzyme”) and in small molecule hydrogen bonds between amide groups and sp^3^-hybridized oxygens [amide•O(sp^3^)–C] (“Reference”). Geometric parameters are defined in fig. S30.

| **(A)** | **Clan** | **Protease** | ***n*** | **1° oxyanion hole**  **hydrogen bond** | | | | | | **2° oxyanion hole**  **hydrogen bond** | | | | | |
| --- | --- | --- | --- | --- | --- | --- | --- | --- | --- | --- | --- | --- | --- | --- | --- |
|  |  |  |  | *d*_HB_ (Å) | | *α*_HB_ (°) | | *ϕ*_HB_ (°) | | *d*_HB_ (Å) | | *α*_HB_ (°) | | *ϕ*_HB_ (°) | |
|  |  |  |  | *mode* | *s.d.* | *mode* | *s.d.* | *mode* | *s.d.* | *mode* | *s.d.* | *mode* | *s.d.* | *mode* | *s.d.* |
| Enzyme | PA | Chymotrypsin | 21 | 3.0 | 0.1 | 125 | 4 | 81 | 6 | 2.7 | 0.1 | 134 | 4 | 77 | 3 |
|  |  | Elastase | 7 | 3.1 | 0.4 | 125 | 10 | 65 | 12 | 2.7 | 0.1 | 135 | 12 | 75 | 20 |
|  |  | Streptogrisin B | 16 | 3.0 | 0.2 | 124 | 3 | 67 | 4 | 2.6 | 0.1 | 131 | 7 | 83 | 4 |
|  |  | Thrombin | 10 | 3.1 | 0.3 | 129 | 15 | 80 | 11 | 3.0 | 0.2 | 139 | 11 | 79 | 29 |
|  |  | Trypsin | 79 | 2.9 | 0.1 | 129 | 6 | 76 | 10 | 2.7 | 0.1 | 127 | 5 | 78 | 5 |
|  | SB | Proteinase K | 1 | 2.9 | – | 139 | – | 34 | – | 3.3 | – | 67 | – | 56 | – |
|  |  | Subtilisin | 22 | 3.0 | 0.1 | 124 | 4 | 75 | 8 | 2.8 | 0.1 | 101 | 5 | 87 | 3 |
|  | SC | Dipeptidyl peptidase IV | 6 | 3.0 | 0.1 | 131 | 2 | 68 | 2 | 2.7 | 0.1 | 117 | 2 | 74 | 3 |
|  |  | Prolyl aminopeptidase | 3 | 2.9 | 0.1 | 141 | 4 | 71 | 3 | 3.0 | 0.2 | 101 | 4 | 56 | 8 |
|  | SE | D-Ala-D-Ala peptidase | 2 | 2.8 | 0.1 | 130 | 7 | 80 | 7 | 2.8 | 0.0 | 128 | 4 | 73 | 2 |
|  | *combined* | | 167 | 2.9 | 0.2 | 125 | 7 | 76 | 10 | 2.7 | 0.1 | 128 | 14 | 78 | 11 |
| Reference | amide•carbonyl (CSD) | | 60018 | 2.9 | 0.2 | 123 | 20 | 5 | 27 |  |  |  |  |  |  |

| **(B)** | **Clan** | **Protease** | ***n*** | **1° oxyanion hole**  **hydrogen bond** | | | | | | **2° oxyanion hole**  **hydrogen bond** | | | | | |
| --- | --- | --- | --- | --- | --- | --- | --- | --- | --- | --- | --- | --- | --- | --- | --- |
|  |  |  |  | *d*_HB_ (Å) | | *α*_HB_ (°) | | *ϕ*_HB_ (°) | | *d*_HB_ (Å) | | *α*_HB_ (°) | | *ϕ*_HB_ (°) | *d*_HB_ (Å) |
|  |  |  |  | *mode* | *s.d.* | *mode* | *s.d.* | *mode* | *s.d.* | *mode* | *s.d.* | *mode* | *s.d.* | *mode* | *s.d.* |
| Enzyme | PA | Alpha-Lytic Protease | 10 | 2.9 | 0.1 | 101 | 2 | 6 | 6 | 2.6 | 0.1 | 148 | 4 | 28 | 13 |
|  |  | Chymotrypsin | 13 | 2.9 | 0.2 | 102 | 11 | 12 | 10 | 2.9 | 0.2 | 164 | 15 | 18 | 15 |
|  |  | Elastase | 9 | 2.7 | 0.1 | 110 | 7 | 11 | 4 | 2.6 | 0.1 | 145 | 7 | 24 | 11 |
|  |  | Streptogrisin B | 3 | 3.2 | 0.2 | 97 | 13 | 23 | 6 | 2.8 | 0.1 | 140 | 1 | 54 | 22 |
|  |  | Thrombin | 25 | 2.8 | 0.2 | 106 | 12 | 7 | 10 | 3.0 | 0.2 | 150 | 15 | 29 | 11 |
|  |  | Trypsin | 13 | 3.0 | 0.3 | 100 | 12 | 10 | 17 | 2.7 | 0.2 | 143 | 13 | 34 | 17 |
|  | SB | Kexin | 4 | 3.1 | 0.6 | 98 | 20 | 8 | 18 | 2.7 | 0.2 | 119 | 6 | 43 | 17 |
|  |  | Proteinase K | 1 | 2.9 | – | 101 | – | 7 | – | 2.6 | – | 124 | – | 33 | – |
|  |  | Sedolisin | 7 | 2.8 | 0.1 | 106 | 5 | 12 | 8 | 3.2 | 0.1 | 131 | 3 | 61 | 6 |
|  | SC | Dipeptidyl peptidase IV | 1 | 3.1 | – | 111 | – | 30 | – | 2.7 | – | 141 | – | 9 | – |
|  |  | Prolyl endopeptidase | 6 | 2.8 | 0.1 | 108 | 4 | 12 | 6 | 2.6 | 0.1 | 128 | 5 | 42 | 10 |
|  |  | Serine carboxypeptidase A | 2 | 2.8 | 0.3 | 107 | 3 | 16 | 5 | 2.6 | 0.0 | 139 | 9 | 47 | 5 |
|  | SE | D-Ala-D-Ala peptidase | 3 | 2.7 | 0.0 | 104 | 2 | 9 | 6 | 2.7 | 0.0 | 142 | 10 | 46 | 16 |
|  | *combined* | | 97 | 2.9 | 0.3 | 104 | 11 | 10 | 11 | 2.7 | 0.2 | 144 | 14 | 27 | 17 |
| Reference | | amide•O(sp^3^)-C (CSD) | 3876 | 3.1 | 0.3 | 119 | 17 | 35 | 26 |  |  |  |  |  |  |

**Table S16**. Energies of the oxyanion hole hydrogen bond geometries in the serine protease GSA- and TSA-bound states and their differences, calculated by mapping geometric measurements from pseudo-ensembles on knowledge-based energy functions (shown in fig. S33 to 35). The geometric parameters, **(A)** *d*_HB_, **(B)** *α*_HB_ and **(C)** *ϕ*_HB_, defined in fig. S30, and their values are listed in table S15.

| **(A)** | ***d*_HB_** | | | | | | | | | | | | | | |
| --- | --- | --- | --- | --- | --- | --- | --- | --- | --- | --- | --- | --- | --- | --- | --- |
| **Clan** | **Protease** | **1° oxyanion hole hydrogen bond**  **(kcal/mol)** | | | | | | **2° oxyanion hole hydrogen bond**  **(kcal/mol)** | | | | | | **1° + 2°** | |
|  |  | Δ*E*_GS_† | | Δ*E*_TS_† | | ΔΔ*E*^‡^# | | Δ*E*_GS_ | | Δ*E*_TS_ | | ΔΔ*E*^‡^ | | ΔΔ*E*^‡^ | |
|  |  | *mean* | *s.d* | *mean* | *s.d* | *mean* | *s.d* | *mean* | *s.d* | *mean* | *s.d* | *mean* | *s.d* | *mean* | *s.d* |
| PA | Trypsin | 0.3 | 0.4 | 1.2 | 0.9 | -0.8 | 1.0 | 2.1 | 1.2 | 1.1 | 1.0 | 1.0 | 1.6 | 0.2 | 1.9 |
|  | Thrombin | 1.5 | 0.8 | 1.1 | 1.1 | 0.4 | 1.3 | 0.9 | 0.7 | 0.8 | 0.8 | 0.1 | 1.0 | 0.5 | 1.7 |
|  | Elastase | 1.4 | 1.0 | 1.9 | 1.3 | -0.5 | 1.6 | 1.5 | 1.1 | 2.6 | 1.2 | -1.1 | 1.6 | -1.6 | 2.3 |
|  | Chymotrypsin | 0.9 | 0.5 | 0.8 | 0.9 | 0.2 | 1.0 | 2.0 | 1.2 | 0.9 | 1.0 | 1.1 | 1.5 | 1.3 | 1.8 |
|  | Streptogrisin B | 0.6 | 0.6 | 1.0 | 0.6 | -0.4 | 0.8 | 4.3 | 0.7 | 1.4 | 0.9 | 2.9 | 1.1 | 2.4 | 1.4 |
| SB | Subtilisin | 0.4 | 0.3 | – | – | – | – | 0.5 | 0.4 | – | – | – | – | – | – |
|  | Proteinase K | 0.0 | – | 0.2 | – | -0.2 | – | 1.9 | – | 3.6 | – | -1.7 | – | -1.9 | – |
| SC | Dipeptidyl peptidase IV | 0.9 | 0.4 | 1.3 | – | -0.4 | 0.4 | 3.0 | 1.4 | 1.3 | – | 1.6 | 1.4 | 1.3 | 1.4 |
|  | Prolyl aminopeptidase | 1.1 | 1.5 | – | – | – | – | 1.3 | 0.5 | – | – | – | – | – | – |
| SE | D-Ala-D-Ala peptidase | 0.7 | 0.7 | 2.2 | 0.6 | -1.5 | 0.9 | 0.4 | 0.0 | 1.7 | 0.3 | -1.3 | 0.3 | -2.8 | 0.9 |

| **(B)** | ***α*_HB_** | | | | | | | | | | | | | | |
| --- | --- | --- | --- | --- | --- | --- | --- | --- | --- | --- | --- | --- | --- | --- | --- |
| **Clan** | **Protease** | **1° oxyanion hole hydrogen bond**  **(kcal/mol)** | | | | | | **2° oxyanion hole hydrogen bond**  **(kcal/mol)** | | | | | | **1° + 2°** | |
|  |  | Δ*E*_GS_ | | Δ*E*_TS_ | | ΔΔ*E*^‡^ | | Δ*E*_GS_ | | Δ*E*_TS_ | | ΔΔ*E*^‡^ | | ΔΔ*E*^‡^ | |
|  |  | *mean* | *s.d* | *mean* | *s.d* | *mean* | *s.d* | *mean* | *s.d* | *mean* | *s.d* | *mean* | *s.d* | *mean* | *s.d* |
| PA | Trypsin | 0.2 | 0.2 | 0.6 | 0.5 | -0.3 | 0.5 | 0.2 | 0.2 | 0.6 | 0.5 | -0.4 | 0.5 | -0.7 | 0.7 |
|  | Thrombin | 0.4 | 0.5 | 0.3 | 0.3 | 0.1 | 0.6 | 0.4 | 0.1 | 0.7 | 0.6 | -0.4 | 0.6 | -0.3 | 0.8 |
|  | Elastase | 0.3 | 0.2 | 0.2 | 0.1 | 0.1 | 0.2 | 0.3 | 0.1 | 0.5 | 0.3 | -0.2 | 0.3 | -0.1 | 0.4 |
|  | Chymotrypsin | 0.1 | 0.1 | 0.6 | 0.4 | -0.5 | 0.4 | 0.4 | 0.1 | 1.1 | 0.8 | -0.8 | 0.8 | -1.2 | 0.9 |
|  | Streptogrisin B | 0.1 | 0.1 | 0.8 | 0.5 | -0.7 | 0.5 | 0.3 | 0.1 | 0.3 | 0.0 | 0.0 | 0.1 | -0.7 | 0.5 |
| SB | Subtilisin | 0.1 | 0.1 | – | – | – | – | 1.8 | 0.2 | – | – | – | – | – | – |
|  | Proteinase K | 0.4 | – | 0.8 | – | -0.4 | – | 3.1 | – | 0.0 | – | 3.1 | – | 2.7 | – |
| SC | Dipeptidyl peptidase IV | 0.3 | 0.0 | 0.2 | – | 0.2 | 0.0 | 0.3 | 0.2 | 0.3 | – | 0.0 | 0.2 | 0.2 | 0.2 |
|  | Prolyl aminopeptidase | 0.4 | 0.0 | – | – | – | – | 1.6 | 0.3 | – | – | – | – | – | – |
| SE | D-Ala-D-Ala peptidase | 0.2 | 0.2 | 0.6 | 0.1 | -0.4 | 0.2 | 0.2 | 0.2 | 0.4 | 0.3 | -0.2 | 0.3 | -0.6 | 0.4 |

| **(C)** | ***ϕ*_HB_** | | | | | | |
| --- | --- | --- | --- | --- | --- | --- | --- |
| **Clan** | **Protease** | **1° oxyanion hole  hydrogen bond**  **(kcal/mol)** | | **2° oxyanion hole**  **hydrogen bond**  **(kcal/mol)** | | **1° + 2°** | |
|  |  | Δ*E*_GS_ | | Δ*E*_TS_ | | ΔΔ*E*^‡^ | |
|  |  | *mean* | *s.d* | *mean* | *s.d.* | *mean* | *s.d.* |
| PA | Trypsin | 0.94 | 0.03 | 0.95 | 0.02 | 1.89 | 0.04 |
|  | Thrombin | 0.94 | 0.03 | 0.84 | 0.26 | 1.78 | 0.26 |
|  | Elastase | 0.91 | 0.05 | 0.89 | 0.07 | 1.80 | 0.09 |
|  | Chymotrypsin | 0.96 | 0.03 | 0.95 | 0.02 | 1.91 | 0.04 |
|  | Streptogrisin B | 0.91 | 0.01 | 0.96 | 0.02 | 1.87 | 0.02 |
| SB | Subtilisin | 0.94 | 0.02 | 0.97 | 0.02 | 1.91 | 0.03 |
|  | Proteinase K | 0.79 | – | 0.89 | – | 1.68 | – |
| SC | Dipeptidyl peptidase IV | 0.92 | 0.02 | 0.96 | 0.03 | 1.88 | 0.04 |
|  | Prolyl aminopeptidase | 0.92 | 0.02 | 0.87 | 0.04 | 1.79 | 0.04 |
| SE | D-Ala-D-Ala peptidase | 0.95 | 0.02 | 0.93 | 0.00 | 1.88 | 0.02 |

† Δ*E*_GS_ and Δ*E*_TS_ are the differences between the average energy value calculated from observed geometries (in the GSA-bound and TSA-bound pseudo-ensembles, respectively) and the energy minimum of their corresponding energy functions, with positive values indicating destabilization.

\# ΔΔ*E*^‡^ is the difference between Δ*E*_GS_ and Δ*E*_TS_ (Δ*E*_GS_ – Δ*E*_TS_).

**Table S17.** Hydrogen bond geometric parameters for the His•Asp hydrogen bond of the catalytic triad in serine protease pseudo-ensembles (“Enzyme”) and in analogous hydrogen bonds in small molecules and PDB structures (“Reference”). The geometric parameters, **(A)** *d*_HB_, **(B)** *α*_HB_ and **(C)** *ϕ*_HB_, are defined in fig. S37.

| **(A)** | **Clan** | **Protease** | **E•GSA**  *n* | **E•TSA**  *n* | ***d*_HB_ (Å)** | | | | | |
| --- | --- | --- | --- | --- | --- | --- | --- | --- | --- | --- |
|  |  |  |  |  | **E•GSA** | | **E•TSA** | | **E•TSA - E•GSA** | |
|  |  |  |  |  | *mode* | *s.d.* | *mode* | *s.d.* | Δ | *p-*value |
| Enzyme† | PA | Alpha-Lytic Protease | 0 | 6 | – | – | 2.77 | 0.04 | – | – |
|  |  | Chymotrypsin | 21 | 13 | 2.64 | 0.07 | 2.72 | 0.10 | 0.08 | 3E-01 |
|  |  | Elastase | 7 | 9 | 2.69 | 0.13 | 2.66 | 0.11 | -0.03 | 4E-01 |
|  |  | Streptogrisin B | 16 | 3 | 2.82 | 0.07 | 2.72 | 0.01 | -0.10 | 2E-02 |
|  |  | Thrombin | 9 | 25 | 2.64 | 0.07 | 2.70 | 0.13 | 0.07 | 2E-01 |
|  |  | Trypsin | 79 | 13 | 2.74 | 0.07 | 2.66 | 0.11 | -0.08 | 2E-01 |
|  |  | *combined* | 132 | 69 | 2.74 | 0.08 | 2.71 | 0.11 | -0.03 | 3E-01 |
|  | SB | Kexin | 0 | 4 | – | – | 5.32 | 0.03 | – | – |
|  |  | Proteinase K | 1 | 1 | 2.73 | – | 2.70 | – | -0.02 | – |
|  |  | Sedolisin | 0 | 6 | – | – | 2.64 | 0.05 | – | – |
|  |  | Subtilisin | 22 | 0 | 2.72 | 0.04 | – | – | – | – |
|  |  | *combined* | 23 | 7 | 2.72 | 0.04 | 2.64 | 0.05 | -0.08 | 6E-04 |
|  | SC | Dipeptidyl peptidase IV | 6 | 1 | 2.74 | 0.05 | 2.82 | – | 0.09 | – |
|  |  | Prolyl aminopeptidase | 3 | 0 | 2.55 | 0.01 | – | – | – | – |
|  |  | Prolyl endopeptidase | 0 | 6 | – | – | 2.77 | 0.13 | – | – |
|  |  | Serine carboxypeptidase A | 0 | 2 | – | – | 2.69 | 0.01 | – | – |
|  |  | *combined* | 9 | 9 | 2.73 | 0.10 | 2.76 | 0.12 | 0.02 | 4E-02 |
|  | *combined* | | 164 | 85 | 2.73 | 0.08 | 2.71 | 0.11 | -0.02 | 6E-01 |
| Reference | imidazole•carboxylate (CSD)‡ | | 240 | 219 | 2.75 | 0.12 | 2.69 | 0.14 | -0.06 | 7E-02 |
|  | His•Asp (PDB)¶ | | 659 | 1114 | 2.76 | 0.41 | 2.73 | 0.39 | -0.03 | 3E-03 |

| **(B)** | **Clan** | **Protease** | ***α*_HB_ (°)** | | | | | |
| --- | --- | --- | --- | --- | --- | --- | --- | --- |
|  |  |  | **E•GSA** | | **E•TSA** | | **E•TSA - E•GSA** | |
|  |  |  | *mode* | *s.d.* | *mode* | *mode* | Δ | *p-*value |
| Enzyme† | PA | Alpha-Lytic Protease | – | – | 114 | 2 | – | – |
|  |  | Chymotrypsin | 116 | 3 | 122 | 4 | 5.09 | 7E-03 |
|  |  | Elastase | 114 | 3 | 116 | 3 | 2.27 | 5E-01 |
|  |  | Streptogrisin B | 110 | 2 | 113 | 0 | 3.42 | 4E-03 |
|  |  | Thrombin | 118 | 3 | 117 | 7 | -0.40 | 1E+00 |
|  |  | Trypsin | 112 | 3 | 118 | 3 | 5.20 | 2E-08 |
|  |  | *combined* | 113 | 3 | 117 | 5 | 4.31 | 7E-08 |
|  | SB | Kexin | – | – | 94 | 2 | – | – |
|  |  | Proteinase K | 105 | – | 105 | – | 0.04 | – |
|  |  | Sedolisin | – | – | 121 | 6 | – | – |
|  |  | Subtilisin | 107 | 4 | – | – | – | – |
|  |  | *combined* | 107 | 4 | 121 | 7 | 13.54 | 4E-04 |
|  | SC | Dipeptidyl peptidase IV | 90 | 2 | 82 | ­– | -8.34 | – |
|  |  | Prolyl aminopeptidase | 101 | 1 | – | – | – | – |
|  |  | Prolyl endopeptidase | – | – | 102 | 3 | – | – |
|  |  | Serine carboxypeptidase A | – | – | 108 | 1 | – | – |
|  |  | *combined* | 90 | 6 | 105 | 8 | 15.50 | 2E-02 |
|  | *combined* | | 113 | 3 | 117 | 7 | 4.86 | 3E-05 |
| Reference | imidazole•carboxylate (CSD)‡ | | 126 | 14 | 123 | 13 | -3.07 | 1E-01 |
|  | His•Asp (PDB)¶ | | 115 | 20 | 118 | 19 | 3.39 | 4E-02 |

| **(C)** | **Clan** | **Protease** | ***ϕ*_HB_ (°)** | | | | | |
| --- | --- | --- | --- | --- | --- | --- | --- | --- |
|  |  |  | **E•GSA** | | **E•TSA** | | **E•TSA - E•GSA** | |
|  |  |  | *mode* | *s.d.* | *mode* | *s.d.* | Δ | *p-*value |
| Enzyme† | PA | Alpha-Lytic Protease | – | – | 7 | 3 | – | – |
|  |  | Chymotrypsin | 5 | 4 | 8 | 6 | 2.59 | 4E-02 |
|  |  | Elastase | 4 | 3 | 3 | 7 | -0.54 | 9E-01 |
|  |  | Streptogrisin B | 9 | 4 | 2 | 1 | -7.71 | 8E-02 |
|  |  | Thrombin | 11 | 4 | 10 | 8 | -1.30 | 7E-01 |
|  |  | Trypsin | 10 | 3 | 10 | 7 | 0.16 | 1E-01 |
|  |  | *combined* | 10 | 4 | 7 | 7 | -3.27 | 2E-01 |
|  | SB | Kexin | – | – | 71 | 4 | – | – |
|  |  | Proteinase K | 4 | – | 5 | – | 0.86 |  |
|  |  | Sedolisin | – | – | 22 | 5 | – | – |
|  |  | Subtilisin | 3 | 3 | – | – | – | – |
|  |  | *combined* | 3 | 3 | 22 | 7 | 19.32 | 3E-09 |
|  | SC | Dipeptidyl peptidase IV | 0 | 0 | 0 | – | 0.00 | – |
|  |  | Prolyl aminopeptidase | 26 | 2 | – | – | – | – |
|  |  | Prolyl endopeptidase | – | – | 22 | 3 | – | – |
|  |  | Serine carboxypeptidase A | – | – | 45 | 4 | – | – |
|  |  | *combined* | 0 | 13 | 23 | 14 | 22.96 | 2E-02 |
|  | *combined* | | 10 | 4 | 7 | 9 | -3.06 | 8E-06 |
| Reference | imidazole•carboxylate (CSD)‡ | | 9 | 17 | 8 | 17 | -1.39 | 4E-01 |
|  | His•Asp (PDB)¶ | | 14 | 27 | 12 | 26 | -1.79 | 2E-02 |

† Only enzymes structures with His•Asp distance (*d*_HB_) < 4 Å were included.

‡ Neutral imidazole•carboxylate hydrogen bonds under “E•GSA” and protonated imidazole•carboxylate hydrogen bonds under “E•TSA”

¶ His•Asp hydrogen bonds found in high quality PDB structures from the Top2018 library (*107*), where “E•GSA” contains structures crystallized at pH > 8 and “E•TSA” contains structures crystallized at pH < 6.

**Table S18**. Energy differences resulting from the changes in the catalytic triad His•Asp hydrogen bond geometries in going from the GSA-bound to TSA-bound states, calculated by mapping geometric measurements from pseudo-ensembles on knowledge-based energy functions. The geometric parameters, *d*_HB_, *α*_HB_ and *ϕ*_HB_, defined in fig. S37 and their values are listed in table S17.

| **Clan** | **Protease** | ***d*_HB_**  **(kcal/mol)** | | | | | | ***α*_HB_**  **(kcal/mol)** | | | | | | ***ϕ*_HB_**  **(kcal/mol)** | | | | | |
| --- | --- | --- | --- | --- | --- | --- | --- | --- | --- | --- | --- | --- | --- | --- | --- | --- | --- | --- | --- |
|  |  | Δ*E*_GS_† | | Δ*E*_TS_¶ | | ΔΔ*E*^‡^# | | Δ*E*_GS_ | | Δ*E*_TS_ | | ΔΔ*E*^‡^ | | Δ*E*_GS_ | | Δ*E*_TS_ | | ΔΔ*E*^‡^ | |
|  |  | *mean* | *s.d.* | *mean* | *s.d.* | *mean* | *s.d.* | *mean* | *s.d.* | *mean* | *s.d.* | *mean* | *s.d.* | *mean* | *s.d.* | *mean* | *s.d.* | *mean* | *s.d.* |
| PA | Chymotrypsin | 0.4 | 0.2 | 0.4 | 0.5 | 0.0 | 0.5 | 0.1 | 0.1 | 0.1 | 0.1 | 0.0 | 0.1 | 0.1 | 0.1 | 0.3 | 0.1 | -0.2 | 0.1 |
|  | Elastase | 0.4 | 0.3 | 0.4 | 0.3 | 0.0 | 0.4 | 0.0 | 0.0 | 0.1 | 0.0 | 0.0 | 0.1 | 0.1 | 0.1 | 0.2 | 0.1 | -0.1 | 0.1 |
|  | Streptogrisin B | 0.3 | 0.3 | 0.0 | 0.0 | 0.3 | 0.3 | 0.2 | 0.1 | 0.1 | 0.0 | 0.0 | 0.1 | 0.1 | 0.1 | 0.1 | 0.1 | 0.0 | 0.1 |
|  | Thrombin | 0.3 | 0.3 | 0.4 | 0.4 | -0.1 | 0.5 | 0.1 | 0.1 | 0.1 | 0.2 | 0.0 | 0.2 | 0.2 | 0.1 | 0.3 | 0.1 | -0.1 | 0.2 |
|  | Trypsin | 0.2 | 0.2 | 0.4 | 0.3 | -0.2 | 0.4 | 0.2 | 0.1 | 0.1 | 0.0 | 0.1 | 0.1 | 0.2 | 0.1 | 0.3 | 0.1 | -0.1 | 0.1 |
| SB | Proteinase K | 0.1 | – | 0.1 | – | 0 | – | 0.2 | – | 0.1 | – | 0.0 | – | 0.0 | – | 0.2 | – | -0.2 | – |
|  | Subtilisin | 0.2 | 0.1 | – | – | – | – | 0.1 | 0.1 | – | – | – | – | 0.1 | – | – | – | – |  |
| SC | Dipeptidyl peptidase IV | 0.2 | 0.1 | 0.3 | 0.0 | -0.1 | 0.1 | 1.1 | 0.2 | 1.6 | – | -0.5 | 0.2 | 0.2 | 0.0 | 0.2 | 0.0 | -0.1 | 0.0 |
|  | Prolyl aminopeptidase | 0.9 | 0.0 | – | – | – | – | 0.4 | 0.0 | – | – | – | – | 0.3 | 0.1 | – | – | – | – |

† ΔE_GS_ energy values were calculated by mapping His•Asp geometric parameters measured in GSA-bound pseudo-ensembles on knowledge-based energy functions derived from His•Asp hydrogen bonds found in the *Top2018* library (*107*) crystallized under pH > 8 (shown in fig. S35A). The small molecule distributions are similar to the PDB distributions but were not used for energy function derivation here because of their small sample sizes.

¶ ΔE_TS_ energy values were calculated by mapping His•Asp geometric parameters measured in TSA-bound pseudo-ensembles on knowledge-based energy functions derived from His•Asp hydrogen bonds found in the *Top2018* library (*107*) crystallized under pH < 6 (shown in fig. S35B).

#ΔΔ*E*^‡^ is the difference between Δ*E*_GS_ and Δ*E*_TS_ (Δ*E*_GS_ – Δ*E*_TS_).

**Table S19.** Hydrogen bond geometric parameters for the His•Ser hydrogen bond of the catalytic triad in serine protease pseudo-ensembles (“Enzyme”) and in analogous hydrogen bonds in small molecules and PDB structures (“Reference”). The geometric parameters, **(A)** *d*_HB_, **(B)** *α*_HB_ and **(C)** *ϕ*_HB_, are defined in fig. S40.

| **(A)** | **Clan** | **Protease** | **E•GSA**  *n* | **E•TSA**  *n* | ***d*_HB_ (Å)** | | | | | |
| --- | --- | --- | --- | --- | --- | --- | --- | --- | --- | --- |
|  |  |  |  |  | **E•GSA** | | **E•TSA** | | **E•TSA - E•GSA** | |
|  |  |  |  |  | *mode* | *s.d.* | *mode* | *s.d.* | Δ | *p-*value |
| Enzyme† | PA | Alpha-Lytic Protease | 0 | 6 | – | – | 3.01 | 0.06 | – | – |
|  |  | Chymotrypsin | 21 | 13 | 2.65 | 0.22 | 2.99 | 0.42 | 0.34 | 1E-03 |
|  |  | Elastase | 7 | 9 | 2.66 | 0.19 | 2.83 | 0.20 | 0.17 | 1E-02 |
|  |  | Streptogrisin B | 16 | 3 | 2.64 | 0.07 | 3.09 | 0.11 | 0.45 | 3E-09 |
|  |  | Thrombin | 10 | 25 | 3.03 | 0.18 | 2.80 | 0.23 | -0.23 | 4E-01 |
|  |  | Trypsin | 79 | 13 | 2.67 | 0.09 | 2.86 | 0.16 | 0.18 | 3E-10 |
|  |  | *combined* | 133 | 69 | 2.66 | 0.15 | 2.87 | 0.26 | 0.20 | 2E-21 |
|  | SB | Kexin | 0 | 4 | – | – | 3.22 | 0.20 | – | – |
|  |  | Proteinase K | 1 | 1 | 3.09 | – | 3.20 | – | 0.11 | – |
|  |  | Sedolisin | 0 | 7 | – | – | 3.06 | 0.77 | – | – |
|  |  | Subtilisin | 22 | 0 | 2.71 | 0.06 | – | – | – | – |
|  |  | *combined* | 23 | 12 | 2.70 | 0.10 | 3.16 | 0.58 | 0.47 | 8E-09 |
|  | SC | Dipeptidyl peptidase IV | 6 | 1 | 2.71 | 0.10 | – | – | – | – |
|  |  | Prolyl aminopeptidase | 3 | 0 | 2.66 | 0.09 | – | – | – | – |
|  |  | Prolyl endopeptidase | 0 | 6 | – | – | 2.95 | 0.15 | – | – |
|  |  | Serine carboxypeptidase A | 0 | 2 | – | – | 2.85 | 0.18 | – | – |
|  |  | *combined* | 9 | 9 | 2.69 | 0.11 | 2.94 | 0.18 | 0.25 | 3E-03 |
|  | *combined* | | 165 | 94 | 2.67 | 0.14 | 2.92 | 0.25 | 0.25 | 1E-29 |
| Reference | imidazole•hydroxyl (CSD)‡ | | 102 | 92 | 2.77 | 0.40 | 2.81 | 0.26 | 0.04 | 3E-01 |
|  | His•Ser (PDB)¶ | | 659 | 815 | 2.78 | 3.20 | 2.77 | 0.44 | -0.01 | 3E-01 |

| **(B)** | **Clan** | **Protease** | ***α*_HB_ (°)** | | | | | |
| --- | --- | --- | --- | --- | --- | --- | --- | --- |
|  |  |  | **E•GSA** | | **E•TSA** | | **E•TSA - E•GSA** | |
|  |  |  | *mode* | *s.d.* | *mode* | *s.d.* | Δ | *p-*value |
| Enzyme† | PA | Alpha-Lytic Protease | – | – | 89 | 3 | – | – |
|  |  | Chymotrypsin | 98 | 8 | 88 | 12 | -10 | 3E-02 |
|  |  | Elastase | 97 | 8 | 105 | 6 | 8 | 9E-01 |
|  |  | Streptogrisin B | 101 | 5 | 90 | 4 | -11 | 5E-04 |
|  |  | Thrombin | 74 | 12 | 90 | 10 | 16 | 2E-01 |
|  |  | Trypsin | 100 | 5 | 93 | 12 | -6 | 1E-01 |
|  |  | *combined* | 99 | 8 | 90 | 11 | -9 | 1E-06 |
|  | SB | Kexin | – | – | 98 | 12 | – | – |
|  |  | Proteinase K | 99 | – | 93 | – | -7 | – |
|  |  | Sedolisin | – | – | 107 | 18 | – | – |
|  |  | Subtilisin | 107 | 4 | – | – | – | – |
|  |  | *combined* | 107 | 4 | 103 | 16 | -4 | 3E-02 |
|  | SC | Dipeptidyl peptidase IV | 93 | 4 | – | – | – | – |
|  |  | Prolyl aminopeptidase | 90 | 5 | – | – | – | – |
|  |  | Prolyl endopeptidase | – | – | 95 | 9 | – | – |
|  |  | Serine carboxypeptidase A | – | – | 90 | 3 | – | – |
|  |  | *combined* | 93 | 4 | 95 | 8 | 2 | 6E-01 |
|  | *combined* | | 99 | 8 | 91 | 11 | -8 | 3E-07 |
| Reference | imidazole•hydroxyl (CSD)‡ | | 122 | 19 | 121 | 12 | -2 | 3E-01 |
|  | His•Ser (PDB)¶ | | 124 | 29 | 124 | 31 | 1 | 6E-01 |

| **(C)** | **Clan** | **Protease** | ***ϕ*_HB_ (°)** | | | | | |
| --- | --- | --- | --- | --- | --- | --- | --- | --- |
|  |  |  | **E•GSA** | | **E•TSA** | | **E•TSA - E•GSA** | |
|  |  |  | *mode* | *s.d.* | *mode* | *s.d.* | Δ | *p-*value |
| Enzyme† | PA | Alpha-Lytic Protease | – | – | 8 | 2 | – | – |
|  |  | Chymotrypsin | 2 | 6 | 3 | 4 | 1 | 4E-01 |
|  |  | Elastase | 4 | 6 | 5 | 3 | 1 | 3E-01 |
|  |  | Streptogrisin B | 1 | 2 | 8 | 2 | 7 | 1E-04 |
|  |  | Thrombin | 13 | 6 | 10 | 5 | -3 | 1E-01 |
|  |  | Trypsin | 1 | 3 | 3 | 5 | 2 | 6E-04 |
|  |  | *combined* | 2 | 5 | 6 | 5 | 4 | 1E-05 |
|  | SB | Kexin | – | – | 2 | 4 | – | – |
|  |  | Proteinase K | 6 | – | 7 | – | 0 | – |
|  |  | Sedolisin | – | – | 29 | 8 | – | – |
|  |  | Subtilisin | 1 | 2 |  |  |  |  |
|  |  | *combined* | 1 | 2 | 8 | 12 | 7 | 2E-05 |
|  | SC | Dipeptidyl peptidase IV | 8 | 3 | – | – | – | – |
|  |  | Prolyl aminopeptidase | 4 | 2 | – | – | – | – |
|  |  | Prolyl endopeptidase | – | – | 7 | 2 | – | – |
|  |  | Serine carboxypeptidase A | – | – | 2 | 1 | – | – |
|  |  | *combined* | 6 | 3 | 6 | 4 | 0 | 5E-01 |
|  | *combined* | | 2 | 4 | 6 | 7 | 4 | 3E-09 |
| Reference | imidazole•hydroxyl (CSD)‡ | | 12 | 24 | 5 | 15 | -7 | 2E-05 |
|  | His•Ser (PDB)¶ | | 10 | 27 | 9 | 28 | -1 | 6E-03 |

† Only structures with His•Ser distance (*d*_HB_) < 4 Å were included

‡ Neutral imidazole•hydroxyl hydrogen bonds under “E•GSA” and protonated imidazole•ether/hydroxyl hydrogen bonds under “E•TSA”

¶ His•Ser hydrogen bonds found in high quality PDB structures from the Top2018 library (*93*), where “E•GSA” contains structures crystallized at pH > 8 and “E•TSA” contains structures crystallized at pH < 6.

**Table S20**. Energy differences resulting from the changes in the catalytic triad His•Ser hydrogen bond geometries in going from the GSA-bound to TSA-bound states, calculated by mapping geometric measurements from pseudo-ensembles on knowledge-based energy functions. The geometric parameters, *d*_HB_, *α*_HB_ and *ϕ*_HB_, defined in fig. S40, and their values are listed in table S19.

| **Clan** | **Protease** | ***d*_HB_**  **(kcal/mol)** | | | | | | ***α*_HB_**  **(kcal/mol)** | | | | | | ***ϕ*_HB_**  **(kcal/mol)** | | | | | |
| --- | --- | --- | --- | --- | --- | --- | --- | --- | --- | --- | --- | --- | --- | --- | --- | --- | --- | --- | --- |
|  |  | Δ*E*_GS_† | | Δ*E*_TS_¶ | | ΔΔ*E*^‡^# | | Δ*E*_GS_ | | Δ*E*_TS_ | | ΔΔ*E*^‡^ | | Δ*E*_GS_ | | Δ*E*_TS_ | | ΔΔ*E*^‡^ | |
|  |  | *mean* | *s.d.* | *mean* | *s.d.* | *mean* | *s.d.* | *mean* | *s.d.* | *mean* | *s.d.* | *mean* | *s.d.* | *mean* | *s.d.* | *mean* | *s.d.* | *mean* | *s.d.* |
| PA | Chymotrypsin | 0.5 | 0.1 | 0.5 | 0.1 | 0.0 | 0.1 | 0.7 | 0.7 | 0.5 | 0.3 | 0.3 | 0.8 | 0.1 | 0.2 | 0.1 | 0.1 | 0.0 | 0.2 |
|  | Elastase | 0.4 | 0.1 | 0.4 | 0.1 | 0.0 | 0.1 | 1.0 | 0.9 | 0.4 | 0.3 | 0.6 | 0.9 | 0.2 | 0.2 | 0.1 | 0.1 | 0.1 | 0.2 |
|  | Streptogrisin B | 0.4 | 0.1 | 0.5 | 0.1 | -0.1 | 0.2 | 0.9 | 0.7 | 0.8 | 0.1 | 0.1 | 0.7 | 0.1 | 0.1 | 0.1 | 0.0 | 0.0 | 0.1 |
|  | Thrombin | 0.6 | 0.1 | 0.5 | 0.1 | 0.1 | 0.1 | 0.6 | 0.4 | 0.6 | 0.4 | 0.0 | 0.5 | 0.3 | 0.2 | 0.2 | 0.1 | 0.1 | 0.2 |
|  | Trypsin | 0.5 | 0.1 | 0.4 | 0.2 | 0.0 | 0.2 | 0.7 | 0.7 | 0.5 | 0.3 | 0.2 | 0.8 | 0.1 | 0.1 | 0.2 | 0.1 | -0.1 | 0.1 |
| SB | Proteinase K | 0.5 | – | 0.7 | – | -0.1 | 0.0 | 0.7 | – | 1.0 | – | -0.2 | – | 0.2 | – | 0.2 | – | 0.0 | – |
|  | Subtilisin | 0.5 | 0.1 | – | – | – | – | 0.3 | 0.4 | – | – | – | – | 0.1 | 0.1 | – | – | – | – |
| SC | Dipeptidyl peptidase IV | 0.5 | 0.1 | 0.4 | 0.0 | 0.1 | 0.1 | 0.2 | 0.2 | 0.2 | 0.0 | 0.1 | 0.2 | 0.2 | 0.1 | 0.4 | 0.0 | -0.2 | 0.1 |
|  | Prolyl aminopeptidase | 0.5 | 0.1 | – | – | – | – | 0.8 | 0.8 | – | – | – | – | 0.1 | 0.1 | – | – | – | – |

† ΔE_GS_ energy values were calculated by mapping His•Ser geometric parameters measured in GSA-bound pseudo-ensembles on knowledge-based energy functions derived from His•Ser hydrogen bonds found in the *Top2018* library (*92*) (shown in fig. S38A). The small molecule distribution is similar to the PDB distribution (fig. S38A) but they were not used for energy function derivation here because of their small sample sizes.

¶ ΔE_TS_ energy values were calculated by mapping His•Ser geometric parameters measured in TSA-bound pseudo-ensembles on knowledge-based energy functions derived from His•Ser hydrogen bonds found in the *Top2018* library (*6*) (shown in fig. S38B).

\# ΔΔ*E*^‡^ is the difference between Δ*E*_GS_ and Δ*E*_TS_ (Δ*E*_GS_ – Δ*E*_TS_).

**Table S21.** Enzymes that perform nucleophilic attack reaction and contain oxyanion holes curated from the Mechanism and Catalytic Site Atlas (M-CSA) database (*24*).

| **Reaction*** | **Oxyanion hole type†** | **Superfamily (CATH) ‡** | **Name** | **Nucl. §** | **Ref PDB** | **Chain**  **ID** | **Nucl. residue ID** | **Oyanion hole**  **H-bond donor 1**¶ | **Oyanion hole H-bond donor 2**¶ | **Oyanion hole H-bond donor 3**¶ | **Protease**# |
| --- | --- | --- | --- | --- | --- | --- | --- | --- | --- | --- | --- |
| Nucleophilic  aromatic substitution | Other | 3.90.226.10 | 4-chlorobenzoyl-CoA  dehalogenase | Asp | 1nzy | A | 145 | Phe:64:N | Gly:114:N | – | N |
| Nucleophilic addition on carbonyl | N+1-type NE | 3.30.360.10 | glyceraldehyde-3-phosphate dehydrogenase (NAD(P)+) (phosphorylating) | Cys | 1cf2 | A | 140 | Asn:141:N | Asn:141:ND2 | – | N |
|  |  | 3.40.1090.10 | phospholipase A2  (group group IVA) | Ser | 1cjy | A | 228 | Gly:229:N | Gly:197:N | Gly:198:N | N |
|  |  | 3.40.366.10 | malonyl-CoA-acyl  carrier protein transacylase | Ser | 1mla | A | 92 | Leu:93:N | Gln:11:N | – | N |
|  |  | 3.40.50.180 | protein-glutamate  methylesterase (CheB) | Ser | 1chd | A | 164 | Thr:165:N | Met:283:N | – | N |
|  |  | 3.40.50.1820 | 2-hydroxymuconate-  semialdehyde hydrolase | Ser | 1uk7 | A | 103 | Phe:104:N | Ser:34:N | – | N |
|  |  |  | 6-deoxyerythronolide-B  synthase | Ser | 1kez | A | 142 | Ala:143:N | – | – | N |
|  |  |  | acetylcholinesterase | Ser | 1mah | A | 203 | Ala:204:N | Gly:121:N | Gly:122:N | N |
|  |  |  | alpha-amino-acid  esterase | Ser | 1mpx | A | 174 | Tyr:175:N | Tyr:82:OH | – | N |
|  |  |  | carboxylesterase | Ser | 2o7r | A | 169 | Ala:170:N | Gly:93:N | Gly:92:N | N |
|  |  |  | carboxypeptidase D | Ser | 1whs | A | 146 | Gly:53:N | Tyr:147:N | – | P |
|  |  |  | cephalosporin-C  deacetylase | Ser | 1l7a | A | 181 | Gln:182:N | Tyr:91:N | – | N |
|  |  |  | chloride peroxidase  (cofactor free) | Ser | 1a7u | A | 98 | Met:99:N | Phe:32:N | – | N |
|  |  |  | cutinase | Ser | 1agy | A | 120 | Gln:121:N | Ser:42:N | – | N |
|  |  |  | esterase | Ser | 1zoi | A | 97 | Thr:98:N | Trp:31:N | – | N |
|  |  |  | myristoyl-ACP-specific  thioesterase | Ser | 1tht | A | 114 | Leu:115:N | – | – | N |
|  |  |  | palmitoyl[protein]  hydrolase (type 2) | Ser | 1pja | A | 111 | Gln:112:N | Leu:45:N | – | N |
|  |  |  | para-nitrobenzyl esterase | Ser | 1qe3 | A | 189 | Ala:190:N | Gly:106:N | Ala:107:N | N |
|  |  |  | triacylglycerol lipase  (EstA) | Ser | 1r4z | A | 77 | Met:78:N | Ile:12:N | – | N |
|  |  |  | triacylglycerol lipase  (Pseudomonas family) | Ser | 1tah | A | 87 | Glu:88:N | Leu:17:N | – | N |
|  |  |  | triacylglycerol lipase  (pancreatic) | Ser | 1hpl | A | 152 | Leu:153:N | Phe:77:N | – | N |
|  |  |  | triacylglycerol lipase  (type B carboxylestrase) | Ser | 1thg | A | 217 | Ala:218:N | Ala:132:N | – | N |
|  |  | 3.40.50.880 | GMP synthase  (glutamine-hydrolysing) | Cys | 1gpm | A | 86 | Tyr:87:N | Gly:59:N | – | N |
|  |  |  | anthranilate synthase | Cys | 1qdl | B | 84 | Leu:85:N | Gly:56:N | – | N |
|  |  |  | intracellular protease | Cys | 1g2i | A | 100 | His:101:N | Gly:70:N | – | P |
|  |  | 3.60.110.10 | N-carbamoyl-D-amino-acid  hydrolase | Cys | 1fo6 | A | 172 | Asn:173:N | Lys:127:NZ | – | N |
|  | N-type NE | 1.10.1500.10 | glutaminase | Ser | 1mki | A | 74 | Ser:74:N | Val:271:N | – | N |
|  |  | 2.10.109.10 | repressor LexA | Ser | 1jhf | A | 119 | Ser:119:N | Glu:152:N | – | P |
|  |  |  | signal peptidase I | Ser | 1t7d | B | 90 | Ser:90:N | Ser:88:OG | – | P |
|  |  | 2.20.210.10 | ubiquitinyl hydrolase 1  (peptidase C19 type) | Cys | 1nbf | A | 223 | Cys:223:N | Asn:218:ND2 | – | P |
|  |  | 2.40.10.10 | classical-complement-pathway  C3 C5 convertase | Ser | 2odq | A | 659 | Ser:659:N | – | – | P |
|  |  |  | ubiquitinyl hydrolase 1  (peptidase C30 type) | Cys | 2bx4 | A | 145 | Cys:145:N | Cys:143:N | – | P |
|  |  | 3.40.309.10 | aldehyde dehydrogenase  (NAD+) (class 2) | Cys | 1o04 | A | 302 | Cys:302:N | Asn:167:ND2 | – | N |
|  |  |  | betaine-aldehyde  dehydrogenase | Cys | 1a4s | A | 297 | Cys:297:N | Asn:166:N | – | N |
|  |  |  | glyceraldehyde-3-phosphate dehydrogenase (NADP+) | Cys | 2esd | A | 284 | Cys:284:N | Asn:154:ND2 | – | N |
|  |  |  | Ulp1 peptidase | Cys | 2bkr | A | 163 | Cys:163:N | Trp:26:NE1 | – | P |
|  |  |  | adenain | Cys | 1nln | A | 122 | Cys:122:N | Gln:115:NE2 | – | P |
|  |  | 3.40.47.10 | acetyl-CoA  C-acyltransferase | Cys | 1afw | A | 125 | Cys:125:N | Gly:405:N | – | N |
|  |  |  | beta-ketoacyl-  [acyl carrier protein]  synthase I | Cys | 1dd8 | A | 163 | Cys:163:N | Phe:392:N | – | N |
|  |  |  | beta-ketoacyl-  [acyl-carrier-protein]  synthase II | Cys | 1kas | A | 163 | Cys:163:N | Phe:400:N | – | N |
|  |  |  | naringenin-chalcone  synthase | Cys | 1cgk | A | 164 | Cys:164:N | – | – | N |
|  |  | 3.40.50.1110 | 1-alkyl-2-acetylglycerophosphocholine  esterase | Ser | 1wab | A | 47 | Ser:47:N | Gly:74:N | Asn:104:ND2 | N |
|  |  |  | esterase (estA) | Ser | 1esc | A | 14 | Ser:14:N | Gly:66:N | Asn:106:ND2 | N |
|  |  |  | lysophospholipase | Ser | 1j00 | A | 10 | Ser:10:N | Gly:44:N | Asn:73:ND2 | N |
|  |  |  | rhamnogalacturonan  acetylesterase | Ser | 1pp4 | A | 9 | Ser:9:N | Gly:42:N | – | N |
|  |  |  | asparaginase | Thr | 3eca | A | 12 | Thr:12:N | Thr:89:N | – | N |
|  |  |  | glutamin-(asparagin-)ase | Thr | 1djo | A | 1020 | Thr:1020:N | Thr:1100:N | – | N |
|  |  | 3.40.50.1460 | caspase-1 | Cys | 2fqq | A | 285 | Cys:285:N | Gly:238:N | – | P |
|  |  |  | caspase-3 | Cys | 1cp3 | A | 163 | Cys:163:N | Gly:121:N | – | P |
|  |  |  | caspase-9 | Cys | 1nw9 | B | 287 | Cys:287:N | Gly:238:N | – | P |
|  |  |  | gingipain R | Cys | 1cvr | A | 244 | Cys:244:N | Gly:212:N | – | P |
|  |  | 3.40.50.850 | N-carbamoylsarcosine  amidase | Cys | 1nba | A | 177 | Thr:173:OG1 | Cys:177:N | – | N |
|  |  |  | nicotinamidase | Cys | 1im5 | A | 133 | Cys:133:N | Ala:129:N | – | N |
|  |  | 3.40.630.20 | pyroglutamyl-peptidase I | Cys | 1-Aug | A | 144 | Cys:144:N | Arg:91:NH | – | P |
|  |  | 3.40.710.10 | beta-lactamase  (Class A) | Ser | 1btl | A | 70 | Ser:70:N | Ala:237:N | – | N |
|  |  |  | beta-lactamase  (Class C) | Ser | 1xx2 | A | 64 | Ser:64:N | Ser:318:N | – | N |
|  |  |  | beta-lactamase  (Class D) | Ser | 1m6k | A | 67 | Ser:67:N | Ala:215:N | – | N |
|  |  | 3.50.80.10 | D-aminoacyl-tRNA deacylase | Thr | 1j7g | A | 80 | Thr:80:N | Phe:79:N | Gln:78:NE2 | N |
|  |  | 3.90.1300.10 | fatty acid amide hydrolase | Ser | 1mt5 | A | 241 | Ser:241:N | Gly:240:N | Gly:239:N | N |
|  |  |  | peptide amidase | Ser | 1ocl | A | 155 | Ser:155:N | Gly:154:N | Gly:153:N | N |
|  |  | 3.90.1360.10 | protein-glutamine  gamma-glutamyltransferase  (bacterial) | Cys | 1iu4 | A | 64 | Cys:64:N | Trp:272:NE1 | – | N |
|  |  | 3.90.260.10 | protein-glutamine gamma-glutamyltransferase  (eukaryotic) | Cys | 1ggt | A | 314 | Cys:314:N | Trp:279:NE1 | – | N |
|  |  | 3.90.70.10 | L-peptidase | Cys | 1qol | B | 51 | Cys:51:N | Asn:46:ND2 | – | P |
|  |  |  | bleomycin hydrolase | Cys | 1gcb | A | 73 | Cys:73:N | Gln:67:NE2 | – | P |
|  |  |  | calpain-2 | Cys | 1kfu | A | 105 | Cys:105:N | Gln:99:NE2 | – | P |
|  |  |  | cathepsin S | Ser | 1glo | A | 25 | Cys:25:N | Gln:19:NE2 | – | P |
|  |  |  | staphopain | Cys | 1x9y | A | 243 | Cys:243:N | Gln:237:NE2 | – | P |
|  | Other | 1.10.10.2660 | E1 ubiquitin-activating  enzyme | Cys | 3cmm | A | 600 | Asn:781:N | Asp:782:N | – | N |
|  |  | 1.10.1040.10 | UDP-glucose 6-  dehydrogenase | Cys | 1dli | A | 260 | Lys:204:NZ | Asn:208:ND2 | – | N |
|  |  | 2.160.20.10 | pectinesterase | Asp | 1gq8 | A | 157 | Gln:135:NE2 | Gln:113:NE2 | – | N |
|  |  | 2.170.16.10 | GyrA intein  (Class 1 intein) | Ser | 1am2 | A | 1 | Thr:72:OG1 | Asn:74:ND2 | – | P |
|  |  |  | GyrA intein  (Class 1 intein) | Asn | 1am2 | A | 198 | Thr:72:OG1 | Asn:74:ND2 | – | P |
|  |  | 2.40.230.10 | phospholipase A1 | Ser | 1qd6 | C | 144 | Gly:146:N | water | – | N |
|  |  | 3.60.20.10 | HslU---HslV peptidase | Thr | 1ht1 | E | 1 | Gly:45:N | – | – | P |
|  |  |  | Amidophosphoribosyl-  transferase | Cys | 1ecf | A | 1 | Asn:101:ND2 | Gly:102:N | – | N |
|  |  |  | asparagine synthase  (glutamine-hydrolysing) | Cys | 1ct9 | A | 1 | Asn:74:ND2 | Gly:75:N | – | N |
|  |  |  | glutamate synthase  (NADPH) | Cys | 1ea0 | A | 1 | Asn:231:ND2 | Gly:232:N | – | N |
|  |  |  | glutamate synthase  (ferredoxin) | Cys | 1ofd | A | 1 | Asn:227:ND2 | Gly:228:N | – | N |
|  |  |  | glutamine-fructose-6-  phosphate transaminase  (isomerizing) | Cys | 1jxa | A | 1 | Asn:98:ND2 | Gly:99:N | – | N |
|  |  |  | glutaryl-7-amino-  cephalosporanic-acid  acylase | Ser | 3s8r | A | 170 | Val:239:N | Asn:413:ND2 | – | N |
|  |  |  | penicillin amidase  (peptidase S45 family) | Ser | 1pnl | B | 1 | Asn:241:N | Ala:69:N | – | N |
|  |  |  | proteasome endopeptidase  complex | Thr | 1ryp | I | 1 | Gly:47:N | Arg:19:N | – | P |
|  |  | 3.60.20.30 | N4-(beta-N-acetyl-  glucosaminyl)-L-  asparaginase | Thr | 1apy | B | 183 | Thr:234:OG1 | Gly:235:N | – | N |
|  |  | 3.60.60.10 | penicillin amidase  (peptidase C59 family) | Cys | 3pva | A | 1 | Tyr:82:N | Asn:175:ND2 | – | N |
|  |  | 3.60.70.12 | D-stereospecific  aminopeptidase  (peptidase S58 family) | Ser | 1b65 | A | 250 | Asn:218:N | Tyr:146:N | – | P |
|  |  | 3.60.90.10 | adenosylmethionine  decarboxylase  (prokaryotic) | Ser | 1vr7 | A | 63 | Ser:55:OG | – | – | P |
|  |  | 3.90.226.40 | 3-hydroxyisobutyryl-CoA  hydrolase | Glu | 3bpt | A | 169 | Gly:146:N | Gly:98:N | – | N |
| S_N_2 | N+1-type NE | 3.40.50.1000 | phosphoserine  phosphatase | Asp | 1l7n | A | 11 | Phe:12:N | Gly:100:N | – | N |
|  |  | 3.90.190.10 | protein-tyrosine-phosphatase  non-receptor class | Cys | 1ytw | A | 403 | Arg:404:N | – | – | N |
|  | N-type NE | 3.40.720.10 | alkaline phosphatase | Ser | 1alk | A | 102 | Ser:102:N | Arg:166:NH1 | Arg:166:NH2 | N |
|  | Other | 3.30.428.10 | bis(5'-adenosyl)-triphosphatase | His | 5fit | A | 96 | Gln:83:NE2 | – | – | N |
|  |  | 3.40.50.1240 | fructose-2,6-bisphosphate  2-phosphatase | His | 2bif | A | 256 | Arg:255:NE | Arg:255:NH2 | – | N |

* The type of nucleophilic attack reaction that the annotated nucleophilic residue of the enzyme performs.

**†** Classified into “N-type nucleophilic elbow (NE)”, “N+1-type nucleophilic elbow (NE)”, and “Other” (all non-nucleophilc elbow oxyanion holes).

‡ Homologous superfamilies, as defined in the CATH Protein Structure Classification database (*110*).

**§** Nucleophilic amino acid residue.

¶ Oxyanion hole donor identity, specified in the format of (amino acid):(residue number):(atom name). “-” indicate that the hydrogen bond donor is missing or cannot be identified.

\# “P” for proteases and “N” for non-protease enzymes.

**Table S22.** Serine and cysteine proteases curated by Buller and Townsend (*27*).

| **Oxyanion hole type†** | **Clan** | **Super-family (CATH)** | **Name** | **Nucl.^‡^** | **Ref PDB** | **Chain ID** | **Nucl. residue ID** | **Oyanion hole H-bond donor 1§** | **Oyanion hole H-bond donor 2§** |
| --- | --- | --- | --- | --- | --- | --- | --- | --- | --- |
| N+1-type nucl. elbow | PC | 3.40.50.880 | PH1704 | Cys | 1G2I | A | 100 | His:101:N | - |
|  | PC | 3.40.50.880 | aspartyl dipeptidase | Ser | 1FYE | A | 120 | Ala:121:N | Gly:88:N |
|  | SK | 3.90.226.10 | ClpP | Ser | 1TYF | A | 97 | Met:98:N | Gly:68:N |
|  | SS | 3.50.30.60 | LD-carboxypeptidase | Ser | 1ZRS | A | 115 | Asp:116:N | – |
| N-type nucl. elbow | CA | 3.90.70.10 | Papain | Cys | 1POP | A | 25 | Cys:25:N | Gln:19:NE2 |
|  | CD | 3.30.70.1470 | Caspase 8 | Cys | 1QTN | A | 360 | Cys:360:N | His:317:NE2 |
|  | CE | 1.10.418.20 | Ulp1 | Cys | 1EUV | A | 580 | Cys:580:N | Gln:574:NE2 |
|  | CF | 3.40.630.20 | Pyrrolidone carboxylate peptidase | Cys | 1IU8 | A | 139 | Cys:139:N | – |
|  | CL | 2.40.440.10 | LD-transpeptidase | Cys | 1ZAT | A | 442 | Cys:442:N | Gly:441:N |
|  | CM | 2.30.30.710 | HCV NS2 protease | Cys | 2HD0 | A | 184 | Cys:184:N | - |
|  | CN | 3.90.70.110 | Equine encephalitis alphavirus nsP2 | Cys | 2HWK | A | 477 | Cys:477:N | Val:476:N |
|  | CO | 3.40.50.1460 | Diamino endopeptidase | Cys | 2HBQ | A | 285 | Cys:285:N | Gly:238:N |
|  | CP | 3.90.1720.30 | deSUMOylase DeSI-1 | Cys | 2WP7 | A | 108 | Cys:108:N | – |
|  | PA | 1.10.1840.10 | SARS protease | Cys | 3SND | A | 145 | Cys:145:N | Gly:143:N |
|  | SF | 2.170.230.10 | Signal peptidase | Ser | 1B12 | A | 90 | Ser:90:N | – |
|  | SJ | 3.30.230.110 | VP4 protease | Ser | 4IZJ | A | 633 | Ser:633:N | – |
|  | ST | 1.20.1540.10 | Rhomboid-1 | Ser | 4QO0 | A | 201 | Ser:201:N | Asn:154:ND2 |
| Other | SH | 3.20.16.10 | CMV protease | Ser | 1NKM | A | 132 | Arg:165:N | – |

**†** Classified into “N-type nucleophilic elbow”, “N+1-type nucleophilic elbow”, and “Other” (all non-nucleophilc elbow oxyanion holes).

‡ Nucleophilic residue.

**§** Oxyanion hole donor identity, specified in the format of (amino acid):(residue number):(atom name). “-” indicate that the hydrogen bond donor is missing or cannot be identified.

**Table S23.** Number of enzyme nucleophiles found for each type of nucleophilic attack reactions from the combined dataset (including all enzymes in tables S3, 21 and 22).

| **Reaction mechanism** | **Electrophilic group** | **Leaving group** | **Number of**  **Nucleophilic elbows**  **(N- or N+1-type)** | **Number of**  **non-nucleophilic**  **elbow oxyanion holes** | **Total number** |
| --- | --- | --- | --- | --- | --- |
| Nucleophilic aromatic  substitution | Chlorobenzoate | Chloride ion | 0 | 1 | 1 |
| Nucleophilic addition on carbonyl | Aldehyde | Alkene (eliminated) | 1 | 0 | 1 |
|  | Aldehyde | Hydride | 4 | 1 | 5 |
|  | Amide (N-carbamoyl) | Ammonia | 2 | 0 | 2 |
|  | Amide (N-glycosylated amino acid) | Amine | 0 | 1 | 1 |
|  | Amide  (amino acid) | Ammonia | 8 | 5 | 13 |
|  | Amide (lactam) | Amine | 3 | 0 | 3 |
|  | Amide (other) | Amine | 2 | 3 | 5 |
|  | Amide (other) | Ammonia | 1 | 0 | 1 |
|  | Amide (peptide) | Amine | 52 | 7 | 59 |
|  | Carboxylate | Water | 1 | 0 | 1 |
|  | Ester | Alcohol | 16 | 2 | 18 |
|  | Ester (adenylated) | Phosphate | 0 | 1 | 1 |
|  | Thioester | Thiol | 9 | 1 | 10 |
|  | *combined* | | 99 | 21 | 120 |
| S_N_2 | Phosphoryl (phosphoic ester) | Alcohol | 3 | 1 | 4 |
|  | Phosphoryl (triphosphate) | Phosphate | 0 | 1 | 1 |
|  | *combined* | | 3 | 2 | 5 |
| *combined* | | | 102 | 24 | 126 |

**Table S24.** Pairwise RMSD values obtained from the local alignments of nucleophilic elbow atoms from different enzymes in their GSA-bound and acylenzyme states.

| **Oxyanion hole**  **type†** | **Nucl.**  **rotamer** | **Reaction**  **mechanism** | **Sub-**  **ensemble** | **Ref**  **PDB** | **Name** | *n* | **RMSD (Å)** | |
| --- | --- | --- | --- | --- | --- | --- | --- | --- |
|  |  |  |  |  |  |  | *mean* | *s.d.* |
| N+1-type NE  (7 atoms) | *trans* | Nucleophilic  addition  on carbonyl | Acylenzyme | 4BCB | ClpP | 18 | 0.19 | 0.05 |
|  |  |  |  |  | Dipeptidyl peptidase IV | 7 | 0.17 | 0.04 |
|  |  |  |  |  | Prolyl endopeptidase | 2 | 0.09 | 0.13 |
|  |  |  |  |  | *combined* | 27 | 0.18 | 0.06 |
|  |  |  | E•GSA | 5EIE | Dipeptidyl peptidase IV | 6 | 0.16 | 0.06 |
|  |  |  |  |  | LD-carboxypeptidase | 1 | 0.74 | – |
|  |  |  |  |  | Prolyl aminopeptidase | 3 | 0.20 | 0.08 |
|  |  |  |  |  | Prolyl endopeptidase | 2 | 0.03 | 0.04 |
|  |  |  |  |  | acetylcholinesterase | 12 | 0.29 | 0.10 |
|  |  |  |  |  | aspartyl dipeptidase | 1 | 0.17 | – |
|  |  |  |  |  | carboxypeptidase D | 1 | 0.15 | – |
|  |  |  |  |  | chloride peroxidase (cofactor free) | 2 | 0.17 | 0.00 |
|  |  |  |  |  | *combined* | 28 | 0.23 | 0.15 |
| N-type NE  (6 atoms) | *g+* | Nucleophilic  addition  on carbonyl | Acylenzyme | 1B12 | Rhomboid-1 | 1 | 0.25 | – |
|  |  |  |  |  | Signal peptidase | 2 | 0.27 | 0.01 |
|  |  |  |  |  | aldehyde dehydrogenase (NAD+) (class 2) | 1 | 0.41 | – |
|  |  |  |  |  | combined | 4 | 0.30 | 0.08 |
|  |  |  | E•GSA | 4YR1 | aldehyde dehydrogenase (NAD+) (class 2) | 2 | 0.68 | 0.12 |
|  |  |  |  |  | asparaginase | 1 | 0.32 | – |
|  |  |  |  |  | *combined* | 3 | 0.56 | 0.22 |
|  | *g–* | Nucleophilic  addition  on carbonyl | Acylenzyme | 2AGE  (Trypsin) | Caspase 8 | 5 | 0.31 | 0.04 |
|  |  |  |  |  | Chymotrypsin | 4 | 0.22 | 0.08 |
|  |  |  |  |  | D-Ala-D-Ala peptidase | 6 | 0.07 | 0.01 |
|  |  |  |  |  | Diamino endopeptidase | 8 | 0.32 | 0.03 |
|  |  |  |  |  | Elastase | 11 | 0.13 | 0.06 |
|  |  |  |  |  | LD-transpeptidase | 3 | 0.44 | 0.02 |
|  |  |  |  |  | SARS protease | 17 | 0.27 | 0.03 |
|  |  |  |  |  | Thrombin | 5 | 0.20 | 0.10 |
|  |  |  |  |  | Trypsin | 7 | 0.11 | 0.04 |
|  |  |  |  |  | beta-ketoacyl-[acyl carrier protein] synthase I | 4 | 0.27 | 0.01 |
|  |  |  |  |  | beta-lactamase (Class A) | 1 | 0.15 | – |
|  |  |  |  |  | beta-lactamase (Class C) | 6 | 0.13 | 0.03 |
|  |  |  |  |  | beta-lactamase (Class D) | 2 | 0.10 | 0.00 |
|  |  |  |  |  | caspase-1 | 8 | 0.32 | 0.03 |
|  |  |  |  |  | fatty acid amide hydrolase | 6 | 0.16 | 0.06 |
|  |  |  |  |  | glutamin-(asparagin-)ase | 4 | 0.16 | 0.01 |
|  |  |  |  |  | ubiquitinyl hydrolase 1 (peptidase C30 type) | 16 | 0.30 | 0.09 |
|  |  |  |  |  | *combined* | 113 | 0.23 | 0.10 |
|  |  |  | E•GSA | 3M7Q  (Trypsin) | Alpha-Lytic Protease | 2 | 0.15 | 0.05 |
|  |  |  |  |  | Chymotrypsin | 20 | 0.08 | 0.03 |
|  |  |  |  |  | D-Ala-D-Ala peptidase | 2 | 0.11 | 0.04 |
|  |  |  |  |  | Elastase | 6 | 0.21 | 0.19 |
|  |  |  |  |  | Proteinase K | 1 | 0.16 | – |
|  |  |  |  |  | SARS protease | 5 | 0.33 | 0.11 |
|  |  |  |  |  | Streptogrisin B | 26 | 0.11 | 0.06 |
|  |  |  |  |  | Subtilisin | 23 | 0.07 | 0.03 |
|  |  |  |  |  | Thrombin | 10 | 0.21 | 0.14 |
|  |  |  |  |  | Trypsin | 84 | 0.08 | 0.04 |
|  |  |  |  |  | asparaginase | 28 | 0.21 | 0.09 |
|  |  |  |  |  | beta-lactamase (Class A) | 4 | 0.40 | 0.26 |
|  |  |  |  |  | beta-lactamase (Class D) | 1 | 0.89 | – |
|  |  |  |  |  | glutamin-(asparagin-)ase | 8 | 0.17 | 0.02 |
|  |  |  |  |  | ubiquitinyl hydrolase 1 (peptidase C30 type) | 16 | 0.66 | 0.42 |
|  |  |  |  |  | *combined* | 236 | 0.16 | 0.20 |
|  |  | S_N_2 | E•GSA | 3M7Q  (Trypsin) | alkaline phosphatase | 9 | 0.22 | 0.18 |

**†** Classified into “N-type nucleophilic elbow (NE)” and “N+1-type nucleophilic elbow (NE)”.

**Table S25.** Nucleophilic elbow geometries in proteases and non-protease enzymes in their GSA-bound states.

| **Type**† | **Reaction**  **mechanism** | **Enzyme(s)** | **N**  **structures** | ***d*_attack_**‡ **(Å)** | | ***a*_attack_**‡ **(°)** | | ***ϕ*_attack_**‡ **(°)** | | **1° *ϕ*_HB_§ (°)** | | **2° *ϕ*_HB_§ (°)** | |
| --- | --- | --- | --- | --- | --- | --- | --- | --- | --- | --- | --- | --- | --- |
|  |  |  |  | *mode* | *s.d* | *mode* | *s.d* | *mode* | *s.d* | *mode* | *s.d* | *mode* | *s.d* |
| P | Nucleophilic addition  on carbonyl | *combined* | 217 | 2.69 | 0.41 | 93 | 9 | 84 | 12 | 74 | 14 | 78 | 19 |
| N | Nucleophilic addition  on carbonyl | 1-alkyl-2-acetylglycero-  phosphocholine esterase | 1 | 4.01 | – | 98 | – | 75 | – | 74 | – | 16 | – |
|  |  | acetylcholinesterase | 12 | 2.43 | 0.16 | 78 | 6 | 80 | 7 | 45 | 13 | 26 | 7 |
|  |  | aldehyde dehydrogenase  (NAD+) (class 2) | 2 | 2.82 | 0.30 | 73 | 2 | 45 | 30 | 62 | 11 | – | – |
|  |  | asparaginase | 29 | 2.81 | 0.17 | 89 | 6 | 86 | 17 | 85 | 13 | 85 | 14 |
|  |  | beta-lactamase  (Class A) | 4 | 3.21 | 0.23 | 50 | 25 | 21 | 33 | 57 | 17 | 15 | 19 |
|  |  | beta-lactamase  (Class D) | 1 | 3.27 | – | 50 | – | 13 | – | 65 | – | 26 | – |
|  |  | chloride peroxidase  (cofactor free) | 2 | 2.46 | 0.15 | 97 | 1 | 87 | 3 | 83 | 4 | 64 | 4 |
|  |  | glutamin-(asparagin-)ase | 8 | 2.78 | 0.08 | 91 | 4 | 86 | 2 | 86 | 4 | 78 | 3 |
|  |  | malonyl-CoA-acyl  carrier protein transacylase | 1 | 3.83 | – | 87 | – | 83 | – | 54 | – | 35 | – |
|  |  | *combined* | 60 | 2.77 | 0.31 | 88 | 11 | 84 | 21 | 84 | 20 | 83 | 29 |
|  | S_N_2 | alkaline phosphatase | 9 | 2.39 | 0.20 | 86 | 4 |  |  |  |  |  |  |
|  |  | protein-tyrosine-phosphatase  non-receptor class | 4 | 3.44 | 0.47 | 74 | 6 |  |  |  |  |  |  |
|  |  | *combined* | 13 | 2.49 | 0.45 | 85 | 6 |  |  |  |  |  |  |

† “P” for proteases and “N” for non-protease enzymes.

‡ Defined in Fig. 2A.

**§** *ϕ*_HB_ is defined in Fig. 5F. 1° refers to the hydrogen bond donor within the nucleophilic elbow, 2° refers to the other hydrogen bond donor(s) that is outside of the nucleophilic elbow (defined in Fig. 6A).

**Table S26.** Enzymes that perform nucleophilic attack and contain oxyanion holes yet do not have nucleophilic elbows and potential mechanistic constraints that prevent them from using nucleophilic elbows.

| **Mechanistic constraint** | **Name** |
| --- | --- |
| Nucleophilic amino acid does not allow N or N+1 arrangement | 3-hydroxyisobutyryl-CoA hydrolase |
|  | bis(5'-adenosyl)-triphosphatase |
|  | fructose-2,6-bisphosphate 2-phosphatase |
| Oxyanion multiple covalent bonds away from electrophilic atom | 4-chlorobenzoyl-CoA dehalogenase |
| Nucleophile attacks the backbone carbonyl of neighboring residue,  making its backbone amide inaccessible | adenosylmethionine decarboxylase (prokaryotic) |
|  | GyrA intein (Class 1 intein) |
| Nucleophile backbone amide used as general base | glutamine-fructose-6-phosphate transaminase (isomerizing) |
|  | glutamate synthase (ferredoxin) |
|  | proteasome endopeptidase complex |
|  | amidophosphoribosyltransferase |
|  | penicillin amidase (peptidase C59 family) |
|  | N4-(beta-N-acetylglucosaminyl)-L-asparaginase |
|  | glutaryl-7-aminocephalosporanic-acid acylase |
|  | asparagine synthase (glutamine-hydrolysing) |
|  | glutamate synthase (NADPH) |
|  | D-stereospecific aminopeptidase (peptidase S58 family) |
|  | HslU---HslV peptidase |
|  | GyrA intein (Class 1 intein) |
|  | penicillin amidase (peptidase S45 family) |
| Unidentified | CMV protease |
|  | UDP-glucose 6-dehydrogenase |
|  | E1 ubiquitin-activating enzyme |
|  | phospholipase A1 |
|  | Pectinesterase |
